# Single-cell genomics and regulatory networks for 388 human brains

**DOI:** 10.1101/2024.03.18.585576

**Authors:** Prashant S. Emani, Jason J. Liu, Declan Clarke, Matthew Jensen, Jonathan Warrell, Chirag Gupta, Ran Meng, Che Yu Lee, Siwei Xu, Cagatay Dursun, Shaoke Lou, Yuhang Chen, Zhiyuan Chu, Timur Galeev, Ahyeon Hwang, Yunyang Li, Pengyu Ni, Xiao Zhou, PsychENCODE Consortium, Trygve E. Bakken, Jaroslav Bendl, Lucy Bicks, Tanima Chatterjee, Lijun Cheng, Yuyan Cheng, Yi Dai, Ziheng Duan, Mary Flaherty, John F. Fullard, Michael Gancz, Diego Garrido-Martín, Sophia Gaynor-Gillett, Jennifer Grundman, Natalie Hawken, Ella Henry, Gabriel E. Hoffman, Ao Huang, Yunzhe Jiang, Ting Jin, Nikolas L. Jorstad, Riki Kawaguchi, Saniya Khullar, Jianyin Liu, Junhao Liu, Shuang Liu, Shaojie Ma, Michael Margolis, Samantha Mazariegos, Jill Moore, Jennifer R. Moran, Eric Nguyen, Nishigandha Phalke, Milos Pjanic, Henry Pratt, Diana Quintero, Ananya S. Rajagopalan, Tiernon R. Riesenmy, Nicole Shedd, Manman Shi, Megan Spector, Rosemarie Terwilliger, Kyle J. Travaglini, Brie Wamsley, Gaoyuan Wang, Yan Xia, Shaohua Xiao, Andrew C. Yang, Suchen Zheng, Michael J. Gandal, Donghoon Lee, Ed S. Lein, Panos Roussos, Nenad Sestan, Zhiping Weng, Kevin P. White, Hyejung Won, Matthew J. Girgenti, Jing Zhang, Daifeng Wang, Daniel Geschwind, Mark Gerstein

**Author notes:** Co-first authors, these authors contributed equally to this work (Prashant S. Emani, Jason J. Liu, Declan Clarke, Matthew Jensen, Jonathan Warrell, Chirag Gupta, Ran Meng, Che Yu Lee, Siwei Xu, Cagatay Dursun, Shaoke Lou).

## Abstract

Single-cell genomics is a powerful tool for studying heterogeneous tissues such as the brain. Yet, little is understood about how genetic variants influence cell-level gene expression. Addressing this, we uniformly processed single-nuclei, multi-omics datasets into a resource comprising >2.8M nuclei from the prefrontal cortex across 388 individuals. For 28 cell types, we assessed population-level variation in expression and chromatin across gene families and drug targets. We identified >550K cell-type-specific regulatory elements and >1.4M single-cell expression-quantitative-trait loci, which we used to build cell-type regulatory and cell-to-cell communication networks. These networks manifest cellular changes in aging and neuropsychiatric disorders. We further constructed an integrative model accurately imputing single-cell expression and simulating perturbations; the model prioritized ∼250 disease-risk genes and drug targets with associated cell types.

**Summary Figure:** 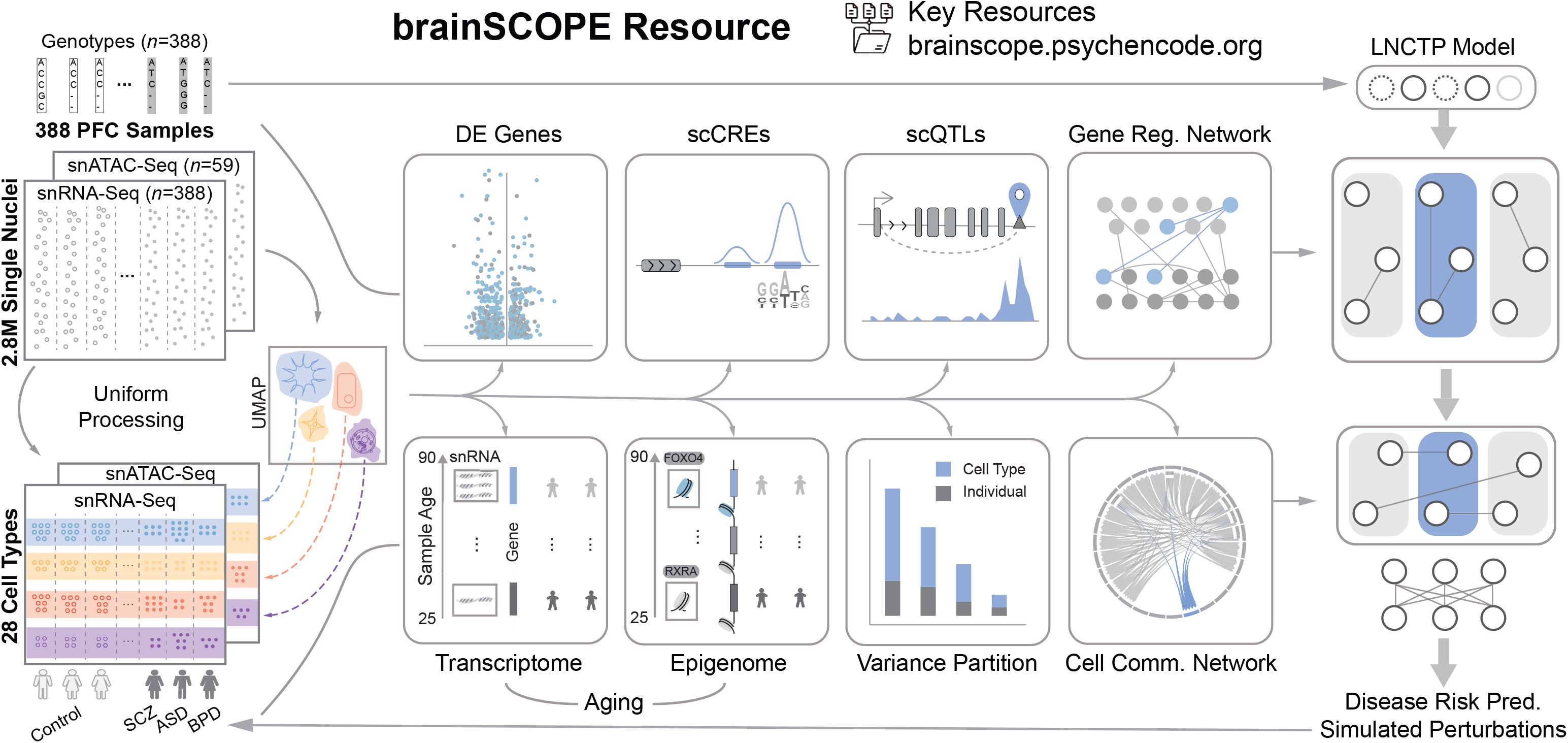

## Introduction

Genetic variants linked to neuropsychiatric disorders affect brain functions on multiple levels, from gene expression in individual cells to complex brain circuits between cells (*1–3*). At every level, they manifest themselves differently depending on the cell type in question. Previously, groups such as GTEx (Genotype-Tissue Expression), PsychENCODE, and ROSMAP (Religious Orders Study/Memory and Aging Project) assembled cohorts large enough to link variants to their effects on gene expression in bulk tissue, generating comprehensive eQTL (expression quantitative trait locus) catalogs for the brain (*4–6*). While useful, these tissue-level results do not reflect the specific cell types involved; moreover, they do not provide strong evidence that eQTLs act in cell-type-specific fashion (*7–10*).

Recently, dramatic technological advances have allowed the measurement of gene expression and chromatin accessibility at the single-cell level (*11–13*). The resulting datasets have shown that the brain has a particularly large number of distinct cell types; cell-type complexity, in fact, is one of the brain’s distinguishing features (*12*). Many brain cell types have been rigorously defined, particularly by the Brain Initiative Cell Census Network (BICCN) (*12*, *14*, *15*). Using these, we can potentially refine our understanding of how variants and gene regulation affect brain phenotypes, including neuropsychiatric disorders (*16*). However, up to now we have not had sufficiently large cohorts, with a wide enough range of brain phenotypes, to make statistically meaningful associations between variants, regulatory elements, and expression and to develop comprehensive models of brain gene regulation at the single-cell level.

To address this gap, the PsychENCODE consortium generated single-cell sequencing data from adult brains with multiple neuropsychiatric disorders in the human prefrontal cortex, using single-nucleus (sn) assays such as snRNA-seq, snATAC-seq, and snMultiome.

Leveraging these data and integrating them with other published studies (*12*, *17–19*), we created a uniformly processed single-cell resource at the population level. This resource, which we call brainSCOPE (brain Single-Cell Omics for PsychENCODE), comprises >2.8 M nuclei from 388 individual brains, including 333 newly generated samples and 55 from external sources (figs. S1-S2). It enables us to assess 28 distinct brain cell types that can be registered against previously identified canonical cell types (*12*, *19*). Using the resource, we identified an average of ∼85K cis-eQTLs per cell type and ∼550K cell-type-specific cis-regulatory elements, which were enriched for variants associated with brain-related disorders. Using our regulatory elements and eQTLs, we inferred cell-type-specific gene regulatory networks (which show great changes across cell types) as well as cell-to-cell communication networks. Moreover, we precisely quantified expression variation in the population, finding, for instance, that common neuro-related drug targets like *CNR1* demonstrate a high degree of cell-type variability and low inter-individual variability and that the transcriptomes of specific neurons are highly predictive of an individual’s age. Finally, we developed an integrative model to impute cell-type-specific functional genomic information for individuals from genotype data alone. Using this model, we prioritized many known and some additional disease genes, now with information about their specific cell type of action. We further associated this prioritization with potential drug targets and simulated the effects of perturbing the expression of particular genes.

All sequencing data, derived analysis files, and computer codes are available from the brainSCOPE resource portal brainscope.psychencode.org, figs. S3-S5; (*20*)); these include gene expression matrices from snRNA-seq data, regulatory regions from snATAC-seq data, variability metrics for all genes, single-cell QTL callsets, regulatory and cell-to-cell communication networks, and the integrative model and its prioritization outputs.

### Constructing a single-cell genomic resource for 388 individuals

We compiled and analyzed population-scale single-cell multiomics data from the human prefrontal cortex (PFC) for a cohort consisting of 388 adults. The individuals in our cohort are diverse in terms of biological sex, ancestry, and age, and include 182 healthy controls as well as individuals with schizophrenia, bipolar disorder, autism spectrum disorder (ASD), and Alzheimer’s disease (AD) (Fig. 1A; fig. S1; data S1-S2; table S1; (*20*)). We used various filters on the total cohort of 388 for different downstream analyses (fig. S2; data S3). In total, to build the resource, we uniformly processed 447 snRNA-seq, snATAC-seq, and snMultiome datasets from within PsychENCODE and external studies with >2.8M total nuclei (after QC and filtering from a raw number of nearly 4M; figs. S6, S7; table S2; (*20*)). Our processing required harmonizing datasets derived from different technologies and modalities; for instance, we generated uniform genotypes, including SVs, from combining whole-genome sequencing (WGS), SNP array, and snRNA-seq data (figs. S1, S8; (*20*)). We also generated custom datasets to bridge studies, in particular, snMultiome sequencing of controls (*20*).

**Figure 1.**
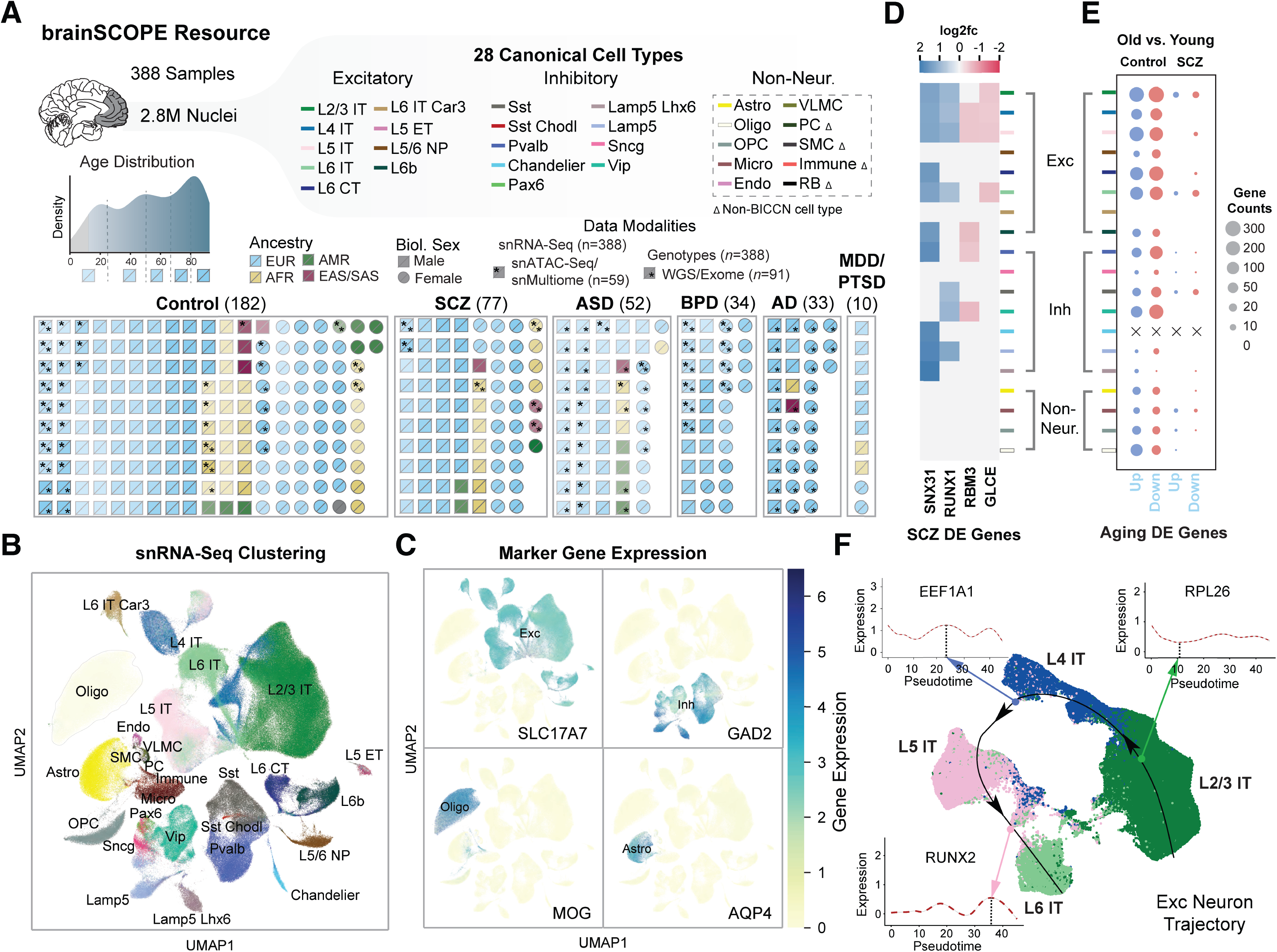
Constructing a single-cell genomic resource for 388 individuals. (A) Overview of the integrative single-cell analysis performed on 388 adult prefrontal cortex samples. (Top) Schematic for 28 cell types grouped by excitatory (Exc), inhibitory (Inh), and non-neuronal cell types (table S3); color labels for each subclass are used consistently throughout all figures (table S4). Dashed box indicates cell types defined with Ma-Sestan marker genes (*19*), with Δ indicating cell types unique to Ma-Sestan (Bottom) Schematic showing all samples labeled by disease, biological sex, ancestry, age, and available data modalities, including a distribution plot for sample ages (gray indicates pediatric samples excluded from most analyses). **(B)** UMAP plot for clustering of 28 harmonized cell types from snRNA-seq data derived from 72 samples in the SZBDMulti-seq cohort (using this study as an example of pan-cohort cell typing; see **fig. S10** for UMAPs of other studies). **(C)** UMAP plots highlighting expression of key marker genes in four broad cell types (excitatory: *SLC17A7*; inhibitory: *GAD2*; oligodendrocytes: *MOG*; and astrocytes: *AQP4*). **(D)** Differential expression (log2-fold change) of four schizophrenia-related genes across cell types in samples from individuals with schizophrenia (blue for upregulation, red for downregulation). **(E)** Numbers of DE genes upregulated (blue) and downregulated (red) in older (>70 years) control (left) and schizophrenia (right) individuals per cell type when compared with younger individuals in each group (<70 years). “X” indicates no DE genes were observed for a particular cell type. **(F)** UMAP plot showing predicted trajectory for excitatory IT neurons in adult control samples from the SZBDMulti-seq cohort. The predicted trajectory proceeds along the cortical layer dimension from L2/3 to L6 in the prefrontal cortex. Inset highlights log-normalized gene expression in cells along the pseudotime axis for three genes.

We developed a cell-type annotation scheme that harmonizes the BICCN reference atlas (*12*) and published analyses specifically focusing on the PFC (labeled “Ma-Sestan” here (*19*); Fig. 1B; figs. S9-S11; (*20*)). In particular, we leveraged the deep sampling of neurons from BICCN and of non-neuronal cells from Ma-Sestan. This resulted in a set of 28 cell subclasses, which we will hereafter refer to as “cell types,” most of which are robustly represented across all cohorts (tables S3-S4). For select downstream analyses that require increased power, we grouped excitatory and inhibitory neuron types into larger “excitatory” and “inhibitory” classes to yield seven major cell groupings. Overall, we assessed a total of 2,557,291 high-quality annotated nuclei from the snRNA-seq data (table S2). We validated our annotation scheme by assessing the expression of key marker genes (Fig. 1C).

Using these datasets, we first calculated cell-type fractions in each sample (figs. S12- S14; data S4; (*20*)). Fractions based on raw cell counts in snRNA-seq show great consistency with those inferred from bulk RNA-seq using deconvolution (fig. S12; data S5-S6). We further found that some cell types demonstrate cell-fraction differences in neuropsychiatric traits (fig. S13). For example, as previously suggested, the Sst cell fraction is different in individuals with bipolar compared to controls (*21*, *22*) (FDR<0.05, two-sided Welch’s t-test). To more broadly quantify differences relevant to population-wide traits, we computed lists of cell-type-specific differentially expressed (DE) genes for each disorder based on established approaches (*23*) (figs. S15-S18; data S7; (*20*)). Fig. 1D shows a representative plot for DE genes in schizophrenia, highlighting many previously known risk genes in a cell-type-specific context (*24*, *25*). We also found that individuals with schizophrenia differ from controls with respect to the number of aging DE genes, which may reflect the increased expression variability in schizophrenia patients (Fig. 1E; fig. S19).

Our snRNA-seq data also recapitulates the spatial relationships among cell types in the PFC. Fig. 1F shows a cell-trajectory analysis (*26*, *27*) across four subclasses of excitatory neurons in controls. We found smoothed patterns of gene-expression variation along the cortical-depth axis (specifically for L2/3, L4, L5, and L6 IT; figs. S20-S22; (*20*)). These findings expand on previous MERFISH-based results for 258 genes in the mouse motor cortex, now showing that cortical depth is related to gene expression variation for thousands of genes (*28*, *29*). Overall, we found 76 genes with significant variation (FDR<0.05, Wald test) across cortical layers, including several genes involved in neural development such as *SEMA6A*, *RUNX2*, *SOX6*, and *PROX1* (figs. S20-S22; list in table S5; data S8).

### Determining regulatory elements for cell types from snATAC-seq

In addition to snRNA-seq data, our resource contains 59 samples with snATAC-seq data, including 40 snMultiome datasets. After strict quality control, we extracted 273,502 deeply sequenced nuclei, allowing us to learn cell embeddings simultaneously from transcriptomic and epigenetic information (table S1; (*20*)). As a result, we recovered 28 distinct PFC cell types consistent with the snRNA-seq annotation and validated these with the chromatin accessibility of marker genes (Figs. 2A-2B; figs. S23-S24). Further, uniform snATAC-seq processing identifies a total of 562,098 open-chromatin regions across all datasets, representing a much larger number of regions than those identified in previous brain studies (Fig. 2C; (*20*)) (*2*, *30*). Following the ENCODE (Encyclopedia of DNA Elements) convention (*31*), we call these scCREs (single-cell candidate cis-Regulatory Elements). About half of these are cell-type-specific and located distal to genes (fig. S25). We validated the functionality of select scCREs using targeted STARR-seq (Fig. 2D; (*20*, *32*)).

**Figure 2.**
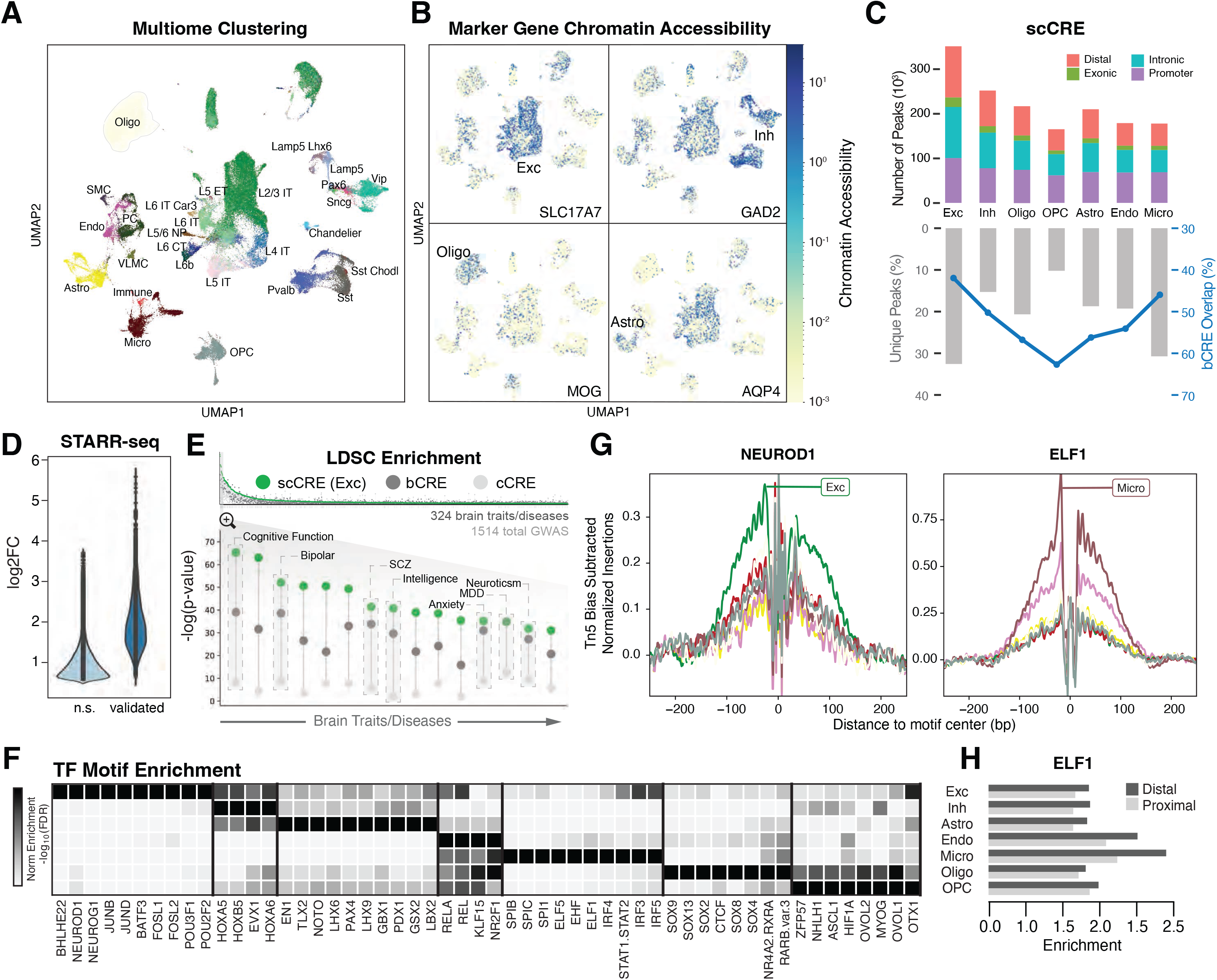
Determining regulatory elements for cell types from snATAC-seq. (A) UMAP plot for clustering of 28 harmonized cell types from snMultiome data derived from 40 individuals. **(B)** UMAP plots highlighting chromatin accessibility of key marker genes for four broad cell types (see Fig. 1C). **(C)** (Top) Counts of open chromatin regions from combined snATAC-seq and snMultiome peaks across cohorts by gene context (promoter, intronic, exonic, or distal). (Bottom) Percentage of unique ATAC peaks found in each cell type. Blue line indicates the percentage of ATAC peaks that overlap with b-cCREs derived from bulk data. **(D)** Change in enhancer activity among open chromatin regions using STARR-seq assays of predicted enhancers, comparing the log2-fold expression change of validated regions to non-validated regions (n.s.). **(E)** (Top) LDSC enrichment across GWAS summary statistics for UK BioBank traits and diseases, including brain-related traits (gray bars), cCREs (white circles), b-cCREs (gray circles), and snATAC-seq peaks in excitatory neurons (scCREs, green circles). (Bottom) LDSC enrichment (log-scaled p-values for LDSC analysis as explained in (*20*)) for select brain traits and disorders. Trait names are listed in table S6. **(F)** Enrichment (log-scaled FDR) of TF binding motifs among cell-type-specific snATAC-seq peaks. **(G)** Differential activity of *ELF1* in proximal and distal regions across cell types. **(H)** Cell-type-specific location of TF binding for *NEUROD1* (left) and *ELF1* (right) across cell types (colors defined in Fig. 1A), based on snATAC-seq footprinting analysis.

Using bulk data, we also developed a reference set of >400K open-chromatin regions, representing brain-tissue candidate cis-Regulatory Elements (b-cCREs; (*20*)). The b-cCREs were generated in a comparable fashion to ENCODE cCREs, which are not tissue-specific (*31*). As expected, they show strong overlap with scCREs (Fig. 2C).

To identify how our cell-type-specific regulatory elements relate to genetic associations, we performed a LDSC (linkage-disequilibrium score regression) analysis (*20*, *33*). In general, we found stronger LDSC enrichment for brain phenotypes in b-cCREs compared to cCREs (Fig. 2E; fig. S26; data S9-S10; table S6). Furthermore, we found additional enrichment when comparing cell-type-specific scCREs in excitatory neurons to b-cCREs, highlighting how snATAC-seq allows for better linkage between regulatory regions and brain phenotypes (Fig. 2E) (*34–37*).

Next, we explored transcription factor (TF) usage across major brain cell types (fig. S27; (*20*)). Fig. 2F shows that major brain cell types clearly use distinct TFs. For instance, *CUX1*, *NEUROG1*, and *PAX3* are mostly active in excitatory neurons, whereas *SPL1* and *SPI1* are specific to microglia. We further observed differences between proximal and distal regulation, for example, in *ELF1* (Fig. 2G; data S11). We were able to validate many TF activities with footprinting (*38*) (Fig. 2H; fig. S27).

### Measuring transcriptome and epigenome variation across the cohort at the single-cell level

Single-cell data across a large cohort offers a unique opportunity to study the sources of expression variation in the brain (Fig. 3A; figs. S28-S30; (*20*)) (*39*). We partitioned the variation in expression of each gene based on the relative contribution of individual and cell-type variability while correcting for covariates (data S12-S13). This allowed us to determine relative contributions to variability based on the function of each gene. For example, brain-specific genes, such as those associated with central nervous system (CNS) morphogenesis and neurotransmitter reuptake, demonstrate a high degree of cell-type variability and a lower inter-individual variation (Figs. 3B-3C; fig. S31; data S14). Conversely, genes associated with common molecular or cellular processes, tend to have lower cell-type variation and higher individual variation (for instance, carbohydrate homeostasis and ATP generation; Fig. 3B).

**Figure 3.**
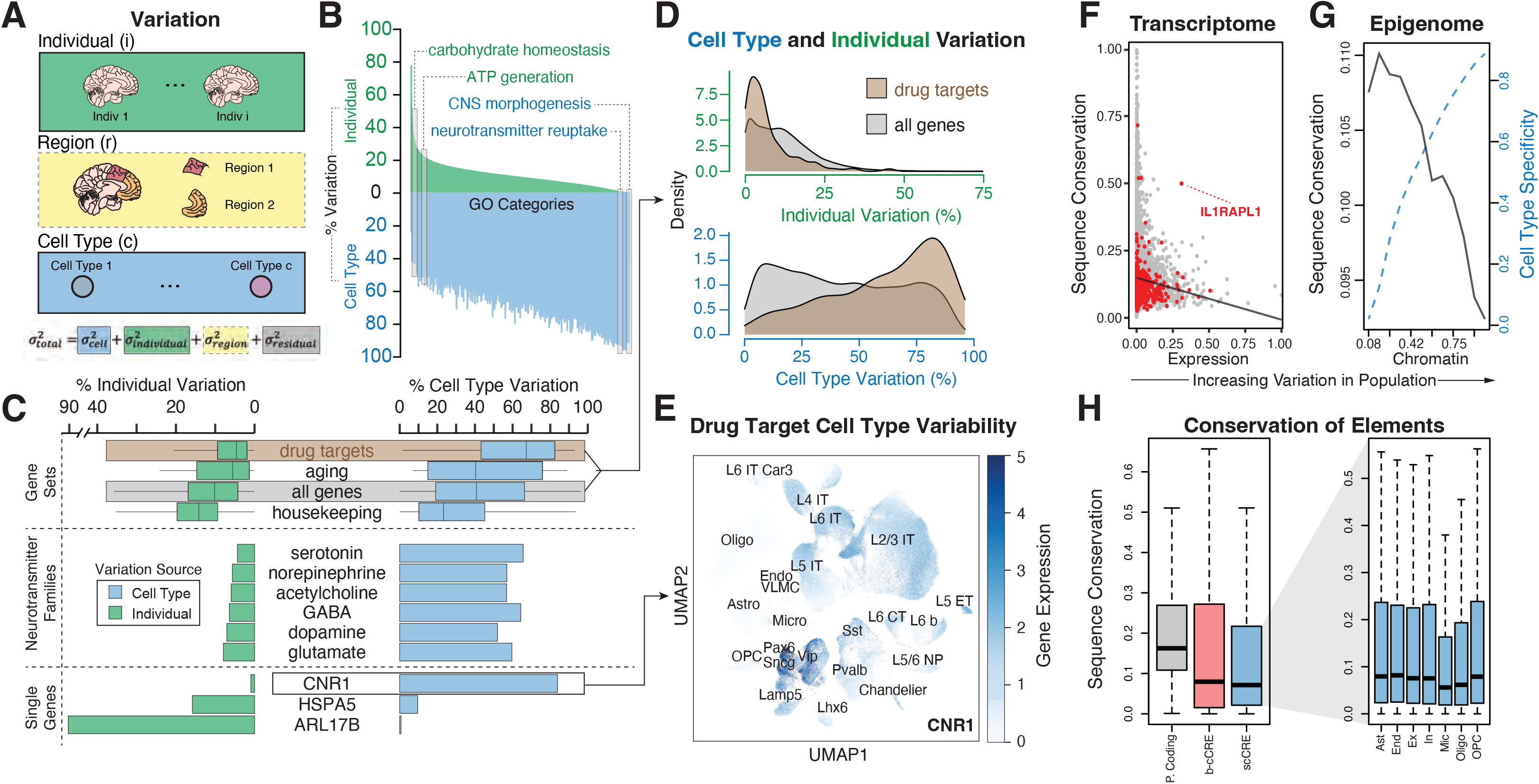
Measuring transcriptome and epigenome variation across the cohort at the single-cell level. (A) Schematic for the calculation of overall gene-level variance partition by integrating individual, brain region, and cell-type-specific variation. Variation analysis using different brain regions (denoted with a dashed gray box) was performed on a subset of individuals, shown in fig. S28. **(B)** Percent expression variation attributed to individuals (green) and cell types (blue) for GO categories, with select GO categories highlighted. **(C)** Percent inter-individual and cell type variation for specific genes and gene sets, including neurotransmitter families and drug targets. **(D)** Distribution of individual variation and cell type variation in drug target genes versus all genes. **(E)** UMAP plot of example drug target, *CNR1*, demonstrating cell-type-specific expression patterns that contribute to high cell type variability. We also assessed other genes such as serotonin receptor genes in fig. S31C. **(F)** Comparison of observed expression variation of individual genes with predicted conservation scores (phastCons). Red dots indicate outlier genes. Black line shows a trend of decreasing conservation as expression variation increases. **(G)** Increased cell-type specificity (dashed blue line) and decreased conservation (black line) observed as the population variability of scCREs increases. **(H)** (Left) Conservation of protein-coding regions, b-cCREs, and scCREs. (Right) Conservation of scCREs by cell type.

Furthermore, within families of CNS-specific genes, some neurotransmitter families manifest higher inter-individual variation compared to others (for example, glutamate vs serotonin, *p*- value=3.7x10^-6,^ one-sided t-test; Fig. 3C; fig. S31). We also identified a few outliers with very large inter-individual variation such as *ARL17B*, likely resulting from copy-number variation (*40*, *41*).

An additional application of quantifying expression variability is characterizing drug-target genes. In particular, we selected 280 common CNS-related drug-target genes and showed that, overall, they have high cell-type variability and low individual-level variability (Fig. 3C; fig. S32A) (*42*). That said, some of the 280 exhibit much higher inter-individual variation than others; *HSPA5* and *CNR1* provide a good illustration (Figs. 3C-3E; fig. S32B). Also, two adrenergic receptor family genes, *ADRA1A* and *ADRA1B*, demonstrate high cell-type variation but distinctly different cell-type expression patterns (fig. S32C).

Next, we found that genes with lower expression variability have higher sequence conservation (Fig. 3F; figs. S33-S34; (*20*)). However, some genes not following this trend serve as interesting exceptions (that is, highly conserved genes with high expression variance). The gene deviating most from the trend is *IL1RAPL1* (Fig. 3F; fig. S34B), an interleukin-1 receptor-family gene inhibiting neurotransmitter release (*43*); *IL1RAPL1* is highly expressed in the brain and has been implicated in intellectual disability and ASD (*44*).

We also leveraged our snATAC-seq profiles to deconvolve population-scale chromatin data (fig. S33; (*20*)). Similar to the transcriptome, open chromatin regions with higher sequence conservation have less variability in their chromatin openness (Fig. 3G; fig. S35). Furthermore, an increase in variability is concurrently observed with an increase in cell-type specificity. These patterns held when we jointly considered a gene and its linked upstream regulatory region; that is, a more variably expressed gene is associated with a more variable upstream chromatin region, and both of these are less conserved at the sequence level. (fig. S34A; (*20*)). Finally, we found that microglia scCREs exhibit the least sequence conservation, consistent with previous studies (Fig. 3H) (*19*, *45*, *46*).

### Determining cell-type-specific eQTLs from single-cell data

To evaluate cell-type expression variation in more detail, we used our processed snRNA-seq data to identify single-cell cis-eQTLs (hereafter referred to as “scQTLs”). We followed the same general procedure used by GTEx (*5*), including conservative filtering at the cell-type level when generating pseudobulk data (*20*). We used this set of scQTLs as our “core callset,” with the objective of facilitating consistent comparisons with those from existing datasets (such as GTEx and PsychENCODE bulk data) (data S15). Note the sparsity intrinsic to snRNA-seq data reduces power, particularly for rarer cell types (fig. S36; table S7; (*20*)) (*47*). To ameliorate the low power, we developed a Bayesian linear mixed-effects model to identify more scQTLs for rare cell types as an additional callset (Fig. 4A; figs. S36C, S37; table S8; (*20*)). We also generated further alternative callsets and a merge of results from all approaches (figs. S38- S39). These callsets include results based on linkage-disequilibrium pruning (table S9; (*20*)), regression across pseudo-time trajectories (below and (*20*)), and conditional analysis (giving rise to ∼1 signal per eGene, where an eGene is a gene involved in an eQTL; (*20*)). Finally, we identified a limited number of cell-type-specific isoform-usage QTLs (iso-QTLs), taking into account limitations in isoform identification from short-read snRNA-seq data (∼134K candidate iso-QTLs with 1389 associated “isoGenes”; figs. S40-S42; data S16; (*20*)).

**Figure 4.**
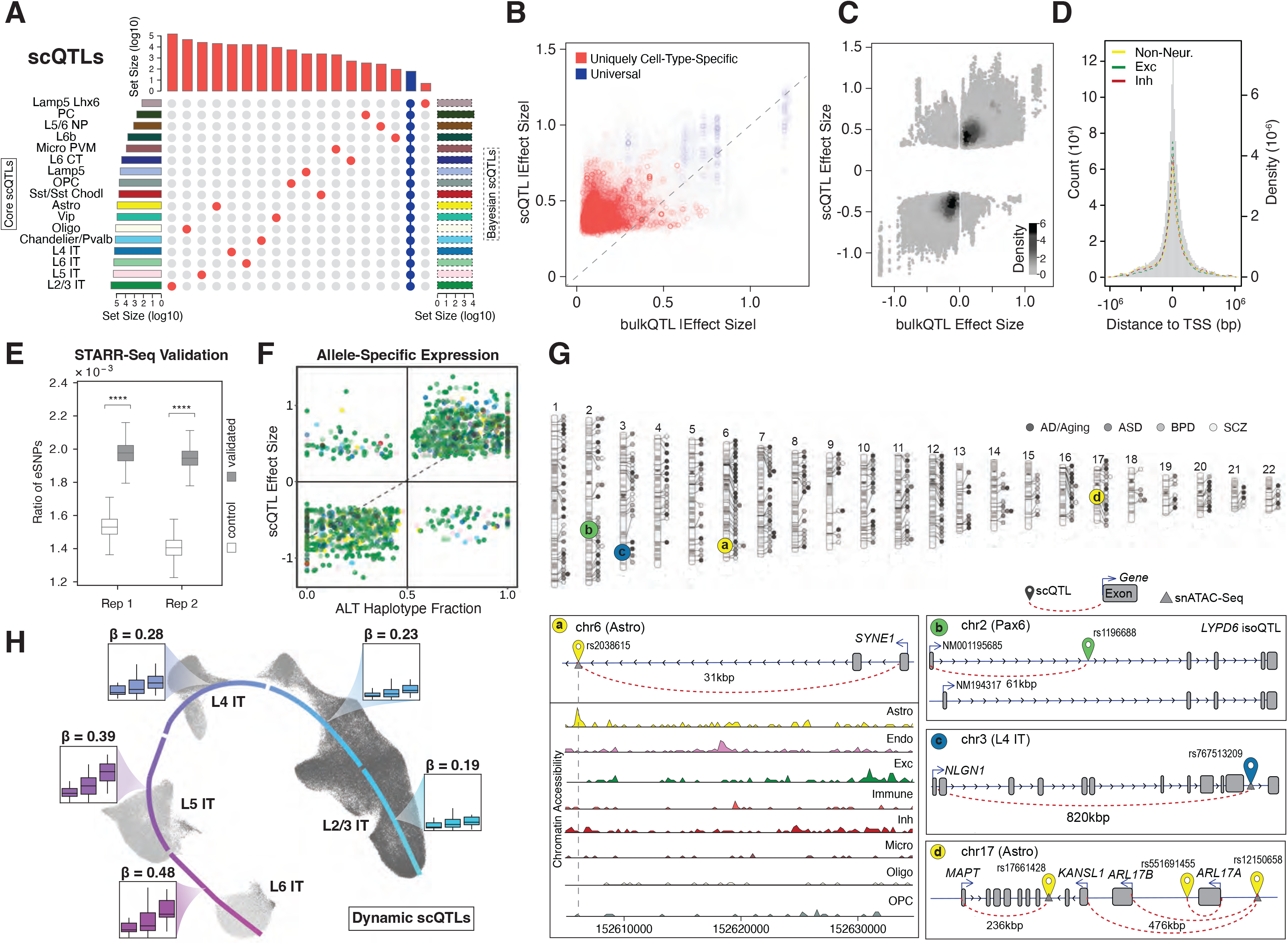
Determining cell-type-specific eQTLs from single-cell data. (A) Partial UpSet plot with identified scQTLs (from the core analysis) that are unique to individual cell types (red) or present across all cell types (blue). Left histogram summarizes the log-scaled total number of core scQTLs per cell type. Right histogram summarizes the log-scaled total number of Bayesian scQTLs per cell type. More complete plots are presented in figs. S36C and S42. **(B)** Scatter plots comparing absolute eQTL effect sizes between single-cell and bulk RNA datasets, highlighting QTLs shared across >14 cell types (blue) and unique to one cell type (red). **(C)** Density plot comparing eQTL effect sizes between single-cell and bulk RNA datasets. **(D)** Histogram with the distribution of scQTLs by distance from eGene transcription start site, with normalized distributions highlighted for the union of scQTLs across excitatory, inhibitory, and non-neuronal cell types. **(E)** Boxplot showing a significantly higher enrichment of eSNPs in active STARR-seq peaks compared to the control group (*p*<1.0x10^-4^, Mann-Whitney U Test). Two replicates are shown. **(F)** Scatter plot comparing scQTL effect sizes with allelic ratios of ASE eGenes, or the fraction of ASE gene reads originating from the haplotype with the scQTL alternative allele. ASE genes were identified in 21 MultiomeBrain cohort samples; cell types are represented with the color scheme used in **(A)**. **(G)** (Top) Chromosome ideogram for the location of eGenes in all cell types related to four brain disorders. (Bottom) Schematics for specific instances of scQTLs for disease-related eGenes. Left schematic shows astrocyte- specific eSNP for *SYNE1* along with chromatin accessibility (snATAC-seq) tracks for eight cell types. Top-right schematic shows the isoQTL for *LYPD6* in Pax6 inhibitory neurons, leading to altered expression of isoforms with different start codons. Middle and bottom-right schematics show SNP-gene pairs for scQTLs associated with *NLGN1* in L4 IT neurons and *MAPT* in astrocytes, respectively. **(H)** UMAP plot for predicted trajectory of excitatory neurons in samples from the SZBDMulti-seq cohort. Box plots highlight the expression of *EFCAB13*, stratified by eSNP genotype in each sample, for cell types in each cortical layer; effect size (β) values for the eSNP increase over pseudotime. Additional information is shown in fig. S53.

Overall, we identified an average of ∼85K scQTLs and ∼690 eGenes per cell type in our core set, resulting in ∼1.4M scQTLs when totaled over cell types (Fig. 4A; fig. S36A, S43; table S7; (*20*)). Many of the scQTLs are uniquely cell-type-specific (i.e. not in any other cell type), but ∼47% appear in more than one cell type (Fig. 4A; fig. S36). About 30% of the scQTLs overlap with bulk cis-eQTLs (*4*). Among these “overlappers,” the direction of effect is consistent (Fig. 4B), but the magnitude of the scQTL effect size is greater than that of the matched bulk eQTL (Fig. 4C; fig. S44; table S10). We posit a “dilution effect” as an explanation, wherein scQTL effect sizes may be diluted in bulk data when they occur only in a relatively small number of cell types. This line of reasoning is supported by comparing scQTLs appearing in a few cell types to those observed in many (Fig. 4B; fig. S36A). Overall, we found cell-type-specific QTLs were likely difficult to detect in bulk measurements, which is borne out by the fact that more than two-thirds of our scQTLs are not found in bulk despite much larger sample sizes available in bulk.

Our scQTLs are strongly enriched in narrow regions around the transcription start sites (Fig. 4D; figs. S45-S46). We validated some of our core scQTLs by comparing them with functional elements identified by STARR-seq, mut-STARR-seq, and massively parallel reporter assays (MPRA) (Fig. 4E; figs. S47-S48; (*20*, *32*)). As further validation, we were able to identify allele-specific expression (ASE) at the single-cell level in samples with WGS-based phased variants (Fig. 4F; fig. S49; (*20*)). Determination of single-cell ASE is particularly challenging due the sparsity of the data (*48–52*). Here, we compared the magnitude of the ASE effect at an SNV with the corresponding effect size of the scQTL involving the same SNV, finding significant correlation as expected (Fig. 4F; fig. S49; p< 2.0x10^-16^, Fisher’s exact test).

Overall, we identified 330 scQTLs for eGenes related to brain disorders (Fig. 4G; figs. S50-S51; data S17). For example, we found scQTLs for *SYNE1*, a candidate autism and schizophrenia gene (*53*, *54*), and *NLGN1*, a candidate gene for multiple brain disorders encoding a ligand for neurexin signaling (*55*). We also found multiple scQTLs within the complex 17q21.31 locus related to brain disorders, including an astrocyte-specific scQTL for the Tau protein gene *MAPT* and a multi-cell type scQTL for the neurodegenerative-disorder risk gene *KANSL1* (Fig. 4G) (*40*). We further highlight an iso-QTL for *LYPD6*, which inhibits acetylcholine-receptor activity in Pax6-type inhibitory neurons (*56*) (Fig. 4G).

Finally, we developed a Poisson-regression model that incorporates a continuous trajectory and a pseudotime-genotype interaction term to further expand our scQTLs, allowing for the calculation of “dynamic scQTLs” that exhibit a changing effect size along the pseudotime trajectory (figs. S52-S53; data S18-S19; (*20*)) (*57*). In particular, for 1692 of the 6255 unique eGenes in four types of excitatory neurons, we found a corresponding dynamic scQTL (with a non-zero interaction term); Fig. 4H and fig. S52 show examples. Moreover, many of these dynamic scQTLs imply widespread QTL effects in cell types where we do not discover a scQTL with our core approach (fig. S52).

### Building a gene regulatory network for each cell type

By integrating multiple data modalities, including scQTLs, snATAC-seq, TF-binding sites, and gene co-expression, we constructed gene-regulatory networks (GRNs) for PFC cell types (Fig. 5A; figs. S54-S58; data S20; (*20*)). In particular, we linked TFs to potential target genes based on their co-expression relationships from snRNA-seq data (*58*, *59*), and mapped scQTLs to connect promoters and enhancers (data S21). We make these networks available in a variety of easy-to-use formats (*20*). For instance, we applied a network-diffusion method that provides the key regulators of a given target gene -- specifically, the aggregate regulatory score of each TF for that target (figs. S59-S60).

**Figure 5.**
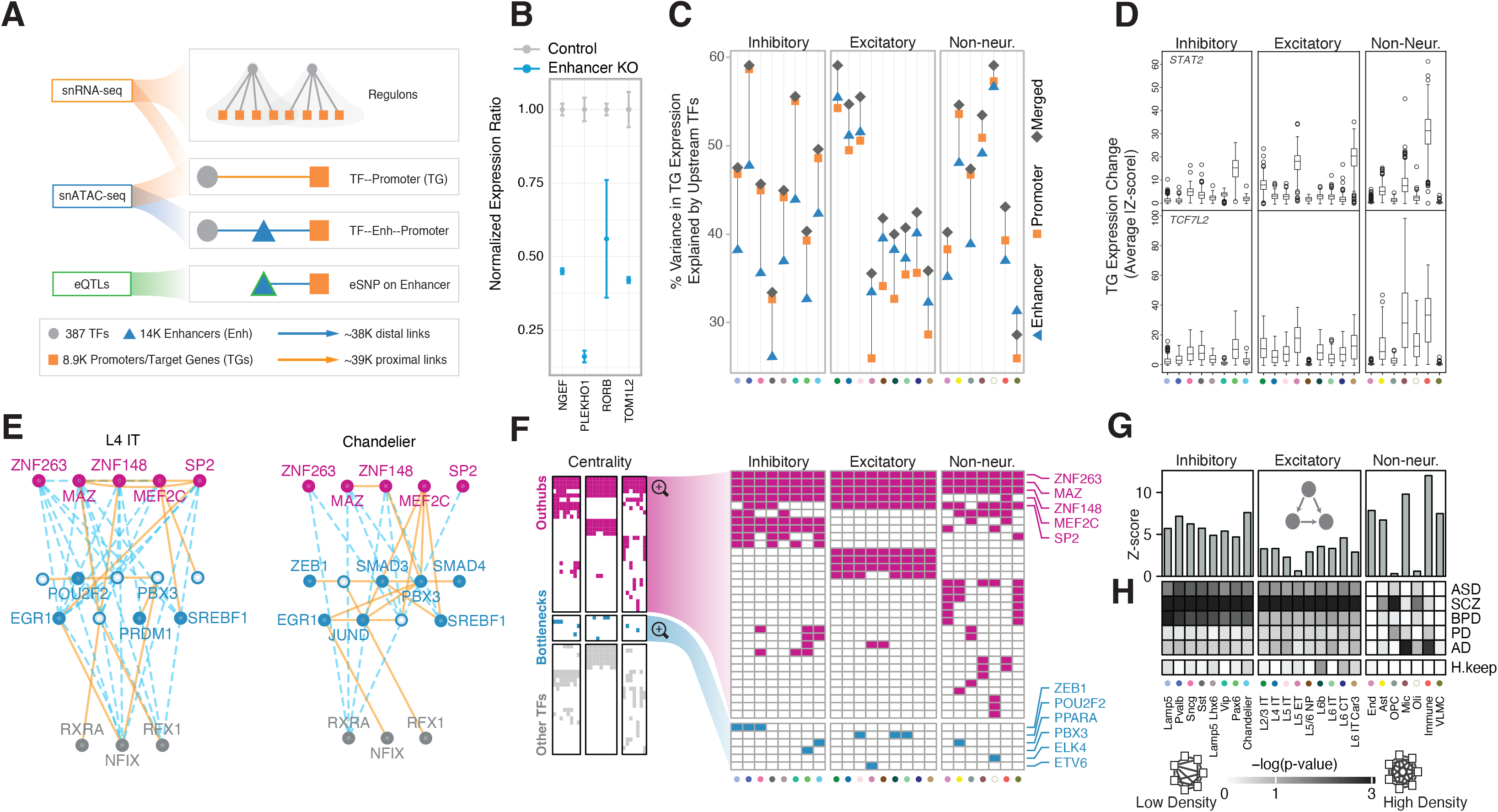
Building a gene regulatory network for each cell type. (A) Schematic for the construction of cell-type-specific GRNs based on snRNA-seq, snATAC-seq, and scQTL datasets. **(B)** Change in expression of four genes after CRISPR-mediated knockout of enhancers identified in cell-type-specific GRNs (blue bars) compared with control samples (gray bars). **(C)** Percent variance in target gene expression explained by the networks. Orange squares, blue triangles, and gray diamonds indicate variance explained by promoter, enhancer, and merged GRNs, respectively. **(D)** Changes in expression (average Z-score) of target genes in cell-type-specific regulons among samples with loss-of-function variants that disrupt the TFs *TCF7L2* and *STAT2*. **(E)** Network graphs depicting a subset of the excitatory (L4 IT) and inhibitory (Chandelier) GRNs that show differential usage of enhancers and promoters. Nodes (TFs) are colored in pink, blue, or gray to represent out-hubs, bottlenecks, and in-hubs, respectively. Nodes without blue fill represent TFs that are absent as bottlenecks in that cell type. Solid orange lines indicate proximal links; distal links are indicated by dashed blue lines. **(F)** Panel representing the full set of TFs (y-axis) that act as hubs or bottlenecks in different cell types (x-axis). Cells are colored if the TF is found to be a pure hub (magenta) or bottleneck (cyan) in the corresponding cell type. (Note hubs here are out-hubs.) The right panel zooms in to highlight hubs (top) and bottlenecks (bottom). **(G)** Motif enrichment analysis bar plots showing a stronger enrichment of transcriptional feed-forward loops (illustrated inset) in inhibitory neurons (left) and most non-neuronal cell types (right) compared with excitatory neurons (center). **(H)** Co-regulatory network changes of disease gene sets across cell types. The white-to-black gradient shows low to high probability (log p-value obtained by random sampling, N=10,000)) of a disease gene set or housekeeping genes (H.keep, y-axis) forming a dense subnetwork in the corresponding cell type (x-axis) (*20*). Cell types on the x-axis in panels C, D, and F are colored uniquely according to names in panel G.

We experimentally validated a subset of these linkages using CRISPR knockouts (Fig. 5B; fig. S61; data S22; (*20*)). Overall, we found that TF expression in the GRNs explain an average of 52% of the variation in expression of target genes, with merged networks explaining more variance than just the promoter or enhancer connections (Fig. 5C; fig. S62). Additionally, mapping loss-of-function (LOF) mutations in individuals to select TFs (“natural knock-outs”) provided further validation by showing the expected change in expression of their target genes in a cell-type-specific manner (fig. S63; (*20*)). Overall, 77% of TFs with LOF variants, including *TCF7L2* and *STAT2,* lead to the expected expression alteration within their cell-type-specific regulons (Fig. 5D; fig. S63).

Our analyses of GRNs uncovered complex network rewiring across the cell types (Fig. 5E; figs. S64-S66; data S23; (*20*)). In particular, the most highly connected TFs (“hubs”) are largely shared across cell types, suggesting their involvement in common machinery used by all brain cells (Fig. 5F). In contrast, bottlenecks (key connector TFs) have much more cell-type-specific activity (Fig. 5F; fig. S67A). Furthermore, the targets of bottleneck TFs are enriched for cell-type-specific functions, such as myelination and axon ensheathment for oligodendrocytes (*60*) (fig. S67B; data S24). Additionally, cell-type-specific GRNs greatly differ in the usage of network motifs, such as feed-forward loops (Fig. 5G). These particular motifs, which are thought of as a noise-filtering mechanism (*61*), are notably enriched in certain non-neuronal cell types.

Finally, disease genes for a particular disorder tend to be co-regulated in a cell-type-specific manner (Fig 5H; figs. S65,S68; (*20*)). For instance, gene sets related to schizophrenia form relatively dense subnetworks in neurons, whereas the AD subnetwork is actively co-regulated just in microglia and immune cells (fig. S69) (*37*, *62*, *63*).

### Constructing a cell-to-cell communication network

To further understand cellular signaling and regulation, we leveraged publicly available ligand-receptor pairs (*64*) in combination with our snRNA-seq data to construct a cell-to-cell communication network (Fig. 6A; table S11-S12, fig. S70; (*20*)). As expected, we observed three broad ligand-receptor usage patterns among excitatory, inhibitory, and glial cell types, indicating that these cell types use distinct signaling pathways in their communication. For instance, in both incoming and outgoing communication, we observed that all nine glial cell types are grouped together based on their ligand-receptor interactions, with growth-factor genes as some of the top contributing ligand-receptor pairs (*65–67*) (Fig. 6B).

**Figure 6.**
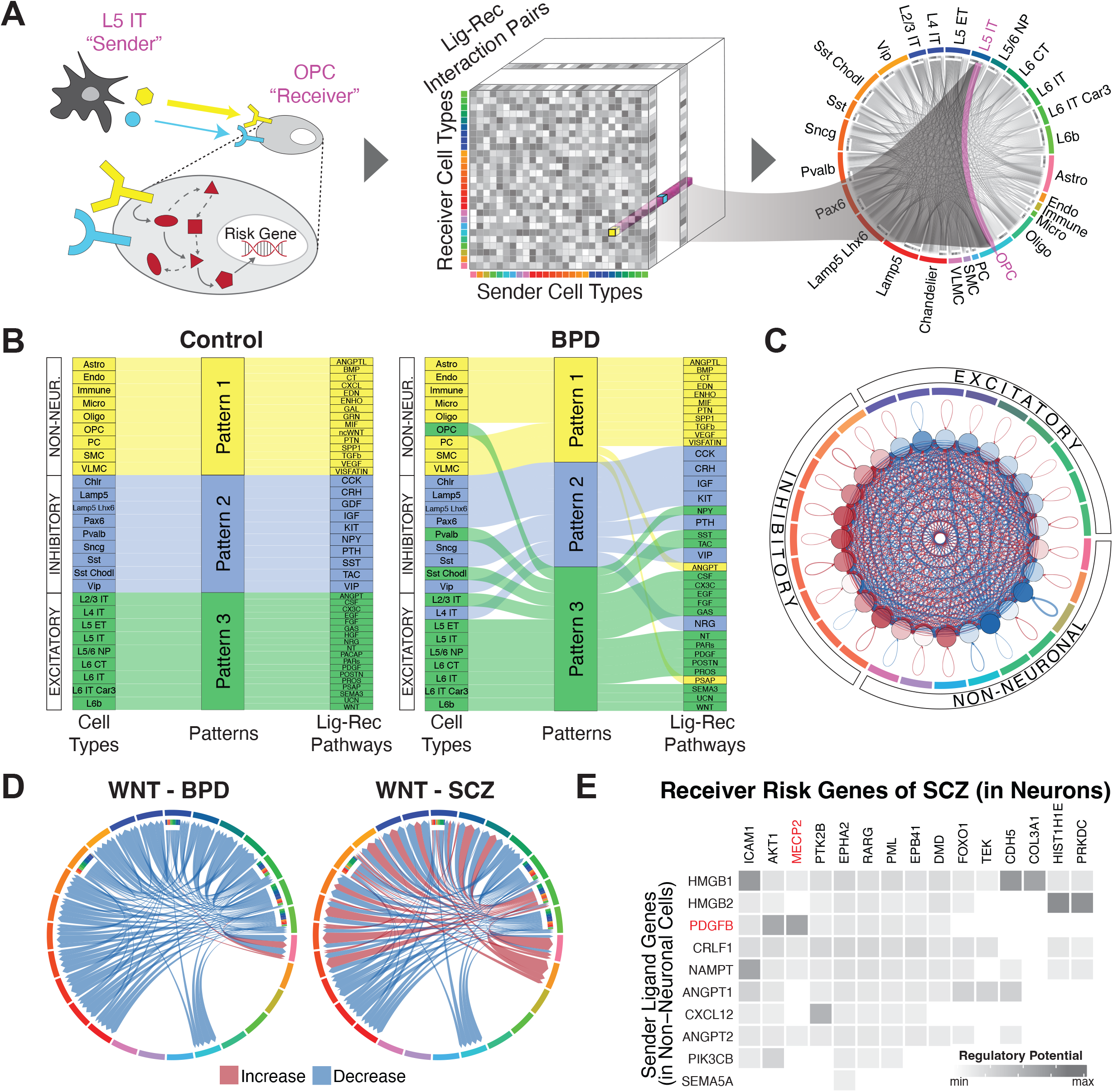
Constructing a cell-to-cell communication network. (A) Schematic for the construction of cell-to-cell communication networks, based on a matrix of co-expressed ligand- receptor gene pairs in signaling pathways between sender and receiver cell types. Circos plot on the right shows the strength of all identified cell-to-cell interactions, highlighting L5 IT to OPC cell types as an example. Note that this model does not consider the synaptic connectivity between neurons. **(B)** Sankey plots for differential clustering of incoming interactions in receiver cells across cell types and ligand-receptor signaling pathways for control (left) and bipolar disorder (right) samples. For example, inhibitory Sst and Sst Chodl cell types were assigned to Pattern 2 in controls, along with the SST-SSTR signaling pathway. This makes sense, as these cell types are predominantly characterized by the *SST* gene. However, in BPD samples, Sst Chodl cells switched from Pattern 2 to 3, along with the SST-SSTR signaling pathway. **(C)** Circos plot showing differential strength of all cell-to-cell interactions between individuals with schizophrenia and control individuals. Red edges indicate increased interaction strength in schizophrenia samples, while blue edges indicate weaker interaction strength. **(D)** Circos plots showing changes in cell-to-cell interaction strengths for ligand-receptor genes in the Wnt signaling pathway between individuals with bipolar disorder (left) and schizophrenia (right) compared with control individuals. **(E)** Predicted likelihoods that ligand genes in non-neuronal cells (y-axis) regulate schizophrenia-associated risk genes (x-axis) in neuronal cell types, with the neurological risk gene *MECP2* highlighted in red.

We next explored how cell-cell communication patterns are altered in individuals with neuropsychiatric disorders, finding that they are greatly changed for schizophrenia and bipolar disorder (Fig. 6B; figs. S71A-B, S72; data S25-S26). In fact, notable inter-mixings occur among the three broad patterns of ligand-receptor usage. For instance, in bipolar disorder, the excitatory pattern (inferred from controls) now also contains OPCs and some inhibitory neurons (Pvalb and Sst Chodl). In individuals with schizophrenia (compared to controls), we also found that excitatory neurons received less incoming signaling, while inhibitory neurons received more (Fig. 6C).

To further highlight network perturbations in disease, we assessed signaling-pathway changes for bipolar disorder and schizophrenia (Fig. 6D). In bipolar, we observed downregulation of the Wnt pathway, consistent with previous findings (Fig. 6D) (*68–71*).

Mechanistically, this downregulation could result in the overactivity of the lithium-targeted GSK3β enzyme in neurons (*72*, *73*). In schizophrenia, the Wnt pathway is downregulated as expected, but we also found increased sender communication strength for L6 IT Car3 neurons, different from bipolar (*74*). We further found downregulation of PTN pathway interactions from glial cells to neurons, consistent with previous studies (*75–77*), and a decrease in signaling to glial cells involving various growth factors (fibroblast, epidermal and insulin) (figs. S71C-E).

These findings support the “glial cell hypothesis,” which posits that deleterious effects on glial cells cascade to neurons (*78*).

Lastly, we extended our extracellular cell-to-cell communication analysis by considering related disruptions to intracellular signaling pathways (Fig. 6E; fig. S73; (*20*)) (*79*). By utilizing disease-risk genes and setting support cells (non-neurons) as the senders and neurons as the receivers, we identified ligand-receptor links connecting known risk genes to potential upstream effectors. For instance, we linked *FOXP1* and its ligand *EBI3* in bipolar disorder and *MECP2* and its ligand *PDGFB* in schizophrenia (*80*, *81*).

### Assessing cell-type-specific transcriptomic and epigenetic changes in aging

We used our population-scale single-cell data to systematically highlight transcriptomic and epigenetic changes due to aging. First, we assessed cell-fraction changes based on deconvolution of bulk data using our single-cell profiles and found that Chandelier and OPC cell types decrease with age, as in previous reports (FDR<0.05, two-sided t-test; Fig. 7A, data S27) (*82*, *83*). This result is consistent with findings from raw cell counts in the single-cell data (FDR<0.05, two-sided t-test; Fig. 7A; data S27; (*20*)). Next, we identified a list of aging DE genes across cell types (Fig. 7B; fig. S74; data S28; (*20*)). This list shows, for instance, that *HSPB1*, which encodes a heat-shock protein and has been previously implicated in longevity, is upregulated in multiple cell types in older individuals (*84*, *85*).

**Figure 7.**
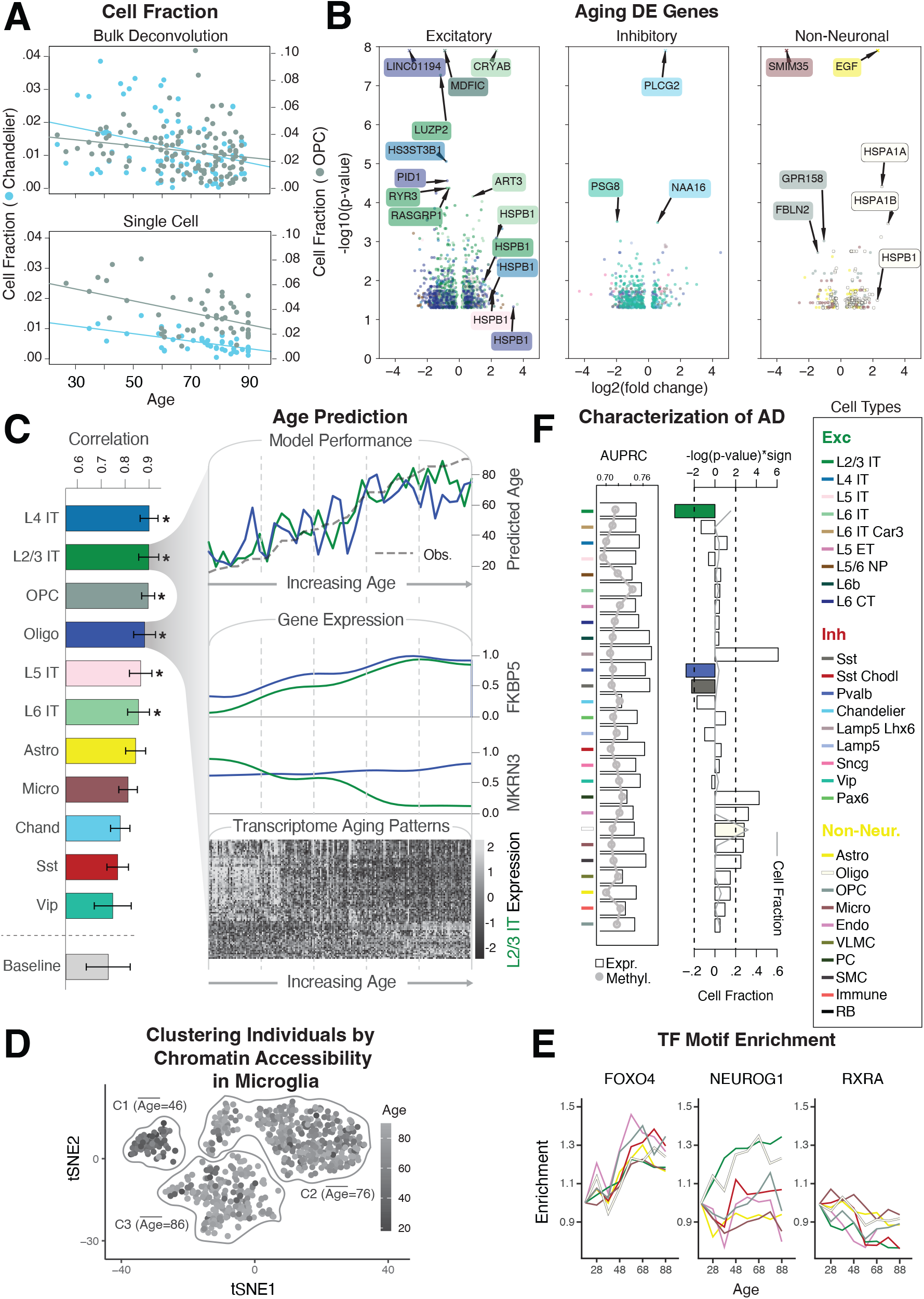
Assessing cell-type-specific transcriptomic and epigenetic changes in aging. (A) Normalized changes in the fraction of OPC (gray) and Chandelier cells (blue) by age, based on bulk RNA-seq deconvolution (top) and single-cell annotation (bottom), with best-fitted lines. **(B)** Log2-fold changes and p-values from DESeq2 (*20*) for differentially expressed genes in older vs. younger individuals (±70 years) among excitatory, inhibitory, and non-neuronal cell types. Values with -log(p)>8 are shown as crosses. **(C)** (Left) Pearson correlation values between model prediction of age and observed age for each cell type and baseline model (covariates). (Top right) Predicted and observed age for oligodendrocytes and L2/3 IT neurons along the age spectrum. (Middle right) Transcriptomic profiles along the age spectrum of two key genes (*MKRN3* and *FKBP5*) related to aging. (Bottom right) Genes demonstrate an increase (light gray) or decrease (dark gray) in expression along the age spectrum. **(D)** tSNE plot of chromatin peaks showing how chromatin patterns in microglia stratify younger and older individuals into three distinct clusters. **(E)** Examples of TF binding motifs that display distinct enrichment patterns across cell types and age. **(F)** (Left) Predictive accuracy (AUPRC) of cell-type-specific expression (bars) and methylation signatures (gray line) towards AD status. (Right) Enrichment of cell fraction changes among individuals with AD. L2/3 IT, Pvalb, and Sst (colored bars) are significantly associated with a decreased cell fraction in AD (log-p value, t-test). Gray line shows the overall median cell fraction of each cell type in AD individuals.

To further explore the relationship between the transcriptome and aging, we constructed a model to predict an individual’s age from their single-cell expression data (Fig. 7C; figs. S75A- B; (*20*)). The model shows that the transcriptomes of six cell types (L2/3 IT, L4 IT, L5 IT, L6 IT, Oligodendrocytes, and OPC) have strong predictive value (Fig. 7C; fig. S75C). It also shows that many individual genes contribute to the model, highlighting broad transcriptome changes in aging. From these, we selected two particularly predictive genes previously associated with aging, *FKBP5* and *MKRN3*, and observed a clear correlation between their expression and aging (Fig. 7C; fig. S76) (*86–88*).

We also investigated the effects of age on the epigenome using our scCREs to deconvolve bulk chromatin accessibility for 628 individuals into those for specific cell types (Fig. 7D; fig. S77). The resulting scCRE activity patterns in certain cell types, particularly microglia, cluster individuals into distinct age groups (Fig. 7D; fig. S77; (*20*)). We further expanded our analysis to highlight how patterns of enriched TF motifs in active scCREs change with age in a cell-type-specific fashion (Fig. 7E; fig. S78; (*20*)). Some TFs demonstrate consistent patterns across cell types (*FOXO4* and *RXRA)*, while others exhibit more cell-type-specific patterns (*NEUROG1*).

Finally, we extended our analysis to identify cell-type-specific changes in neurodegenerative disease. We obtain cell-type fractions by using our single-cell expression profiles to deconvolve 638 bulk RNA-seq samples, containing AD cases and controls (fig. S79A; (*20*)) (*89*). Certain glial fractions show a significant increase in AD (p<0.005, t-test), while several neuronal fractions decrease, especially Sst, Pvalb, and L2/3 IT, in line with previous studies (*90*) (Fig. 7F). We compared this result with that from directly comparing cell-type-specific gene-expression and methylation signatures to determine case-control status (*91*), finding that the fractions and signatures capture independent information (fig. S79B; data S29; (*20*)).

### Imputing gene expression and prioritizing disease genes across cell types with an integrative model

We incorporated many of the preceding single-cell datasets and derived networks into an integrative framework to model and interpret the connections between genotype and phenotype. We term our modeling framework a Linear Network of Cell Type Phenotypes (LNCTP; Fig. 8A; (*20*)). This framework serves four tasks: (1) to impute cell-type-specific and bulk tissue gene expression from genotype; (2) to predict the risk of disorders based on input genotypes; (3) to highlight genes and pathways contributing to particular phenotypes in their specific cell type of action; and (4) to simulate perturbations of select genes and quantify their impact on overall gene expression or trait propensity. The LNCTP has several visible layers associated with components of the resource described above, including genotypes at scQTL and bulk eQTL sites, cell-type-specific and bulk tissue-based GRNs, cell-type fractions, cell-to-cell communication networks, gene co-expression modules, and sample covariates (*20*).

**Figure 8.**
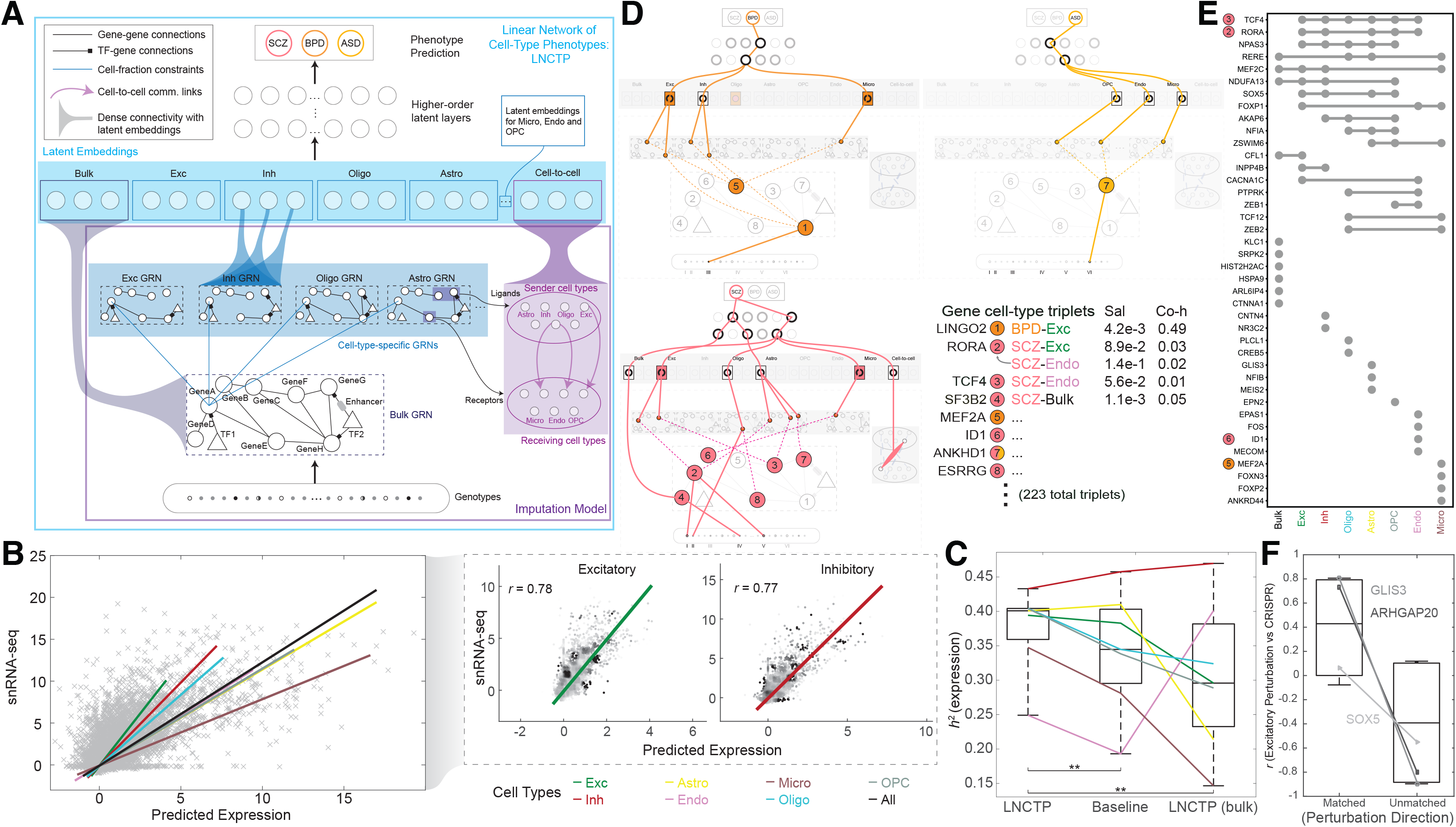
Imputing gene expression and prioritizing disease genes across cell types with an integrative model. (A) LNCTP schematic. Bulk and cell-type gene expression levels were imputed from genotype using a conditional energy-based model incorporating GRNs and cell-to-cell networks. Cell-type-specific nodes with dense connectivity were then incorporated into a deep linear model to predict phenotypes in each sample and prioritize cell types and genes for each trait. **(B)** (Left) Imputed single-cell expression values from LNCTP compared with observed snRNA-seq expression values, with best-fit lines for all cells and individual cell types. (Right) Correlations among imputed expression values for genes in excitatory and inhibitory neurons with best-fit lines. **(C)** Comparison of explained variance in gene expression from the LNCTP model with a baseline model using deconvolved, imputed bulk expression data and a model that includes only bulk expression data. Colored lines indicate the performance of individual cell types in each model (**p<0.01, two-tailed paired t-test over gene:cell-type pairs). **(D)** Schematic for LNCTP model interpretation, showing relationships between prioritized intermediate phenotypes for schizophrenia (SCZ, pink), bipolar disorder (BPD, orange) and ASD (light orange). Gene:cell-type:disease triplets are associated with salience (Sal) and coheritability (Co-h) values (*p<0.05, **p<0.01, two-tailed t-test; data S30). Significant cell type, GRN, and cell-to-cell associations are shown at the latent-embedding layers (p<0.05, two-tailed t-test; data S31-S32). Tree structures connect representative subgraphs (feature combinations) in each model (figs. S80-S81). The schematic also highlights QTL variants linked to the prioritized genes: (I) eQTL (bulk) chr15:60578052, (II) scQTL (Oligo) chr1:216891970, (III) eQTL (bulk) chr9:27902874, (IV) scQTL (Oligo) chr11:66017740, (V) eQTL (bulk) chr15:61553688, and (VI) scQTL (Astro) chr3:158668177. (Note, we shorten the readthrough transcript ANKHD1- EIF4EBP3 to ANKHD1 in ASD.) **(E)** UpSet plot for SCZ showing overlap between genes with the highest saliency per cell type or bulk expression, including four genes highlighted in panel D (colored circles). **(F)** Pearson correlations of LNCTP (excitatory neurons in SCZ) and CRISPR perturbation vectors for three example genes, when perturbation directions are matched vs. unmatched, and correlations are calculated across imputed genes (*p<0.05, one-tailed t-test; Fig. S88).

The LNCTP was trained as a conditional energy-based model that represents the joint distribution of the above “visible” variables conditioned on genotype, with additional latent layers (Fig. 8A; (*20*)). It imputes cell-type-specific gene expression from genotype with high cross-validated accuracy: the mean correlation between the imputed and experimentally observed expression profiles is 69% across major cell types and ∼78% in excitatory and inhibitory neurons (Fig. 8B). This corresponds to explaining 38% of the variance in cell-type gene expression (or, equivalently, estimating the heritability of cell-type gene expression *h^2^*), compared to a 34% baseline achieved by combining prior methods for bulk-imputation and cell-type deconvolution (*20*, *92*, *93*). The baseline does not include our derived GRNs and cell-to-cell networks, so the improvement represents the additional predictive performance possible with these networks (Fig. 8C). Moreover, the inclusion of imputed single-cell gene expression data also improves the overall prediction accuracy of disorders (discussed below) and accounts for a larger fraction of common-SNP heritability of these disorders beyond predictions based solely on bulk expression or polygenic risk scores (*94*) (table S13).

We exploited the ability of the LNCTP to impute missing data for discovery of cell-type-specific molecular phenotypes important for neuropsychiatric disorders. Doing so allowed us to link variants with their “intermediate” functional genomic activities, such as cell-type-specific gene expression, pathway activity, and cell-cell communication. We used a hierarchical linear architecture for the trait-prediction portion of the LNCTP, which performed comparably to or better than non-linear architectures (table S14-S15; (*20*)). Moreover, the LNCTP generates a model that is directly interpretable at multiple scales, avoiding many of the difficulties arising in the interpretation of deep neural networks, while maintaining a hierarchical structure. Our linear architecture allowed us to prioritize intermediate phenotypes by both gradient-based saliency, a metric directly derived from weights in the model, and co-heritability, which directly compares the genetic components of two traits. For instance, we can use the LNCTP to calculate the co-heritability of the genetic component of a particular gene’s cell-type-specific expression with respect to schizophrenia or other disorders (fig. S80; (*20*)).

Fig. 8D and fig. S81 provide an overview of key prioritized genes, cell types, and cell-to-cell interactions in various disorders (full lists in data S30-S32). We found 64, 51, 108, and 34 gene/cell-type pairs for schizophrenia, bipolar disorder, ASD, and AD, respectively (*20*). In particular, *TCF4*, the first identified cross-psychiatric disorder locus (*95*), is important for neurons in schizophrenia (*96*), *LINGO2* is important for excitatory neurons in bipolar disorder, and *ANKHD1* is highly weighted in ASD, supporting current hypotheses (*97*, *98*). Fig. 8E shows the associated cell types for the most highly prioritized genes. For example, *RORA* is important in many cell types for schizophrenia (but is, nevertheless, not prioritized in the bulk data; (*20*)). It is associated with retinoic-acid signaling, which has been proposed to be an important determinant of schizophrenia and bipolar risk (*99*). Further, we note the retinoic-acid signaling-associated gene *ESRRG* is prioritized in oligodendrocytes (Fig. 8D).

Overall, prioritized genes associated with bulk expression exhibit only a modest overlap with the prioritized cell-type-specific genes, indicating that integration of single-cell data in the LNCTP permits the prioritization of distinct genes compared to those found with bulk data alone (Fig. 8E). Moreover, as expected, the prioritized genes are enriched for cell-type-specific scQTLs, disease DE genes, and brain-related functional categories (figs. S82-S83). They are also enriched for prior GWAS and literature support as well as bottleneck locations in the regulatory network (figs. S84-S85; data S33). However, several genes specifically prioritized by the LNCTP are not differentially expressed for their respective disorders, including *MEF2A* and *ID1,* perhaps highlighting that they act through network effects (figs. S83, S85) (*100*).

In terms of cell types, excitatory neurons and microglia are prioritized in schizophrenia and bipolar disorder, supporting their importance for conferring genetic risk (*101*), with oligodendrocytes also prioritized in bipolar disorder. Moreover, in schizophrenia, we observed an increase in cell-to-cell interactions between excitatory neurons and microglia as well as a decrease between microglia and oligodendrocytes, consistent with the known glial dysregulation in the disease (Fig. 8D) (*78*).

We further used the LNCTP to perform in silico perturbation analysis, where we perturbed a specific gene’s expression and observed the induced expression changes in other genes (and the ensuing changes in trait propensity). Perturbations of our prioritized genes, as well as known drug targets (retrieved via DrugBank (*102*)) both induce overall expression changes strongly characteristic of case status (figs. S86A-B). As expected, the induced changes more strongly impact genes in close proximity to the perturbed gene in the GRNs (fig. S87; table S16). We synthesized the perturbations into a workflow to suggest potential drugs for repurposing with CLUE (*42*) by matching a perturbation’s effects to drugs inducing changes potentially complementary to those found in a particular disorder (fig. S86C; table S17; (*20*)).

Finally, to independently validate the results of our simulated perturbation analysis, we used data from CRISPR perturbations (CRISPRi and CRISPRa) applied to specific genes in glutamatergic neurons (*103*). Induced gene-expression changes resulting from the CRISPR perturbations are more highly correlated with those resulting from LNCTP perturbations when the direction of the perturbation is matched (versus not matched; Fig. 8F; figs. S88-S89; table S18; (*20*)). Furthermore, they are more aligned with the direction of case-control DE for LNCTP- prioritized genes than for non-prioritized ones (fig. S90). While more comprehensive validation is essential, these results offer promising indications that LNCTP can find verifiable prioritizations of gene/cell-type pairs.

## Discussion

Here, we used population-scale multi-omic data to build a comprehensive single-cell functional genomics resource (brainSCOPE) for investigating brain disorders in adults (Figs. S3-S5; (*20*)). The resource can be summarized at multiple levels: (1) raw data and metadata with a harmonized identifier system for each of the individuals; (2) quantifications of single-cell gene expression (count matrices) with a BICCN-compatible cell-typing system for the PFC; (3) lists of DE genes and differential cell-fractions for various phenotypes; (4) snATAC-seq signal tracks for various cell types and ENCODE-compatible regulatory elements (b-cCREs and scCREs), including lists of validated ones; (5) the variability for each gene and functional category (by individual, cell type, and brain region) and the associated sequence conservation of genes and regulatory elements; (6) a core set of GTEx-compatible scQTLs and other additional sets of QTLs (such as dynamic eQTLs); (7) full GRNs for each cell type, including enhancer-to-gene and TF-to-regulatory element links, and associated files relating each downstream gene to its most significant upstream regulators; (8) cell-to-cell communication networks (expressed as ligand-receptor-by-cell-type matrices); (9) integrative models with code for imputation, perturbation and prioritization of cell-type-specific functional genomics in brain disease; and (10) the resulting prioritized genes, cell types, and cell-to-cell linkages. The brainSCOPE portal also includes visualizer tools for many of the data types (fig. S4).

The resource allows for several important observations. These include the robustness of cell typing to population variation in 388 individuals and the identification, via shared scQTLs and dynamic scQTLs, of common regulatory programs between cell types. Moreover, by partitioning the observed expression variation, we identified certain drug targets demonstrating high variability between cell types but low variation across individuals (e.g., *CNR1*), a fact that is perhaps key to their therapeutic efficacy. We also found that gene-expression changes in certain neurons and glial cells can accurately predict the age of an individual.

Finally, a key outcome of our work is providing a set of promising targets for experimental validation. We see these falling into three classes. Class 1 comprises genes that are prioritized by the LNCTP model but not found by traditional DE analysis. This class is ideal for CRISPR assays seeking to test predicted cell-type and phenotypic effects. Other intriguing candidates are genes that have impacts on cell-to-cell communication spanning multiple cell types (class 2), and genes prioritized in disorders by the LNCTP with further support from DE analysis but lacking prior literature support (class 3). Overall, the LNCTP prioritized gene targets consistent with previous findings, and also suggested new avenues for investigation. We further used the LNCTP to simulate perturbations and make predictions regarding the effects of known drug-gene interactions on resulting phenotypes -- for instance, by perturbing drug-target expression levels. This application will potentially allow for assessing combinations of drugs for targeting multiple genes.

A few limitations should be noted regarding the data used in this study. Firstly, a number of recent works have demonstrated that RNA expression does not completely correlate with protein abundance, and this observation can be even more pronounced in the context of sub- regions within the brain (*104–106*). Another related complication is the uncertainty in the extent to which expression in postmortem tissues accurately reflects the expression patterns in live ones (*107*).

Future efforts could potentially address these limitations. They can also expand our analyses beyond the PFC and integrate functional genomic data from other connecting brain regions (such as the anterior cingulate cortex) to create a comprehensive brain-wide functional genomic atlas. This work could include the incorporation of developmental data as well as experimentally tractable models (such as those from cortical organoids); regulatory network changes over time can then be imputed across developmental axes toward fully mature brain GRNs. We could also incorporate imaging into our integrative model to improve our predictions of brain-associated phenotypes. Finally, more extensive validation of our results would be valuable, such as via targeted CRISPR assays.

Overall, the brainSCOPE resource has the potential to facilitate precision medicine by linking variants to specific cell types and their cell-type-specific impacts -- for example, to help identify the cell type of action for potential therapies. Through our integrative analyses, we provide an extensive collection of inferences and predictions for neuroscientists to verify in new cohorts, populations, assays, and experimental conditions.

## Materials and Methods Summary

The Materials and Methods for each section of the Main Text are available in the Supplementary Materials (*20*), which is organized using the same section headings as in the main text. These include a detailed description of the individuals and datasets assessed in the integrative analysis, protocols used for generating additional sequencing data and replication experiments for the analysis, and all computational and statistical analysis performed for each part of the integrative analysis.

## Acknowledgements

The authors thank the founder of the Allen Institute, P. G. Allen, for his vision, encouragement, and support. Rosemarie Terwilliger and Matthew J. Girgenti thank Keck Microarray Shared Resource (KMSR) and Yale Center for Genome Analysis (YCGA) at Yale University for their assistance with 10x Genomics single cell RNA-seq services.

## Author contributions

The following eleven co-first authors contributed equally to this work: P.S.E., J.J.L., D.C., M.J., J.W., C.G., R.M., C.L., S.Xu, C.D., and S.Lou. The following seven co-second authors contributed equally to this work: Y.Chen, Z.C., T.G., A.Hw., Y.L., P.N., and X.Z. All individually named authors contributed substantially to the paper through either data generation or analysis: data generation, T.E.B., J.B., L.B., L.C., Y.Cheng, M.F., J.F.F., S.G., J.G., N.H., N.L.J., R.K., Ji.L., S.Ma, M.M., S.Maz., J.R.M., D.Q., M.Sh., M.Sp., R.T., K.J.T., B.W., S.Xi., M.J.Ga., E.S.L., P.R., N.Se., K.P.W., and M.J.Gi.; and data analysis, P.S.E., J.J.L., D.C., M.J., J.W., C.G., R.M., C.L., S.Xu, C.D., S.Lou, Y.Chen, Z.C., T.G., A.Hw., Y.L., P.N., X.Z., T.E.B., T.C., L.C., Y.D., Z.D., M.Ga., D.Ga., S.G., E.H., G.E.H., A.Hu., Y.J., T.J., N.L.J., S.K., Ju.L., S.Li., J.M., E.N., N.P., M.P., H.P., A.S.R., T.R.R., N.Sh., K.J.T., G.W., Y.X., A.C.Y., S.Z., D.L., E.S.L., P.R., Z.W., H.W., J.Z., D.W., D.Ge., and M.Ge. The following seven senior authors played instrumental roles in this study: M.J.Ga., D.L., E.S.L., P.R., N.Se., Z.W., K.P.W., and H.W. The following five corresponding authors co-led the analysis: M.J.Gi., J.Z., D.W., D.Ge., and M.Ge.

## Data and materials availability

The brainSCOPE resource was developed from raw sequencing data (snRNA-Seq, snATAC-Seq, snMultiome, and genotype) derived from 12 individual cohorts, including eight PsychENCODE cohorts and four external cohorts. Raw datasets for the PsychENCODE cohorts, as well as protected-access integrated datasets such as imputed genotypes, are available at the PsychENCODE Knowledge portal (*108*). For the external cohorts, AMP-AD raw datasets and imputed genotypes are available at the AD Knowledge Portal (*109*). Girgenti- snMultiome datasets are deposited at NCBI GEO (GSE261983) (*110*). Ma-Sestan and Velmeshev datasets are available from their respective publications (*18*, *19*, *111*). Other key resources and additional datasets used in the integrative analysis are available in the supplementary materials (for smaller datasets) or on the brainSCOPE portal at http://brainscope.psychencode.org (for larger datasets) (*20*). Code used in this manuscript is deposited on GitHub and linked from the brainSCOPE portal (*112*).

## List of supplementary materials

Materials and Methods Figs. S1 to S90 Tables S1 to S18

Data S1 to S33 References (113–249)

