## Supplementary Materials for "Single-cell genomics and regulatory networks for 388 human brains"

**This PDF file includes:**

- Materials and Methods
- Figs. S1 to S90
- Tables S1 to S18
- Captions for Data S1 to S33
- References (*113-249*)

**Introduction of PsychENCODE data and more on the supplement**
**Materials and Methods**
**1 Constructing a single-cell genomic resource for 388 individuals**
**1.1 PsychENCODE Consortium Structure - Description of the PsychENCODE Consortium Structure and Contributions to the Resource**
**1.2 Dataset Overview - Overview of all the datasets combined to create the resource**
**1.3 Portal Overview - Description of supplemental files and datasets available on the brainSCOPE portal**
Key Resource Files
Raw Datasets
Output Files
Data Visualization
Outputs from the ROSMAP study analyses
Code - Using the Dockerized LNCTP Model
***1.4 snMultiome Dataset***
Human postmortem tissue
Nuclei isolation, microfluidic capture, and cDNA synthesis for snMultiome
**1.5 snRNA-seq Processing - Generation of cell-type-specific expression data**
Step 1. Count matrix generation, demultiplexing, and ambient RNA clean-up
Step 2. Per-fastq set/sample processing using Pegasus
Step 3. Per-study aggregation of processed sample and cell-type annotation
**1.6 Genotype Processing - Uniform analysis of genotype datasets**
Variant calling from WGS and RNA-seq
Genotype quality control (QC) and imputation
Genotype PCA analysis and QC
Rare variant and structural variant (SV) annotation and analysis
**1.7 Cell-type Fractions - Calculation of cell-type fractions from snRNA-seq data**
**1.8 Cell-type Fractions - Deconvolution of cell fractions for PsychENCODE Consortium bulk-RNA-seq data**
**1.9 DE Analysis - Differentially Expressed (DE) Genes for disease traits**
**1.10 Trajectory Analysis - Identifying DE genes over cell types by pseudotime**
Overall approach and differential expression analysis
Preprocessing and QC
Trajectory analysis
Differential expression along the trajectories
Post-processing
Additional results from the IT neuron trajectory analysis
**2 Determining regulatory elements for cell types from snATAC-seq**
**2.1 snATAC-seq Processing**
Step 1. Per-sample QC and filtering

Step 2. Per-sample dimensionality reduction and preliminary analysis
Step 3. Data aggregation and further QC
Step 4. Batch effect removal
Step 5. Peak calling
Step 6. Distal-peak-to-gene linkage
Step 7. Motif enrichment and TF footprinting analysis
Step 8. Processed data availability
**2.2 LDSC - Methods for Linkage Disequilibrium Score Regression (LDSC) enrichment of candidate regulatory elements**
**2.3 STARR-seq - Validation of putative enhancers in neural progenitor cells**
**2.4 b-cCREs - Identification of brain-specific cCREs**
**3 Measuring transcriptome and epigenome variation across the cohort at the single-cell level**
**3.1 Variance Partition - Partitioning transcriptome variation among cell types, individuals, and regions**
**3.2 Variance Partition - Quantifying epigenomic variation across cell types**
**3.3 Conservation - Cross-species sequence conservation**
**4 Determining cell-type-specific eQTLs from single-cell data**
**4.1 scQTLs - Cell-type-specific eQTL analysis**
**4.2 Bayesian scQTLs - scQTLs using Bayesian linear mixed effects models**
**4.3 Dynamic scQTLs - Dynamic sc-eQTL analysis**
**4.4 Isoform QTLs - Identify QTLs for isoform expression for each cell type**
**4.5 Allele-specific expression**
**4.6 mutSTARR-seq - Investigating the allelic effects of eQTLs on enhancers**
**4.7 STARR-seq and MPRA Validation - Validation of the scQTLs using STARR-seq and MPRA**
Identifying the eSNPs
Defining the active and control sets for STARR-seq and MPRA
Enrichment analysis of the scQTLs in MPRA and STARR-seq
**5 Building a gene regulatory network for each cell type**
**5.1 GRN construction - Construction of cell-type GRNs**
SCENIC
scGRNom
**5.2 GRN evaluation**
Overlaps of SCENIC results with snATAC peaks
Comparing cell type GRNs with tissue-naive GRNs
Variance in gene expression explained by GRN models
GRN stability
**5.3 CRISPR validation - Validation of TFs and target genes identified in GRNs**
**5.4 Network Characterization - Comparison of GRN structure across cell types**
Centrality analysis
Comparison of cell-type regulons with disease co-expression modules from bulk data

GO enrichment analysis
Motif analysis
Disease co-regulatory networks
5.5 Unifying TF-target Regulons
**6 Constructing a cell-to-cell communication network**
**6.1 Cell-to-Cell Network - Methods to build cell-to-cell networks**
**6.2 Cell-to-Cell Network - Methods to determine latent patterns**
**6.3 Cell-to-Cell Network - Validation of cell-to-cell network with spatial data**
**7 Assessing cell-type-specific transcriptomic and epigenetic changes in aging**
**7.1 Aging Cell Fractions - Single-cell aging cell-type fraction**
**7.2 Aging Cell Fractions - Deconvolution of bulk-RNA-seq data**
**7.3 Aging DE - DE genes in control aging and schizophrenia aging**
Aging DE genes in control group
Aging DE genes in the schizophrenia patient group
Aging DE genes and AD DE genes
**7.4 Aging STEM - Short Time-series Expression Miner analyses**
**7.5 Aging Model - Aging prediction using prioritized genes**
**7.6 Aging Chromatin**
**7.7 AD Model - Associating cell-type fractions and signatures with AD**
Inference of cell-type-specific gene expression and methylation
Building a model to predict AD
Multilayer perceptron
**8 Imputing gene expression and prioritizing disease genes across cell types with an integrative model**
**8.1 LNCTP Priors - Linear Network of Cell-Type Phenotypes imputation priors**
Per-sample, per-cell type metacells
Combining the metacells into z-score distributions
**8.2 LNCTP Framework**
**8.3 LNCTP Motivation - Motivation for linear network architecture**
**8.4 LNCTP Training**
Unary training
GMRF training
DNN training
LNCTP on AD Prediction
**8.5 LNCTP Interpretation**
Imputation of bulk and cell-type expression
Saliency-based cell-type and gene prioritization
Heritability and coheritability estimates
Prioritized subgraph analysis
Polygenic risk score calculations
**8.6 LNCTP Validation - In silico validation of LNCTP-prioritized genes and drug**

**targets**
8.6.1 Prior literature- and GWAS-based analysis of prioritized genes
8.6.2 Network analysis of prioritized genes
8.6.3 DE analysis of prioritized genes
8.6.4 Perturbation analysis of prioritized genes and drug targets
8.6.5 CLUE analysis
8.6.6 Network analysis of LNCTP perturbations
8.6.7 Ablation analysis
8.7 Independent CRISPR validation of LNCTP
8.7.1 Gene expression comparisons to CRISPR experiments
8.7.2 Gene regulatory network (GRN) overlaps with CRISPR differentially expressed genes (DEGs)
8.7.2.1 Diffusion-network-based approach
8.7.2.2 Hop-distance-based approach

### Introduction of PsychENCODE data and more on the supplement

This document provides an organized reference and includes four main sections: Materials and Methods, Supplementary Figures, Supplementary Tables, Supplementary Data. To present the data and results in an organized way, we prepared both the Materials and Methods section and Supplementary Figures section to align with our main text. In the Materials and Methods section, we use main text subheadings as primary headings. The secondary level of heading represents the content of each section, where we have included a ***precise heading*** (Bold and Italic) and detailed heading in a combined fashion ("***Dataset Overview*** - Overview of all the datasets combined to create the resource").

To link and cross-reference between the main text and this supplement as clearly as possible, we also list all precise headings that are linked to each main text section here. All precise headings connected to the main text can be used as a quick guide for finding information.

#### Quick Guide to Finding Information in the Materials and Methods Using [*Precise Heading*]

| Main Text Subheading | Precise Heading |
|---|---|
| Constructing a single-cell genomic resource for 388 individuals | "PsychENCODE Consortium Structure", "Dataset Overview", "*Portal Overview*", "snMultiome Dataset", "snRNA-seq Processing", "Genotype Processing", "Cell-type Fractions", "DE Analysis","Aging DE", "Trajectory Analysis" |
| Determining regulatory elements for cell types from snATAC-seq | "snATAC-seq Processing", "LDSC", "STARR-seq", "b-cCREs", |
| Measuring transcriptome and epigenome variation across the cohort at the single-cell level | "Variance Partition","Conservation" |
| Determining cell-type-specific eQTLs from single-cell data | "scQTLs", "Bayesian scQTLs", "Dynamic scQTLs", "Allele-specific expression", "Isoform QTLs", "STARR-Seq", "mutSTARR-Seq" |
| Building a gene regulatory network for each cell type | "GRN Construction", "GRN evaluation", "CRISPR Validation", "Genotype processing", "Network Characterization", "Unifying TF-target Regulons" |

| | |
|---|---|
| Constructing a cell-to-cell communication network | “Cell-to-Cell Network” |
| Assessing cell-type-specific transcriptomic and epigenetic changes in aging | “Aging Cell Fractions”, “Aging DE”, “Aging STEM”, “Aging model”, "Aging Chromatin", “AD Model” |
| Imputing gene expression and prioritizing disease genes across cell types with an integrative model | “LNCTP Framework”, “LNCTP Priors”, “LNCTP Training”, “LNCTP motivation”, “LNCTP Interpretation”, “LNCTP Validation”, “Independent CRISPR validation of LNCTP” |

The supplementary figures and tables are both numbered based on the order in which they are mentioned in the main text. Note that many associated data files are available with unique file IDs on the brainSCOPE portal: http://brainscope.psychencode.org and https://brainscope.gersteinlab.org. References to the supplementary materials in the main manuscript are contextualized in this document with the tag “**Main manuscript reference**. ”

### Materials and Methods

#### 1 Constructing a single-cell genomic resource for 388 individuals

##### 1.1 *PsychENCODE Consortium Structure* - Description of the PsychENCODE Consortium Structure and Contributions to the Resource

The PsychENCODE Consortium (http://www.psychencode.org/) consists of over 200 researchers across 20+ institutions who are studying the effects of functional genomic elements in individuals with neuropsychiatric disorders (*113*). PsychENCODE consists of several internal committees and working groups. In particular, the following groups contributed to the data collection and integrative analysis presented in the brainSCOPE Resource:

- *Data Generation Center*: Performed single-nucleus RNA sequencing (snRNA-seq), single-nucleus ATAC-sequencing (snATAC-seq), single-nucleus Multiome (snMultiome), and genotyping for 313 prefrontal cortex (PFC) samples in disease and control individuals.
- *Data Analysis Center*: Performed uniform computational processing and integration of sequencing datasets for the 313 PsychENCODE samples with 20 non-PsychENCODE samples generated for this cohort and 55 samples from external sources, and generated key resources for brainSCOPE.
- *Validation Working Group*: Performed CRISPR, massively parallel reporter assay (MPRA), STARR sequencing (STARR-seq) validation, and spatial transcriptomics experiments to validate key resources.
- *Data Coordinating Center*: Maintained raw datasets for brainSCOPE in appropriate data repositories, and constructed web portal data visualization browsers for key resources and ancillary datasets.

##### 1.2 *Dataset Overview* - Overview of all the datasets combined to create the resource

**Main manuscript reference:** First supplementary reference in the first paragraph of "Constructing a single-cell genomic resource for 388 individuals."

The collection of snRNA-seq, snATAC-seq, and snMultiome datasets used in this analysis encompasses control, schizophrenia, bipolar disorder, autism spectrum disorder (ASD), Alzheimer's disease (AD) and post-traumatic stress disorder (PTSD) samples from the PFC from several studies that fall under the scope of the PsychENCODE Consortium (*108*) These studies include the CommonMind Consortium (CMC), UCLA-ASD, SZBDMulti-seq, MultiomeBrain, DevBrain, IsoHuB, PTSDBrainomics, and Lieber Institute for Brain Development (LIBD) studies. For the snRNA-seq component, the library preparation methods varied from study to study, including: 10x Genomics (UCLA-ASD, DevBrain, IsoHuB, PTSDBrainomics, and LIBD); 10x Genomics + MULTI-seq (*108*); 10x Genomics + CellHashing (*114*); and 10x Genomics Multiome (snRNA-seq + snATAC-seq). Additional snATAC-seq data were obtained from the UCLA-ASD study. The analysis also encompasses data from published studies and repositories, including the ROSMAP (*115*) study (as available on the AD Knowledge Portal (*116*)), the Ma-Sestan study (*19*), and the Velmeshev study (*18*), as well as additional snMultiome data (described below) from control individuals (labeled as “Girgenti-snMultiome” in the meta-data file), processed and analyzed exclusively for this paper. All functional genomic samples in the cohort were taken from tissue samples of various regions within the dorsolateral PFC (DLPFC), and all samples in the integrated analysis were from adults (at least 13 years of age, ranging up to 90+ years old).

Metadata for all samples used in this study by cohort as well as available data modalities and uniform clinical and demographic information for each sample are shown below as **data S1,** with additional information for mapping IDs across modalities provided in **data S2**. The datasets used as inputs for each downstream analysis are listed in **data S3** and visualized as a dependency graph in **fig. S2**.

**File:** PEC2_sample_metadata.txt (**data S1**): This file contains clinical and demographic meta-data for each sample in the brainSCOPE Resource.
**File:** PEC2_sample_mapping.xlsx (**data S2**): This file contains mapping of uniform IDs for each sample across cohorts and data modalities (snRNA-seq, snATAC-seq, and genotype data).

#### 1.3 ***Portal Overview*** - Description of supplemental files and datasets available on the brainSCOPE portal

**Main manuscript reference:** First supplementary reference in the last paragraph of “Introduction” and first supplementary reference in the first paragraph of the “Discussion.”

Datasets produced by the PsychENCODE Consortium include raw and analyzed population-scale single-cell multi-omics data in a cohort consisting of 388 individual samples from the adult human PFC of healthy controls and individuals afflicted by neuropsychiatric diseases. These data include snRNA-seq, snATAC-seq, snMultiome, and genotype data integrated and uniformly processed from 12 different cohorts. Together, the raw datasets and processed output files described in the manuscript and supplement comprise the *brainSCOPE* (Brain Single-Cell Omics for PsychENCODE) *Resource*.

All brainSCOPE-related datasets are available at http://brainscope.psychencode.org and https://brainscope.gersteinlab.org. The portal contains lists of available data and links to a data visualization tool called PsychSCREEN, as described below. Screenshots of the main brainSCOPE portal, the protected dataset portal, and the PsychSCREEN browser are available in **figs. S3, S4,** and **S5,** respectively.

##### Key Resource Files

This page of the brainSCOPE portal provides users with a list of select files generated from each major analysis of the paper. It is intended for end users who wish to easily access key results from each of the major paper analyses for use in downstream analyses. Files on this page include sample metadata; cell-level and pseudobulk summary matrices for gene expression by cell type; single-cell cis-regulatory element (scCRE) regions for each cell type;

lists of differentially expressed (DE) genes by condition and cell type; individual and cell type-derived variation for all genes; single-cell quantitative trait loci (QTL) callsets from the primary scQTL analysis; cell-type-specific gene regulatory networks (GRNs); cell-to-cell communication networks; and lists of genes prioritized by disease from the Linear Network of Cell-type Phenotypes (LNCTP) model. A screenshot of this webpage is shown below in **fig. S3** and is available online at http://brainscope.psychencode.org/key_resource_files.html and https://brainscope.gersteinlab.org/key_resource_files.html.

**Raw Datasets**

The page labeled "Raw Data" provides links to snRNA-seq, snATAC-seq, and genotype file sets stored in protected-access data repositories. In particular, raw sequencing datasets include snRNA-seq, snATAC-seq, and genotype data (single-nucleotide polymorphism [SNP] microarray, whole-genome sequencing [WGS], or exome sequencing) for all samples derived from PsychENCODE Consortium cohorts. A screenshot of the main protected access data repository for PsychENCODE datasets is provided below as **fig. S5**. Links are also provided for datasets from external cohorts that were analyzed alongside the PsychENCODE cohorts, including ROSMAP samples that are hosted on the AD Knowledge Portal or Velmeshev samples that are hosted on NCBI Gene Expression Omnibus. Each individual dataset is linked using accession numbers for long-term data archival.

The following data matrix files hosted on the brainSCOPE portal provide links to datasets for each sample on Synapse or other repositories:

**File:** raw_sequencing_data_links.xlsx: This file contains accession numbers and links to all available snRNA-seq, snATAC-seq, and genotype data for each sample that are publicly available to download from Synapse or other repositories.

**Output Files**

The page labeled "Output Files" hosts all processed datasets with open access described in the manuscript and supplement. To better connect the supplementary text with our resource files hosted on the brainSCOPE portal, we also list the resource file name and provide a brief description of these files at the end of each related Materials and Methods section. A screenshot of this webpage is shown below in **fig. S3** and is available online at http://brainscope.psychencode.org/integrative_files.html and https://brainscope.gersteinlab.org/integrative_files.html.

**Data Visualization**

Finally, a link is provided to a tool for interactive visualization of the brainSCOPE Resource data, which is integrated into the PsychSCREEN genomic data browser. The tool includes genome browsers for snATAC-seq peaks, scQTLs, and GRN annotations. We also provide a tool to visualize single-cell expression data for all genes by cell type across subcohorts, including interactive Uniform Manifold Approximation and Projection (UMAP) and dot plots. Finally, summary dot plots for gene variation and DE genes by disease and cell type are provided. This tool is available at https://psychscreen.wenglab.org/psychscreen/single-cell.

Datasets available on PsychSCREEN can also be downloaded at https://psychscreen.wenglab.org/psychscreen/downloads. Screenshots of example data visualizations are available in **fig. S4**, and an explanation of all PsychSCREEN features is provided on the brainSCOPE portal at http://brainscope.psychencode.org/psychscreen_example.html and https://brainscope.gersteinlab.org/psychscreen_example.html.

##### Outputs from the ROSMAP study analyses

Per the data use requirements for the AD Knowledge portal (*116*), we have made the processed snRNA-seq expression datasets (cohort and per-individual) and imputed genotype files for ROSMAP available in the portal.

The ROSMAP h5ad file and the individual-specific gene-by-cell expression matrices are available via the AD Knowledge Portal (https://adknowledgeportal.org). The AD Knowledge Portal is a platform for accessing data, analyses, and tools generated by the Accelerating Medicines Partnership (AMP-AD) Target Discovery Program and other National Institute on Aging (NIA)-supported programs to enable open-science practices and accelerate translational learning. The data, analyses and tools are shared early in the research cycle without a publication embargo on secondary use. Data is available for general research use according to the following requirements for data access and data attribution (https://adknowledgeportal.org/DataAccess/Instructions).

##### Code - Using the Dockerized LNCTP Model

The LNCTP model is currently available as a Docker container and can be accessed publicly on https://hub.docker.com/repository/docker/icefirecloud/lnctp-server/general. This allows users to easily run or modify the model on their local systems. While the model can run on both CPU and GPU, it is recommended to use a GPU for the training phase to expedite the process.

To use the docker file, users must install Docker first, and then use the following steps (for a Linux-based system):

*docker pull icefirecloud/lnctp-server:latest*
*docker run -it icefirecloud/lnctp-server bash*

Now the user should be in the docker container:

*cd lnctp_code_asd*
*python lnctp_test_models.py*
*python lnctp_train_models.py*

Corresponding outputs will be printed during processing. Users may also switch to other folders (SCZ, BPD) and run the corresponding LNCTP scripts for other predictions.

#### 1.4 ***snMultiome Dataset***

**Main manuscript reference:** Fourth supplementary reference in the first paragraph of "Constructing a single-cell genomic resource for 388 individuals."

**Human postmortem tissue**

Human DLPFC (Broadman Area 9/46) samples were collected by the Girgenti laboratory from the NIH NeuroBioBank (https://neurobiobank.nih.gov/) (*117*) \cite{29496155} following the guidelines provided by the Yale Human Investigation Committee. Samples from the NIH NeuroBioBank were chosen to be free of neurodegenerative conditions, stroke, head injury, HIV, COVID, and any known neuropsychiatric conditions. Human tissues were collected and handled in accordance with ethical guidelines and regulations for the research use of human brain tissue set forth by the NIH (http://bioethics.od.nih.gov/humantissue.html) and the World Medical Association Declaration of Helsinki (http://www.wma.net/en/30publications/10policies/b3/index.html). Appropriate informed consent was obtained, and all available non-identifying information was recorded for each specimen. No obvious signs of neuropathological alterations were observed for any of the human specimens considered and analyzed in this study. For all specimens, regions of interest were sampled from frozen tissue slabs or whole specimens stored at -80ºC.

**Nuclei isolation, microfluidic capture, and cDNA synthesis for snMultiome**

To obtain pure and intact nuclear populations, brain tissues were homogenized by a Dounce homogenizer in an ice-cold isolation buffer containing 2M sucrose. All buffers were ice-cold and all reagents used for nuclear isolation were molecular biology grade unless stated otherwise. A total of 20-30 mg of pulverized tissue was added into 5 ml of ice-cold lysis buffer (320 mM sucrose [Sigma #S0389], 5 mM $CaCl_2$ [Sigma #21115], 3 mM $Mg[Ace]_2$ [Sigma #63052], 10 mM Tris-HCl [pH 8; AmericanBio #AB14043], protease inhibitors without EDTA [Roche #11836170001], 0.1 mM EDTA [AmericanBio #AB00502], RNAse inhibitor [80 U/ml; Roche #03335402001], 1 mM dithiothreitol [DTT; Sigma #43186], and 0.1% TX-100 [v/v; Sigma #T8787]). Reagents DTT, RNAse protector, protease inhibitors, and TX-100 were added immediately before use. The suspension was transferred to a Dounce tissue grinder (15 ml volume, Wheaton #357544, autoclaved, RNAse free, ice-cold) and homogenized with loose and tight pestles, 30 cycles each, with constant pressure and without introduction of air. The homogenate was strained through a 40-µm tube top cell strainer (Corning #352340) pre-wetted with 1 ml isolation buffer (1,800 mM sucrose [Sigma #S0389], 3 mM $Mg[Ace]_2$ [Sigma #63052], 10 mM Tris-HCl [pH 8; AmericanBio #AB14043], protease inhibitors without EDTA [Roche #11836170001], RNAse inhibitor [80 U/ml, Roche #03335402001)] and 1 mM DTT [Sigma #43186]. An additional 9 ml of isolation buffer was added to wash the strainer. The final 15 ml of solution was mixed by inverting the tube 10x and carefully pipetted into two ultracentrifuge tubes (Beckman Coulter #344059) onto the isolation buffer cushion (5 ml) without disrupting the phases. The tubes were centrifuged at 30,000 x g for 60 min at 4ºC on an ultracentrifuge (Beckman L7-65) and rotor (Beckman SW41-Ti). Following the ultracentrifugation, the supernatant was carefully and completely removed, and 100 µl of resuspension buffer (250 mM sucrose [Sigma #S0389], 25 mM KCl [Sigma #60142], 5 mM $MgCl_2$ [Sigma #M1028], 20 mM Tris-HCl [pH 7.5; AmericanBio #AB14043; Sigma #T2413], protease inhibitors without EDTA [Roche #11836170001], RNAse inhibitor [80 U/ml; Roche #03335402001], and 1 mM DTT [Sigma #43186]) was added dropwise on the pellet in each tube and incubated on ice for 15 min. Pellets were gently dissolved by pipetting 30x with a 1 ml pipette tip, pooled, and filtered through a 35-μm tube top cell strainer (Corning #352235). Finally, nuclei were counted on a Countess cell counter (ThermoFisher) and diluted to 1 million/ml with sample-run buffer (0.1% bovine serum albumin [Gemini Bio-Products #700-106P], RNAse inhibitor [80 U/ml; Roche #03335402001], and 1 mM DTT [Sigma #43186]) in Dulbecco's phosphate-buffered saline (Gibco #14190).

The nuclei samples were placed on ice and taken to the Yale Center for Genome Analysis core facility, where they were processed with targeted nuclei recovery of 20,000 nuclei per sample on a microfluidic Chromium System (10x Genomics) following the manufacturer's protocol (10x Genomics, Chromium Next GEM Single Cell Multiome ATAC + Gene Expression Reagent Bundle, PN-1000283). Libraries were sequenced with paired-end 150 bp reads on an Illumina NovaSeq 6000 to a target depth of 250 million read pairs per sample.

#### 1.5 ***snRNA-seq Processing*** - Generation of cell-type-specific expression data

**Main manuscript reference:** Second supplementary reference in the first paragraph and first reference in the second paragraph of "Constructing a single-cell genomic resource for 388 individuals."

The snRNA-seq processing pipeline was constructed based on published best practices and new benchmarking metrics for existing methods. The pipeline is mostly implemented in Python, except for the final cell-type annotation steps that involve the R-based program Azimuth. The overall workflow can be summarized in three main steps:

1. Count matrix generation, demultiplexing, and ambient RNA clean-up
2. Per-fastq set/sample processing using Pegasus (https://pegasus.readthedocs.io/en/stable/) (*118*)
3. Per-study aggregation of processed sample and cell-type annotation

In the following section, we present each of the steps in greater detail.

**Step 1. Count matrix generation, demultiplexing, and ambient RNA clean-up**

A schematic for this portion of the workflow is presented in **fig. S6A**.

**Count matrix generation:** The count matrix was generated using CellRanger *count* v6.0(https://support.10xgenomics.com/single-cell-gene-expression/software/pipelines/latest/using/count) (*119*). Each sample was run independently, and no aggregation (using CellRanger *aggr*) was carried out. However, if the same sequencing sample was run through multiple lanes and shared the same sequencing sample identifier with different lane numbers (L001, L002, etc.), then we used the CellRanger *count* feature to automatically pool all contents of the 'same-sample different-lanes' fastqs. We also used the '--include-introns' flag for all samples, as the studies under consideration all involve snRNA-seq, and therefore necessitate a quantification of pre-mRNA (and thus pre-spliced) transcripts as well. Additionally, we specified the chemistry of the sample being quantified, whether based on 10x Genomics v2 or v3 chemistry or ARC chemistry. The same initial procedure for per-cell RNA quantification was followed for the MULTI-seq and CellHashing data as well, with the demultiplex step using the corresponding hashtag oligo (HTO) files described in the following section. Gene expression datasets from snMultiome samples were also processed separately from the snATAC-seq reads, and we used the "ARC-v1" tag to represent chemistry in the pipeline.

**Demultiplexing:** The studies analyzed included data from MULTI-seq and CellHashing multiplexing assays. To process these data, we first quantified the per-cell HTO counts. There are two standard programmatic options for this: using the R-based deMULTIplex package (https://github.com/chris-mcginnis-ucsf/MULTI-seq) (*120*) or using the command-line-based CITE-seq-Count package (*114*, *121*). For reasons of programmatic convenience, we chose to use the CITE-seq-Count package (v1.4), with the command embedded into our Python scripts (*122*). The options provided for this command include: the R1 and R2 fastq files for the HTO reads; a csv file containing the HTO barcodes and an associated name for the hashtag to be included in downstream HTO quantifications; the first (*cbf*) and last (*cbl*) locations for the cell barcodes in the file; the first (*umif*) and last (*umil*) locations for the unique molecular identifiers (UMIs) in the file, which are chemistry specific; and the approximate number of cells expected in the sample or a list of cell barcodes. For the tags file, we used the data providers' lists of HTO barcodes followed by generic names – Hashtag_1, Hashtag_2, etc. We set *cbf* = 1, *cbl* = 16, *umif* = 17, and *umil* = 26 for the v2 chemistry samples and *umil* = 28 for the v3 chemistry samples. We chose to provide an expected number of cells instead of explicit cell barcodes. We supplied the expected number of cells using the *metrics_summary.csv* file output by CellRanger. From that file, the number of cells output by CellRanger's own cell-identification algorithm was extracted and 500 was added to it as a simple buffer (to account for possible undercounting of cells by CellRanger's algorithm).

**Ambient RNA clean-up:** To more carefully separate out true cells from empty droplets with ambient RNA, we used the program *remove-background* (https://cellbender.readthedocs.io/en/latest/index.html) from the *CellBender* package (*122*). We used the program in command-line form wrapped in our Python script. The program is optimized to run on GPUs (there is a substantial difference in the runtime depending on whether GPUs or CPUs are used), and accordingly we implemented this portion of the pipeline on a GPU (adding the --cuda flag). The input to the program is the raw output .h5 file from CellRanger *count* without filtering for cells identified by CellRanger. The options included in the program run are: a target false positive rate (--fpr) of 0.01; the number of training epochs (--epochs) = 150; the rough expected number of cells (--expected-cells) = the output in *metrics_summary.csv* from CellRanger *count*, as in the CITE-seq-Count run; and the total number of droplets to be included (--total-droplets-included) in the analysis, set to be the expected number of cells + 20,000 (chosen to be large enough to encompass many empty droplets for the training). The output of this step is a .h5 file, where the empty droplets are filtered out, leaving just the inferred true cells. This .h5 file is chosen as the input for the downstream analyses in Pegasus.

**Step 2. Per-fastq set/sample processing using Pegasus**

The primary steps for this part of the analysis are outlined in **fig. S6B**. The steps are applied to the CellBender output for each sample separately, although some pooling may have been carried out across parallel lanes for the same sample, as described above. Many of the steps are applied analogously to those in the Pegasus tutorial (*pegasus-tutorials.readthedocs.io/ en/latest/_static/tutorials/pegasus_analysis.html*). After filtering cells based on the lower bounds shown in **fig. S6B**, we removed 1,135 genes included in the MitoCarta v3.0 database (*123*) such as mitochondrial genes and certain genes highly correlated with RNA sample quality (see, for example, Hodge et al. 2019 (*124*). The robust genes were identified and the counts matrix was log-normalized using the default options in Pegasus.

At this stage, if the data were from a multiplexed experiment, the matrices were decomposed into cells from each of the component samples using the Pegasus *demultiplex* algorithm. The inputs to this were the matrix, feature, and barcode files from the CITE-seq-Count run. Only cells that were identified as singlets in the demultiplexing step were retained.

Next, doublets were identified using a combination of two computational methods. These steps were carried out for the demultiplexed samples as well. The rationale is that demultiplexing removes inter-sample doublets, while intra-doublet samples still need to be removed. We found that a combination of the program Scrublet (*125*) in default mode and DoubletDetection *(126)* worked well. The parameters for the DoubletDetection *BoostClassifier* algorithm include: *n_iters* = 25, *use_phenograph* = *False*, and *standard_scaling* = *True*. The subsequent *predict* function uses the parameters *p_thresh* = $1e^{-16}$ and *voter_thresh* = 0.3.

After doublet removal, we aggregated the demultiplexed samples again for robust gene identification, highly variable gene selection (5,000 genes chosen), principal component analysis (PCA), batch correction using Harmony (*127*), nearest-neighbor detection, Leiden clustering, and UMAP dimensionality reduction.

We carried out differential expression analysis using the t-test, comparing the expression of genes in every cluster against all others. Finally, cell-type inference was carried out based on a series of marker gene collections using the *infer_cell_types* function. It should be noted that many of these later steps (from the highly variable gene selection onwards) were not strictly necessary for further analysis. The raw count matrices for each sample, after doublet removal and demultiplexing, were used in the next steps without referencing any of the cell-type annotations in this per-sample stage.

**Step 3. Per-study aggregation of processed sample and cell-type annotation**

A schematic depicting the set of steps for this stage of the workflow is shown in **fig. S7**. First, we aggregated all samples across each study, with the demultiplexed sample names being used as identifiers for the MULTI-seq and CellHashing datasets. We specifically selected raw count matrices for the following analyses. We then proceeded with joint analysis steps on the aggregated AnnData object in Pegasus. Many of the steps remained the same as above (although no doublet detection was carried out at this stage), until the cluster annotation process. One difference is that we increased the Leiden clustering resolution parameter to between 4.0 and 6.0. The intention was to recover as many cells as possible by more finely dividing the clusters, before removing those clusters that failed to be annotated (as described in the following section). The number varied simply because the limit for UMAP generation was 64 clusters, and the same resolution parameter resulted in different numbers of clusters for different studies.

**Cluster annotation:** Cluster annotation proceeded in a two-step process (**fig. S9**). We first used the Pegasus' *infer_cell_types* function to associate the Leiden clusters with reference cell types based on the hybrid marker gene sets obtained from merging excitatory and inhibitory neuronal subclass markers from the BRAIN Initiative Cell Census Network (BICCN) taxonomy (**fig. S11)** (*128*) and non-neuronal subclasses from Ma-Sestan (file *Documentation_merged_subclass_markers.json)* (*19*). Note that four of the non-neuronal subclasses are unique to the Ma-Sestan dataset: *Immune*, *RB*, *PC*, and *SMC*. The final subclass annotations (also detailed in **table S3,** color annotations detailed in **table S4**) included the following:

**Excitatory Neurons:** *L2/3 IT*, *L6 IT*, *L4 IT*, *L5 IT*, *L6 IT Car3*, *L5 ET*, *L6 CT*, *L5/6 NP*, *L6b* (L# signifies the cortical layer context; IT = intra-telencephalic, ET = extra-telencephalic, CT = cortico-thalamic, NP = near-projecting)

**Inhibitory Neurons:** *Lamp5*, *Pax6*, *Sncg*, *Vip*, *Lamp5 Lhx6*, *Chandelier*, *Pvalb*, *Sst*, *Sst Chodl*

**Non-Neuronal Cells:** *Astro* (Astrocytes), *Endo* (Endothelial cells), *VLMC* (Vascular Leptomeningeal cells), *Micro* (Microglia), *Oligo* (Oligodendrocytes), *OPC* (Oligodendrocyte precursor cells), *Immune* (immune cells), *RB* (Red Blood lineage cells), *PC* (*Pericytes*), SMC (*Smooth Muscle Cells*)

These annotations, in turn, were used at this stage of the analysis because of their purported robustness across brain regions and species (see, for example, the discussion in (*129*)). Given these annotations, and the most likely assignments (sometimes a many-to-one assignment of subclasses to clusters), we used Pegasus' *infer_cluster_names* and *annotate* functions to assign a single best-fit subclass assignment. We then removed all unassigned clusters. Our rationale in doing so was that these unassigned clusters mainly consisted of low-expression cells, likely reflecting cellular debris. For example, we found that some clusters were enriched for synaptic and transmembrane protein expression, while the overall number of genes expressed was very small. We speculated that these clusters included extracellular debris and accordingly removed them. We thus chose a conservative approach to cluster inclusion in our downstream analyses.

However, because of the ambiguity of the resolution in cluster-based assignment strategies, we did not finalize the cell assignments based on the marker gene analysis. Instead, the marker gene analysis was used for the aforementioned cluster removal. The remaining clusters were then processed through Seurat/Azimuth (*130*) pipelines to assign subclass annotations. The advantage of the Azimuth approach is that it uses cell-based assignment; that is, each cell is individually assessed against a reference cell atlas to find the most likely assignment of cell type. Specifically, the .h5ad objects from the Pegasus processing (after the aforementioned cluster removal) were read into Seurat objects using functions in the *SeuratDisk* package, and processed as follows: first, the AnnData object was converted into a .h5Seurat object; second, the .h5Seurat object was loaded using the *LoadH5SeuratObject* function; third, due to issues with reading the raw counts through this mechanism, the raw counts for each study were independently exported into .npz format and reassigned to the Seurat object in the "counts" slot; fourth, the dataset was processed using the *SCTransform* function in Seurat (*130*); finally, the *FindTransferAnchors* and *TransferData* functions were used to add subclass assignments as metadata in the Seurat object. We exported these assignments to a separate file for integration with the Pegasus objects downstream. To transfer anchors, we performed the same preprocessing steps on the raw counts for the reference atlas cells from the BICCN (118,291 cells pre-annotated with the reference neuronal subclasses) and the Ma-Sestan study (172,120 cells pre-annotated with the reference subclasses). Note that the marker genes for the subclasses were derived from the same dataset.

**Merging of the results from the two reference atlases.** We first annotated all the cells with the BICCN and Ma-Sestan schemes separately. To reconcile the results, we performed the following steps:

1. We assessed the cell-type label for each annotation scheme and assigned cell types to the appropriate cell class (excitatory, inhibitory, or glial).
2. If the cell classes were the same for the BICCN and the Ma-Sestan schemes, we retained the cell barcode. If the cell classes were different, we removed the cell barcode. We have found that cells that are mismatched in cell class between the two schemes tend to be of lower technical quality, measured in terms of the number of genes expressed and the number of UMI counts.
3. For all the cell barcodes that were retained, if the cell class was excitatory or inhibitory, we kept the BICCN label in the final annotation; if the cell class was glial, we kept the Ma-Sestan annotation.
4. These new annotations were merged with the .h5ad objects by filtering out cells that could not be reconciled at the cell-class level.

Once the subclasses were assigned, we added them to the metadata in the Pegasus .h5ad (AnnData) objects. The .h5ad objects were rerun through dimensionality reduction and reclustered so that the UMAP coordinates reflected the post-filtration groups of cells. We have provided these annotated objects for download via Synapse (see above). The matrix “raw.X" contains the raw UMI counts and “X” contains the log-normalized counts. The annotations of the .obs dataframe of the AnnData object include marker gene annotations (“anno”) of clusters as well as Azimuth labels (“azimuth”, “subclass”). The final annotations are contained under the “subclass” column.

We also have provided the individual IDs associated with each cell and sample. This is especially relevant for the multiplexed datasets. In addition to the .h5ad files, we have uploaded to Synapse and the brainSCOPE portal tab-delimited expression matrix files for each individual in the analyses, where genes with HUGO Gene Nomenclature Committee (HGNC) symbols mark the rows and the cells are arrayed along the columns. The column headers are the Azimuth labels of the cells. We emphasize the fact that each individual was assigned a separate matrix; this means that if parallel samples were sequenced for the same individual in the same cohort, the final matrix pools all cells from these samples.

Additional notes follow for specific datasets:

1. The UCLA-ASD dataset includes cells from BA44/45 and BA4/6 regions.
2. For the UCLA-ASD dataset, when comparing the genotypes from WGS, exome or SNP array assays with genotypes obtained from snRNA-seq reads, we found that the samples 10BW, 63BW, 65BW, and 37BW had potential mismatches. These samples were removed in the downstream analysis.

We provide links to uniformly processed individual-level gene expression matrices and complete expression datasets (*.h5ad files) for downstream analysis on the brainSCOPE portal.

In addition, we provide several files related to the cell-typing scheme used in our analysis, along with the source code used to process snRNA-seq data:

**File:** [sample]-annotated_matrix.txt.gz: These files represent expression matrices for individual samples in our cohort. Each column is labeled with a cell type from our harmonized classification scheme (see below), while each row represents the normalized expression values for a single gene (noted with an HGNC symbol).
**File**: BICCN_mat.RDS: This R Data Object contains the PFC cell-type annotation scheme from the BICCN Consortium.
**File**: Ma_Sestan_mat.rds: This R Data Object contains the PFC cell-type annotation scheme from Ma-Sestan.
**File**: BICCN_meta_share.RDS: This file contains the harmonized PFC cell-type annotation scheme used for analysis in the study (the merger between BICCN_mat.rds and Ma_Sestan_mat.rds).
**File**: Azimuth_mapping.R: This R script is used to annotate cell types in an Azimuth object.
**File:** reconcile_annotations.py: This Python script is used to generate the harmonized cell typing scheme from the input matrices.
**File**: PsychENCODE_scRNA_pipeline-main.tar.gz: File contains source code used for processing snRNA-Seq datasets.

#### 1.6 ***Genotype Processing*** - Uniform analysis of genotype datasets

**Main manuscript reference:** Third supplementary reference in the first paragraph of "Constructing a single-cell genomic resource for 388 individuals"; second supplementary reference in the second paragraph of "Building a gene regulatory network for each cell type."

In order to generate a uniform set of genotyped variants across all 12 cohorts in our study for downstream analysis, we integrated the genotypes of 383 individuals from a diverse range of data sources (SNP microarray, exome and WGS, and directly from the snRNA-seq samples) for filtering and imputation.

**Variant calling from WGS and RNA-seq**

For samples with available exome and WGS data (derived from the ROSMAP, UCLA-ASD, Velmeshev, Girgenti-snMultiome, and DevBrain cohorts), we re-processed next-generation sequencing data to uniformly identify variants. In particular, we used the GATK4 Best Practices pipeline v1.0 (*131*) to identify variants from either raw fastq files or pre-aligned bam files **(fig. S8A)**. Using this pipeline, we first aligned samples to the reference genome (hg38) using BWA v.0.7.15 (*132*), and then marked duplicate reads using PicardTools v.2.16.0. We then used GATK v.4 to perform HaplotypeCaller and Base Quality Score Recalibration steps to identify variants at a per-sample level, and further performed Variant Quality Score Recalibration and Joint Genotyping across all samples. We used the default parameters for each pipeline based on data type (exome or WGS) to generate final variant call format (VCF) callsets for each cohort.

For all non-multiplexed snRNA-seq samples, including 29 samples without available WGS or SNP array data, we further identified variants directly from the snRNA-seq samples. To do this, we used the GATK4 RNA germline variant calling pipeline (**fig. S8A**), which aligns the snRNA-seq data to the reference genome (hg19) using STAR v.2.5.3a (*133*) and incorporates an additional step to split reads spanning splice junctions before calling variants using HaplotypeCaller. At this step, we performed JointGenotyping as in the DNA-based GATK Best Practices (as the RNA-based Best Practices pipeline does not provide for joint genotyping of variants across samples), and then filtered the variants for Fisher strand bias <30 and QD (quality/depth) score >2.0.

**Genotype quality control (QC) and imputation**

In addition to the VCF files generated from the WGS or snRNA-seq data, we obtained available microarray or next-generation sequencing-based variant callsets for each cohort (**fig. S8A**). For each individual dataset, we first lifted hg18 or hg19 datasets over to hg38, performed strand-flipping and excluded ambiguous SNPs for array-based datasets using snpflip (https://github.com/biocore-ntnu/snpflip), and fixed alleles to the hg38 reference genome using plink2 (*134*). SNPs with a Hardy-Weinberg equilibrium $<1\times10^{-6}$ and missing in >5% of samples were removed, and all individuals were confirmed to not have missing genotypes (>5%) or high or low rates of heterozygosity (>|3SD| from the cohort mean).

For genotype imputation, we aimed to maximize the number of final variants available for each sample without having any missing data. To do this, we first merged genotype data for samples across most cohorts, while genotype data for the snRNA-seq-based variants and genotypes from Velmeshev. (variants restricted to coding regions), MultiomeBrain (variants derived from pre-processed VCF files), and ROSMAP Affymetrix array (lifted over from hg18) datasets were imputed separately. SNPs with <90% call rate and minor allele frequency (MAF) <0.05 across all samples were removed, leaving a total of 114,000 SNPs for imputation in the combined cohort dataset. Genotypes were imputed on the NIH TOPMed Imputation Server (Mimimac 4) using default parameters, Eagle 2.4 phasing, and an $R^2$ threshold of 0.3 (*135*) (**fig. S8A**). Post-imputation, we merged the imputed snRNA-seq, MultiomeBrain, Velmeshev, and ROSMAP Affymetrix genotypes with the rest of the cohorts, and filtered out imputed variants with MAF<0.05 and those not present in all samples. Our final imputed genotype dataset contains 1.95 million SNPs for 383 samples, comparable to previous population-scale QTL analyses (*4*).

VCF files for imputed genotypes of all PsychENCODE Consortium samples (excluding ROSMAP samples, select WGS samples, and snMultiome samples generated in this study), including callsets that both include and exclude samples with snRNA-seq-derived genotypes, are available with approved access within the PsychENCODE Consortium data portal on Synapse (see above section for “Raw dataset availability”). Raw sequencing files for ROSMAP samples are available with separate approved access through the AMP-AD Knowledge Portal on Synapse.

**Genotype PCA analysis and QC**

We applied Peddy software (*136*) on the final set of imputed genotypes to calculate genotype PCAs for each sample and to predict genetic ancestry based on 1,000 Genomes samples (**fig. S8A**). As these predictions showed strong concordance with self-reported ancestry for samples with available meta-data, we used the genetic ancestry as a covariate in a downstream analysis (**fig. S1E**). Additionally, for samples with both genotype and non-multiplexed snRNA-seq data, we performed a QC check for sample swaps and inconsistencies with sample meta-data. In particular, we used the "Risk Assessment" program from the privaseq3 toolkit (*137*) to match the genotypes derived from RNA-seq samples with those from WGS or SNP microarray data. All of the snRNA-seq samples in our final cohort showed the highest matching scores with their respective WGS/array genotypes; samples with potential mismatches were excluded from downstream analysis. We also ensured consistency for biological sex (based on coverage on the X and Y chromosome) and predicted ancestry (based on the Peddy results) among the snRNA-seq data, genomic data, and sample metadata across all samples.

VCF files for imputed genotypes of all PsychENCODE Consortium-derived samples (excluding AMP-AD samples and select callsets outside the scope of the consortium) are available for researchers with approved access within the PsychENCODE Consortium data portal. Two VCF files are available, one with and one excluding samples with snRNA-Seq derived genotypes. Imputed genotypes for the AMP-AD samples used in this study are available on the AD Knowledge portal.

**Rare variant and structural variant (SV) annotation and analysis**

In addition to the above work to generate a robust set of common single-nucleotide variants (SNVs) for downstream analysis, we also assessed rare deleterious SNVs and SVs for their roles in disrupting gene regulation in a cell-type-specific manner (**fig. S8B**). To identify rare SNVs and small insertions/deletions (indels) from 82 samples with exome or WGS data (**fig. S1D**), we annotated GATK-derived variant calls using the Annovar v.06-2020 package (*138*). We specifically selected variants annotated for disrupting exonic, splice-site, and promoter regions (or those within 1 kbp of the transcriptional start site [TSS]) that are present in <1% of the GnomAD v.3.0 general population panel (*139*) (**fig. S8B**). Additionally, likely damaging missense and promoter variants were selected for downstream analysis by filtering for a Combined Annotation Dependent Depletion Phred-like score >10.0 (*140*). Overall, we identified an average of 13,503 rare variants per individual, including 84 rare loss-of-function (LOF), 455 rare deleterious missense, and 16 rare splicing variants per individual.

We also used the genotyping software PanGenie *(141)* to identify both rare and common SVs in 48 samples with available WGS fastqs. PanGenie is a kmer-based genome inference algorithm that uses a high-quality phased SV panel from 64 Human Genome Structural Variation Consortium/1000 Genomes samples (*142*) to genotype SVs in short-read data. After genotyping each sample, we annotated SVs >50 bp in length for gene overlap and population frequency within the reference panel (**fig. S8B**). Using this method, we identified an average of 18,669 genomic deletions and 26,579 genomic insertions with any allele frequency per sample.

Finally, we used the rare SNVs identified in these samples to assess gene regulatory "knockouts" in the context of the cell-type-specific GRNs. Briefly, we first overlapped rare LOF coding variants in each sample with lists of transcription factors (TFs) identified in each cell-type-specific regulon, and found 112 TFs with rare variants in at least one of the 82 samples with rare variant data. For each disrupted cell-type-specific regulon, we then compared the expression values of each downstream target gene in annotated cells among samples with and without the mutation, and calculated basic Z-scores comparing expression across the groups of cells. A broad threshold of $|Z|>2.5$ was used to define genes with putative outlier expression (*143*); we found that 79/103 regulons tested (77%) had downstream genes with average absolute Z-scores of >2.5 (**fig. S63**), indicating possible global disruption of downstream genes due to LOF variants in the TF.

VCF files for the rare SNVs and SVs of all PsychENCODE Consortium samples (excluding AMP-AD samples and select WGS callsets outside the scope of the consortium) will be available for researchers with approved access within the PsychENCODE Consortium data portal. AMP-AD variant datasets are available on the AD Knowledge portal.

#### 1.7 ***Cell-type Fractions -*** Calculation of cell-type fractions from snRNA-seq data

**Main manuscript reference:** First supplementary reference in the third paragraph of "Constructing a single-cell genomic resource for 388 individuals."

Single-cell fraction statistics were calculated based on the harmonized cell annotation scheme. The distribution of the raw cell fraction of each cell type in each individual is shown in **fig. S13**, and the distributions for each individual in one example cohort (SZBDMulti-seq) are shown in **fig. S14.** We compared cell-type fractions between ASD and control samples (using samples from the DevBrain, Velmeshev, and UCLA-ASD cohorts), schizophrenia and control samples (using samples from the SZBDMulti-seq, CMC, and MultiomeBrain cohorts), and bipolar disorder and control samples (using samples from the SZBDMulti-seq and MultiomeBrain cohorts). Cell-type fractions between disease and control samples were compared using Welch's t-test, where outliers with >1.5 interquartile range (IQR) were removed. Only cell types with a median fraction per sample larger than 0.5% were compared. The nominal p-value was corrected by Benjamini–Hochberg false discovery rate (FDR), and adjusted p-values <0.05 were considered as significant. Computed cell fractions for each individual and cell type are described in the following file on the brainSCOPE portal:

**File**: cell_type_fraction_count_with_meta.csv (**data S4**): This file contains normalized cell fractions calculated for each individual from the snRNA-seq data. Columns indicate cell type, cell counts, cell fraction, and relevant meta-data for each individual.

#### 1.8 ***Cell-type Fractions -*** Deconvolution of cell fractions for PsychENCODE Consortium bulk-RNA-seq data

In addition to snRNA-seq data, we deconvolved cell fractions based on bulk-RNA-seq datasets previously published by the PsychENCODE Consortium *(4)* against snRNA-seq samples in the current dataset. Specifically, we collected the CMC single-cell cell-type annotations and raw count matrices alongside the PsychENCODE Consortium bulk-RNA-seq raw count matrix. We used BisqueRNA to infer cell-type fractions of the bulk-RNA-seq data (*144*). The single-cell raw counts were first log normalized, while the bulk-seq raw counts were quantile normalized, before being input into BisqueRNA.

As with the single-cell-derived fractions, these results are described in a file on the brainSCOPE portal:

**File:** bisque.PEC_CMC.rev.txt (**data S5**): This file contains normalized cell fractions calculated from deconvolved bulk-RNA-seq data.
**File:** cell_fraction_corr.txt (**data S6**): This file lists correlations between cell fractions from single-cell and bulk deconvolution datasets for each cell type.

#### 1.9 ***DE Analysis*** - Differentially Expressed (DE) Genes for disease traits

**Main manuscript reference:** Second supplementary reference in the third paragraph of "Constructing a single-cell genomic resource for 388 individuals."

This section only includes method details for disease traits DE genes. For DE analysis on control aging and schizophrenia aging, see more details in ***7.3 Aging DE*** and supporting experiments in **fig. S19**.

We calculated DE genes for each cell type regarding different diseases. Cell-type-specific pseudobulk gene expression profiles were first generated from the snRNA-seq data by calculating the sum of the raw gene counts per cell type per individual.

**Filtering:** Counts per million (CPM) normalization was used to filter out lowly expressed genes. Genes were removed if their CPM normalized expressions were >0.5 in <30% of the samples. Genes were also removed if the raw count sums of all individuals were <20. Additionally, within each cell type, individuals with <50 cells detected were removed from the calculation. After filtering, cell types containing less than 16 samples were also excluded from the differential expression calculation.

**Deseq2**: For each cell type, differential expression analysis was performed with the raw counts using the standard pipeline for the Deseq2 likelihood ratio test (*23*), with age, gender, genotype ancestry, PMI, average UMI per cell, and disease status as covariates. Contrasts were made between disease and healthy status. Multiple testing corrections were performed, and genes with an adjusted p-value<0.05 were defined as differentially expressed between contrast conditions.

We also compared the DE gene list calculated by DESeq2 with DE genes calculated by Dreamlet *(145)* in **figs. S16-S18**. We plotted the $\log_2$ fold change from these two sets for each cell type using a scatter plot. We also calculated the Pearson correlation of the $\log_2$ fold change.

The log fold changes for DE genes with a p-value less than 0.05 were found to be in concordance between the two methods.

We also describe the **Dreamlet** method here: DE analysis was performed using the Dreamlet package that uses linear mixed models with raw count matrices as inputs. Covariates selected for each cohort were the following: age, gender, log UMI count, and PMI. Obtained p-values from Dreamlet were further corrected for FDR within each cell group independently using p.adjust from the R stats package and applying Benjamini–Hochberg correction.

The input files for the analysis are available in **data S3**, and we provide the pseudobulk expression matrices for each cell type on the brainSCOPE portal. DE genes for disease traits are available in several files on the brainSCOPE portal. Each file contains the gene name, average expression, $\log_2$ fold change, standard error, test statistic, p-value, and FDR-corrected p-value among individuals with the disease:

**File:** ASD_DEGcombined.csv (**data S7**): Sets of DE genes between control individuals and individuals with ASD for each cell type. (**Data S7** contains all significant DE genes (p<0.05, DESeq2 likelihood ratio test); full results available on the brainSCOPE portal.)
**File:** Bipolar_DEGcombined.csv (**data S7**): Sets of DE genes between control individuals and individuals with bipolar disorder for each cell type. (**Data S7** contains all significant DE genes (p<0.05, DESeq2 likelihood ratio test); full results available on the brainSCOPE portal.)
**File:** Schizophrenia_DEGcombined.csv (**data S7**): Sets of DE genes between control individuals and individuals with schizophrenia for each cell type. (Data S7 contains all significant DE genes (p<0.05, DESeq2 likelihood ratio test); full results available on the brainSCOPE portal.)
**File:** [celltype].expr.bed.gz: Pseudo-bulk snRNA-Seq expression matrices for 24 cell types, listing logCPM normalized expression values for individuals who pass quality control for each cell type.
**File:** *_ASD_table.csv: Different version of DEGs between control individuals and individuals with ASD for 20 cell types, used for LNCTP model comparisons.
**File:** *_Bipolar_disorder_table.csv: Different version of DEGs between control individuals and individuals with bipolar disorder for 19 cell types, used for LNCTP model comparisons.
**File:** *_Schizophrenia_table.csv: Different version of DEGs between control individuals and individuals with schizophrenia for 21 cell types, used for LNCTP model comparisons.

#### **1.10** ***Trajectory Analysis*** - Identifying DE genes over cell types by pseudotime

**Main manuscript reference:** First supplementary reference in the last paragraph of “Constructing a single-cell genomic resource for 388 individuals.”

##### Overall approach and differential expression analysis

We implemented the following approach to identify genes that consistently vary in a continuous fashion across the dimension of cortical depth. First, we generated embeddings of the excitatory IT cells (consisting of L2/3 IT, L4 IT, L5 IT, and L6 IT neurons) in lower-dimensional representations, and then constructed smooth pseudotime trajectories in this lower-dimensional space. Genes that demonstrated variation in a statistically significant fashion over the trajectory were selected for downstream analysis (Wald test, FDR $\leqslant 0.05$). We analyzed each study cohort separately for such genes, filtering out studies with more than one trajectory (thus resulting in trajectories that may include only two cell types) or that were noisy, and subsequently identified those genes that were also found in the trajectories for all remaining cohorts.

The input datasets for this analysis were the annotated h5ad files from the harmonized snRNA-seq processing. Additionally, during the process of identifying genes that varied smoothly across the trajectories, we removed genes that were differentially expressed in only one cell type. To generate a list of DE genes for each cell type, we separately ran the h5ad files through the following processing steps using the Seurat package (*146*, *147*):

1. Read the h5ad file into R and transpose the raw count matrix; convert this transpose first into .csr format and then into dgCMatrix format (in order to generate a Seurat object); and create a Seurat object with the raw count matrix and the metadata from the h5ad file.
2. Run the function *SCTransform* (*148*) to normalize the matrix, with the number of variable features set to 10,000.
3. Subset the L2/3 IT, L4 IT, L5 IT, and L6 IT neurons.
4. Run PCA and UMAP analyses and then subset out the variable features.
5. Run Seurat's *FindMarkers* function (default settings with a Wilcoxon rank sum test) in a "one versus all" fashion for the four cell types.

In the following section, we provide further details of the pipeline.

##### Preprocessing and QC

After reading the data from each study cohort into a Seurat object in the manner described above, the Seurat object was subset to (a) include only L2/3 IT, L4 IT, L5 IT, and L6 IT neurons; (b) keep only control samples; and (c) keep only samples ≥20 years of age. We then filtered out low-quality cells with a total number of ≤1,000 UMI counts and ≤500 expressed genes. Importantly, we retained only protein-coding genes for downstream analyses; while our method does not inherently vary based on this filtering step, we chose to do so to simply focus on the protein-coding component. Subsequently, the normalization step used the *SCTransform* function, where variable features included were those with a residual variance >1.3. After running PCA, the program Harmony (*127*) performed batch-correction with a maximum number of 50 iterations. A UMAP embedding was generated with the top 30 principal components (PCs). Finally, the Seurat object was subset to include only the variable features.

##### Trajectory analysis

We used the program *Slingshot* (*26*) to generate pseudotime trajectories. We input the UMAP coordinates as the lower-dimensional representation used in *Slingshot*, and set the starting cluster for the trajectories to be the cluster associated with L2/3 IT cell type. Note that we did not use any form of clustering to identify cell-type clusters in the UMAP representation; rather, we fed *Slingshot* the labels inferred in our annotation pipeline. The *Slingshot* function built a graph based on the data, with the clusters as nodes. A minimum spanning tree was computed for the graph, and the tree was smoothed via principal curve analysis. Pseudotimes were determined through orthogonal projections of the points onto the smoothed trajectories.

##### Differential expression along the trajectories

For the assessment of differential expression of genes along the pseudotime trajectories, we used the package *tradeSeq* (*27*). The *tradeSeq* package offers the function *fitGAM,* which fits a generalized additive model (GAM) using the count matrix and *Slingshot* trajectories. Specifically, the function considers each gene and smoothes log counts of the gene expression along the pseudotime trajectories using cubic splines. The GAM allows us to account for several covariates in the process: we included for each donor their biological sex, age, sample PMI, total cell counts, and cell-type proportions. The last two covariates were included to remove any bias associated with particular samples not capturing sufficient total numbers of cells or having certain cell types be underrepresented. The covariates were included in the form of a model matrix in R.

A suite of statistical tests may be computed on these smooth functions to determine which genes are differentially expressed along a certain lineage, between two lineages, or across certain conditions. Our analysis made use of *tradeSeq*'s associationTest function, which evaluates the statistical significance of the variation of a gene across the trajectory by testing whether the spline fit parameters vary significantly across the trajectory. We applied a filter on the FDR of 0.05 on the gene set from each cohort based on the Wald test implemented by the association Test function.

##### Post-processing

For post-processing, we worked on a set of processed data cohorts that underwent two stages of selection. In the first stage, we selected cohorts that had a single trajectory. In the second, we further subset the cohorts, after visual inspection of the trajectories, to only include ones that showed smooth trajectories that were less impacted by the occurrence of clusters where cells from multiple cell types were intermixed. Two data cohorts demonstrated single trajectories but had considerable intermixing of IT neuron types, so we removed these cohorts from the final overlap across cohorts. The final cohorts included Velmeshev, SZBDMulti-seq, CMC, IsoHuB, and MultiomeBrain. We identified a common set of genes (76 for the filtered cohorts, 5 TFs in both sets) that varied significantly along the trajectories in each cohort and that consistently occurred in the results for all five cohorts considered. We note that including the two cohorts with noisy single trajectories reduced the number of overlapping genes by 10 (from 76 to 66). The list of common genes across trajectories is available below as **table S5**, and on the brainSCOPE portal as a file (the last genes in the file are ten noise-sensitive genes).

##### Additional results from the IT neuron trajectory analysis

We conducted downstream analyses of the 76 genes identified as significant in the IT neuron trajectory analysis, connecting them with the results in other sections of this paper as well as published results. The goal was to identify biological functions that might be enriched in the gene sets. For example, there were seven ribosomal proteins in our significant gene set.

Upon inspection of the cell expression averages across the pseudotime trajectories (**fig. S21**), we found that the patterns across the spatial trajectory for these seven genes were remarkably similar.

Next, we identified genes in our set that overlapped with the GRNs. Five TFs (*TFEC*, *RUNX2*, *MAF*, *PROX1*, *ERG*) occurred in the IT neuron cell-type-specific GRNs. *ERG* TF showed up only in the L4 IT cell-type-specific GRN.

We subsequently searched for an overlap with other relevant categories of genes:

(a) **Ligands and receptors in the cell-to-cell communication network:** The genes *SEMA6A* and *PENK* overlapped with the ligands, while *TACR3* and *RXFP2* overlapped with the receptors.

(b) **DE genes for schizophrenia, bipolar disorder, ASD, and aging:** We found overlaps of the significant (p_adjusted < 0.05, Wald test) genes with the DE genes set (DEGs) for schizophrenia, ASD, and aging.

**Schizophrenia**

Subclass = Pvalb, Total # of DEGs = 115, Overlapping DEGs = *CARD18*
Subclass = L5.IT, Total # of DEGs = 396, Overlapping DEGs = *NPNT, LRIG3*
Subclass = L6.IT, Total # of DEGs = 176, Overlapping DEGs = *LONRF3*
Subclass = OPC, Total # of DEGs = 20, Overlapping DEGs = *RPL32, ENPP1*
Subclass = Astro, Total # of DEGs = 38, Overlapping DEGs = *TMEM132C*
Subclass = L2.3.IT, Total # of DEGs = 166, Overlapping DEGs = *CRYAB, CNDP1*
Subclass = Vip, Total # of DEGs = 22, Overlapping DEGs = *CCDC178*

**Bipolar Disorder**

No overlapping genes were found.

**ASD**

Subclass = Vip, Total # of DEGs = 76, Overlapping DEGs = *CRYAB*
Subclass = L4.IT, Total # of DEGs = 185, Overlapping DEGs = *RPS23*
Subclass = Sncg, Total # of DEGs = 195, Overlapping DEGs = *MAF*
Subclass = L2.3.IT, Total # of DEGs = 828, Overlapping DEGs = *EEF1A1*
Subclass = Micro, Total # of DEGs = 267, Overlapping DEGs = *APOE, ANKRD62, ARHGAP6*

**Aging**

Subclass = Sst, Total # of DEGs = 73, Overlapping DEGs = *ADAMTS6, EYA4, SCN7A*
Subclass = L2.3.IT, Total # of DEGs = 572, Overlapping DEGs = *LONRF3, RYR3, RUNX2*
Subclass = Vip, Total # of DEGs = 289, Overlapping DEGs = *CCN2, VRK2, CCDC178*
Subclass = L6.IT, Total # of DEGs = 269, Overlapping DEGs = *CRYAB, CA8, RASGEF1B*
Subclass = L4.IT, Total # of DEGs = 145, Overlapping DEGs = *SEMA6A*

(c) **Functional categories of genes as identified by gene ontology (GO)** (*149*, *150*): We further explored the functional categories of the significant genes, to identify any emerging patterns. We found many of the genes were associated with cell-type differentiation, neurogenesis, and gliogenesis (**data S8**).

Finally, given the patterning of the cortical layers during development, we considered the expression of the significant genes as a function of developmental stage as quantified by the bulk RNA-seq assays from the BrainSpan study (*151*). As shown in **fig**. **S22**, several genes in the list showed considerable variation across the developmental trajectory, with some decreasing consistently from the earliest stages onwards (*EEF1A1* in **fig. S22A**), some decreasing around the time of birth (several of the ribosomal proteins in **fig. S22A**), and some increasing dramatically around the time of birth (*APOE*, *CRYAB*, *CXCL14,* and *IGFBP7* in **fig. S22C**).

The list of significant genes (FDR<0.05, Wald test, overlapped across five cohorts) is also available on the brainSCOPE portal:

**File:** IT_neuron_trajectory_sig_gene_set.tsv (**table S5**): Overlap set of genes (HGNC symbols) across five cohorts, identified as significantly varying (FDR < 0.05, overlapped across five cohorts, Wald test for each cohort) across the IT neuron trajectories in each cohort separately.

### 2 Determining regulatory elements for cell types from snATAC-seq

#### 2.1 ***snATAC-seq Processing***

**Main manuscript reference:** Second supplementary reference in the first paragraph of "Constructing a single-cell genomic resource for 388 individuals"; first and second supplementary references in the first paragraph, and first reference in the last paragraph of "Determining regulatory elements for cell types from snATAC-seq."

The snATAC-seq data processing and analysis workflow is largely based on existing and published approaches, and heavily relies on the Signac package in R (*152*). We ran this pipeline on snATAC-seq datasets from the UCLA-ASD cohort, as well as snMultiome datasets from the MultiomeBrain cohort and snMultiome datasets generated specifically for this paper. In general, this pipeline consists of the following steps:

1. Per-sample QC and filtering
2. Per-sample dimensionality reduction and preliminary analysis
3. Data aggregation and further QC
4. Batch effect removal
5. Peak calling
6. Distal-peak-to-promoter linkage
7. Motif enrichment analysis

These steps are largely derived from two perspectives: per-sample analysis and data integration. Here, the emphasis is to ensure that the processed snATAC-seq data are of high enough quality for further downstream analysis.

**Step 1. Per-sample QC and filtering**

To be consistent with the snRNA-seq data processing, before any filtering using snATAC-seq data, we first removed the barcodes with the high possibility of being doublets or having poor quality on the snRNA-seq side, using the list of barcodes after Pegasus processing with the subset() function in Signac v.1.5.0 (*152*). Then, for each sample, we created a chromatin assay object using the CreateChromatinAssay function with other metadata, including fragments, n_counts, and n_frags, properly added. Then, we filtered each sample to keep only cells with a sequencing depth of >1,000 and a TSS enrichment of ≥2. We set the cutoff threshold to 2 due to the fact that post-mortem brain cells tend to have lower quality, especially in terms of the TSS enrichment. Setting a threshold too large (for instance, 4) may result in a dramatic loss of sample size to study.

**Step 2. Per-sample dimensionality reduction and preliminary analysis**

After initial curation of each sample, we conducted dimensionality reduction analysis in a number of ways. Using the snATAC-seq data only, we first performed term frequency inverse document frequency (TFIDF) and latent semantic indexing with 30 dimensions using the RunTFIDF and RunSVD functions in Signac. Next, including snRNA-seq information, we attempted to create a joint embedding with the FindMultiModalNeighbors function. With the two sets of embeddings, we performed UMAP visualization, resulting in two sets of figures, which allowed us to visually inspect the quality of each sample and decide on a sample level to keep. We further overlaid the TSS enrichment, number of fragments, and number of genes per cell on these UMAP visualizations as an additional sanity check procedure to ensure the cell clusters were reasonably formed. The resulting sample objects were stored individually in an RDS file.

**Step 3. Data aggregation and further QC**

We first loaded and merged each sample object into one Signac multiome object using the merge function. After merging all samples, we calculated basic statistics of the combined dataset, such as the mean TSS enrichment and mean snATAC-seq sequencing depth. Here, another round of dimensionality reduction using both methods (with and without snRNA-seq data) was performed, and the results were visualized and compared. Any outlier cells or clusters were removed manually after this step.

**Step 4. Batch effect removal**

With the aggregated dataset mostly clean, we aligned all samples and removed as much batch effect as possible using Harmony with 100 iterations. After running Harmony, we had an embedding with the samples aligned (*127*). We then conducted another UMAP visualization, colored by sample to confirm that the samples were indeed aligned.

**Step 5. Peak calling**

After the dataset was filtered and curated, we used the CallPeaks function in the Signac package to call the cell-type-specific peaks. We used the default parameters and Macs2 2.2.6 *(153)*. In this step, instead of using subclass labels, we chose to split the data into the following generalized cell types: excitatory neurons, inhibitory neurons, astrocytes, endothelial cells, microglia, oligodendrocytes, oligodendrocyte precursor cells, and immune cells. We received from this call a list of merged peaks with cell-type-specific annotations. From there, we separated cell-type-specific peaks using the annotation function, which gave us a list of cell-type-specific anchored peaks for each cell type. Using hg38 annotations from the UCSC Genome Browser, we further split the peak set of each cell type into four categories: promoter, intronic, exonic, and distal. If a peak intersected with the promoter region of a gene, we categorized it as promoter, and similarly for the exonic and intronic regions. If a peak did not have any overlap with the above three categories, we classified it as a distal peak.

**Step 6. Distal-peak-to-gene linkage**

We generated a peak-to-gene linkage graph employing the addPeak2GeneLinks function in ArchR, utilizing two distinct parameter sets. The first configuration involved a relaxed approach, setting a maximum distance of $\pm$250,000 base pairs (=250 kbp) and a Pearson's correlation cutoff of 0.1. In the second, more lenient configuration, we extended the maximum distance to $\pm$500 kbp and adjusted the Pearson's correlation cutoff to 0.45. The latter, more lenient, distance of 500 kbp was used in the generation of the cell-type-specific gene regulatory networks (GRNs; see supplementary section 5 below). All other parameters were maintained at their default values. Subsequently, we imported the motif-to-peak graph from JASPAR2020 motif annotation using the getPeakAnnotation function in ArchR. Combining these two graphs, we constructed a motif-peak-gene linkage graph, establishing an any-to-any relationship. Specifically, we established an edge from a motif to a gene if there was a concurrent edge from the motif to a peak and from that peak to the designated gene.

**Step 7. Motif enrichment and TF footprinting analysis**

Given a set of called peaks and the curated dataset, we performed motif enrichment analysis using ArchR (*38*). First, we added the peak set to the dataset and generated the corresponding count matrix, using the addPeakSet and addPeakMatrix functions with default parameters. Then, we added the differentially accessed marker peaks using getMarkerFeature functions, with the cell type being the general cell type described in the peak calling step, the bias being TSS enrichment and $\log_{10}$(nFrags), and the testing method being the Wilcoxon test. We used the JASPAR2020 *(154)* database for motif annotations, and calculated the motif enrichment scores in terms of normalized $-\log_{10}(P_{adj})$ using the found peaks and cutoffs of FDR $\leq$0.1 and $\log_2$FC $\geq$1. A heatmap was drawn from this result with a manually selected set of motifs. Furthermore, we selected a set of TF motifs to calculate motif footprints using ArchR's getPositions, addGroupCoverages, and getFootprints functions, according to default parameters and the general cell types described above.

**Step 8. Processed data availability**

The identified snATAC-seq peaks, signal tracks based on snATAC-Seq and snMultiome datasets, and associated analyses are available on the brainSCOPE portal as follows:

**File:** [celltype].PeakCalls.bed: These BED files consist of a set of seven cell-type-specific ATAC-seq peaks identified in our analysis of snMultiome and snATAC-seq data, along with a file showing the union of peaks across all cell types (All.celltypes.Union.PeakCalls.bed).
**File:** [celltype].bigwig: These bigWig files consist of seven cell-type-specific signal tracks for chromatin accessibility, as well as a track for merged open chromatin signal across all cell types. Three sets of signal tracks are available for each snATAC-Seq and snMultiome cohort assessed (Girgenti-snMultiome, MultiomeBrain, and UCLA-ASD).
**File:** *tf_enrich.csv (**data S11**): These files list the enrichment Z-scores of scCREs in proximal and distal regions for 634 TF binding sites.

#### 2.2 ***LDSC*** - Methods for Linkage Disequilibrium Score Regression (LDSC) enrichment of candidate regulatory elements

**Main manuscript reference:** First supplementary reference in the third paragraph of "Determining regulatory elements for cell types from snATAC-seq."

We downloaded 11,908 genome-wide association study (GWAS) summary statistics for 4,585 traits, including 3,582 for males, 3,741 for females, and 4,585 for both sexes, from the UK Biobank (http://www.nealelab.is/uk-biobank/). We also obtained 17 summary statistics from PGC (Psychiatric Genomics Consortium, https://pgc.unc.edu/for-researchers/download-results/) and six summary statistics from PASS *(155)*. We parsed these statistics into a format that could be recognized by the LDSC pipeline (*156*). In order to control the statistical power of the summary statistics, we filtered the summary statistics with the threshold of >5,000 samples. We divided all traits into 19 biological system-based categories derived from the Human Phenotype Ontology (*157*): behavioral, blood/blood-forming tissues, cardiovascular, constitutional symptom, digestive system, ear, endocrine system, eye, genitourinary system, growth, head/neck, immune system, integument, musculoskeletal, metabolism/homeostasis, neoplasm, nervous system, and respiratory. We defined 'behavioral' and 'nervous system' traits as brain-related traits, and all other categories as non-brain-related traits. We then ran the LDSC pipeline against nine main types of annotations: UCLA-ASD snATAC-seq data for Astro, Endo, Exc, Inh, Micro, OPC, Oligo, adult brain cis-regulatory elements (b-cCREs), and ENCODE adult cCREs v4 (*31*). LDSC enrichment results for all brain-related triats are shown in **fig. S26**. To ensure genome version consistency, all summary statistics and annotations were lifted over to GRCh37. Linkage disequilibrium (LD) scores used in the pipeline were derived from (*158*).

All LDSC results for the nine tested sets of regulatory elements and relevant metadata files are available on the brainSCOPE portal:

**File:** ukbb-all-traits-pval.csv (**data S9**): This file contains the -log(p-value) of LDSC enrichment for UKBiobank GWAS summary statistics in snATAC-seq peaks of seven cell types (Astro, Endo, Exc, Inh, Micro, OPC, Oligo) as well as adult b-cCREs regions. The index is trait ID. The column 'UK Biobank trait' refers to the trait name/description in UKBB. The column 'HPO phenotype category' refers to the phenotype ontology category. The column 'brain' refers to whether the trait is brain-related.

**File:** PGC_PASS-all-traits-pval.csv (**data S10**): This file contains the -log(p-value) of LDSC enrichment for PGC and PASS GWAS summary statistics in snATAC-seq peaks of seven cell types (Astro, Endo, Exc, Inh, Micro, OPC, Oligo) as well as adult b-cCREs regions. The column ‘brain’ refers to whether the trait is brain-related.
**File:** IDtoTraitName.txt: This file contains a matrix to convert trait ID into the full UKBiobank trait name.
**File:** cluster.trait.txt: This file contains a matrix that assigns each UK BioBank trait to a Human Phenotype Ontology category, such as behavioral, eye, or respiratory. The column 'HPO phenotype category' is used to distinguish brain- and non-brain-related traits. We defined the “behavioral” and “nervous system” categories as brain-related traits.

#### 2.3 ***STARR-seq -*** Validation of putative enhancers in neural progenitor cells

**Main manuscript reference:**Third supplementary reference in the first paragraph of “Determining regulatory elements for cell types from snATAC-seq.”

We performed two rounds of capture STARR sequencing (CapSTARR-seq) in primary human neural progenitor cells (phNPCs) isolated from the fetal cortex (*159*), each containing two biological replicates. Detailed protocols for the CapSTARR-seq assays are available in the PsychENCODE Consortium publication by Gaynor and colleagues *(32)*. This approach implemented a hybridization-based capture method to isolate specific candidate regions to test for enhancer activity through STARR-seq. Our first CapSTARR-seq experiment interrogated 22,400 candidate regions selected based on bulk ATAC-seq, DNase-seq, and ChIP-seq data from the PFC. Our second experiment interrogated 56,215 candidate regions selected based on bulk ATAC-seq data from the PFC, ChIP-seq and DNase seq data from NPCs, developmental data, eQTL and transcription-wide association study data from the fetal brain, and GWAS data (*4*, *100*, *160*–*162*). Two replicates were performed for each experiment. Combined, these panels identified 8,148 regions with enhancer activity in at least one replicate and 6,612 regions with enhancer activity across both replicates *(32)*. We identified 2,288 predicted target genes for our enhancer regions, many of which were implicated in neuronal pathways.
A list of validated enhancers from the STARR-seq experiments is available on the brainSCOPE portal in the following file:

**File:** starrseq_enhancers_merged.bed: BED file that contains the merged set of validated STARRseq enhancers.

#### 2.4 ***b-cCREs*** - Identification of brain-specific cCREs

**Main manuscript reference:** First supplementary reference in the second paragraph of “Determining regulatory elements for cell types from snATAC-seq.”

To curate candidate cis-regulatory elements specific to the brain from adult samples, we followed a multi-step approach. First, we identified all non-disease brain ENCODE DNase-seq experiments (96 adult samples). From each DNase-seq experiment, we selected V4 cCREs with a Z-score > 1.64. V4 cCREs were obtained directly from ENCODE, accessible at screen.encodeproject.org (*31*). Subsequently, we removed cCREs with limited experimental support and retained only those elements that were supported by at least five experiments. This filtering resulted in a collection of 253,638 adult b-cCREs. We selected this threshold to maximize phyloP conservation across the retained elements.

A list of validated enhancers from the STARR-seq experiments is available on the brainSCOPE portal in the following file:

**File:** adult_bcCREs.bed: BED file containing the list of identified b-cCREs.

### 3 Measuring transcriptome and epigenome variation across the cohort at the single-cell level

#### 3.1 ***Variance Partition*** - Partitioning transcriptome variation among cell types, individuals, and regions

**Main manuscript reference:** First supplementary reference in the first paragraph, and first supplementary reference of the last paragraph of “Measuring transcriptome and epigenome variation across the cohort at the single-cell level.”

In order to assess biological sources of expression variation, we leveraged the tool variancePartition (*163*) and applied it to our population-scale snRNA-seq data. We filtered the total set of genes down to ~13,000 genes based on minimum quality requirements such as sufficient expression. We used pseudobulk expression (similar to expression processing for eQTLs) of each gene for a given individual and cell type as a sample for this analysis. Together, all samples across individuals and cell types were run through variancePartition to assess the percent variation associated with cell type, individual, and residual. In addition, we found the total variance across all samples for a given gene.

With the same strategy described above, we also considered the partitioning of variation among different brain regions, cell types, and donors. Six individuals with snRNA-seq data from three brain regions (PFC, parietal lobule, and occipital lobule) from the Gandal-UCLA cohort *(164)* were processed using our snRNA-seq pipeline and hybrid annotation scheme, as described above (see section “snRNA-seq Processing”). Average gene expression was calculated based on the single-cell gene expression matrices for each individual, and we calculated the total variation and percent variation for each gene using variancePartition.

Next, we compiled a list of all GO annotations for each gene from geneontology.org *(150)*, with a specific focus on curating gene sets/families associated with neurotransmitters (such as serotonin). Since each gene set/family consists of a list of associated genes, we took the mean total variance as well as the mean partitioned variance (by cell type, individual, and residual) to provide a breakdown of the gene sets and families per their associated variation by percent. We compared various neurotransmitter families and other gene sets in regards to their partitioned variance.

To plot the UMAPs to show the serotonin receptor genes *HTR2A* and *HTR2C*, we used the SZBDMulti-seq dataset and drew a heatmap of the gene expression across cell types (**fig. S31C**). The annotations are consistent with the UMAP labels in **Fig. 1B**.

In addition to the GO annotations, we investigated the variation of drug target-related genes. We selected 280 neuro-related drug target genes from the CLUE database and calculated their inter-individual and inter-cell type variation (*42*). We further highlight some of these examples in **fig. S32**.

Associated output and metadata files for the variancePartition analysis are available on the brainSCOPE portal, as follows:

**File:** variancePartition_output.csv (**data S12**): This file contains the summary of the variancePartition results for all genes (~13k genes that meet the minimum QC requirements). Percent variations derived from various factors are given, as well as the total absolute variation of each gene.

**File:** variance-partition-092623.csv (**data S13**): This file contains a summary of the variancePartition results for all genes. Percent variations derived from various factors (individuals, cell types, brain regions) are given, as well as the total absolute variation of each gene.

**File:** gene_map_df_02252023.txt (**data S14**): This file contains the data frame for mapping genes to their respective gene families using GO annotations.

#### 3.2 ***Variance Partition*** - Quantifying epigenomic variation across cell types

To obtain population-scale variation on open chromatin regions in a cell-type-specific manner, we leveraged 628 bulk ATAC-seq signals previously generated by the PsychENCODE Consortium *(4)*. We used cell-type-specific open chromatin regions from our snATAC-seq dataset as anchors and found the variation of the bulk ATAC signals within those regions. Cell-type specificity was determined by using the merged set of anchors and then determining how many of the seven cell types intersected with the merged anchors.

#### 3.3 ***Conservation*** - Cross-species sequence conservation

**Main manuscript reference:** First supplementary reference in the third paragraph, and second supplementary reference of the last paragraph of “Measuring transcriptome and epigenome variation across the cohort at the single-cell level.”

We used phastCons annotations as a measure of cross-species sequence conservation for all analyses (*165*). PhastCons citations are available for download at http://hgdownload.cse.ucsc.edu/goldenpath/hg38/phastCons100way/.
Specifically, we used bigwigaverageoverbed to calculate phastCons scores across various annotations. Protein-coding gene annotations were derived from GENCODE release 43 of the human genome. In addition to coding gene peak regions, we computed phastCons scores for b-cCRE regions and cell-type-specific snATAC-seq open chromatin regions.

Several output files associated with the conservation analysis are available on the brainSCOPE portal, as follows:

**File:** gene_coding_conservation_phastcons.bed: This BED file lists the PhastCons scores for all coding genes.
**File:** bcCRE_conservation_phastcons.bed: This BED file lists the PhastCons scores for all b-cCRE regions.
**File:** scATAC_conservation_phastcons.bed: This BED file lists the PhastCons scores for all identified snATAC-seq peaks.

### 4 Determining cell-type-specific eQTLs from single-cell data

#### 4.1 ***scQTLs*** - Cell-type-specific eQTL analysis

**Main manuscript reference:** First, second, fourth and sixth supplementary reference in the first paragraph, and first supplementary reference of the second paragraph of "Determining cell-type-specific eQTLs from single-cell data."

To evaluate the association between genotypic variation and gene expression variation within distinct cell types, we used snRNA-seq data to identify cis-eQTLs within 17 distinct cell types ("scQTLs"). To do so, we first merged genotype, UMI expression, and covariate data across all 12 cohorts. Within this merged cohort, we excluded any samples meeting the following criteria:
- Some samples were duplicated across different cohorts; we took one instance of such samples to avoid data redundancy in our analysis.
- Samples missing genotype data were excluded.
- We excluded any samples with missing covariates.

For each cell type, we first generated pseudo-bulk matrices by averaging the UMI counts across all nuclei within a given sample for that cell type. For a given individual to be included in the pseudobulk matrix of a given cell type, we enforced that the cell type must be represented by >50 nuclei. Additionally, we enforced that each individual must have >300 nuclei in total. We then normalized the psuedobulk expression matrix for each cell type by performing a log(CPM+1) transformation. We filtered out lowly expressed genes by removing genes in which the fraction of samples with non-0 expression is ≤10%. These filtered matrices were then used as the psuedobulk expression matrices for our scQTL analyses. For genotype data, we removed all variants with MAF<0.05 from the uniformly processed imputed genotypes.

We additionally performed a simulated statistical power analysis using powerEQTL (*166*), and found that a minimum of 200 samples per cell type would be required to detect significant eQTLs ($p<0.05$, one-way unbalanced ANOVA) with 80% power (**fig. S36D**), given the parameters of 50 nuclei/sample, 80 million SNP/gene pairs tested, >0.05 SNP MAF, SNP effect size of 0.13 and standard deviation (SD) of 0.13, 0.5 intra-group correlation, and an error rate of 0.5.

We adopted a highly standardized approach for finding scQTLs. In particular, after generating pseudobulk matrices for each cell type, we followed the same general procedure as that adopted by GTEx (*5*). By using a 1 MB cis window up- and downstream of the TSS of each gene, we implemented QTLtools (*167*) using biological sex, age, diagnosis, cohort, five genotype PCs (to control for ancestry), and 100 expression PCs as covariates (to control for hidden batch effects). We used 100 expression PCs in particular, as this was found to optimize the total number of significant eGenes across cell types (**fig. S36E**). Details regarding significance detection in eGene searches are given in the next paragraph.

Our scheme for defining significance is detailed by the approach taken by GTEx (*5*) in their application of FastQTL (*168*), and we summarize this framework here. QTLtools was used to perform permutation-based searches to first identify a set of eGenes for each cell type (at an FDR threshold of 0.05). Our results are based on performing 10,000 permutations, though we note that the gene-level p-values (significance was determined using a fitted beta distribution derived from random permutations, as detailed in (*168*) and (*167*)) obtained do not differ substantially if only 1,000 permutations had been used (**fig. S39**). eGenes were determined to be associated with at least one significant eSNP within its cis-window using permutation tests (with 10,000 permutations). FDR values were estimated using Storey q-values (*169*) and we designated all such genes as “eGenes” with an FDR threshold of 0.05. These corrections were applied at the level of each cell type individually. In addition, for each cell type separately, in order to call all significant eSNPs associated with each eGene, we defined a genome-wide threshold ($p_t$) as the beta distribution-derived p-value (also called the “beta-adjusted p-value” within QTLtools) associated with the least significant gene among all eGenes within a given cell type. For each eGene, a nominal p-value threshold was determined using the eGene's associated beta distribution $f_g$ by setting this eGene-specific nominal p-value threshold to $F_g^{-1}(p_t)$, with $F_g^{-1}$ being the inverse cumulative beta distribution for that eGene. Any SNP within the 1 MB cis-window of the eGene having a nominal p-value less than the eGene-specific nominal p-value threshold $F_g^{-1}(p_t)$ was considered to be a significant eSNP associated with that eGene. All such significant pairs of eSNPs and eGenes were taken to be significant scQTLs.

We assessed the functional and disease relevance of eGenes from our primary scQTL callset using GO enrichment analysis (**fig. S51**) and disease-associated gene set overrepresentation analysis (**fig. S50**). GO enrichment analysis was performed using g:Profiler (*170*). Additionally, we annotated eGenes with sets of high-confidence genes related to ASD (Tier 1 genes in the Simons Foundation Autism Research Initiative [SFARI] Gene database), schizophrenia, bipolar disorder, AD, and aging (*171–175*). eGenes with disease annotations were visualized across the human genome in **Fig. 4G** using the PhenoGram package (https://ritchielab.org/software/phenogram-downloads).

Various scQTL callsets are available on the brainSCOPE portal, including the primary scQTL callset and several modified callsets calculated with different parameters. These include those that used LD-based filtering of genotypes before calculation. LD pruning was performed using the software PLINK (*176*) (v. 1.90-beta5.3) with the following parameters:

a) window size of 50 SNPs
b) LD calculated between each pair of SNPs in the window
c) one of a pair of SNPs removed if the LD is >0.5
d) the window shifted five SNPs forward to repeat the procedure

Each set of QTLs contains one file per cell type, with columns (in order) given as:

1. The common gene name associated with the eGene
2. The gene chromosome
3. Start position of the gene
4. Start position of the gene (provided again as output from the software)
5. The gene strand
6. The number variants in the cis window for this gene
7. The distance between the variant and the gene start position
8. The variant ID
9. The variant chromosome
10. The start position of the variant
11. The start position of the variant (provided again as output from the software)
12. The nominal p-value of the association between the variant and the gene
13. The $r^2$ of the linear regression
14. The beta (slope) of the linear regression
15. Indicator variable denoting whether this variant was the best hit for this gene

A given eGene may be associated with many eSNPs via scQTLs. However, the effect of a given eSNP may be conditionally dependent on the effects of other eSNPs for the same eGene. Thus, we separately also performed a conditional analysis using QTLtools (*167*), wherein the objective was to identify the number of independent signals for each eGene (where a signal is a group of eSNPs with conditionally dependent effects on gene expression, and with effects that are conditionally independent of eSNPs that do not belong to the same signal). The approach implemented here aims to (a) identify the set of independent signals, (b) find the best candidate eSNP for each signal, and then to (c) assign all other eSNPs to their appropriate signal.

For this conditional QTL analysis, the columns in the data files are as follows:

1. The gene ID or if one of the grouping options is provided, then gene group ID
2. The gene chromosome
3. Start position of the gene
4. (Dummy variable)
5. The gene strand
6. The number variants in the cis window for this gene.
7. The distance between the variant and the gene start positions.
8. The most significant variant ID (here, the most significant variant is that with the lowest nominal p-value, with each p-value being derived from the coefficient associated with the linear regression of the gene's normalized expression on the genotype dosage associated with that variant, while including covariates in the multivariate linear regression). This p-value is thus derived from a t-test on the coefficient associated with the variant within the multivariate linear regression. We applied the --normal flag when running QTLtools \cite{28516912}. More detailed descriptions of deriving p-values from the coefficients in linear regression models can be found in the works by Casella et al (*177*)\cite{Berger, R. L., & Casella, G. (2001). Statistical inference. Duxbury} and James et al (*178*) \cite{James, Gareth, et al. An introduction to statistical learning. Vol. 112. New York: Springer, 2013.});

9. The most significant variant's chromosome (here, this is simply the chromosome associated with the most significant variant; please see details within item #8 of this list)
10. The start position of the most significant variant (here, this is simply the locus associated with the most significant variant; please see details within item #8 of this list)
11. (Dummy variable)
12. The rank of the association. This tells you if the variant has been mapped as belonging to the best signal (rank=0), the second best (rank=1), etc. As a consequence, the maximum rank value for a given gene tells you how many independent signals there are (for example, rank=2 means 3 independent signals).
13. The nominal forward p-value of the association between the most significant variant and the gene. Here, significance is defined similarly to how it is defined within item #8 of this list, but with the dependent variable being the residualized gene expression during the forward conditional pass. Further details can be found in \cite{28516912}.
14. The r squared of the forward linear regression.
15. The beta (slope) of the forward linear regression.
16. Whether or not this variant was the forward most significant variant. (To derive significance, see description within item #13 of this list).
17. Whether this variant was significant. (To derive significance, see description within item #13 of this list; details are provided in \cite{28516912}).
18. The nominal backward p-value of the association between the most significant variant and the gene. For details regarding significance in the backward pass, please see \cite{28516912}
19. The r squared of the backward linear regression.
20. The beta (slope) of the backward linear regression.
21. Whether or not this variant was the backward most significant variant. For details regarding significance in the backward pass, please see \cite{28516912}
22. Whether this variant was significant. (To derive significance, see description within item #13 of this list; details are provided in \cite{28516912}).

Specific callsets and associated meta-data files are available on the brainSCOPE portal as follows:

**File:** [celltype]_sig_QTLs.dat: These files represent the primary set of cell-type-specific scQTLs identified for 24 cell types, processed with the standardized (GTEx-based) pipeline.
**File:** [celltype]_sig_eGenes.dat (**data S15**): These files list the significant eGenes for 17 cell types with significant scQTL results, used as inputs for functional validations. For a discussion of how significance was derived, please refer to the paragraph above that starts with "Our scheme for defining significance is detailed by the approach taken by GTEx…"
**File:** [celltype].SNPs.dat: These files list all variant IDs used per cell type in the scQTL calculations.
**File:** [celltype].cov.100_expr_PCs.bed: These files represent sample-level covariates used for QTL calculations in each cell type. Each column represents an individual, while rows represent binary variables for biological sex, age of death, diagnosis, cohort, five genotype PCs, and up to 100 expression PCs.

**File:** core_scqtl_processing_code.tar: TAR file containing the code used for calculating scQTLs using the standardized (GTEx-based) pipeline.
**File:** [celltype]_sig_QTLs.dat: These files represent cell-type-specific scQTLs identified for 17 cell types, processed with the standard pipeline but utilizing LD variant selection.
**File:** [aggregated_celltype]_sig_QTLs.dat: These files represent cell-type-specific scQTLs identified for three PFC cell type groupings (inhibitory neurons, excitatory neurons, and non-neuronal cells), processed using the standard pipeline.
**File:** conditional.[celltype].txt**:** These files represent cell-type-specific scQTLs processed with a conditional regression-based QTL identification pipeline.
**File:** conditional_top_variants_per_signal_[celltype].txt**:** These files represent cell-type-specific scQTLs processed with a conditional regression-based QTL identification pipeline, and contain the top variants for each eGene identified in the analysis.
**File**: scQTL_disease_overlap.txt (**data S17**): This file contains a list of eGenes found in any cell type from the primary analysis that are annotated for four brain-related diseases and traits. "X" annotations in each of the four disease columns indicate if an eGene is associated with a disease.

#### 4.2 ***Bayesian scQTLs -*** scQTLs using Bayesian linear mixed effects models

**Main manuscript reference:** Third supplementary reference in the first paragraph of "Determining cell-type-specific eQTLs from single-cell data."

In addition to providing scQTLs based on standard approaches and methods, we also provide supplementary scQTL callset based on Bayesian analysis. Specifically, we quantify the relationship between genotype dosage and gene expression using a Bayesian linear mixed effects model. Our description and implementation follow closely those given by Hoff *(179).*

Before discussing how a Bayesian linear mixed effects model works, it is helpful if we first briefly review the basic principles behind standard QTL approaches. In any QTL analysis using linear regression, the key value of interest is the slope associated with the regression of a given gene's expression (*expr*) on the genotype dosage associated with a particular variant (*g*), while simultaneously controlling for a set of q covariates ($cov_1$, $cov_2$, … $cov_q$):

$$expr \sim \beta_0 + \beta_g g + \beta_{cov\_1} cov_1 + \beta_{cov\_2} cov_2 + \dots + \beta_{cov\_q} cov_q$$

In the context of this multivariate regression, the key value of interest is the parameter $\beta_g$, which quantifies the association between the genotype dosage *g* and expression. A QTL search under this model is framed as a hypothesis test, with $H_0$ denoting the scenario wherein $\beta_g = 0$. A standard QTL search entails estimating the value of $\beta_g$ using a least squares linear regression, and then evaluating the evidence against $H_0$ using a p-value (namely, larger absolute values of $\beta_g$ that deviate from 0 provide greater evidence in favor of $H_1 \neq 0$).

A Bayesian-based approach to this problem treats the parameter $\beta_g$ as an unknown value that can be described by a probability distribution. In particular, this probability distribution is first given as a prior $p(\beta_g)$. It is then updated after evaluating our dataset, using gene expression values across a large cohort of individuals, along with their corresponding genotypes and covariates. This updated distribution is the posterior probability distribution $p(\beta_g|dataset)$ associated with the parameter $\beta_g$. The posterior distribution $p(\beta_g|dataset)$ is defined over the range of all possible $\beta_g$ values (negative ∞ to positive ∞ in our case). $p(\beta_g|dataset)$ optimally describes our available information for $\beta_g$, given our prior $p(\beta_g)$, our dataset, and a sampling model. Instead of evaluating the strength of evidence for or against $H_0$ using p-values, a Bayesian-based approach evaluates the degree to which 0 lies within or outside of the posterior $p(\beta_g|dataset)$. A scenario wherein the value 0 lies very much within the high-density region of $p(\beta_g|dataset)$ provides evidence against the presence of a QTL (**fig. S37A**), whereas a scenario wherein 0 lies in one of the very far tails (or far from high-density regions of $p(\beta_g|dataset)$) provides evidence that $\beta_g \neq 0$, and thus provides evidence in favor of a QTL (**fig. S37B**).

In the context of this study, we searched for QTLs in m=24 cell types. Thus, the general model for our analyses takes the following general form:

$$
\begin{aligned}
expr_{Astro} &\sim \beta_{0(Astro)} + \beta_{g(Astro)}\, g + \beta_{cov\_1(Astro)}\, cov_1 + \ldots + \beta_{cov\_q(Astro)}\, cov_q \\
expr_{Oligo} &\sim \beta_{0(Oligo)} + \beta_{g(Oligo)}\, g + \beta_{cov\_1(Oligo)}\, cov_1 + \ldots + \beta_{cov\_q(Oligo)}\, cov_q \\
&\;\;\vdots \\
expr_{OPC} &\sim \beta_{0(OPC)} + \beta_{g(OPC)}\, g + \beta_{cov\_1(OPC)}\, cov_1 + \ldots + \beta_{cov\_q(OPC)}\, cov_q
\end{aligned}
$$

For a given cell type, the set of parameters can be represented by a vector $\boldsymbol{\beta}$. For example, $\boldsymbol{\beta}_{Astro}$ is a vector representing $\langle \beta_{0(Astro)}, \beta_{g(Astro)}, \beta_{cov\_1(Astro)}, \beta_{cov\_2(Astro)}, \ldots \beta_{cov\_q(Astro)} \rangle$. The set of explanatory variables can likewise be represented in a vector $\boldsymbol{X}$, with $\boldsymbol{X}$ being the vector $\langle 1, g, cov_1, cov_2, \ldots cov_q \rangle$. Likewise, for a given cell type, we have N expression values, where N denotes the number of individuals for whom expression data are available (or the sample size used in a regression for a given cell type). These expression values may therefore also be represented by vectors ***expr***. Thus, a more compact representation of the set of equations above may be given as:

$$
\begin{aligned}
\boldsymbol{expr}_{Astro} &\sim \boldsymbol{\beta}_{Astro}^{T}\, \boldsymbol{X}_{Astro} \\
\boldsymbol{expr}_{Oligo} &\sim \boldsymbol{\beta}_{Oligo}^{T}\, \boldsymbol{X}_{Oligo} \\
&\;\;\vdots \\
\boldsymbol{expr}_{OPC} &\sim \boldsymbol{\beta}_{OPC}^{T}\, \boldsymbol{X}_{OPC}
\end{aligned}
$$

Our objective is to approximate posterior distributions associated with the parameter vectors, with these posterior distributions given by $p(\boldsymbol{\beta}_{Astro}|\boldsymbol{X}_{Astro}, \boldsymbol{expr}_{Astro})$, $p(\boldsymbol{\beta}_{Oligo}|\boldsymbol{X}_{Oligo}, \boldsymbol{expr}_{Oligo})$, … $p(\boldsymbol{\beta}_{OPC}|\boldsymbol{X}_{OPC}, \boldsymbol{expr}_{OPC})$. From these posterior distributions, we can easily determine the marginal posterior distributions that are truly of interest in the context of our analysis – namely, the posterior distributions associated with the parameters that describe the relationship between genotype dosage and expression within each cell type (represented by the parameters $\beta_{g(Astro)}$, $\beta_{g(Oligo)}$, ... $\beta_{g(OPC)}$).

In order to approximate the posterior distributions p($\boldsymbol{\beta}_{Astro}$|$\boldsymbol{X}_{Astro}$, $\boldsymbol{expr}_{Astro}$), p($\boldsymbol{\beta}_{Oligo}$|$\boldsymbol{X}_{Oligo}$, $\boldsymbol{expr}_{Oligo}$), … p($\boldsymbol{\beta}_{OPC}$|$\boldsymbol{X}_{OPC}$, $\boldsymbol{expr}_{OPC}$), we must first define prior distributions for the parameter vectors in each cell type, p($\boldsymbol{\beta}_{Astro}$), p($\boldsymbol{\beta}_{Oligo}$), … p($\boldsymbol{\beta}_{OPC}$). We set each cell type's parameter vector prior as a multivariate normal (MVN) distribution that itself is parameterized by a mean vector **Θ** and a covariance matrix **Σ**:

$$
\begin{aligned}
p(\boldsymbol{\beta}_{Astro}) &\sim MVN(\boldsymbol{\Theta}, \boldsymbol{\Sigma}) \\
p(\boldsymbol{\beta}_{Oligo}) &\sim MVN(\boldsymbol{\Theta}, \boldsymbol{\Sigma}) \\
&\;\;\vdots \\
p(\boldsymbol{\beta}_{OPC}) &\sim MVN(\boldsymbol{\Theta}, \boldsymbol{\Sigma})
\end{aligned}
$$

Note that the same vector **Θ** and matrix **Σ** are used for all cell types. Importantly, **Θ** and **Σ** are not known in advance, and specific values are thus not assigned to them. Instead, **Θ** and **Σ** are also modeled using a Bayesian approach, and therefore each have their own respective prior distributions, p(**Θ**) and p(**Σ**). We use an MVN distribution for p(**Θ**) and an inverse Wishart distribution for p(**Σ**):

$$
\begin{aligned}
p(\boldsymbol{\Theta}) &\sim MVN(\boldsymbol{\mu}_0, \boldsymbol{\Lambda}_0) \\
p(\boldsymbol{\Sigma}) &\sim \text{inverse-Wishart}\ (\eta_0, \mathbf{S}_0^{-1})
\end{aligned}
$$

The model described here is a Bayesian linear mixed effects model (or a "normal hierarchical regression model"), and it may be represented diagrammatically using the following hierarchical structure:

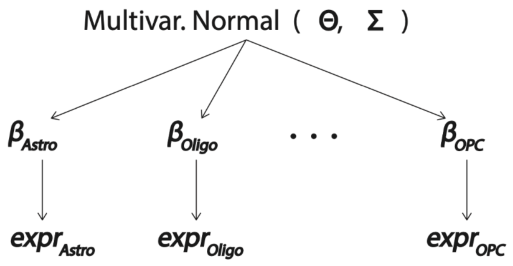


Treating the common mean vector **Θ** and covariance matrix **Σ** as unknown parameters that are estimated under a Bayesian framework provides important behavior for this model: namely, it provides a mechanism for sharing information across cell types. By contrast, if we were to alternatively set **Θ** and **Σ** to specific fixed values in advance, then $\boldsymbol{\beta}_{Astro}$, $\boldsymbol{\beta}_{Oligo}$, … $\boldsymbol{\beta}_{OPC}$ would be conditionally independent parameter vectors (each having the same priors), so there would be no mechanism to share effects across cell types. The notion of incorporating shared effects between cell types may be justified by considering what the parameters within each of the vectors $\boldsymbol{\beta}_{Astro}$, $\boldsymbol{\beta}_{Oligo}$, … $\boldsymbol{\beta}_{OPC}$ represent. Consider the single parameter $\beta_{g(Astro)}$ as an example. $\beta_{g(Astro)}$ represents the effect of a particular variant on gene expression, while controlling for covariates. $\beta_{g(Astro)}$ may be found to be a very large positive value, but if the sample size associated with astrocytes is small, then this may reduce our confidence that $\beta_{g(Astro)}$ truly has a very large value. In this regard, the large estimate for $\beta_{g(Astro)}$ may be an artifact that stems from the fact that a low sample size confers high variance when estimating a parameter. By contrast, the remaining cell types (oligodendrocytes, OPC, etc.) may have parameter estimates of $\beta_{g(Oligo)}$, … $\beta_{g(OPC)}$ that are very close to 0, and these cell types may have much larger sample sizes. Their larger sample sizes provide us with confidence that the true underlying values $\beta_{g(Oligo)}$, … $\beta_{g(OPC)}$ are 0. Given that the underlying biology among the various cell types of the PFC are not completely independent, this in turn may further reduce our confidence that $\beta_{g(Astro)}$ truly has a large positive value. By sharing information between cell types, we may use the near-0 estimates of $\beta_{g(Oligo)}$, … $\beta_{g(OPC)}$ to inform our estimate for $\beta_{g(Astro)}$, effectively helping to mitigate the effect that the small sample size in astrocytes has on yielding an extreme estimate for $\beta_{g(Astro)}$.

Looking back to the hierarchical diagram above, we see that our full model has the unknown parameters **Θ**, **Σ** , $\boldsymbol{\beta}_{Astro}$, $\boldsymbol{\beta}_{Oligo}$, … and $\boldsymbol{\beta}_{OPC}$ (for clarity of exposition, we omit discussion of another parameter σ, which describes the variance associated with gene expression at a particular genotype value within a given cell type; see *(179)* for further details). $\boldsymbol{\mu}_0$, $\boldsymbol{\Lambda}_0$, $\eta_0$, and $\mathbf{S}_0^{-1}$ are hyperparameters for which we pre-specify precise values (see details below). Our objective is thus to approximate the joint posterior distribution of all unknown parameters:

$$p(\boldsymbol{\Theta}, \boldsymbol{\Sigma}, \boldsymbol{\beta}_{Astro}, \boldsymbol{\beta}_{Oligo}, \dots \boldsymbol{\beta}_{OPC} \mid \boldsymbol{X}_{Astro}, \boldsymbol{X}_{Oligo}, \boldsymbol{X}_{OPC}, \boldsymbol{expr}_{Astro}, \boldsymbol{expr}_{Oligo}, \boldsymbol{expr}_{OPC})$$

We approximate this joint posterior distribution using Gibbs sampling. Gibbs sampling is a Markov chain in which we sample from the full conditional distribution of each parameter in turn, while conditioning on the data as well as the most recently sampled values for all of the other parameters. Under our assumed priors given above, these full conditional distributions are given as follows (for the $\boldsymbol{\beta}$ vectors, the full conditional distribution is only provided for $\boldsymbol{\beta}_{Astro}$, but full conditional distributions associated with the other vectors $\boldsymbol{\beta}_{Oligo}$, … $\boldsymbol{\beta}_{OPC}$ have the same general form):

$$p(\boldsymbol{\beta}_{Astro} \mid \boldsymbol{X}_{Astro}, \boldsymbol{expr}_{Astro}, \boldsymbol{\Theta}, \boldsymbol{\Sigma}, \sigma) \sim \mathrm{MVN}(\boldsymbol{\mu}_{Astro}, \boldsymbol{\Sigma}_{Astro})$$

$$\boldsymbol{\mu}_{Astro} = [\boldsymbol{\Sigma}^{-1} + \boldsymbol{X}_{Astro}^{T}\boldsymbol{X}_{Astro}\sigma^{-2}]^{-1} (\boldsymbol{\Sigma}^{-1}\boldsymbol{\Theta} + \boldsymbol{X}_{Astro}^{T}\boldsymbol{expr}_{Astro}\sigma^{-2})$$

$$\boldsymbol{\Sigma}_{Astro} = [\boldsymbol{\Sigma}^{-1} + \boldsymbol{X}_{Astro}^{T}\boldsymbol{X}_{Astro}\sigma^{-2}]^{-1}$$

$$p(\boldsymbol{\Theta} \mid \boldsymbol{\Sigma}, \boldsymbol{\beta}_{Astro}, \boldsymbol{\beta}_{Oligo}, \dots \boldsymbol{\beta}_{OPC}) \sim \mathrm{MVN}(\boldsymbol{\mu}_m, \boldsymbol{\Lambda}_m)$$

$$\boldsymbol{\mu}_m = [\boldsymbol{\Lambda}_0^{-1} + m\boldsymbol{\Sigma}^{-1}]^{-1} [\boldsymbol{\Lambda}_0^{-1}\boldsymbol{\mu}_0 + m\boldsymbol{\Sigma}^{-1}\underline{\beta}]$$

$$\boldsymbol{\Lambda}_m = [\boldsymbol{\Lambda}_0^{-1} + m\boldsymbol{\Sigma}^{-1}]^{-1}$$

$\underline{\beta}$ = mean value of the parameter vectors $\boldsymbol{\beta}_{Astro}$, $\boldsymbol{\beta}_{Oligo}$, … $\boldsymbol{\beta}_{OPC}$

$$p(\boldsymbol{\Sigma} \mid \boldsymbol{\Theta}, \boldsymbol{\beta}_{Astro}, \boldsymbol{\beta}_{Oligo}, \dots \boldsymbol{\beta}_{OPC}) \sim \text{inverse-Wishart}\ (\eta_0 + [\mathbf{S}_0 + \mathbf{S}_\theta]^{-1})$$

Our implementation relies on first using the significant eGenes from running 10,000 permutations (For a discussion of how significance was derived, and how it's used throughout this paragraph, please see supplementary section 4.1 scQTLs - Cell-type-specific eQTL analysis; especially the paragraph above that starts with "Our scheme for defining significance is detailed by the approach taken by GTEx..."). This is the first step in the 2-step multiple testing correction scheme used to find our core set of scQTLs. We used the significant eGenes identified from these permutations. For each significant eGene (FDR<0.05) in each cell type, we evaluated the top eSNP per eGene. This top eSNP is the most significant eSNP for the eGene, with significance given by the eSNP's associated nominal p-value. Using this eSNP-eGene pair (which we term an "anchor QTL" for the Bayesian approach), we implemented the Bayesian linear mixed effects model across all 24 cell types, using 10 expression PCs when searching for Bayesian QTLs (see **fig. S36C, S36E, S37**, and **table S8**). Any covariates with very high collinearity were removed from the regression. To set the priors in our model, we used the least squares regression estimates within each cell type to set $\boldsymbol{\mu_0}$ (thus, $\boldsymbol{\mu_0}$ was set to be the mean among the "standard" least squares regression estimates from each of the 24 cell types). Likewise, $\boldsymbol{\Lambda_0}$ was set to equal the covariance matrix associated with the least squares regression estimates across the 24 cell types. $\mathbf{S_0}$ was also set to equal the covariance among the least squares regression estimates.

In order to approximate the posterior distributions associated with the parameter vectors that model the relationship between the eSNP (as well as the covariates) with the eGene's expression, we then ran a Gibbs sampler consisting of 25,000 steps. We thinned the sequence to mitigate the strong dependencies between adjacent steps in the Gibbs sampler (which is a Markov chain) by saving every 10th set of parameter values in the Gibbs sampler.

To determine whether a given eSNP-eGene pair within a given cell type was a Bayesian QTL, we evaluated whether the value $\beta_g = 0$ (for that cell type) appeared within the highest posterior density (HPD) associated with $\beta_g$ for each cell type. The intuition behind this approach is given in **fig. S37**. In our implementation, we consider a given eSNP-Gene pair not to be a Bayesian QTL if the 0.9999999999 HPD overlaps the value $\beta_g = 0$. In practice, this means that a given eSNP-eGene pair is considered to be a Bayesian QTL if every one of the 2,500 sampled $\beta_g$ values is >0 (designating a QTL with a positive effect size) or if every one of the 2,500 sampled $\beta_g$ values is <0 (designating a QTL with a positive effect size).

The results from this analysis are given in **fig. S36C** and **table S8**. Three key features may be highlighted:

1) The number of scQTLs identified using this Bayesian framework is substantially less than the number of scQTLs identified using the standard linear regression-based approach (ie, our core scQTL callset). In this regard, the Bayesian framework offers a more conservative set of results, and it is largely a consequence of the fact that our framework relies on a small set of anchor QTLs as a base (as detailed above, these anchor QTLs are the eSNP-eGene pairs that link the most significant eSNP with a significant eGene). However, the diminished number of scQTLs identified under the Bayesian framework may also partially result from a phenomenon whereby a significant scQTL using the least squares linear regression becomes non-significant under the Bayesian approach. This may occur, for instance, if a significant effect size occurs in only one cell type; the other cell types (with non-significant effects) may effectively pull the one strong effect size toward 0, thereby causing it to be non-significant. This is a result of the shared effects between the cell types.

2) The number of QTLs identified using the Bayesian approach is relatively stable across cell types. This is also a direct consequence of the shared effects that are built into the Bayesian linear mixed effects model.

3) There are several cell types (namely, L5.ET, Pax6, SMC, Endo__VLMC, L6.IT.Car3, Sncg, and Immune) for which we were unable to identify any significant QTLs using least squares linear regression (see **fig. S36C**). However, a limited number of significant scQTLs have been identified using the Bayesian framework detailed here. Thus, this framework may be especially valuable for low-abundance cell types in which small sample sizes make it otherwise difficult to identify significant eSNP-eGene associations using standard linear regression-based approaches.

An UpSet plot delineating the extent of overlap among various scQTL types (standard, Bayesian, conditional, and LD-pruned) is provided in **fig. S38**.

As with the primary ("core") scQTL analysis, the scQTL callsets for the Bayesian analysis are available on the brainSCOPE portal. Columns indicate cell type, eGene, eSNP, and the posterior mean and standard deviation associated with posterior beta effect sizes from the Gibbs sampler. We also provide a similar file with overlapping scQTL results across methods on the brainSCOPE portal.

**File:** bayesian_scQTLs.dat: These files represent cell-type-specific scQTLs processed with the Bayesian model-based QTL identification pipeline.
**File:** Bayesian_QTL_code.tar.gz: TAR file contains code used for calculating scQTLs with the Bayesian model-based pipeline.
**File**: metaQTLs.txt: This file lists a combined set of QTLs discovered in each analysis (primary, conditional, Bayesian, and dynamic) for all cell types. Each scQTL entry lists analysis type, cell type, gene and SNP identifiers, and associated statistics.

#### 4.3 ***Dynamic scQTLs*** - Dynamic sc-eQTL analysis

**Main manuscript reference:** Fifth supplementary reference in the first paragraph, and first supplementary reference of the last paragraph of "Determining cell-type-specific eQTLs from single-cell data."

To further expand our sc-eQTL analysis to a true single-cell resolution, we also developed a Poisson mixed effect (PME) model that incorporates continuous cell-state scores generated from trajectory analysis and the interaction between genotype and trajectory. With this model, we can calculate scQTLs at the single-cell level and assign a unique effect size to each cell, resulting in what we refer to as "dynamic sc-eQTLs". The total effect size ($\beta_{total}$) of the dynamic sc-eQTLs varies across single cells, reflecting the dynamic nature of gene regulation. Furthermore, the $\beta_{total}$ is partially related to the continuous trajectory, which allows us to extrapolate eQTLs to cells in which we may not detect these eQTLs with the standard pseudo-bulk approach.

To ensure rigorous control over potential batch effects, we limited our dynamic sc-eQTL analysis to a single cohort, the SZBDMulti-seq cohort, which comprises 24 healthy donors, 24 schizophrenia patients, and 24 bipolar patients. We excluded four individuals with <1,000 cells, resulting in a total of 312,131 neurons (L2/3 IT, L4 IT, L5 IT, and L6 IT) from 68 SZBD-Kellis individuals. To correct for batch effects, we integrated the data using SCALEX (*180*).

Next, we conducted a Slingshot trajectory analysis using the same pipeline as described in the previous "Trajectory analysis" section. We extracted pseudo-time from the gene expression patterns that varied along the cortical depth axis, specifically from L2/3 IT to L6 IT neurons (**fig. S53**), and incorporated it into our subsequent PME model.

In our PME model, we modeled the raw UMI counts of a gene as a function of genotype, adjusting for trajectory cell state score and multiple individual and cell-level confounders including age, biological sex, genotype PCs, total UMI counts, and gene expression PCs. Individual-level random effects were also included in the model:

$$\log(E_i) = \theta + \beta_G X_{d,geno} + \beta_{age} X_{d,age} + \beta_{sex} X_{d,sex} + \sum_{k=1}^{5} \beta_{gPC_k} X_{d,gPC_k} + \beta_{nUMI} X_{i,nUMI} + \beta_{ptime} X_{i,ptime} + \sum_{k=1}^{80} \beta_{ePC_k} X_{i,ePC_k} + (\phi_d | d)$$

Here, E denotes the expression level of a gene in cell i, θ denotes the intercept, and all βs denote fixed effects as indicated (nUMI: number of UMI counts, gPC: genotype PC, ePC: single nucleus mRNA expression PC, ptime: pseudo-time cell state score generated from trajectory analysis) for covariates at cell i or donor d level. The donor is also modeled as a random effect.

To test whether there is a significant interaction between the trajectory and the genotype, we further include the interaction term Gxptime in our model:

$$\log(E_i) = \theta + \beta_G X_{d,geno} + \beta_{age} X_{d,age} + \beta_{sex} X_{d,sex} + \sum_{k=1}^{5} \beta_{gPC_k} X_{d,gPC_k} + \beta_{nUMI} X_{i,nUMI} + \beta_{ptime} X_{i,ptime} + \sum_{k=1}^{80} \beta_{ePC_k} X_{i,ePC_k} + \beta_{Gxptime} X_{d,geno} X_{i,ptime} + (\phi_d | d)$$

Overall, 6,225 top eQTLs (eGene – most significant eSNP pairs) identified in at least one IT neuron cell type were tested with our PME model. Given the much more limited sample size (a reduction from around 300 to 68 individuals), we still successfully reproduced 4,186 (67.2%) top eQTLs from the primary scQTL dataset, among which 1,692 showed significant interactions. The significance was tested via the likelihood ratio test (**fig. S52**).

Similar to the main scQTL analysis, the dynamic scQTL callsets are represented as individual text files for each assessed cell type on the brainSCOPE portal:

**File:** full_dynamic_eqtl.tsv: This file lists the full set of eQTLs used as input for the PME model (6,225 top eQTLs identified with the pseudo-bulk approach).
**File:** sig_dynamic_eqtl.tsv (**data S18**): This file lists the 4,186 significant dynamic eQTLs identified with the PME model (SNP term only).
**File:** sig_ptime_dynamic_eqtl.tsv (**data S19**): This file lists the 1,692 significant dynamic eQTLs identified with the PME model (SNP and interaction terms).

**File:** SZBD.ptime.tsv: This file lists the pseudo-time values for all tested excitatory neuron cells from samples in the SZBDMulti-Seq cohort (generated via SCALEX and Slingshot).

#### 4.4 ***Isoform QTLs*** - Identify QTLs for isoform expression for each cell type

**Main manuscript reference:** Seventh supplementary reference in the first paragraph of "Determining cell-type-specific eQTLs from single-cell data."

In addition to identifying cell-type-specific eQTLs from our datasets, we identified QTLs for isoform-specific expression ("isoQTLs") using cell-type-specific single-cell datasets. This analysis is challenging, as the 3' sequencing direction of the 10X snRNA-seq technology leads to reduced sequencing rates in isoforms that differ at the 5' end of the gene, compounding the sparsity inherent to single-cell expression data (*181*). To overcome this issue, we used the SCASA algorithm for isoform quantification, which fits read-count data onto an alternating expectation maximization model to identify and quantify single-cell expression for clusters of isoforms that can be detected in each sample (*181*).

After running SCASA for all 388 snRNA-seq samples with default parameters, we transformed the expression values of individual cells into pseudo-bulk data across cell types, similar to the main expression data. Briefly, we performed the following steps: matched cells with their annotated cell types; discarded cells without annotations, those with <500 UMIs, or those with <200 isoforms expressed; removed cells of the same cell type if there were <10 total cells per individual; summed the non-zero read counts across samples and cell types; normalized expression values to CPM without log transformation; and removed isoform clusters where >95% of individual values were 0 or NA. After filtering, we were able to assess an average of 42,662 nuclei across 187 individuals per cell type, with wide variability across cell types (between 871 and 287,831 nuclei and 11-320 individuals) (**fig. S40A**). We next used the software package sQTLseekeR2-nf (*182*) to identify isoQTLs for each individual cell type in our dataset. Due to low sample sizes and cell counts, we merged select similar cell types for this analysis (chandelier with Pvalb, endothelial with VLMC, and Sst with Sst_chodl), as was done in the main scQTL analysis, resulting in 23 total cell types. sQTLseekerR2 models the effects of genotype (imputed variants with >0.05 MAF) towards isoform expression (CPM values), using cohort, biological sex, age of death, disorder, nuclei count, three expression PCs (calculated using the prcomp package in R), and three genotype PCs (calculated using PLINK2) as covariates. Similar to the scQTL calculations described above, sQTLseekeR2 first identified nominally significant isoSNPs ($p<0.05$, one-way factorial test) located within 5 kbp of the canonical gene TSS ("isoGene") after performing additional QC steps (such as removing genes without alternative isoforms in the cohort), and found the associated isoGenes for each isoSNP that passed multiple testing corrections. We then implemented the permuted mode of sQTLseekeR2 to identify beta distribution-derived p-values and FDR values for each isoGene, and calculated individual nominal p-value thresholds for each isoGene for isoSNP discovery. We also excluded isoQTLs at complex genetic loci with high variation, namely the HLA locus and *ARL17B* at the 17q21.31 locus, from downstream analysis.

Due to low sample size, difficulty in detecting isoforms from 10X sequencing data, and noise from unspliced transcripts in the nucleus, our power to detect isoQTLs in specific cell types is diminished. As such, we report a total of 1,389 putative isoGenes corresponding to 133,861 isoSNPs across 22 of the 23 cell types with beta distribution-derived p-values <0.05, without filtering for FDR at the isoGene level (**fig. S40B**). Note that SST Chodl cells did not yield any isoQTLs passing nominal significance (p>0.05, beta distribution). With this approach, end users can set their own FDR threshold for candidate isoQTLs. For instance, we found 86 significant isoGenes at FDR<0.25, 32 isoGenes at FDR<0.1, and 21 isoGenes at FDR<0.05 (**figs. S40C-D**). In particular, the 21 isoQTLs with FDR<0.05 that we considered for downstream analysis, which corresponded to 575 total isoSNPs (or 527 unique isoSNPs), were present in nine cell types with larger sample sizes and nuclei counts. Most of these isoQTLs were unique to a particular cell type, with only 48/527 isoSNPs (9%) present in multiple cell types (**fig. S41A**). We also found that 302/527 (57.3%) of the isoSNPs overlapped with bulk RNA-seq-based isoQTLs in the adult brain (**fig. S41B)**. Finally, we observed a bias towards isoSNPs contributing to complex 3' events (390/575, 74%), likely due to the 3' bias inherent in 10X Genomics single-cell RNA-seq (**fig. S41C**). Examples of isoQTLs are shown alongside scQTLs in **Fig. 4G** and presented as a Sashimi plot in **fig. S42** *(183).*

Lists of isoSNPs and isoGenes that passed filtering thresholds are available on the brainSCOPE portal, as described below. Text files for isoGenes in individual cell types contain columns for (a) isoGene name, (b) number of cis variants tested, (c) mean LD in the region, (d) top isoSNP ID, (e) p-value, (f-g) two fitted beta distribution parameters, (h) number of permutations, (i) permutation-based p-value, (j) beta distribution-based p-value, (k) runtime, and (l) Benjamini-Hochberg FDR correction value. Text files for isoGenes in individual cell types contain columns for (a) isoGene name, (b) isoSNP name, (c) F test statistic, (d) number of genotype groups, (e) maximum difference in relative expression difference among groups, (f-g) alternate transcript IDs, (h) number of individuals with missing, reference, heterozygous, and homozygous alternate genotypes, (i) nominal p-value, and (j) nominal FDR value.

**File:** [celltype]_permuted_fdr_correct.nominal05.txt: These files represent genes in cell-type-specific isoform usage QTLs (isoGenes) in 22 cell types (permuted beta distribution-derived p<0.05, with FDR values listed for each isoGene).
**File:** [celltype]_significant_sCQTLs_p0.05.txt (**data S16**): These files represent all SNPs associated with isoGenes (permuted beta distribution-derived p<0.05 filter) for cell-type-specific isoform usage QTLs (isoSNPs), filtered for isoGene-specific nominal p-value <0.05, in 22 cell types.

#### 4.5 *Allele-specific expression*

**Main manuscript reference:** Second supplementary reference in the third paragraph of "Determining cell-type-specific eQTLs from single-cell data."

We identified genes with allele-specific expression (ASE) in the 21 MultiomeBrain cohort samples to compare the effect sizes and allelic fraction of cell-type-specific eQTL variants and corresponding allele-specific expression eGenes, respectively (**Fig. 4F, fig. S49**).

The BAM output files were first converted to pseudo-bulk fastq files containing snRNA-seq reads corresponding to each cell type using bamtofastq v1.4.1

(https://support.10xgenomics.com/docs/bamtofastq). We then remapped the reads to personal genome fasta sequences constructed with vcf2diploid (*184*), and assessed allelic imbalance at each heterozygous SNV and gene using the AlleleSeq2 pipeline (*184–186*).

The full scASE files will be available for researchers with approved access within the PsychENCODE Consortium protected data portal. Anonymized files are available on the brainSCOPE portal as listed below.

**File**: MultiomeBrain_scASE_hetSNVs.tsv (protected) and MultiomeBrain_scASE_hetSNVs_anonymized.tsv: These files list hetSNVs that confer ASE in individual cell types, with haplotype-specific read counts.
**File**: MultiomeBrain_scASE_genes.tsv (protected) and MultiomeBrain_scASE_genes_anonymized.tsv: These files contain ASE genes with haplotype-specific read counts.

#### 4.6 ***mutSTARR-seq*** - Investigating the allelic effects of eQTLs on enhancers

**Main manuscript reference:** First supplementary reference in the third paragraph of "Determining cell-type-specific eQTLs from single-cell data."

We used a mutation STARR-seq approach (MutSTARR-seq) to investigate the allelic effects of eQTLs on activity of our putative enhancers. To do so, we first examined the overlap between our CapSTARR-seq results (see above section "STARR-seq analysis") and variants identified from scQTL analysis (see above section "scQTL analysis"). We prioritized 47 scQTLs and synthesized gene fragments with and without the scQTL variant to assess the enhancer activity of each fragment through STARR-seq. We found that four enhancer regions had significantly different enhancer activity with the eQTL variant, but only one of these regions remained significant following correction for multiple testing (Chi-squared test, adjusted p-value = 0.005). This region was chr17:45894107-45894607, and the alternate allele for this region showed increased enhancer activity. This region is predicted to target 12 different genes and has been previously implicated in several neurodegenerative diseases (*40*).

Lists of tested enhancers from our mutSTARR-seq experiments are available on the brainSCOPE portal in the following file:

**File:** starrseq_enhancers_merged_sig_qtls_fdr_0.05.bed: This file includes the panel design used in mut-STARRseq by intersecting STARR-Seq-validated enhancers with cell-type-specific eQTLs.

#### 4.7 ***STARR-seq and MPRA* Validation -** Validation of the scQTLs using STARR-seq and MPRA

**Main manuscript reference:** First supplementary reference in the fourth paragraph of "Determining cell-type-specific eQTLs from single-cell data."

##### Identifying the eSNPs

Our comparative enrichment analysis was carried out using eSNPs. The eSNPs are those variants that appear as part of significant scQTLs. In particular, these significant scQTLs are the primary set of QTLs identified for 17 cell types, processed with a standardized/conservative pipeline (available in the file "[celltype]_sig_QTLs.dat" on the brainSCOPE resource page).

##### Defining the active and control sets for STARR-seq and MPRA

(1) STARR-seq : We obtained candidate regions of STARR-seq with a fold-change score based on two replicates in primary human neural progenitor cells (phNPCs) isolated from the fetal cortex (*159*, *187*). Next, we obtained active enhancer regions in these two replicates using STARRPeaker *(188)*, totaling 6,202 regions for replicate 1 and 6,484 for replicate 2. For each replicate, we selected candidate regions with fold-change scores falling within the 25% quartile range as their respective control regions (**figs. S48A-B**). As a result, we obtained a total of 7,276 regions for both replicates.

(2) MPRA: We obtained the MPRA dataset of primary cells from the study by *(189)*. We used a more strict criterion to define the activity of MPRA by its RNA/DNA ratio. First, we categorized MPRA peaks into three groups: active enhancers (is_enhancer=1), silencers (is_silencer=1), and controls (is_enhancer=0 and is_silencer=0) based on their initial labels. Then, we computed the activity scores of MPRA using abs(RNA/DNA - 1). Next, we selected the 25% quartile of MPRA (6,221) in the control group as the control set. Additionally, we identified the MPRA peaks (6,575 peaks) with activity scores above the maximum active score in the control group as the active set (**fig. S48C**).

##### Enrichment analysis of the scQTLs in MPRA and STARR-seq

We randomly selected 6,000 peaks with replacements from the active set and control set in MPRA or STARR-seq and used BEDTools *(190)* to intersect the eSNPs dataset with both the active and control peak sets. We calculated the ratio of eSNPs located within the active and control sets. This process was repeated 2,000 times, allowing us to statistically test the enrichment of eSNPs in the active and control sets. The results show that eSNPs are more enriched in the active set than in the control set (**fig. S48D**).

### 5 Building a gene regulatory network for each cell type

#### 5.1 ***GRN construction*** - Construction of cell-type GRNs

**Main manuscript reference:** First supplementary reference in the first paragraph of "Building a gene regulatory network for each cell type."

We used snATAC-seq and snRNA-seq data from healthy individuals across all cohorts to predict proximal and distal regulatory links between TFs and their potential target genes. We used the metacell algorithm to identify homogeneous and robust groups of cells, which helped to reduce noise and variability in the snRNA-seq data (*191*). The values within the resulting metacell matrices were converted to CPM units and standardized to z-scores within each cohort. The resulting z-score matrices were combined and used for GRN construction using complementary approaches, as described below. We note that red blood lineage cells yielded very few metacells and were excluded from the GRN analysis. Additionally, Sst and Sst_Chodl cells were merged to improve statistical power. Our approach resulted in the identification of cell-type-specific GRNs, highlighting the utility of single-cell-resolution datasets for uncovering gene regulation patterns across cell types.

**SCENIC**

Single-Cell rEgulatory Network Inference and Clustering (SCENIC) *(58)* is a commonly used computational method for reconstructing GRNs and determining cell state from snRNA-seq. First, SCENIC uses cis-regulatory motif enrichment analysis (RcisTarget) on gene co-expression modules (which can be inferred from GENIE3 or GRNboost) to infer regulons. Given that gene co-expression modules may contain many false positives and indirect targets, RcisTarget then evaluates and identifies the modules in which the upstream regulator's binding motif is enriched across the target genes, resulting in regulons that consist only of direct targets. Lastly, the activity level of these regulons in each cell is quantified using an area under the recovery curve (AUC)-based enrichment score using AUCell.

For the present study, the SCENIC pipeline (R version) was applied to the metacell expression matrix. We first used GRNBoost2 in Python (*192*) to infer gene co-expression networks across all cell types. Then, we identified regulons with *runSCENIC_1_coexNetwork2modules* and *runSCENIC_2_createRegulons* functions with default settings. Specifically, the cisTarget motif enrichment analysis limited the regions for TF searching to a distance of 10 kbp centered on the TSS or 500 bp upstream of the TSS. Next, the revealed regulons (with at least 10 genes) were scored for their activity level in each cell using the *runSCENIC_3_scoreCells* function, and the AUCell enrichment was calculated based on the top 1% of the number of detected genes per cell. Lastly, we used the regulon specificity score (RSS) (*193*) metric to assess cell-type specificity of the discovered regulons. RSS, which is based on Jensen-Shannon divergence, is a measure of the difference in regulon activity across different cell types. For each cell type, we ranked the predicted regulons by their RSS values and selected the top 100 regulons with the highest scores. Finally, within each regulon, we sorted TF-target gene links based on the '*CoexWeight'* from GRNboost2 outputs and selected links with the top 20% scores to construct the cell-type GRNs.

**scGRNom**

A limitation of the SCENIC approach is that it does not take into account the dynamic and functional information that can be gained from snATAC-seq or open chromatin regions. snATAC-seq is a powerful tool for identifying accessible chromatin regions and can provide important information on gene regulation and functional genomic elements at a cell-type resolution. Without snATAC-seq data, the SCENIC approach may not be able to accurately predict gene expression patterns and identify regulatory relationships between TFs and target genes.

To compensate for this, we applied the scGRNom pipeline (*59*) to capture regulatory links within accessible promoter regions in the snATAC-seq data. Briefly, the scGRNom uses a *prior*, or reference network, depicting likely interactions between TFs and target genes based on TF binding site (TFBS) mapping within accessible or open promoters. However, under this scenario, multiple TFBSs can be mapped within a single promoter region, with potentially many of them having no regulatory effects. The scGRNom pipeline filters such links by estimating expression-based relationships between every target gene and all TFs linked to its promoter in the reference network. To do this, scGRNom uses elastic-net regression, a machine learning method that we have previously tested to model TF-target gene expression in bulk brain GRNs *(4)*. For the present analysis, we first mapped putative TFBSs in the JASPAR2020 database *(154)* within promoter peak regions in the snATAC-seq data to create reference networks for each cell type. Then, the reference network and normalized metacell expression matrix were supplied to the scGRNom function *scGRNom_getNt*, along with the following parameters: *cutoff_by = 'quantile', cutoff_percentage = 0.2,* and *train_ratio = 0.7*.

For subclass annotations, reference networks were replicated from the parent cell type or closely matching subclass cell type (for example, excitatory and inhibitory neuron reference networks used for L2/3 IT and Pvalb, respectively; microglia and endothelial reference networks used for immune cells and VLMC, respectively). We excluded *p*ericytes and *s*mooth muscle cells from this analysis to avoid using snATAC-seq peaks from biologically distinct cell types.

We merged high-confidence TF-enhancer-promoter interactions from snATAC-seq (top 100 TFs; intersection of all cohorts) with the union of top-scoring TF-target gene links from SCENIC (top 20% of links within the top 100 regulons) and scGRNom (top 20% TFs for each target gene) to construct merged GRNs for the 24 cell types. The 20% threshold was determined by evaluating SCENIC results on a benchmark derived from TF-promoter links in our ATAC-seq (see below section “GRN evaluation”). We discern activating and repressing edges by examining the sign of Pearson’s correlation between the metacell expression profiles of each TF-target gene pair. Note that we retain edge weights as RSS scores for SCENIC runs, absolute regression coefficient from scGRNom runs, and binary value of 0 or 1 depicting presence/absence of motif-enhancer-peak links in snATAC derived links. Finally, scQTLs were mapped to enhancers and promoters in the merged GRNs using the *findOverlaps* function of the GenomicRanges package (*194*).

We have constructed two sets of cell-type GRNs for the analyses in this paper: (1) GRNs with the pediatric samples (age < 13 years) filtered out and utilizing all the snATAC-seq data (**GRN-A**); (2) GRNs including the pediatric samples and using only the UCLA-ASD snATAC-seq peaks (**GRN-B**). The GRN-B set was used in the LNCTP (see section 1.8).

We provide the cell-type-specific GRNs and meta-data files related to their construction (including MetaCell expression values) on the brainSCOPE portal, as follows:

**File:** [celltype]_GRN.txt: These text files detail cell-type GRNs for 24 cell types (**GRN-A**). The files list the TF, enhancer location, promoter location, target gene, interaction type (proximal or distal), correlation, activating/repressing, edge-weight, and cell type.
**File:** [celltype].txt: These text files detail an alternate version of cell-type GRNs for the 24 cell types that were used as inputs for the LNCTP model (**GRN-B**). (These GRNs differ from the main files in that only UCLA-ASD snATAC-seq peaks were used to generate the GRNs and that samples with an age < 13 years were retained.)
**File**: SCENIC_RSS.csv (**data S20**): This file shows SCENIC-derived cell-type RSS (regulon specificity scores) for all TFs, used as inputs for constructing the final GRNs.
**File:** [celltype].eqtl_edge.txt (**data S21**): These files list the edges in the cell-type GRNs that link enhancer/promoter elements to scQTLs.

#### 5.2 *GRN evaluation*

We aimed to capture both distal and proximal regulatory links between TFs and target genes by integrating snATAC-seq and snRNA-seq data using complementary strategies. Proximal links between TFs and target genes (TF → promoter) were identified using two different approaches: the SCENIC pipeline and scGRNom. We reasoned that using only one technique would likely miss important regulatory interactions. The SCENIC pipeline captures cell-type-specific regulon activities from cell-type expression data but remains devoid of epigenomic data. The scGRNom pipeline fills this gap by utilizing a prior network built using enhancer promoter interactions in snATAC-seq data. However, utilizing snATAC-seq data from parent cell types for subclass cell annotations could limit scGRNom's ability to capture differences between certain sub-cell types. Therefore, taking a union of highly scoring TF-target gene links from both these methods could potentially construct a more robust GRN. Thus, we combined the most highly scoring proximal links from SCENIC and scGRNom with distal links from snATAC data to construct the cell-type GRNs. On average, each cell-type GRN comprises 387 TFs, ~9000 target genes, and 14,000 enhancers. These genes are connected via an average of 39,000 proximal and 38,000 distal links (**fig. S54**).

**Overlaps of SCENIC results with snATAC peaks**

As part of our integrative approach to construct cell-type-specific GRNs, we also mapped TFBSs to gene promoters using our snATAC-seq data. These binding sites were then used to evaluate the outputs of the SCENIC pipeline, which identifies cell-type-specific regulons based on gene expression data. To evaluate the accuracy of the SCENIC regulons for each cell type, we used TFBSs in ATAC peaks within gene promoter regions as a benchmark. We calculated precision as the proportion of SCENIC links that were also present in the snATAC-seq benchmark. Through our analysis, we found that the best precision was achieved when we selected the top 20% links based on the co-expression weight from GRNboost and the top 100 regulon ranks from SCENIC. We used this threshold to filter SCENIC runs in the merged GRNs. It is worth noting that we did not perform this evaluation for the scGRNom complement of the GRNs. This is because scGRNom uses TF-target gene promoter links from snATAC-seq data as a prior to build a GRN from snRNA-seq data. Instead, we chose a threshold of the top 20% targets for each TF based on our experience developing the algorithm and tests with other brain datasets (*59*, *195*).

Furthermore, we observed that the cell-type GRNs show patterns that reflect the known biology of cell-type relationships in the brain. For example, the GRNs of excitatory neuron subtypes were more similar to each other than to inhibitory neuron subtypes, and vice versa (**figs. S55B, S56**), which is not surprising given their distinct functions and developmental origins. This pattern was maintained even when the snATAC-seq links were removed from the GRNs, suggesting that the snRNA-seq data alone are sufficient to capture the cell-type relationships. Additionally, the stability of the pattern across different edge-trimming thresholds for both scGRNom and SCENIC indicates that the results are robust to variations in the analysis parameters (**fig. S55A).**

**Comparing cell type GRNs with tissue-naive GRNs**

We also compared these cell-type-specific GRNs to cell-type-naive regulatory networks available in the DoRothEA database *(196)*. The DoRothEA database provides a comprehensive list of TF-target gene interactions, scored on a scale from A to E, with A indicating the most reliable interactions supported by strong literature evidence, and E representing the least reliable interactions based on computational predictions. As anticipated, we observed only minimal overlaps with the DoRothEA network, with the majority of overlaps found in the E category of edges **(fig. S66**). This observation suggests that bulk GRNs are insufficient in capturing the intricacies of cell-type-specific regulatory signals.

**Variance in gene expression explained by GRN models**

Gene expression is a product of regulation, which includes TFs as regulators. Regulators often contribute to explaining a certain amount of the observed variability in gene expression. We formulate a network-based regression method to estimate the percentage of variation in gene expression explainable by our cell type GRN models. Operationally, for each target gene, we fit a linear regression model using the expression levels of all linked regulators (TFs) as predictors, weighted by their respective edge weights (Pearson's correlation). The percentage of variation explained by the regression model is then obtained by comparing the amount of variability captured by the model (SS_residual) with the total variability in the data (SS_total). These analyses indicate that the GRNs account for approximately 52% of the variance in gene expression on average (**fig. S62**). Notably, when focusing solely on TF→TG edges mediated through enhancers or promoters, this percentage decreases by 5-10%, suggesting that merging enhancer and promoter links into a unified GRN substantially contributes to capturing more accurate GRN models.

Overall, these tests allowed us to evaluate the stability and robustness of our cell-type GRNs and ensure their accuracy in capturing both distal and proximal regulatory links. Our findings provide confidence in the accuracy and biological relevance of the constructed GRNs.

**GRN stability**

We assessed the stability of these cell-type GRNs across three random splits of the CMC cohort. In each split, we randomly selected 80% of the donors within the cohort and independently applied our GRN inference pipeline. Note that it was not possible to create two equal and non-overlapping splits of any cohort because our GRN pipeline uses the metacell algorithm to normalize each cohort and integrate them within a Z-score space. Applying the metacell pipeline to too few individuals yields very low power to run the GRN inference pipeline. Thus, we decided to use three splits of 80% randomly chosen donors from the CMC cohort.

Within this framework, the regulon specificity scores (generated by SCENIC) from the splits are substantially correlated (**fig. S57**). We also found that the average overlap between the hub TFs is ~71% across the three splits (**fig. S58**). This level of agreement between the splits exceeds what has been observed in similar analyses of bulk datasets (*100*).

#### 5.3 ***CRISPR validation*** - Validation of TFs and target genes identified in GRNs

**Main manuscript reference:** First supplementary reference in the second paragraph of "Building a gene regulatory network for each cell type."

We overlapped the predicted regulatory genes and enhancers identified from cell-type-specific GRNs with psychiatric disease-associated active enhancers and their nearby genes *(32)*. We selected four enhancers that have the same predicted target gene for further functional validation: *NGEF, RORA*, *PLEKHO1*, and *TOML2*. These enhancers were knocked out in primary human NPCs through ribonucleoprotein-mediated CRISPR/Cas9 genome editing in two biological replicates. DNA genotyping and Sanger sequencing were used to confirm the deletion of the enhancers at the expected loci. TaqMan quantitative PCR (qPCR) was used to examine the relative expression level change of target genes before and after enhancer knockout (KO). Detailed protocols for the CRISPR/Cas9 assays are available in the PsychENCODE Consortium publication by Gaynor and colleagues *(32)*.

We validated the CRISPR/Cas9 deletion of candidate enhancers with Sanger sequencing *(32)*. After enhancer KO, Taqman qPCR assays showed that the average expression of the target gene was diminished after enhancer KO. In particular, the relative expression level of *NGEF* decreased to 0.45 (SD between biological replicate 1 and biological replicate 2: ± 0.01, after EH37E1198822 KO), *RORB* decreased to 0.56 (SD: ± 0.2, after EH37E1000386 KO), *PLEKHO1* decreased to 0.16 (SD: 0.02, after EH37E0114246 KO), and *TOM1L2* decreased to 0.42 (SD: ± 0.01, after EH37E0426064 KO). These results demonstrate that these active enhancers regulate transcription of the target gene tested.

Additional data related to these experiments are located in the following file on the brainSCOPE portal:

**File:** crispr_validation_results.xlsx (**data S22**): This file details expression of the target genes before and after CRISPR KO of the linked enhancers, as predicted by peak2gene linkages.

#### 5.4 ***Network Characterization*** - Comparison of GRN structure across cell types

**Main manuscript reference:** First supplementary reference in the third paragraph, and first supplementary reference in the last paragraph of "Building a gene regulatory network for each cell type."

##### Centrality analysis

We used three measures of network centrality to assess the importance of TFs in a directed cell-type GRN. Specifically, we computed the in-degree and out-degree of a given TF in a GRN, which represent the number of incoming and outgoing edges, respectively. We also calculated the betweenness centrality by counting the number of shortest paths that pass through a TF among all possible shortest paths. These centrality scores were sorted in decreasing order, and we selected the top 5% of TFs based on out-degree, in-degree, and betweenness centrality. From these lists, we identified pure out-hubs, which are TFs that solely exhibit out-hub behavior and are not present in the top decile of the other two centralities. We identified pure bottlenecks and in-hubs in a similar fashion. These lists of TFs are shown in **data S23**. The centrality scores were calculated using the igraph package (https://igraph.org/) in R. All hubs and bottlenecks are located in the data files referenced below and available on the brainSCOPE portal:

**File:** inhubs.txt (**data S23**): This matrix lists whether a TF was identified as an in-hub (1) or not (0) in each cell type.
**File:** outhubs.txt (**data S23**): This matrix lists whether a TF was identified as an out-hub (1) or not (0) in each cell type.
**File:** bottlenecks.txt (**data S23**): This matrix lists whether a TF was identified as a bottleneck (1) or not (0) in each cell type.

##### Comparison of cell-type regulons with disease co-expression modules from bulk data

Our cell-type GRNs link TFs to their potential target genes based on co-expression relationships in scRNA-seq data and binding sites identified in scATAC-seq data. We aimed to investigate whether the group of target genes associated with a particular TF, or regulons, shares overlap with co-expressed disease gene modules previously identified in bulk data. Our specific interest focused on investigating whether genes within cell-type regulons exhibit co-expression behaviors that are disease specific.

To test this hypothesis, we obtained disease co-expression modules previously identified from bulk data (*100*). We then conducted statistical enrichment analysis of these bulk disease modules within cell-type regulons (generated by SCENIC) using hypergeometric tests and counted the number of regulons with significant p-values. We observed that non-neuronal regulons are particularly enriched for ASD and depleted for bipolar disorder **(fig. S65)**. Additionally, certain excitatory neuronal regulons, such as L6.IT, showed relatively higher enrichment for schizophrenia. Overall, this analysis demonstrates the utility in identifying potential regulators of disease modules, information not readily extractable from co-expression modules.

**GO enrichment analysis**

We obtained the human GO biological process annotations that were propagated along 'is_a' and 'part_of' relationships from a previous study (*197*). These annotations were filtered to retain only those terms that annotated between 5 and 1,000 genes. We then applied the resulting set of GO biological process terms for an enrichment analysis. To determine the statistical enrichment of a list of query genes, such as bottleneck genes, within a given GO biological process term, we used hypergeometric tests. The background for these tests consisted of all genes present in the corresponding GRNs. We report only those enrichment tests with an FDR of less than 0.1.

GO enrichment results for identified bottleneck genes are available as a file on the brainSCOPE portal:

**File:** GO_enrichment_bottlenecks.txt (**data S24**): GO enrichment among bottleneck genes in cell-type-specific GRNs. Columns represent GO terms, p-value, FDR, signature, gene set count, overlap count, background, count cell type, TF, and enrichment score.

**Motif analysis**

In order to identify particular structural patterns in the cell-type GRNs, we conducted a motif analysis. Specifically, we examined triplets of TF subgraphs that were grouped into 13 isomorphic classes. To determine the frequency of each class in each GRN, we used the Mfinder tool (*198*). We used default parameters for the analysis, with the exception of the number of random networks (r), which was set to 1,000. From these results, we calculated the z-score of the distribution.

**Disease co-regulatory networks**

The cell-type GRNs that we generated are directed and unweighted graphs with edges connecting TFs to target genes. However, in order to investigate the co-regulation strength among known disease risk genes (which include non-TF genes), we converted these directed GRNs into undirected networks that also include connections between target genes. To accomplish this, we calculated the overlap between predicted TFs for every pair of target genes within each GRN, quantifying the overlap using the Jaccard Index. The Jaccard Index values of each cell type were then used to populate a gene-gene adjacency matrix $C$, where each entry $C_{ij}$ represents the level of coregulation between gene $i$ and gene $j$. We then subsetted matrix $C$ to contain only disease risk genes, and calculated the weighted density of the resulting disease subnetwork (*199*). This procedure is illustrated in **fig. S69.**

To determine the statistical significance of these observations, we estimated a pseudo-p-value for each gene set-disease combination by randomly selecting gene sets from the background of all genes present in $C$, and counting the number of times the co-regulation score of the randomly selected gene sets was greater than or equal to the observed score in 999 independent trials. For our calculations, we obtained ASD risk genes from the SFARI gene database (*171*) (release 02-02-2023) and filtered for high-confidence genes (score = 1). Schizophrenia, AD, and Parkinson’s disease risk genes were extracted from the DisGeNet database and screened for manually curated sources (source = "CTD_human" or "GWASCAT"). A set of housekeeping genes was used as a control.

**Modular organization of cell-type GRNs**

We were also interested in understanding the modular organization of the cell-type GRNs, as functionally similar disease genes often tend to cluster and function as modules. To identify clusters or modules of co-regulated genes within each cell type, we used the Louvain clustering algorithm (with the modularity parameter set to 1) on the gene-gene adjacency matrix *C* (discussed above). On average, we found 10 modules per cell type, with a median of 562 genes per module and 8,280 genes in total. These modules are summarized in **fig. S68.**

We used the normalized mutual information (NMI) as a metric to gauge the similarity between these cell-type modules and disease modules previously identified in bulk data (*100*). Under this framework, we found that all cell-type modules in our dataset are relatively more similar to bipolar disorder-related modules compared to ASD and schizophrenia. In conclusion, our analysis shows that cell-type GRNs are highly modular and suggests that disease genes are more likely to cluster in a cell-type-specific manner.

We have listed the cell type-specific module assignments of genes in the following file available on the brainSCOPE portal:

**File:** Gene_module_mappings.csv: This file lists genes in the first column, module memberships in the second column, and cell types in the third column. The modules represent sets of co-regulated genes. Users are advised to filter modules that are too large and too small for downstream analysis.

#### 5.5 *Unifying TF-target Regulons*

**Main manuscript reference:** Second supplementary reference in the first paragraph of "Building a gene regulatory network for each cell type."

The brainSCOPE portal provides access to cell-type GRNs. In addition, we created a unified GRN by merging all cell-type GRNs. This GRN comprises 587,385 links, with 26.9% of which are manifest in only one cell type **(fig. S64)**. The unified GRN is similar but not the same as a bulk GRN in that it weighs rare cell types more highly. This additional GRN resource presents opportunities to develop tractable solutions for retrieving disease genes regulated across the brain broadly, offering a more streamlined approach to managing extensive lists of disease genes and reducing complexity. Note that analyzing disease gene rewiring within this pseudo-bulk GRN will inevitably result in the loss of cell-type-specific signals and thus is not an explicit focus of this paper.

To develop a more sophisticated way of relating a set of input genes to their upstream regulators, we have also applied a network diffusion method. This method, given a target gene, provides the key regulators—specifically, the aggregate regulation score of each TF for that target. This approach allows us to chain together larger networks involving multiple TFs, surpassing simple combined regulons.

To illustrate the relationship between TFs and their targets, utilizing the all-inferred TF-target relations in cell-type-specific GRNs along with their relations in unified GRN, we applied MultiXrank, a random walk with restart (RWR) algorithm capable of utilizing multilayer graphs *(200)*. We individually applied the algorithm to each cell-type-specific GRN and to the integrated GRN, consisting of 24 layers composed of cell-type-specific GRNs, using default parameters. We calculated the diffusion scores by running the algorithm for each TF as a starting point in the RWR algorithm for a given GRN. Thus, the resulting diffusion scores represent the TF-target relations utilizing the topology of either cell type-specific GRN or multilayer-GRN based on the input graph. We used arithmetic averages to aggregate the diffusion scores across multiple layers when we applied the MultixRank algorithm to the unified multilayer GRN.

Overall, we have generated an easy-to-use data file containing diffusion scores for each TF-target relationship for the unified GRN and cell-type-specific GRN, wherever such a relationship exists **(fig. S59)**. **fig. 60** demonstrates to users how to identify the upregulators for an example gene, *EGFR*. The user can identify the top regulators of *EGFR* gene in the cell-type-specific GRNs (**fig. S60A**) and in the unified GRN, where cell-type specificity is absent (**fig. S60B**).

Text files for the unified GRN and diffusion scores are available on the brainSCOPE portal:

**File:** unified_GRN.txt: This matrix lists TF-target links in the unified GRN derived by taking a union of all cell-type GRNs. Links found in a cell type are marked as 1, and are otherwise marked as 0.
**File:** unified_GRN_diffuision.txt: This file lists diffusion scores between TFs and target genes for the unified GRN and for each cell type-specific GRN. Users can easily search for a relevant downstream gene (such as those related to a DE experiment) as a row in the file to find all the regulatory scores for upstream TFs.

### 6 Constructing a cell-to-cell communication network

#### 6.1 ***Cell-to-Cell Network*** - Methods to build cell-to-cell networks

**Main manuscript reference:** First supplementary reference in the first paragraph, and first supplementary reference in the last paragraph of “Constructing a cell-to-cell communication network.”

The advent of single-cell transcriptomics provides us the opportunity to better deconvolute molecular and cellular signaling interactions in health and disease. Here, we applied the standard workflow of CellChat (v1.5.0) utilizing single-cell gene expression of ligands and receptors to infer a cell-to-cell communication network (*64*). The normalized count matrix for the SZBDMulti-seq dataset was used in our cell-to-cell analysis (C2C analysis), with the required cell-type annotations coming from the ‘subclass’ metadata column. The number of inferred ligand-receptor pairs depends on the method for calculating the average gene expression per cell group. We used the default CellChat robust mean method called ‘trimean’, with the cutoff ‘trim’ parameter set to the default value of 0.1. Three separate analyses were done for each of the psychiatric conditions in the SZBDMulti-seq dataset, namely control, bipolar disorder, and schizophrenia.

Next, we moved from individual cell-to-cell communication analyses to differential analyses between conditions. We began our analyses by normalizing the three-dimensional matrices (sender cell types x receiver cell types x ligand-receptor interactions) in each of the CellChat objects so that all matrices had the same sum to account for batch effects in the snRNA-seq data. For example, we found upregulation of the MIF ligand-receptor pathway in smooth muscle cells in schizophrenia. We then used the advanced computation and pattern recognition approach non-negative matrix factorization (NMF) on the snRNA-seq data to compare cell-type signaling differences between patients with psychiatric disorders and controls. Of note, we opted to not use the default Lee-Seung method for NMF optimization, which was first applied to two-dimensional images, but instead use the Brunet algorithm, which was developed specifically for biological data (*201*). Moreover, we used random seeding and 200 runs for our optimization parameters, and set the rank to three patterns. Finally, a differential circle plot for each disorder versus control was generated, with nodes colored by input for the overall signaling difference, while individual chord plots were generated to visualize the individual pathway differences.

Finally, we confirmed our findings with another C2C analysis software package named NicheNet (v1.1.1) (*79*). NicheNet inputs were as follows: sender cells included astrocytes, endothelial cells, immune cells, microglia, oligodendrocytes, oligodendrocyte precursor cells, pericytes, smooth muscle cells, and vascular leptomeningeal cells; receiver cells included all the neuronal cell types. Target risk genes were those identified from PsychENCODE Consortium analyses (*172*, *173*). The following filters were used in the NicheNet analysis: ‘n_ligands’ as 20, ‘n_targets’ as 400, and ‘cutoff’ as 0.25. The top 10 ligands and 15 target genes were then selected for our downstream C2C analysis.

Cell-to-cell communication networks per control and diseased individuals, represented as CellChat and NicheNet objects as well as summarized text files listing ligand-receptor patterns per cell type, are available as RDS and h5 files on the brainSCOPE portal as follows:

**File:** cellchat_C2C_network_[disorder].txt (**data S25**): These files contain sets of ligand-receptor signaling patterns across cell types in control, schizophrenia, and bipolar disorder individuals. Files list all interactions between ligand-receptors in different cell types, along with the strength of interaction and annotations for interaction type and pathway.
**File:** cellchat_C2C_network_netP_[disorder].txt (**data S26**): These files contain sets of ligand-receptor signaling patterns across cell types summarized by signaling pathway in control, schizophrenia, and bipolar disorder individuals.
**File:** SZBD-Kellis_annotated-CON_cellchat.rds.gz: Network of ligand-receptor signaling patterns across cell types in control individuals.
**File:** SZBD-Kellis_annotated-SZ_cellchat.rds.gz: Network of ligand-receptor signaling patterns across cell types in schizophrenia individuals.
**File:** SZBD-Kellis_annotated-BD_cellchat.rds.gz: Network of ligand-receptor signaling patterns across cell types in bipolar disorder individuals.
**File:** SZBD-Kellis_annotated-BD_CON.h5seurat: Changes in cell-to-cell networks between individuals with bipolar disorder and control individuals, represented as a NicheNet Seurat h5 object.
**File:** SZBD-Kellis_annotated-SCZ_CON.h5seurat: Changes in cell-to-cell networks between individuals with schizophrenia and control individuals, represented as a NicheNet Seurat h5 object.

#### 6.2 ***Cell-to-Cell Network*** - Methods to determine latent patterns

The patterns in the NMF (Non-negative Matrix Factorization) can be thought of as broad clustering of cell types and signaling pathways. Simultaneously, cell types were broadly clustered into the same pattern if they share the use of certain signaling pathways; signaling pathways were broadly clustered into the same pattern if they share the same sender cell types.

There are three patterns, both biologically and mathematically based. Biologically, it is known that there is a broad typing of excitatory, inhibitory, and non-neuronal cell types. We chose three categories and determined whether our 24 cell types could be properly divided among the broad categories. Mathematically, we calculated silhouette and cophenetic scores, implemented in the NMF R Package (*202*), using default parameters. We determined that rank 3 was the most stable. For both of these scoring metrics, the higher the score, the better their reproducibility across repeated experiments. We illustrate this in **fig. S72**.

#### 6.3 ***Cell-to-Cell Network*** - Validation of cell-to-cell network with spatial data

To perform validation for our communication network, we emphasized the importance of spatial distance in ligand-to-receptor gene signaling. That is, the cell type expressing the ligand gene (sender cell type) must be “close” in proximity to the cell type expressing the receptor gene (receiver cell type) in order for cell-to-cell communication to occur. The farther apart the cell types are, the less likely they are communicating.

From ten neurotypical controls, spatially resolved transcriptomics data were generated using 10x Genomics Visium across the anterior, middle, and posterior DLPFC (n = 30). To deconvolute the cell types in the spatial data, we generated snRNA-seq data of these tissue blocks using 10x Genomics Chromium (n = 19). Raw cell-type deconvolution was performed by multiple teams of PsychENCODE (led by Keri Martinowich, Leonardo Collado-Torres, Kristen Maynard, and Stephanie Hicks) (*203*). To obtain the normalized cell-type values, we grouped by sample ID and then divided by the column sums (where columns are raw counts for each spot for each cell type). We show the spatial locations of specific snRNA-seq-labeled cell types, with layer specificity in **fig. S70**.

Due to the difference in annotations between the spatially resolved data and our own data, pooling of a few cell types was required. Specifically, because inhibitory cell types had no informative PFC layer registration (per the spatial team above), all inhibitory clusters were merged into a single cluster. Endothelial cells, smooth muscle cells, and pericytes were further merged into the endomural cluster. L5 IT and L5 ET cells were collectively labeled as excit_l5, while L6 IT, L6 IT Car3, L6 CT, and L6b cells were labeled as excit_l6 (**Table S11**). After such merging, only the common cell types (11/13) were kept for validation of our communication network. After harmonizing annotations between the spatial data and our own data, we calculated the spatial distances among all pairs of cell types.

We performed three correlations between the spatial distance matrix (obtained from our spatial data) and the communication matrix (from our original analysis). We found that Pearson, Kendall, and Spearman correlations gave corresponding coefficients of -0.179, -0.158, and

-0.241 at p-values of 0.06, 0.01, and 0.01, respectively (**Table S12**). The negative correlation values of all three tests validate the spatial requirement of our communication network.

### 7 Assessing cell-type-specific transcriptomic and epigenetic changes in aging

#### 7.1 ***Aging Cell Fractions*** - Single-cell aging cell-type fraction

**Main manuscript reference:** First supplementary reference in the first paragraph of "Assessing cell-type-specific transcriptomic and epigenetic changes in aging."

To characterize single-cell cell-type fractions by age, we used single-cell cell-type fractions from our harmonized cell annotation scheme. We used control samples from the CMC cohort in this analysis. We modeled the association between cell fractions and age using a generalized linear model with biological sex and genotype ancestry as covariates (**data S27B**). Outliers were removed from the analysis if they were smaller or larger than 1.5 IQR.

Scatter plot in **Fig. 7A** (bottom) shows the cell fraction change over age in OPCs (grey) and Chandelier cells (blue) with best-fit lines showing the change trend. These two cell types showed significant decreases across age in the bulk RNA-seq deconvolution data (see section 7.2 for details) and single-cell annotations (FDR<0.05, two-sided t-test).

#### 7.2 ***Aging Cell Fractions -*** Deconvolution of bulk-RNA-seq data

**Main manuscript reference:** First supplementary reference in the first paragraph of "Assessing cell-type-specific transcriptomic and epigenetic changes in aging."

We used quantile-normalized bulk RNA-seq datasets available for >800 individuals in the ROSMAP cohort (*17*), available on the AMP-AD Knowledge Portal *(116),* to calculate cell-type fractions based on age and for downstream analysis in modeling AD status. Transcripts for each sample were quantified from aligned RNA-seq BAM files, using the *htseq-count* command in the HTSeq software package (*204*) and transcript annotations from GENCODE v87 gtf files. We used these individual-level files to generate a bulk RNA-seq expression matrix, and collected this matrix and snRNA-seq cell-type annotations from available ROSMAP samples as inputs for the BisqueRNA software, which infers cell-type fractions of the bulk RNA-seq data (*144*). The single-cell raw counts were first log-normalized, while the bulk-seq raw counts were quantile normalized, before being input into BisqueRNA for AD patient and control samples, respectively. Signature matrices for different cell types generated by BisqueRNA were used for downstream analyses.

We have exhibited the correlation between cell-type fractions and the aging process through the bulk deconvolution results focusing specifically on data from the CMC cohort. The alterations in the fractions of various cell types as revealed by bulk deconvolution align with the cell fraction estimations obtained from CMC's snRNA-seq data (**data S27**)**.** This concurrence is further supported by previous reports; for instance, cell types like Chandelier cells and OPCs demonstrate a significant decline (FDR<0.05, two-sided t-test) as individuals age *(82, 83).*

#### 7.3 ***Aging DE*** - DE genes in control aging and schizophrenia aging

**Main manuscript reference:** Second supplementary reference in the first paragraph of "Assessing cell-type-specific transcriptomic and epigenetic changes in aging."

For the aging DE gene calculation, we focused on the SZBDMulti-seq and CMC cohorts that were balanced over age as well as between healthy/schizophrenia disease groups.

**Aging DE genes in control group**

For the control aging DE gene analysis presented in the main text, we only used healthy control samples, which we further split into young (25-70 years) and old (70-90 years) individuals. The number of old and young individuals are balanced. Pseudo bulk gene expression per cell type per individual was generated through the sum of the raw gene counts. The filtering strategy is the same as the "DE Analysis" section (see supp."DE Analysis"). After filtering, the DEseq2 (likelihood ratio test) standard pipeline (*23*) was used for DE gene calculation with cohort, biological sex, genotype-derived ancestry, PMI, and average cell UMI count as covariates. Contrasts were made between the old and young groups. Multiple testing corrections were performed, and genes with an adjusted p-value<0.05 were defined as differentially expressed between contrast conditions.

**Aging DE genes in the schizophrenia patient group**

For aging DE genes in the schizophrenia patient groups, we only used disease individuals from the SZBDMulti-seq and CMC cohorts. We further split the samples into young (25-70 years) and old (70-90 years) individuals. The numbers of old and young individuals in the schizophrenia patient groups were balanced. The total number of samples used in this analysis was comparable to the number of individuals used in the control aging analysis (**fig. S19A**). The calculation followed the same aging DE gene pipeline mentioned above. Compared with the control aging group, only a small number of aging DE genes were identified in the schizophrenia patient group (**fig. S19A**).

To further validate the observations, we conducted a permutation analysis, randomly excluding five samples from each comparison group (control old, control young, schizophrenia old, schizophrenia young. **fig. S19B**) Despite the reduced sample size, a consistent pattern emerged: the schizophrenia patient group exhibited fewer aging DE genes compared with the control group, reinforcing our findings (**fig. S19B**).

In addition to the permutation analysis, we explored technical and biological covariates that might potentially affect the results. We explored the distribution of the number of cells per individual and the UMI count. We did not observe a substantial change in the number of cells between different groups. We do observe a difference in UMI per individual among the groups. Differences in the UMI among groups could arise for biological or technical reasons; thus, we included this as a covariate when we characterized the aging DE genes.

**Aging DE genes and AD DE genes**

We also compared our healthy aging DE gene list with AD DE genes of the PFC by their $\log_2$ fold change across distinct brain cell types. The AD DE gene list, published by Mathys, Hansruedi, et al. (2019), highly overlaps with samples in the ROSMAP cohort (*115*). **Data S28** lists an inner join of both sets of DE genes with their $\log_2$ fold change in AD and aging.

The DE genes are available on the brainSCOPE portal. Each file contains the gene name, average expression, $\log_2$ fold change, standard error, test statistic, p-value, and FDR-corrected p-value among older individuals.

**File**: Aging_DEGcombined.csv: Sets of DE genes in older and younger individuals for 20 cell types.
**File**: Aging-schizophrenia_DEGcombined.csv: Sets of DE genes in older and younger individuals in Schizophrenia group for 20 cell types.

#### 7.4 ***Aging STEM*** - Short Time-series Expression Miner analyses

Short Time-series Expression Miner (STEM) is a Java application designed to cluster, compare, and visualize short time series gene expression data derived from microarray experiments, typically involving eight time points or fewer (*205*). Researchers can leverage STEM to identify significant temporal expression patterns and pinpoint the genes linked to these patterns.

Note that STEM methods have been used in a variety of publications involving single-cell analyses *(206, 207)*. If one has a "hard-coded" time (instead of pseudotime from a trajectory analysis), STEM is essentially the same as the newer method scSTEM *(208)*, which aims to analyze time series data by pseudotime point. Since we are focusing on continuous change of gene expression with age within each individual cell type, the original STEM approach is more appropriate.

Samples from SZBDMulti-seq are normalized by CPM and then grouped into six different age groups (30-40, 40-50, 50-60, 60-70, 70-80, and 80-90 years old). In this algorithm, pseudobulk gene expression time series data were clustered to different model profiles to which its time series most closely matched based on the correlation coefficient. We successfully identified multiple groups of genes that align with various continuous model profiles for each cell type. Notably, these profiles capture the continuous nature of gene expression changes over time. We believe that incorporating STEM analysis enriches our understanding of the dynamic interplay between gene expression and aging. Detailed information about the short time series models (by STEM software) for each cell type, as well as the genes associated with specific model profiles, can be accessed in **fig. S74** and the brainSCOPE portal.

**File:** [celltype].txt: Sets of differentially-expressed genes for sample age derived from STEM analysis for 17 cell types.

#### 7.5 ***Aging Model*** - Aging prediction using prioritized genes

**Main manuscript reference:** First supplementary reference in the second paragraph of "Assessing cell-type-specific transcriptomic and epigenetic changes in aging."

In our study, we account for the fact that individual variability might manifest in the number and types of cells present. Recognizing that certain cell types might only be prevalent in a subset of individuals, our initial step involves filtering out cell types that are present in less than 70% of the population. This QC step results in 11 distinct cell types. We then sought to establish a sample pool wherein all 11 cell types are uniformly present. This resulted in an intersection that incorporated approximately 180 individuals. For our predictive modeling, we used the XGBoost regression framework, crafting a model for each of the identified cell types to predict age across each of our sample pools. Given the importance of age distribution, we applied stratified sampling for our train-test split. Specifically, we grouped ages using a bin width of 10 years, ensuring that both training and test samples were proportionately drawn from each bin. This culminated in an 80-20% train-test division. To gauge the efficacy and precision of our models, we implemented multiple evaluation metrics: Pearson correlation, Spearman correlation, mean absolute error, and root mean squared error. The XGBoost models were trained using three-fold cross-validation, adopting the negative mean squared error as the scoring function. For robustness, this train-test split process was repeated ten times, and we present both the mean and SD of our results.

To derive deeper insights from our model predictions, we incorporated the SHAP (SHapley Additive exPlanations) method. SHAP values, rooted in cooperative game theory, offer a principled framework to interpret and attribute the contributions of individual features to machine learning predictions. In our age prediction, a positive SHAP value for a specific feature indicates its role in driving predictions towards higher ages, while a negative value suggests the opposite. By analyzing these values, one gains a clearer understanding of the factors influencing age predictions, potentially revealing cell-type-specific age expression markers that correlate with the aging processes. Specifically, we spotlighted the top ten genes characterized by the highest mean absolute SHAP values for L2/3 IT and oligodendrocytes, showcasing them in **fig. S76**. We also visualized the RNA expression of the SHAP prioritized genes, as shown in **Fig. 7C**. Finally, we provide both summarized results and the full set of results from the SHAP model, along with source code used to run the model, on the brainSCOPE portal:

**File:** [celltype]_shap_summary_stratify.csv: These files contain age prediction gene prioritization results per cell type, based on the SHAP model. The first column lists gene or covariate (cohort, disorder, ancestry, or biological sex), and the second column lists the SHAP score.
**File:** shap_xgboost.tar.gz: TAR file containing source code and full results for SHAP gene expression prediction model for aging.

#### 7.6 *Aging Chromatin*

**Main manuscript reference:** First and second supplementary references in the third paragraph of "Assessing cell-type-specific transcriptomic and epigenetic changes in aging."

In order to determine cell-type-specific aging open chromatin regions, we first compiled bulk ATAC-seq samples from 628 individuals. Next, we deconvolved their signals to cell-type-specific ATAC-seq peaks from snATAC-seq datasets. This was done by using bigwigaverageoverbed to obtain the average signal per individual represented on each cell-type-specific peak. Next, we performed a PCA for each cell type, retaining 50 PCs to dimensionally reduce the matrix consisting of cell-type-specific peaks and signals from the 628 individuals. We further used t-distributed stochastic neighbor embedding for additional dimensionality reduction, and reduced the matrix to two dimensions. We performed k-means clustering on the resulting two-dimensional embedding to find distinct clusters. Each point is colored based on age, showing that there are differences in age for the different clusters identified. Overall, we found that oligodendrocytes and microglia show the highest stratification of age based on clusters from the embedding of ATAC-seq peaks.

The following files that list the integrated ATAC-seq peaks and associated meta-data are available on the brainSCOPE portal:

**File:** bwaob_output_col5_[cell-type].matrix: These matrices list open chromatin peaks generated from the deconvolution of bulk ATAC-seq datasets signal to cell-type-specific signals for seven cell types.
**File:** bulk_ATAC_samples_ordered.cleaned.txt: This file lists the column (sample) names for the deconvoluted signal matrices.
**File:** pec_atacseq_metadata_08142021.csv: This file lists the biosample metadata for samples in the bulk ATAC-seq dataset.

#### 7.7 ***AD Model -*** Associating cell-type fractions and signatures with AD

**Main manuscript reference:** First and second supplementary references in the last paragraph of "Assessing cell-type-specific transcriptomic and epigenetic changes in aging."

##### Inference of cell-type-specific gene expression and methylation

Once we obtained the cell-type fraction for each bulk sample (see above section "Aging cell fractions," we used the bMIND software package (*91*) to infer cell-type-specific gene expression and cell-type-specific methylation for each sample. The DEGs from bulk-RNA-seq and snNA-seq among ROSMAP individuals were selected for cell-type-specific expression inferences; the differential methylation regions were selected for cell-type-specific methylation inferences. Methylation datasets for 740 ROSMAP individuals (*17*) were downloaded from the AMP-AD Knowledge Portal (*116*) and integrated with the bulk- and snRNA-seq analyses to build a predictive model.

##### Building a model to predict AD

First, we explored the cell-type-specific associations between AD phenotype, defined as AD case and control, and cell-type fractions. P-values for changes in each cell-type fraction were calculated based on a one-tailed student's t test between the control and AD group. Then, we studied the contribution of gene expression and methylation features from each cell type for the prediction of AD status using RandomForest and multiple layer perceptron (MLP) deep learning models. The area under the precision recall curve (AUPRC) was used to evaluate the performance of cell-type features.

##### Multilayer perceptron

The MLP we used is a neural network model composed of two dense layers. The first layer applies a rectified linear unit activation function, while the second layer uses a sigmoid activation function to produce the final output. The input is flattened into a one-dimensional vector before being fed into the first layer. To evaluate the performance of the model, we performed 25 experiments (total of 5 runs with corresponding seeds, where each run has its own 5-fold cross-validation based on the corresponding seed). We split the data using a 3:1:1 ratio for training:validation:testing subsets. We explored various model configurations of different inputs, labels, and parameters, and provide the most important ones here. To enhance the performance, we integrated the MLP model using the Adaboost method. We used the MLP classifier as the base estimator. During training, we adjusted the weights of the target classes for each MLP classifier based on the previous classifier's performance. Specifically, we assigned a higher weight to the class on which the previous classifier made more mistakes. The final result is the weighted output of the 25 MLP classifiers, with each classifier's weight based on its performance in the previous training.

The following output files related to the AD prediction model are available on the brainSCOPE portal:

**File:** hybrid.admodel.auprc.txt (**data S29**): Text file containing AUPRC values for cell types and data modalities (rf.meth=methylation, rf.expr=expression) from AD model predictions.

### 8 Imputing gene expression and prioritizing disease genes across cell types with an integrative model

#### 8.1 ***LNCTP Priors*** - Linear Network of Cell-Type Phenotypes imputation priors

**Main manuscript reference:** Second supplementary reference in the first paragraph of "Imputing gene expression and prioritizing disease genes across cell types with an integrative model."

To constrain the cell-type expression imputation process, we used the processed snRNA-seq cohorts to quantify priors on the expression values for each gene in each cell type. In doing so, we had to contend with inter-cohort batch effects, as well as the inherent sparsity of snRNA-seq data. The strategy used thus followed these steps:

1. For each cell type in each individual in every cohort, we used the program MetaCell-2 (*209*) to generate “metacells," which aggregate counts across several cells in close proximity (based on expression similarity) to each other. The motivation is to strike a balance between the need to reduce the sparsity of the expression vectors at the individual cell level, and the need to retain some measure of heterogeneity in the cell-type expression values. If we took a simple pseudobulk across all the cells in a given type, we could lose information about cell-type heterogeneity.
2. The gene expression values in each individual were then converted into CPM units, averaged across all meta-cells in that individual and cell type, and subsequently converted into z-scores.
3. The z-score distribution for each cell type across all individuals and study cohorts was used as a prior in the deep learning framework. Specifically, the mean and SD in the z-scores were calculated and incorporated into the corresponding Gaussian Markov random field (GMRF) terms.

The underlying assumptions in this procedure are that: (a) metacells maintain a representative measure of the intra-cell-type heterogeneity, while reducing sparsity; and (b) the z-scores for gene expression are less prone to batch effects, by utilizing the relative expression levels of the genes rather than the absolute expression values. The latter assumption is based on the idea that the relative expression levels of genes within samples are functionally important and can capture essential variation across samples. Moreover, we chose to use z-scoring as a linear transformation as opposed to nonlinear transformations, such as logarithmic scaling, as this enables the GMRF model to apply the linear cell fraction constraints easily to the expression values.

In the following sections, we provide further details about the processing steps for generating metacell-based expression z-scores to be used as inputs into the predictive models.

**Per-sample, per-cell type metacells**

1. Each dataset is filtered, with only those genes containing at least 50 UMIs in total being passed on to the cell-type-specific processing step. Additionally, only genes expressed in at least three cells are passed to the next step.
2. The cell types are treated sequentially. All cells of a particular cell type are separated out for Metacell-2 processing.
3. If, for a cell type in a dataset, there are <30 cells or the total sum of the UMI counts is <30,000, we calculate the simple median of the cell expression vector. This vector is appended as a “metacell” to an existing dataframe.
4. If the above conditions are not satisfied, the full Metacell pipeline is run.
    a. The dataset is converted to an *anndata* object.
    b. Metacell-2’s *divide_and_conquer_pipeline* algorithm is run.
    c. If the process fails, *None* is returned.
    d. The metacells are “collected” (using Metacell-2’s *collect_metacells*), and a dataframe is returned.

5. The returned dataframe, if not *None*, has the cell type appended to the names of the metacells, and is concatenated with the full dataframe containing all metacells from all cell types for that particular dataset.
6. If *None* is returned, the simple median is calculated as described above.

**Combining the metacells into z-score distributions**

Each metacell for each individual and study cohort are run through the following steps.

1. First, the gene expression in each metacell vector is converted from raw UMI counts to CPM.
2. The z-scoring step is run for each cell type separately:
   a. The mean of the gene expression across all metacells is found.
   b. Then, z-scoring is performed on the resulting average vector.
3. All z-scores calculated are then concatenated across the different studies.
4. The mean and SD of the z-scores across all cells of a given type in all studies are calculated.

The file containing metacell-normalized expression data is available on the brainSCOPE portal:

**File:** Metacells_Zscores_all.txt.gz: Normalized MetaCell gene expression values for seven cell types across individuals who pass quality control filters, used as input for GRNs and the LNCTP model.

#### 8.2 *LNCTP Framework*

**Main manuscript reference:** First supplementary reference in the first paragraph of "Imputing gene expression and prioritizing disease genes across cell types with an integrative model."

We define an integrated framework, which we refer to as a 'Linear Network of Cell-type Phenotypes (LNCTP), as an energy model representing the joint distribution of a collection of phenotypes of interest (including cell-type resolved phenotypes at multiple levels), especially brain disorders and traits, conditioned on (a representation of) the genotype. The network is linear in the sense that the expectation of any phenotype conditioned on any subset of other phenotypes is a linear function. As described below, this property ensures that the coheritability of phenotypes in the network can be readily estimated.

We partitioned the variables of the model into: genotypes $(z)$, intermediate phenotypes $(x)$, hidden (latent) factors $(h)$, and high-level/complex traits $(y)$. We further indexed the intermediate phenotypes into those associated with $C$ cell-types, denoted as $x_1$, $x_2$, ...$x_C$, and we used $x_0$ to denote those phenotypes associated with bulk measurements. Additionally, we used a set of intermediate phenotypes $x^{(c2c)}$ associated with cell-to-cell communication strengths, and a set of variables $f_{1...C}$ representing the estimated cell fractions in the bulk observations. For an individual $i$, $z_{in} \in \{0, 1, 2\}$ represents the alternative allele dosage for individual $i$ at a common SNP $n$, (where $n \in 1...N_{SNP}$), and $x_{icg} \in R$ represents a normalized summary of the expression of gene $g$ in cell type $c$ in individual $i$. Here, we used a z-score normalization of the meta-cell outputs as our summary variable. Further, $x^{(c2c)}_{i,L,c}$, $x^{(c2c)}_{i,R,c} \in R$ represent summary features associated with the ligands and receptors, respectively, of cell type $c$ in individual $i$, $f_{ic} \in [0,\ 1]$ represents the fraction of cell type c in the bulk data for individual $i$, $h_{iln} \in R$ represents the activation of hidden node $n \in 1...N_l$ at level $l \in 1...L$ in individual $i$, and $y_i$ represents a high-level phenotype of interest, for instance case/control (all examples we consider are binary).

The probabilistic model for the full LNCTP model is defined as follows:

$$p_{LNCTP}\left(y_i, h_i, x_i | z_i\right) = p_{GMRF}\left(x_i | z_i\right) \cdot p_{DNN}(y_i, h_i | x_i) \tag{1}$$

$$p_{GMRF}\left(x_i | z_i\right) \propto \exp\ (-E_{GMRF}\left(x_i | z_i\right))$$

$$E_{GMRF}\left(x_i | z_i\right) = x^T_{i0\cdot} J x_{i0\cdot} + \sum_g x^T_{i0g} b\left(z, \beta_g\right) + \sum_c \left(x^T_{ic\cdot} J_c x_{ic\cdot} + x^T_{ic\cdot} b_c + x^T_{ic\cdot} J^{(c2c)}_c x_{i,LR,c}\right) + \sum_{c_1, c_2} J^{(c2c)}_{c_1, c_2} x^{(c2c)}_{i,L,c_1} x^{(c2c)}_{i,R,c_2} + \lambda \sum_g \left(x_{i0g} - f(z)^T x_{i,1...C,g}\right)^2$$

$$p_{DNN}\left(y_i, h_i | x_i\right) = p_y\left(y_i | h_{iL\cdot}, W_{L+1}\right) \cdot \prod_{l=2...L} p_h\left(h_{iL\cdot} | h_{i,l-1,\cdot}, W_l\right) \cdot p_h\left(h_{i1\cdot} | x_i, W_1\right)$$

Here, the parameters of the model are $\theta = \{\beta_{1...G}, J_{0...C}, J^{(c2c)}, W_{1...L}\}$ and $\lambda$ acts as a hyperparameter. As suggested by the notation, $p_{GMRF}$ has the form of a Gaussian Markov Random Field (GMRF) conditioned on $z$, while $p_{DNN}$ is a stochastic deep neural network (DNN). Further, the parameters $\beta_{1...G}$ and $J_{0...C}$ reflect the sparsity structure arising from the eQTLs and GRN linkages, respectively (discussed below), where the non-zero elements of $J_c$ occur only between genes connected in the GRN of cell type $c$. To ensure that the model satisfies the linear conditional property mentioned above, we made the following choices for specific distributions: $p_h\left(v_1 | v_2, W\right) = \delta(v_1 | v_2^T W)$, $p_y\left(y | v_1, W\right) = Bernoulli(y | \sigma(v_1^T W))$, $b\left(z, \beta_g\right) = \beta_g^T z_g$, $f_c(z) = \hat{f}_c$. Here, $\delta(a|b)$ is a delta distribution, which is 1 if $a = b$ and 0 otherwise, and $\hat{f}_c$ is the expected value of $f_c$ at the population level. We note that more complex (non-linear) distributions can be modeled by varying these choices: in particular $p_h\left(v_1 | v_2, W\right) = \delta(v_1 | \sigma(v_2^T W))$ models a deterministic DNN with non-linear activation $\sigma$. Further, for a categorical variable $y$, the model forms a generalized linear model, and is hence linear in the logits of the response (or the liability in a probit model).

Finally, we note that we compared three specific architectures for $p_{DNN}(y_i, h_i | x_i)$. ***Dense***: Here, $W_{1...L}$ are a set of dense matrices; ***Sparse Embedding***: Here, L=1, and the nodes of $h_{i1\cdot}$ are partitioned into $(C + 2)$ sets of size $E$, where $E$ denotes the embedding dimensionality, and $W_1$ is sparsely structured so that non-zero connections appear such that nodes from the same cell-type/bulk/cell-to-cell components of $x_i$ are connected only to the same partition of $h_{i1\cdot}$; ***Sparse Embedding* + *Dense***: Finally, we allow L>1, but ensure $W_1$ is structured as above, while $W_{2...L}$ have dense connectivity. This last architecture is the version depicted in **Fig. 8A**. We note that in all architectures, we used batch normalization prior to each layer, $h_{il}$, which we folded into the matrix $W_l$.

#### 8.3 *LNCTP Motivation* - Motivation for linear network architecture

**Main manuscript reference:** First supplementary reference in the third paragraph of "Imputing gene expression and prioritizing disease genes across cell types with an integrative model."

The use of a hierarchical linear architecture for the LNCTP framework ensures that the model can be readily interpreted and related to population genetics quantities as described below. However, we also motivate our choice on both performance and theoretical grounds. Particularly, we observed that models with linear activations performed comparably or better than models with non-linear activations across architectures tested (**table S14**), in agreement with results in neuroscience suggesting that large-scale linear models are competitive in related datasets (*210*). Further, theoretical analyses of deep linear models show that they have specific benefits, such as implicit $l_1$ and $l_2$ regularization (along with structured weight sharing), which may explain their generalization properties (*210*, *211*).

#### 8.4 *LNCTP Training*

**Main manuscript reference:** First supplementary reference in the second paragraph of "Imputing gene expression and prioritizing disease genes across cell types with an integrative model."

We followed the method outlined in (*4*) to create datasets from the bulk expression data, ensuring balance for covariates including age, gender, ethnicity, and cohort. Hence, for each disorder, we generated ten data splits of the following sizes (training/testing): schizophrenia (640/70); bipolar disorder (170/18); ASD (50/12). Further details for training the AD model are described below. To these, we added the harmonized single-cell data, which were used to train the meta-cell priors. Importantly, we did not use the comprehensive set of cell types. Instead, we reduced the dimensionality of the overall network by using a coarse-grained set of cell types: all excitatory neurons were combined under the type “Excitatory,” inhibitory neurons under “Inhibitory”, and separate categories were created for the non-neuronal types “Astrocytes”, “Oligodendrocytes”, “Oligodendrocyte Precursor Cells (OPC)”, “Microglias”, and “Endothelial Cells”. The meta-cell construction followed this coarse-graining approach, with averages calculated across all finer-grained excitatory and inhibitory cell types within each broader category of “Excitatory” and “Inhibitory”, respectively. To create a tractable architecture for training, we further limited the genes in each model to include all TFs, along with high-confidence genes for schizophrenia, and the highest correlating genes with the case/control status on bipolar disorder and ASD. This resulted in a gene set of ~560 genes for each disorder, meaning that the dimensionality of x is ~5,000. We also considered the GRN set **GRN-B**, encompassing all the GRNs constructed for the different cell types, including pediatric samples (age < 13 years) and utilizing only the UCLA-ASD snATAC-seq peaks. The LNCTP was then trained piecewise as described below.

**Unary training**

Local predictors for each gene were trained using a Lasso loss function, to predict the z-score normalized expression from the eQTL SNPs associated with each gene. Hence, we optimized:

$$L_g^{Lasso}(\beta) = \left(x_{i0g} - \beta_{g0} - \sum_n \beta_{gn} z_{ign}\right)^2 + \lambda_{Lasso}\left|\beta_g\right|_1 \quad (2)$$

where $z_{i,g,n=1...N_g}$ are the $N_g$ bulk eQTL SNPs associated with gene $g$, and $\lambda_{Lasso}$ is the Lasso penalty, which is found through 10-fold cross-validation on the training partition.

**GMRF training**

We then trained the $p_{GMRF}(z)$ term in Eq. (1), while fixing the unary terms. The J matrices were initialized to the diagonal matrices: $J_c = diag([\left(\sigma_{c,g=1...N_g}^{meta}\right)^2])$ , where $\left(\sigma_{c,g}^{meta}\right)^2$ is the variance of gene $g$ in cell type $c$ from the z-scored meta-cell data. Further, the bulk unary terms

$$b_{0g}\left(z, \beta_g\right) = \frac{\beta_{g0} - \sum_n \beta_{gn} z_{ign}}{\left(\sigma_{0,g}^{metta}\right)^2} + b_{0g}^{off} \quad (3)$$

were set to:

where $\sigma_{0g}^2$ is the empirical estimate of the bulk variance of gene $g$, and $b_{0g}^{off}$ is a gene-specific offset, initialized to 0, while the cell-type unary terms were initialized to $b_{cg} = \mu_{cg}^{meta}/\left(\sigma_{c,g}^{meta}\right)^2$, where $\mu_{cg}^{meta}$ is the mean of the meta-cell data for gene $g$ in cell type $c$. These settings ensure that the GMRF is initialized to a Gaussian centered on the predictions from Eq. (2) in the bulk variables and at the meta-cell means for the cell-type-specific variables.

The GMRF training then proceeded by performing stochastic gradient descent on the loss for $p_{GMRF}(z)$, where the bulk variables were treated as observed and the cell-specific variables were treated as hidden. Hence, we have:

$$L^{GMRF}(x|z) = \log p_{GMRF}(x_0|z) = \log \int_{x_{1...C}} p_{GMRF}(x_0, x_{1...C}|z) \tag{4}$$

$$\nabla_{b_{cg}} L^{GMRF}(x|z) = E_{p_{GMRF}(x_{1...C}|z,x_0)\delta(z,x_0)}[b_{cg}] - E_{p_{GMRF}(x_0,x_{1...C}|z)\delta(z)}[b_{cg}]$$

$$\nabla_{J_{ij}} L^{GMRF}(x|z) = E_{p_{GMRF}(x_{1...C}|z,x_0)\delta(z,x_0)}[x_i x_j] - E_{p_{GMRF}(x_0,x_{1...C}|z)\delta(z)}[x_i x_j]$$

where the distributions $p_{GMRF}(x_{1...C}|z, x_0)\delta(z, x_0)$ and $p_{GMRF}(x_0, x_{1...C}|z)\delta(z)$ represent the clamped and unclamped GMRF distributions, respectively (see $(212)$ for a derivation of Eq. (4)).

We estimated the required expectations in Eq. (4) via Gibbs sampling; specifically, we used the updates:

$$x_i \sim N(.\,|\sum_j J_{ij} x_j, J_{ii}^{-1}) \tag{5}$$

where $x_i$, $x_j$ range across the nodes of the GMRF (note that, for clarity, we dispensed with the individual, cell-type, and gene indices here; further, we ensured that each GMRF variable received two Gibbs updates per expectation evaluation). We then made a step in the direction of the gradient from Eq. (4), while enforcing the GRN sparsity structure on the $J$ matrices. Since, in general, this may produce a non-positive-semi-definite $J$ matrix, we added multiples of a small value $\epsilon$ to the diagonal until the matrix became positive-semi-definite (hence, performing a projected Stochastic gradient descent update). We trained the GMRF by repeatedly taking gradient steps with a decreasing learning rate, while evaluating $L^{GMRF}$ on a subset of the training data, until this value no longer increased.

**DNN training**

To train the term $p_{DNN}(x)$, we fixed the $x$'s to the estimated bulk and cell-type expression from the GMRF training step, and used these to predict $y$. We optimized DNN modes using the dense, sparse-embedding, and sparse-embedding+dense architectures described above, while varying hyperparameters including the learning rate (0.01, 0.001, 0.0001), number of hidden layers (1, 3, 5, 7, 9), number of units per hidden layer (25, 50, 100, 200), and activation function (linear, RELU, sigmoid, tanh). We optimized all models on all ten data splits and monitored the loss on a subset of the training set (validation set) to determine an early stopping point. The optimal hyperparameter settings for each disorder were selected according to the best performing models on the validation partitions, and the reported performance was evaluated on a separate hold-out test set (per data split).

**LNCTP on AD Prediction**

**Dataset used**

We used the ROSMAP dataset as the main source to evaluate and train the LNCTP on AD prediction. Among the samples, 366 were cases (AD patients) and 179 were controls , composing a total of 545 samples. We split the data into a 3:1:1 ratio for train-valid-test sets, while maintaining the case-control ratio of approximately 67:33 before feeding them into the prediction models.

**AD model - *De novo* model and c2c version**

We evaluated LNCTP's performance in a *de novo* setting. First, since there are 878,363 SNP IDs in the ROSMAP genotypes, we padded it to a dimension of 1,097,784 with 0s (to match the SNP dimensions of the other disorder cohorts) and reordered it according to the mapping alignment. Using this reordered data, we identified and trained the best unary models for predictions. The unary results were then used by the GMRF to generate outputs for the AD samples. These predictions from GMRF include the gene expression predictions after the GMRF imputation, which have 545 rows, corresponding to the total 545 cases. We identified 578 significant genes (by their Ensembl ID; $p<0.05$, 2 tailed t-test for differential expression, cases vs control) based on the bulk imputed differential expression; this set includes TFs and AD high-confidence genes. Subsequently, we passed the GMRF imputed expression values through the WGCNA modules for preprocessing, including assigning modules for genes and skipping genes that do not exist in modules.

Once the data were processed, we ran the TensorFlow models with a selected range of hyperparameters and network architecture options for optimal performance (we did not perform additional replicate experiments for each of the options). In addition, we explored the c2c version of the TensorFlow models; we used GMRF outputs for the c2c version, and applied similar steps as before. The final results and qualitative comparisons with methods from other papers can be found in **table S15**.

#### 8.5 *LNCTP Interpretation*

**Main manuscript reference:** Second supplementary reference in the second paragraph, second supplementary reference in the third paragraph, and first and second supplementary references in the fourth paragraph of "Imputing gene expression and prioritizing disease genes across cell types with an integrative model."

By design, our model permits a variety of interpretation strategies:

##### Imputation of bulk and cell-type expression

For a given individual i, their bulk and cell-specific expression may be estimated directly from the genotype by evaluating $E[x_{i,0}, x_{i,1...C}|z_i]$. This can be evaluated efficiently via:

$$\hat{x}_{i,0} = b(z_i)\Sigma^{0,0} \quad (6)$$

$$\hat{x}_{i,c} = \hat{x}_{i,0}\Sigma^{0,c}$$

where $\Sigma = J^{-1}$. We compared the imputed gene expression from our model as above with imputed estimates formed by first applying Predixcan to impute bulk expression (*92*), and then applying CibersortX (*213*) in high-resolution mode using the mean expression profiles from the metacell analysis as the signature matrix, which we refer to as our 'Baseline model' in **Fig. 8C**.

##### Saliency-based cell-type and gene prioritization

For any individual node in our model, the saliency may be defined as the square of the gradient of the expected output of the network with respect to the node in question; hence:

$$sal(x_{icg}) = \left(\nabla_{x_{icg}} E\left[y'_i|z_i\right]\right)^2 \quad (7)$$

$$sal(h_{iln}) = \left(\nabla_{h_{iln}} E\left[y'_i|z_i\right]\right)^2$$

where we use $y'_i$ to denote the output of the final layer before passing through the sigmoid function (the log-odds ratio for $y$). We can thus use $sal(x_{icg})$ as a measure of the salience of gene $g$ in cell type $c$ in our model, and $\sum_{n\in E_c} sal(h_{i1n})$ as a measure of the salience of cell type $c$, where $E_c$ are the embedding nodes for cell type $c$ at hidden layer 1 of the DNN.

##### Heritability and coheritability estimates

All nodes in our network can be considered either endophenotypes (with either explicit or implicit semantics, corresponding to the $x$ and $h$ nodes, respectively) or high-level phenotypes (case/control status of a complex trait). The linear form of our network makes it straightforward to estimate the coheritability between pairs of (endo)phenotypes. Specifically, we evaluate:

$$h_{x,y} = \frac{C_{A_{x,y}}}{\sqrt{V_{P_x}V_{P_y}}} \quad (8)$$

where $h_{x,y}$ is the coheritability between phenotypes $x$ and $y$, which requires us to estimate $C_{A_{x,y}}$, the covariance between the additive genetic variance of $x$ and $y$, and the square root of the product of the variances of $x$ and $y$. We typically evaluate $h_{x,y}$, where $x$ is an endophenotype in the LNCTP, and $y'$ is the log-odds output. Hence, $C_{A_{x,y}}$ can be evaluated readily across the test set, and $V_{P_x}$, $V_{P_y}$' can be approximated using the variance of each node (alternatively, we may view this as an exact calculation of the coheritability of two additive traits). We used the direct Pearson correlation between $x$ and $y$ (on the test partition) to provide a summary p-value for each intermediate trait $x$ tested. Finally, we used the network outputs to estimate the heritability of each complex disorder on the liability scale, by scaling the outputs using Eq. (11) from *(4)*.

##### Prioritized subgraph analysis

To interpret and prioritize the hidden nodes of our model, we aimed to build a consensus, prioritized subgraph across the models learned for the same disorder across multiple data splits. To do so, for each model, we fixed parameters $A$ and $B$, representing the width and the branching factor of the reduced subnetwork, respectively. Then, for each data-split, at each level of its respective model, we chose the $A$ nodes with the highest absolute coheritability, and joined each of these to the $B$ nodes with the largest absolute connecting weights (in $W_l$) on the previous level. We then used these skeleton networks to produce a consensus graph by successively overlaying the subgraphs from each data split in a randomized order. Specifically, the nodes at the gene and cell-type embedding layers were overlaid deterministically, since these have explicit semantics. For each hidden layer, when a new graph was overlaid with the consensus graph, all $A!$ permutations were performed of the hidden nodes, and the one resulting in the greatest number of overlapping edges with the previous layer was selected (with ties broken arbitrarily). This process produced a weighted consensus subgraph (where the edges are weighted by the number of model reductions in which they appear), and a given internal node at a hidden layer in this consensus graph may be interpreted as a higher-order feature, grouping together the units at lower levels within the sub-tree below the selected node, allowing us to calculate salience and coheritability statistics for selected latent nodes of the network (see **fig. S80**). The subgraph prioritization process is summarized in Algorithm 1.

##### Polygenic risk score calculations

As a baseline for comparing the performance of the LNCTP model, we used a standardized pipeline (*214*) to calculate ASD, schizophrenia, bipolar disorder, and AD polygenic risk scores (PRS) from sets of SNPs overlapping the eQTLs used as LNCTP inputs. We used two sets of SNPs to calculate two AD PRS variants–one with AD-prioritized SNPs as inputs, and the other with schizophrenia-prioritized SNPs as input. We first accessed summary statistics for four recent large-scale GWAS studies for ASD, schizophrenia, bipolar disorder, and AD (*215–218*). We removed SNPs with INFO scores <0.8 as well as ambiguous and duplicate SNPs, and we further log-transformed the effect sizes of ASD SNPs. Next, we selected imputed genotypes from the population-scale samples assessed in the model and filtered for SNPs in eQTLs specific to ASD, bipolar disorder, and schizophrenia. After lifting the data over and fixing alleles to hg38 reference genome coordinates (for all disorders except AD), we performed strand-flipping using snpflip (https://github.com/biocore-ntnu/snpflip). SNPs with MAF<0.05, Hardy-Weinberg equilibrium p-values $<1\times10^{-6}$, or missing in >1% of samples were removed, and individuals missing >1% of genotype calls or with high or low rates of heterozygosity (>|3SD| from the cohort mean) were also removed. An additional strand-flipping step was performed to match alleles between the sample genotypes and summary statistic SNPs. Finally, we used LDPred2 (auto mode) to calculate disease-specific PRS for each sample (*219*), utilizing HapMap3-based haplotypes and 1,000 Genomes centimorgan map units to perform LD. Due to lower power, calculations for AD models did not use maximum likelihood estimates for alpha and variance components, did not allow for changing signs between iterations in the Gibbs sampler, and used a shrinkage multiplication coefficient of 0.8 for the LD matrix. To compare the PRS with the LNCTP models using the accuracy and liability metrics in **table S13**, we used the same ten data splits as above, and set a threshold for each data split by maximizing the accuracy on the training split and averaging over the test accuracies according to the selected thresholds. The heritability on the liability scale was then estimated as above for the LNCTP model.

**Algorithm 1 (Subgraph prioritization)**

**Inputs:** $W_{1...N,1...L}$ linear networks, $X_{1...N,1...S}$ data samples, $A$ output nodes per layer, $B$ output branching factor
**Outputs:** $W'_{1...N,1...L}$ sparsified linear networks, $\omega$ consensus sparsified network, $\mu$ map from individual to consensus networks

**for** $n = 1 \ldots N, s = 1 \ldots S$:
    calculate model activations $\alpha_{n,s}^{1...L}$ by passing $X_{n,s}$ through $W_{n,1...L}$
$W'_{1...N} \leftarrow \text{SparsifyNetworks}(W_{1...N,1...L}, \alpha_{1...N,1...S}^{1...L}, A, B)$
Initialize $\omega$ to empty network, with $L$ layers, $A$ nodes per layer, and no edges
**for** $n = 1 \ldots N$:
    Set $\mu_{n,l=1}$ to the identity map
    **for** $l = 2 \ldots L-1$:
        **if** $n == 1$:
            Set $\mu_{1,l}$ to the identity map
        **else:**
            **for,** $\pi \in \text{perm}(A)$:
                $o_\pi \leftarrow \text{Overlap}(\omega, \pi, W'_n, l)$
                $\mu_{1,l} = \text{argmax}_\pi(o_\pi)$
    **for** $i = 1 \ldots A, j = 1 \ldots A$:
        **if** $W'_{n,l-1,\mu_{1,l-1}(i),\mu_{1,l}(j)} \neq 0$:
            $\omega_{l-1,i,j} = \omega_{l-1,i,j} + 1$

**function** $W'_{1...N} \leftarrow \text{SparsifyNetworks}(W_{1...N,1...L}, \alpha_{1...N,1...S}^{1...L}, A, B)$
    Set $W'_{1...N} = W_{1...N}$
    **for** $n = 1 \ldots N, l = 1 \ldots L-1$:
        $W'_{1...N,a,:} = 0$ for all $a \notin \text{argmax}_C\{\text{sum}_{a' \in C}\left(\text{abs}\left(\text{corr}\left(\alpha_n^{l,a'}, \alpha_n^L\right)\right)\right); |C| = A\}$
    **for** $n = 1 \ldots N, l = L-1 \ldots 2$:
        $W'_{1...N,a,b} = 0$ for all $a \notin \text{argmax}_C\{\text{sum}_{a' \in C}\left(\text{abs}(W'_{n,l-1,a',b})\right); |C| = B\}$
        $W'_{1...N,:,b} = 0$ for all $b$ such that $\text{sum}(W'_{n,l+1,b,:}) = 0$

**function** $o \leftarrow \text{Overlap}(\omega, \pi, W'_n, l)$
    $o = 0$
    **for** $i = 1 \ldots A, j = 1 \ldots A, l = 2 \ldots L$:
        **if** $W'_{n,l-1,\mu_{1,l-1}(i),\mu_{1,l}(j)} \neq 0$ **and** $\omega_{l,i,j} > 0$:
            $o = o + 1$

The following file detailing the LNCTP inputs, as well as the Docker image containing the source code to run LNCTP, is available on the brainSCOPE portal:

**File:** LNCTP_prioritzed_genes_celltypes.xlsx (**data S30**): Excel file listing the genes, cell types, and network elements prioritized by the LNCTP model for schizophrenia, bipolar disorder, ASD, and AD, including salience, coheritability, and p-values for each gene-cell type combination.
**File:** lnctp.tar: TAR file containing Docker image of source code used to perform the LNCTP model analysis. This file is also available to download from the Docker repository at https://hub.docker.com/r/icefirecloud/lnctp-server.

#### 8.6 ***LNCTP Validation*** - In silico validation of LNCTP-prioritized genes and drug targets

We approached the validation of the LNCTP-prioritized gene sets in four parallel ways. First, we identified external support for the prioritized genes in the neuropsychiatric GWAS results and published gene sets associated with schizophrenia, ASD, and major depressive disorder (MDD). Second, we explored several of the visible (non-latent) components of the imputation model to evaluate how these components may have contributed to the prioritization of genes in the LNCTP. In doing so, we suggest some plausible sources of phenotypic association for particular genes. Third, we used the imputed gene expression vectors as the baseline and perturbed certain important categories of genes to quantify the downstream impacts on the case-control status. This served as both validation and a showcase of the utility of LNCTP for *in silico* experimentation. Finally, we performed an ablation analysis, where we compared the performance of the model with and without certain components to assess the importance of including them.

In most of these validation approaches, we primarily used a set of eight key genes prioritized by LNCTP (disorders in which they are prioritized are shown in parentheses; see **Fig. 8D** and **data S30**): *RORA* (schizophrenia, bipolar disorder); *TCF4* (schizophrenia, bipolar disorder); *MEF2A* (schizophrenia, bipolar disorder); *SF3B2* (schizophrenia); *ANKHD1-EIF4EBP3* (ASD); *LINGO2* (bipolar disorder); *ESRRG* (bipolar disorder, ASD); and *ID1* (bipolar disorder). These genes have high saliency in the trait-prediction models and are often found to be prioritized in multiple cell types. They cover all three neuropsychiatric disorders considered in this work. Importantly, these genes also cover some of the classes of genes described in the Discussion section of the main manuscript. Specifically:

**Class 1: LNCTP-prioritized genes that are not found to be significantly differentially expressed.** If we allow for inclusion in both the single-cell (this work) and bulk DE gene sets (*100*), *MEF2A* and *ID1* fall into this class of not being differentially expressed.

**Class 3: Genes prioritized in disorders by LNCTP with DE support but lacking extensive prior literature support.** While *ANKHD1-EIF4EBP3* and *RORA* were prioritized in the PEC 2018 DE gene analysis (*100*), there does not seem to be extensive additional literature support for them.

##### 8.6.1 Prior literature- and GWAS-based analysis of prioritized genes

We used recent GWAS for schizophrenia to identify genes containing fine-mapping SNPs (*217*). Through our investigation, we found that, among the eight key genes, TCF4 contained a fine-mapping SNP specifically associated with schizophrenia. Furthermore, TCF4 also exhibits associations with ASD (*220*) and developmental disorders (from the DDD study) (*221*) based on rare variant analysis.

The genes related to the respective diseases (ASD, MDD, and schizophrenia) that we compared were obtained from various sources. These sources were compiled in our previous study, ((*222*); **data S33**). Here, we classified these genes into 36 categories based on studies focusing on genetics, differential expression, and co-expression related to the aforementioned diseases (**fig. S84A**).

In addition to comparing the key-genes for GWAS fine-mapping and intersection with disorder related genes in previous sources, we compared all prioritized genes in SCZ for such enrichment. Hence, we ran hypergeometric tests for each prioritized gene in each cell-type (and all prioritized genes) to look for the enrichment of fine-mapped and literature associated genes within the prioritized set against the background of all genes in the SCZ model. The results are shown in (**fig. S84B**), showing particularly enrichment in excitatory and inhibitory neuronal gene sets.

##### 8.6.2 Network analysis of prioritized genes

To establish the relationship between LNCTP and some of its main inputs, the GRNs, we investigated the characteristics of the key highlighted TFs in **Fig. 8D**. We analyzed the patterns of the degree statistics of the TFs in the bulk GRN. **fig. S85N** shows the degree distribution (bottleneck connections and number of neighbors in the graph) of the TFs in the bulk GRN; all TFs highlighted in **Fig. 8D** have a higher betweenness degree (**fig. S85A**), but a more dispersed out-degree (**fig. S85B**).

##### 8.6.3 DE analysis of prioritized genes

We further investigated whether the prioritization of genes by LNCTP draws upon differential expression between disorder and control samples. Using the eight key genes as examples, we found that only *TCF4* is within the FDR-corrected significant DE gene set for any cell type. However, several of the key genes had $\log_2$ fold-changes that are almost completely positive or negative across all cell types in specific disorders, such as *TCF4* and *SF3B2* in bipolar disorder, indicating consistent up- or down-regulation.

Next, we examined the occurrence of the key genes in the set of DE genes between schizophrenia/bipolar disorder/ASD and controls in the PEC 2018 cohort (*100*). We identified *RORA* and *SF3B2* in the schizophrenia upregulated gene sets, LINGO2 in the schizophrenia and bipolar disorder upregulated gene sets, *ESRRG* in the schizophrenia and ASD downregulated gene sets, and *ANKHD1-EIF4EBP3* in the ASD upregulated gene set.

Overall, while differential gene expression may contribute to the prioritization of some of the genes, it is unlikely to be the sole characteristic of these genes.

##### 8.6.4 Perturbation analysis of prioritized genes and drug targets

The perturbation analysis consisted broadly of three steps (**fig. S86A**). First, we chose which genes to perturb. We identified three gene sets of interest: the eight key genes prioritized by LNCTP, a set of potential drug targets for the disorders at hand, and a set of background genes whose perturbations can serve as a comparison against the prior two sets. Our gene sets are summarized below:

1. Key genes: *RORA, TCF4, MEF2A, SF3B2, ANKHD1-EIF4EBP3, LINGO2, ESRRG, and ID1*. These are genes meeting a p<0.01 threshold on the coheritability analysis, and simultaneously occurring within a prioritized subgraph, as indicated in **Fig. 8D**.
2. Drug targets: *NFKB1, CACNA1D, FOS, ATF1, CHRNA2, ATF6, ESRRG, NR3C2, JUN,* and *TP53*. These are genes sampled from the 494 targets of neuropsychiatric drugs identified in DrugBank *(102)* that overlapped with the full gene sets for all three neuropsychiatric disorders.
3. Background: *NEU2, BUB1B.PAK6, CST6, F2, HARBI1, INO80E, ARHGAP1*, and *TARS2*. These are genes sampled from those having minimum rank across cell types of at least 2,000 in the schizophrenia model (using saliency to rank the genes; the maximum rank is 4,384, and 14 genes meet this threshold).

Second, we focused on one gene at a time in our chosen set, fixing it to either a high or low value (depending on that gene's difference of expression in cases as opposed to controls) in the bulk GRN segment of the LNCTP imputation model. Lastly, we re-imputed expression values for all other genes. To do this, we ran the imputation using a conditional form of the LNCTP energy model. This is summarized below:

$$p_{LNCTP-perturbed}\left(y_i, h_i, x_{i,\neg(c^*,g^*)} | z_i, x_{ic^*g^*} = \{1,-1\}\right) \propto p_{GMRF}\left(x_{i,\neg(c^*,g^*)} | z_i, x_{ic^*g^*} = \{1,-1\}\right) \cdot p_{DNN}(y_i, h_i | x_i)$$

$$p_{GMRF}\left(x_{i,\neg(c^*,g^*)} | z_i, x_{ic^*g^*} = \{1,-1\}\right) \propto \exp\left(-E_{GMRF}\left(x_{i,\neg(c^*,g^*)} | z_i, x_{ic^*g^*} = \{1,-1\}\right)\right)$$

$$E_{GMRF}\left(x_{i,\neg(c^*,g^*)} | z_i, x_{ic^*g^*} = \{1,-1\}\right) = x_{i0\cdot}^T J x_{i0\cdot} + \sum_g x_{i0g}^T b\left(z, \beta_g\right) + \sum_c \left(x_{ic\cdot}^T J_c x_{ic\cdot} + x_{ic\cdot}^T b_c + x_{ic\cdot}^T J_c^{(c2c)} x_{i,LR,c}\right) +$$
$$\sum_{c_1,c_2} J_{c_1,c_2}^{(c2c)} x_{i,L,c_1}^{(c2c)} x_{i,R,c_2}^{(c2c)} + \lambda \sum_g \left(x_{i0g} - f(z)^T x_{i,1\ldots C,g}\right)^2 + \infty \cdot \delta(x_{ic^*g^*} = \{1,-1\})$$

where the notation is as in Eq. 1, section 8.2, $(c\ *,\ g\ *)$ denotes the perturbed gene and cell type, whose expression is set to 1 or -1, and $\delta(a)$ is a delta function whose value is 0 if expression $a$ is true, and 1 otherwise.

Third, as a further validation step, we investigated whether the LNCTP-prioritized gene perturbations drive the overall gene expression patterns of control samples towards more "case-like" behavior. The underlying idea is that if we perturb individual key genes and observe that the pattern of gene expression for control samples deviates in a manner that begins to resemble the expression patterns for a particular phenotype, then it adds to the evidence that those genes are relevant for that particular phenotype. The saliency calculation is itself such a quantification of how specific components of LNCTP contribute to the phenotype, but we want an independent measure for this validation step. Accordingly, we chose a simpler approach focused on imputed gene expression alone, which ignores the contributions of the latent layers of the trait-prediction component within LNCTP and focuses exclusively on the outputs of the imputation component.

The unperturbed z-scored gene expression vectors are fed into a support-vector classifier (SVC, using the Python *scikit-learn* package *(223)*) that is trained to classify samples into either phenotype cases or controls (**fig. S86A**). We use a linear kernel for the SVC, resulting in a hyperplane separating the cases and controls. The unit normal to this hyperplane broadly defines a direction of separation between controls and phenotypic cases. When the perturbations shift the positions of expression vectors for all samples, we quantify the degree to which the balance between the numbers of cases and controls is modified. The hypothesis is that a perturbation of a key gene in the direction of case-like behavior should drive the entire expression vector for the sample in the direction of case-like behavior, if the perturbed gene is truly relevant to the disorder (**fig. S86B**).

The SVC training accuracies were 0.997 for schizophrenia, 0.857 for bipolar disorder, and 1.0 for ASD (likely due to the smaller number of ASD samples). Note that the case and control numbers are nearly balanced in the unperturbed cohort. Given the trained SVC, we classified the perturbed samples as "cases" or "controls" for the forward and reverse perturbations of the key genes, drug target genes, and background genes. When we measured the net increase in the number of predicted schizophrenia cases upon perturbing the gene, we found an  increase in the numbers of predicted cases for the drug targets and key genes relative to the increase in cases for the background genes, which are more evenly distributed around 0 (**fig. S86C**). We focus here on schizophrenia cases as they form the largest phenotypic group in our cohort. Naturally, there are some caveats in this simple analysis. These results are to be understood as limited by the relatively small numbers of genes considered. We only explored the bulk gene expression in the perturbation process, rather than the cell-type-specific expression. Furthermore, given the fact that the schizophrenia gene sets we start with are those previously identified as high-confidence schizophrenia genes *(4)*, it is difficult to identify a true "background" set of genes. Nonetheless, the results are somewhat indicative of the disease relevance of the key genes.

Files describing the results of the perturbation analysis are available on the brainSCOPE portal. Each .zip file contains individual .csv files for results from forward and reverse gene perturbations for background and drug target genes, with disease-specific reference files included for comparison. The suffix of '_1' or '_-1' at the end of the .csv file names refers to whether the given gene is perturbed up (_1) or down (_-1); it reflects whether that gene's expression value was set to 1 or -1 in the model.

**File: perturbed_expression.zip:** ZIP file containing forward perturbation results of key genes identified by the LNCTP model.
**File: perturbed_expression_background.zip:** ZIP file containing reverse perturbation results of key genes identified by the LNCTP model.
**File: perturbed_expression_background.zip:** ZIP file containing forward perturbation results of background genes.
**File: perturbed_expression_background_reverse.zip:** ZIP file containing reverse perturbation results of background genes.

**File: perturbed_expression_drugs.zip:** ZIP file containing forward perturbation results of drug target genes.
**File: perturbed_expression_drugs_reversed.zip:** ZIP file containing reverse perturbation results of drug target genes.

##### 8.6.5 CLUE analysis

**Main manuscript reference:** First supplementary reference in the seventh paragraph of "Imputing gene expression and prioritizing disease genes across cell types with an integrative model."

We used the query tools available on Clue.io (*42*) to probe the *Connectivity Map* (CMAP) for reference perturbagen signatures that counteract the effects of the perturbation. The *Gene Expression L1000* data, drawn from the platform's latest version, formed the basis of our query parameters. Furthermore, we incorporated the top 150 upregulated genes as well as the top 150 downregulated genes (by log2-fold-change) from each cell type into our search. We established a stringent statistical threshold by considering only those results as significant that had a q-value less than 0.01 (q-values are provided by the database using methods outlined in (*42*)). The significant findings are compiled and presented in a supplemental file (see below for description), provided on the brainSCOPE portal.

For the eight genes in the perturbation analysis in the eight cell types, we found 17,725 compounds and 566 peptides or other biological agents (such as cytokines) (**table S17**). Among these, several well-known drugs used for neuropsychiatric disorders emerged, including dopamine receptor antagonists, dopamine receptor agonists, glutamate receptor antagonists, calcium channel blockers, GABA receptor agonists, and MAP kinase inhibitors.

In addition to these established drugs, our analysis revealed insights related to molecular information and compounds with unknown effects. For instance, we observed that the cytokine IL-1a shows potential in reversing the expression changes of the ID1 gene in microglia. This finding supports previous studies (*224*) highlighting cytokine imbalances in schizophrenia, thus strengthening these hypotheses.

Furthermore, we identified a compound called 10-DEBC, an AKT inhibitor, which exhibits significant effects on reversing the forward perturbations of *TCF4*, *ID1*, *RORA*, *SF3B2*, and *ANKHD1-EIF4EBP3*. A recent study (*225*) revealed that AKT inhibition in the central nervous system can lead to signaling deficits, thereby triggering psychiatric symptoms. While our *in silico* analysis suggests that 10-DEBC could potentially counteract the gene expression effects of several key genes across multiple cell types, the aforementioned study suggests that AKT inhibitors might have adverse psychiatric effects. Understanding and reconciling these seemingly contradictory results in terms of the specific network effects induced by the drug could lead to valuable insights.

Another intriguing discovery was the consistent occurrence of the bromodomain inhibitor in the perturbation reversal results of all eight genes studied. Published research (*226*) indicates that treatment with the bromodomain inhibitor JQ1 significantly rectifies abnormal gene expression in schizophrenia patient-derived neurons.

A file summarizing our CLUE analysis results is provided on the brainSCOPE portal:

**File**: clue_significant.xls: This file contains a list of perturbagens identified through CLUE that elicit expression profiles opposing perturbations induced by key genes prioritized in LNCTP. A summary table listing the number of perturbagens identified per perturbed gene and cell type is also provided.

##### 8.6.6 Network analysis of LNCTP perturbations

To understand how the LNCTP perturbation results are affected by GRNs, we calculated the correlation of the genes' proximities to the perturbed gene in the GRN with the magnitude of the imputed expression results after perturbation. We hypothesized that the magnitude of the imputed gene expression should be inversely correlated with the proximity of the gene to the perturbed gene in the GRNs. The proximity of the genes in the GRNs could be measured by different metrics: in-degree, out-degree, and PageRank scores *(227)*. These metrics are not available for all genes in the GRNs because the GRNs are directed; target genes in the GRNs cannot have a non-zero out-degree or PageRank score when they themselves are the reference nodes, as they are the dead-ends in the graph. **Table S16** shows that all these metrics are inversely correlated with the magnitude of the imputed gene expression after perturbation, although some of them are not statistically significant (two-sided t-test, $p \leq 0.05$).

We further compared PageRank scores of top- and bottom-ranked genes based on their imputed gene expression. We ranked the genes by their absolute imputed expression values and labeled the top and bottom 10% genes. We then compared PageRank scores of the bottom and top affected genes using the Wilcoxon rank sum test. **fig. S87** shows that the overall magnitudes of the LNCTP imputations are inversely correlated with the PageRank scores; hence LNCTP predictions are in line with our initial hypothesis that proximal genes to the perturbed genes in the GRNs are affected to a greater extent by the perturbation.

##### 8.6.7 Ablation analysis

In order to assess the contribution of each component to the LNCTP performance, we compared the model with the following energy forms:

$$E_{GMRF-unrary}\left(x_i|z_i\right) = \sum_g x_{i0g}^T b\left(z, \beta_g\right)$$

$$E_{GMRF-unary+GRNs}\left(x_i|z_i\right) = x_{i0}^T J x_{i0\cdot} + \sum_g x_{i0g}^T b\left(z, \beta_g\right) + \sum_c \left(x_{ic}^T J_c x_{ic\cdot} + x_{ic\cdot}^T b_c + x_{ic}^T J_c^{(c2c)} x_{i,LR,c}\right) + \lambda \sum_g \left(x_{i0g} - f(z)^T x_{i,1\ldots C,g}\right)^2$$

The first includes only the unary terms for the bulk expression imputation (this is equivalent to a TWAS/Predixcan model with a LASSO penalty on the SNP-gene links), while the second also includes the GRN connections, which allow us to impute the cell-type-specific expression without the cell-to-cell network connections. The results are presented in **table S13**.

#### 8.7 *Independent CRISPR validation of LNCTP*

**Main manuscript reference:** First supplementary reference in the last paragraph of "Imputing gene expression and prioritizing disease genes across cell types with an integrative model."

##### 8.7.1 Gene expression comparisons to CRISPR experiments

For an independent validation of the prioritized gene sets from our analyses, we identified published CRISPR experimental data in human brain cell types. One such resource, CRISPRbrain.org (*103*), consolidates data from multiple publications reporting on assays targeting human brain cell types such as glutamatergic neurons, microglia, and astrocytes. We focused on the CRISPRi (CRISPR interference) and CRISPRa (CRISPR activation) results from one paper in this resource, Tian et al., 2020 (*103*). In this study, the authors ran both sets of CRISPR assays in glutamatergic neurons differentiated from human induced pluripotent stem cells (iPSCs) on specific target genes. Subsequently, they performed a transcriptome readout using a method called CROP-seq. This involved obtaining transcriptome data from groups of single cells containing guide RNAs for the target genes, and then determining differential expression relative to cells with control guide RNAs. The experimental setup in this study is analogous to perturbing genes in the LNCTP GRNs and observing the resulting gene expression patterns. Thus, we compared the experimental differential gene expression patterns to those in LNCTP for overlapping sets of genes. Additionally, we examined the agreement of the CRISPR perturbations with the expected differences in expression of some target genes between schizophrenia cases and controls, or whether the CRISPR perturbations on select genes push gene expression patterns towards a more "case-like" or "control-like" behavior in agreement with the fold-change differences observed for those select genes.

We selected 10 gene perturbations from the external CRISPR dataset, which intersected with the genes included in our LNCTP network, representing both prioritized and non-prioritized classes of genes in our analysis. The chosen genes/perturbations were *MEF2C-i* (related to *MEF2A* which is in class 1: LNCTP prioritized genes that are not DE genes); *GLIS3-a* and *HSPA9-i* (class 1: LNCTP prioritized genes that are not DE genes); *PSAP-i* and *WNT3-a* (class 2: genes prioritized by cell-to-cell network analysis); *ARHGAP20-a* (class 3: LNCTP and DE-prioritized gene without extensive prior support); *ATXN7-a*, *ATXN-i*, *FOXC1-i,* and *SOX5-a* (non-prioritized genes). Here, we use -i and -a to denote interference and activation CRISPR perturbations; see also section 8.6 and the Discussion for an in-depth discussion of these classes of prioritized genes. We note that for certain target genes, the induced differential expression in the CRISPR experiments might not align with the expected direction. For example, in the CRISPRa assay targeting MEF2C, we observed that the expression of MEF2C is actually reduced relative to the controls; the log2 fold-change = -0.25. This is counter to the expectation that the activation assay would increase the gene expression of the target gene. Nevertheless, we have included these genes under the assumption that there might be biological and/or technical reasons (such as experimental noise) for this counterintuitive behavior but the overall modification of cellular expression patterns would still be informative.

For each gene, we applied LNCTP perturbations in both positive (activation) and negative (interference) directions, using the method described in section 8.6.4. Note that the

GRN expression values in LNCTP were quantified as z-scores. We applied these perturbations separately to both the gene nodes in the bulk GRN network and the excitatory neuron GRN network, reflecting the application of CRISPR perturbations in glutamatergic neurons. We applied these perturbations to the individuals from each of the data splits for the LNCTP schizophrenia model (combining training/validation/test partitions) and calculated the mean (signed) change in z-score across all other genes in the respective bulk/excitatory GRN. We then calculated the correlation of these z-score changes with the external CRISPR observed log2 fold-change vectors for each data fold. Subsequently, we compared the distributions of the Pearson's correlations for matched perturbations vs. unmatched perturbations. **fig. S88** shows the results of this analysis, presenting both the case where the correlation is taken across all other genes in the GRN, or only across the upper decile of genes, according to their absolute z-score change in the LNCTP perturbed network. In addition, we consider the distribution across all perturbations, and the restriction to perturbations involving genes with at least 10 neighbors in the LNCTP network. As shown in panel A, the correlations of the LNCTP excitatory network and CRISPR perturbations are higher when comparing matched perturbations, and the difference in correlation values is enhanced when the upper decile is chosen (reaching $r$=0.81 correlation for *GLIS3-a,* as shown in **Fig. 8F**) and when distributions are compared for perturbations involving highly connected genes. Panel B shows that a similar separation of correlations in matched versus unmatched directions when considering perturbations in the LNCTP bulk network, although these are slightly attenuated with respect to the excitatory network perturbations (as expected, given that the CRISPR perturbations are in glutamatergic neurons). Panel C then shows that, for the same perturbation and gene sets, by combining the matched with the absolute values of the unmatched Z-score changes for the LNCTP perturbations, we can calculate a 'joint perturbation score' (defined in the caption, and illustrated in panel D), which achieves still higher correlation values with the CRISPR data (median r=0.56 and a maximum r=0.91, for the same perturbations and genes as in panel A, lower right).

For a closer one-to-one correspondence, we also converted the CRISPR expression vectors into z-scores (using the log2 CPM counts and log2 fold-change vectors) and calculated the Pearson's correlations between the signed z-score differences for the CRISPR perturbations and the LNCTP perturbations. In this case, we calculated the correlations for each individual in each data split, resulting in (10*Number of individuals) data points, restricting the perturbations to genes with at least 10 neigbors in the LNCTP network. The results, shown in **fig. S89A,** affirm that the correlations are higher when the CRISPR and LNCTP perturbation directions are matched relative to when they are not. However, we note that the absolute values of the correlations, as well as their differences, are reduced with respect to **fig. S88**; plausibly, this reflects the variation introduced by individual genetic backgrounds (including multiple disorder types) in the LNCTP perturbations, while the population mean is better matched to the observed perturbations in the *in vitro* CRISPR system, with a uniform genetic background. Further, we compared the effect of network distance with respect to the perturbed gene on the accuracy of the LNCTP predictions. Panel B shows how the mean neighborhood size varies with network distance in the LNCTP networks. Panel C shows that we observe an increase in the matched correlation values and the matched versus unmatched difference when looking at the 1-hop and 2-hop neighbors to a given perturbed gene in the excitatory network; as expected, the correlations are strongest for the immediate 1-hop neighbors, and gradually decrease with the

2-hop and larger neighborhoods, while a separation between matched and unmatched conditions remains for all neighborhood sizes. Panel D replicates the analysis of panel C, but uses the 'joint perturbation score' defined above, showing similar behavior.

We further investigated whether the CRISPR perturbation vectors for prioritized genes preferentially align with the unit normal to a Support Vector Classifier (SVC) hyperplane, representing a direction of separation between controls and phenotypic cases, as described in section 8.6.4 above. We trained 100 such SVC models by bootstrapping the samples across each of our 10 data folds (combined). For each CRISPR perturbation, we determined the expected sign of its dot-product with the SVC unit normal by determining whether the perturbed gene was empirically up or downregulated in our bulk RNA-seq data for schizophrenia cases. For instance, since *GLIS3* is upregulated in schizophrenia, a CRISPR-activation perturbation of this gene is expected to produce a positive dot-product with the SVC normal vector. **fig. S90** shows the results of this analysis across all genes, where (A) shows the distribution of dot products for each gene along with the expected signs, and (B) shows that perturbations expected to increase case-status exhibit higher dot-products than those expected to increase control-status. Finally, (C) shows that the mean difference in dot-products for expected case-enhancing vs. expected control-enhancing perturbations is greater for genes in the prioritized classes noted above than for non-prioritized genes.

These results indicate a positive agreement between CRISPR experiments and the perturbations induced *in silico* using the trained LNCTP network. The degree of agreement is impacted by the fact that the CRISPR experiments were performed on a homogenous population of glutamatergic neurons, subject to technical noise, while the perturbations on the trained LNCTP were carried out for many individuals with unique genetic backgrounds. Despite this disparity, the promising correspondence between the two gives us greater confidence in the outputs of our computational approach.

Files describing the results of the perturbation analysis for CRISPR targets are available on the brainSCOPE portal. The .zip files contain individual .csv files for results from gene perturbations for CRISPR target genes, with disease-specific reference files included for comparison. The suffix of '_1' or '_-1' at the end of the .csv file names refers to whether the given gene is perturbed up (_1) or down (_-1); it reflects whether that gene's expression value was set to 1 or -1 in the model. _exc suffixes in the .csv file names indicate that a gene was perturbed in the excitatory neuron network as opposed to the bulk networks.

**File: perturbation_crispr.zip:** ZIP file containing results for perturbation of bulk and excitatory networks with CRISPR target genes.
**File: perturbation_crispr_ref.zip:** ZIP file containing disease-specific reference expression datasets used in perturbation of CRISPR targets.

##### 8.7.2 Gene regulatory network (GRN) overlaps with CRISPR differentially expressed genes (DEGs)

We further evaluated the correspondence between our results and those of the CRISPR experiments by examining the overlaps between target genes of TFs in our GRNs and the DEGs in the CRISPR experiments (FDR < 0.05 from the authors' original analysis (*103*)) where those same TFs are perturbed. The basic hypothesis is that, if there is a strong correspondence between our inferred networks and the CRISPR experiments, there should be an enrichment of the DEGs among the downstream targets of TFs in our GRNs (relative to the upstream genes). We followed two separate approaches to quantify the overlaps, with the results shown in **table S18**.

###### 8.7.2.1 Diffusion-network-based approach

We used the network diffusion approach (from Section 5.5 *Unifying TF-target Regulons*) to create an excitatory-neuron-specific diffusion graph that combines all the weighted GRNs from the excitatory subtypes (L2/3 IT, L4 IT, L5 IT, L6 IT, L6 IT Car3, L5 ET, L6 CT, L5/6 NP, L6b). This is done to match the target glutamatergic cell types in the CRISPR experiments. Next, for each of the 9 TFs in our GRNs that overlap with the CRISPR target gene set – *GLIS3, NR2F2, SOX5, PPARGC1A, TAF1, MEF2C, THAP1, EGR2, SPI1* – we found the strength of the diffusion-based connections of the TF to all genes in our excitatory GRN. We then used several quantile thresholds to determine which genes are 'downstream' of the TF (higher diffusion scores) or 'upstream' of the TF (lower diffusion scores). We compared the overlaps of CRISPR DEGs (FDR<0.05, determined in (*103*)) with the downstream genes versus those with the upstream genes using a Fisher's exact test to determine if there is a significant (nominal p<0.05) enrichment of CRISPR DEGs in the downstream gene set from our diffusion network. The approach used the upstream genes as a background set for the estimation of the statistical significance of the enrichment in the downstream gene set.

The results are provided in **table S18A**. The CRISPR results themselves show considerable variation between the 9 TFs studied: the number of DEGs (FDR<0.05) varies, from 1,997 for SOX5 to 3 for MEF2C (row 1 of **table S18A**); the effect size on the TF itself is not uniformly significant in terms of FDR value (Row 3 of **table S18A**). In fact, some of the target TFs are not even observed in the experiments where their expression is modified (EGR2, SPI1; both are CRISPRa experiments). With regard to the latter point, we have noted above that in some cases (such as *MEF2C*), the gene expression of the intended target of the CRISPR experiment does not change significantly. This could be because of dataset noise or other regulatory feedback processes that compensate for the induced changes in expression. However, both these sources of variability play a role in explaining the correspondence between our GRN downstream genes and CRISPR DEGs.

Overall, we find, for diffusion score quantile thresholds of 0.5-0.8, that the downstream-DEG overlap is significant at the nominal p<0.05 (one-sided Fisher's exact test for greater enrichment in downstream genes) level for *SOX5*, *NR2F2*, *TAF1*, and *GLIS3*. *PPARGC1A* is marginally significant (nominal p<0.1 across all thresholds). For the other TFs, the overlap is not significant. The significant TFs are those for which the number of DEGs is higher and for which the effect sizes in the CRISPR experiment are very significant or near-significant (the table is ranked from right to left in terms of the effect sizes). Thus, it seems to be the case that we get high correspondence with the CRISPR results when a combination of the numbers of DEGs and the effect sizes are high. We believe that these results provide strong independent corroboration of our networks, especially given that we are comparing our results to single-cell CRISPR experiments on induced glutamatergic neurons derived from stem cells (and not tissue-derived neurons from a diversity of individuals).

###### 8.7.2.2 Hop-distance-based approach

We applied a similar overlap analysis to a discrete version of the network, which we term the 'hop-distance-based approach'. The network in this case was generated by calculating the distances between TFs and target gene with the *distances* function in the *R igraph* package (*228*, *229*) applied to a combined excitatory neuron GRN, which in turn was created as the union of all the GRN connections in the excitatory subtypes. In the network, "downstream" and "upstream" directions are determined by whether a target gene is reachable from a TF by following the directed connections in the GRN (downstream) or if the TF is reachable from the gene (upstream); the distance between a TF and up-/down-stream genes are found as the total number of steps needed within the GRN to reach the target (a gene in the downstream case, the TF in the upstream case). We chose to consider all genes within a hop-distance of $\leq 2$ as downstream, and pooled all the downstream genes with hop-distance > 2 and all upstream genes together as "upstream". This is because the high interconnectedness of the network meant that most genes were labeled as downstream in *igraph*. Since the goal is to observe whether more proximal downstream genes show an enrichment in CRISPR DEGs, we chose a hop-distance cutoff of 2 as reasonable.

We note that although the concept of the 'hop' distance does implicitly play a role in the network diffusion approach, a multi-hop path between a TF and a target gene may also produce a strong diffusion score as long as there is a high probability of observing those connections in the GRN. That is why we consider both approaches here, even though the correlation between the two is expected to be high.

The results are shown in **table S18B**. The TFs are ordered as in **table S18A** and the patterns of significance (at the $p<0.05$ and $p<0.01$, one-sided Fisher's exact test for greater enrichment in downstream genes) are essentially the same as in the diffusion-network-based approach: *SOX5*, *NR2F2*, *TAF1*, *GLIS3*, and *PPARGC1A* are significant at the $p<0.05$ level, while *THAP1* overlaps demonstrate a p-value of 0.058. The same conclusions as above hold, where CRISPR experiments with greater impacts on the target TFs and with more DEGs tend to show significant enrichment in the downstream gene set of our networks.

### Supplementary Figures

A

| DLPFC samples by disorder | snRNA-Seq | snATAC-Seq | Genotypes | | | Biol. sex | | Ancestry | | | | |
|---|---|---|---|---|---|---|---|---|---|---|---|---|
| | | | WGS/WES | Array | RNA-based | Female | Male | EUR | AFR | AMR | EAS/SAS | Unknown |
| Control | 182 | 30 | 23 | 137 | 22 | 61 | 121 | 140 | 29 | 8 | 4 | 1 |
| Schizophrenia | 77 | 6 | 6 | 70 | 1 | 29 | 48 | 60 | 11 | 3 | 3 | 0 |
| ASD | 52 | 9 | 23 | 24 | 5 | 10 | 42 | 43 | 4 | 4 | 1 | 0 |
| AD/Cognitive disorder | 33 | 0 | 29 | 3 | 1 | 18 | 15 | 31 | 1 | 0 | 1 | 0 |
| Bipolar disorder | 34 | 10 | 10 | 24 | 0 | 16 | 18 | 34 | 0 | 0 | 0 | 0 |
| MDD/PTSD | 10 | 0 | 0 | 10 | 0 | 2 | 8 | 8 | 2 | 0 | 0 | 0 |
| **brainSCOPE total** | 388 | 59 | 91 | 268 | 29 | 136 | 252 | 316 | 47 | 15 | 9 | 1 |

B

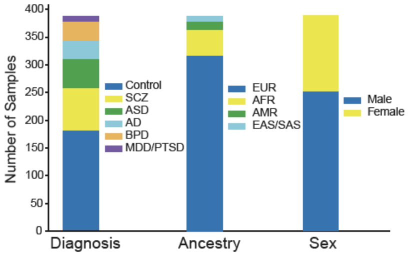


C

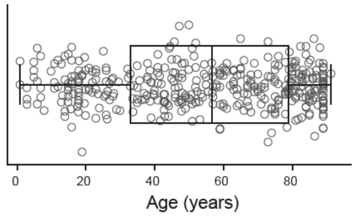


D

| Cohort | Sample size |
|---|---|
| DevBrain | 9 (full WGS) |
| Girgenti-snMultiome | 9 (full WGS) |
| ROSMAP | 37 (full WGS) |
| UCLA-ASD | 2 (full WGS) |
| Velmeshev | 13 (exome only, no SVs) |
| MultiomeBrain | 21 (VCF, no SVs) |
| **Total** | 88 rare SNVs; 54 with SVs |

E

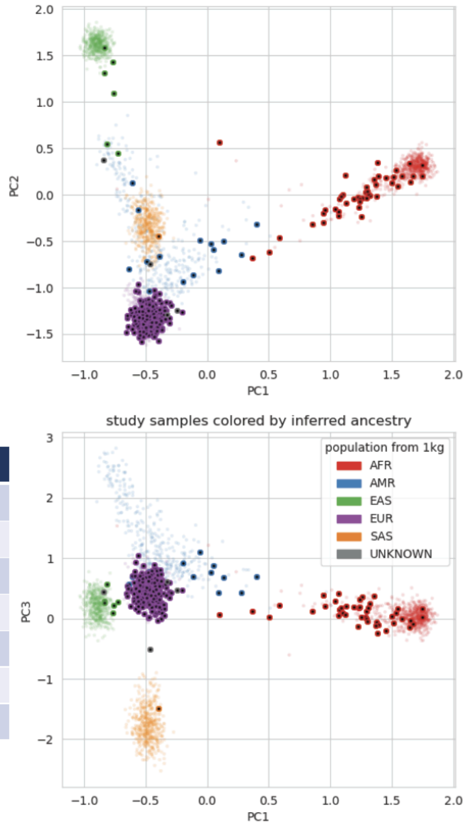


**Fig. S1. Demographic metadata for samples in the brainSCOPE cohort.**

**(A)** Table shows sample counts in the brainSCOPE Resource cohort by data modality, biological sex, ancestry, and disease. Sample ancestries were predicted using Peddy software based on 1,000 Genomes datasets (*113*). **(B)** Bar plot shows the distribution of samples by demographic metadata. **(C)** Box plot shows the distribution of sample ages. **(D)** Table shows sample sizes per cohort with available next-generation sequencing data (WGS and exome) for rare variant and SV calling. **(E)** Scatter plots show sample PCs denoted by ancestry. Translucent points in the background are genotype PCs from 1,000 Genomes samples.

More detail in the supplementary section "***Dataset Overview.***" This supplementary figure relates to **Fig. 1** and main text section "Constructing a single-cell genomic resource for 388 individuals."

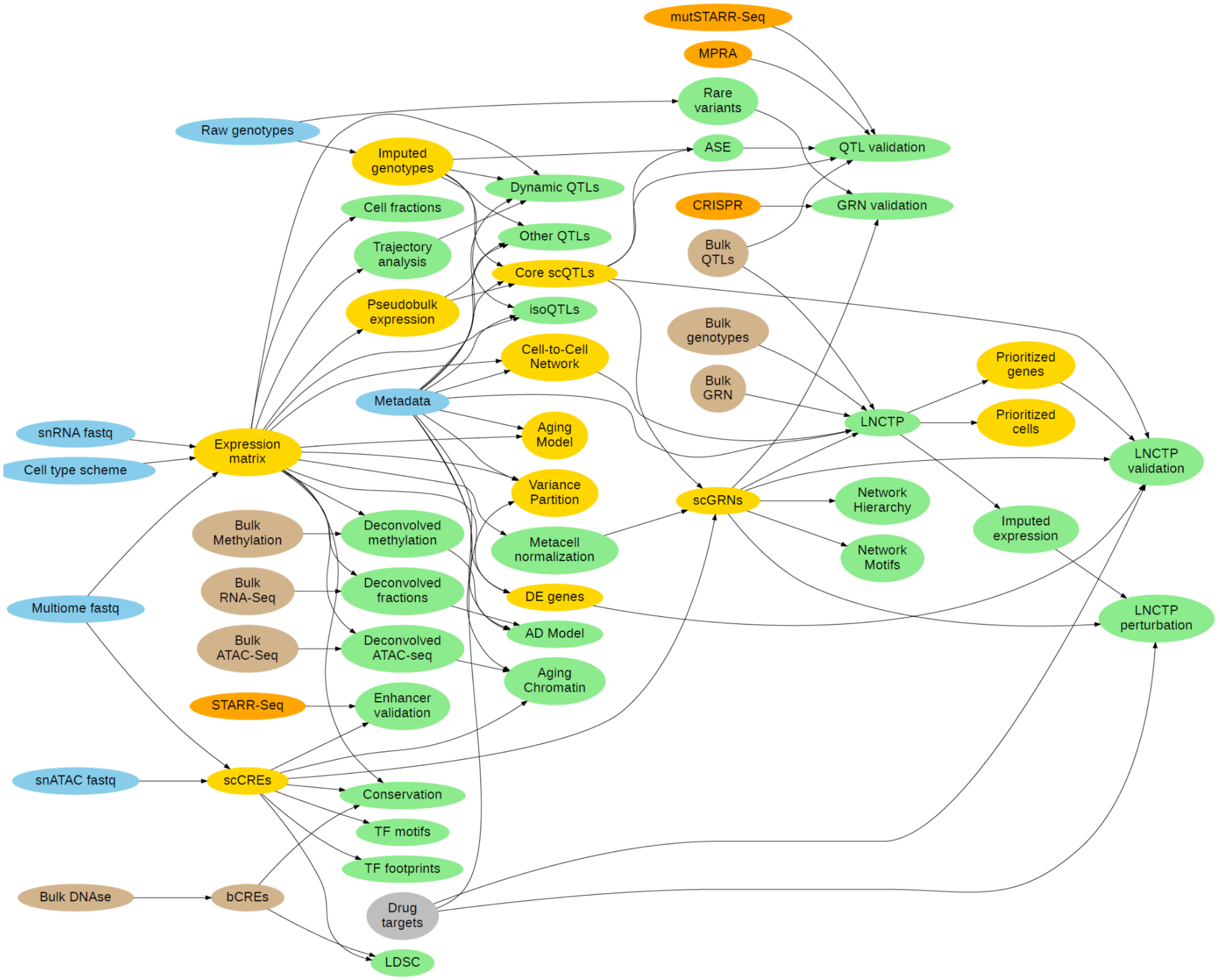


**Fig. S2. Dependency graph for all datasets in the manuscript.**

Blue nodes indicate new single-cell sequencing data and metadata generated for this manuscript, and brown nodes indicate bulk sequencing data used for imputing our single-cell data in a population context. Yellow nodes represent key resources generated from our analyses, while green nodes represent additional datasets available on the brainSCOPE portal. Orange nodes represent functional experiments from the PEC validation group used to validate our results; gray nodes represent external databases used for *in silico* validation.

More details on the input and output files can be found in **data S3**.

Further details are described in the supplementary section "***Dataset Overview***." This supplementary figure relates to **Fig. 1** and main text section "Constructing a single-cell genomic resource for 388 individuals."

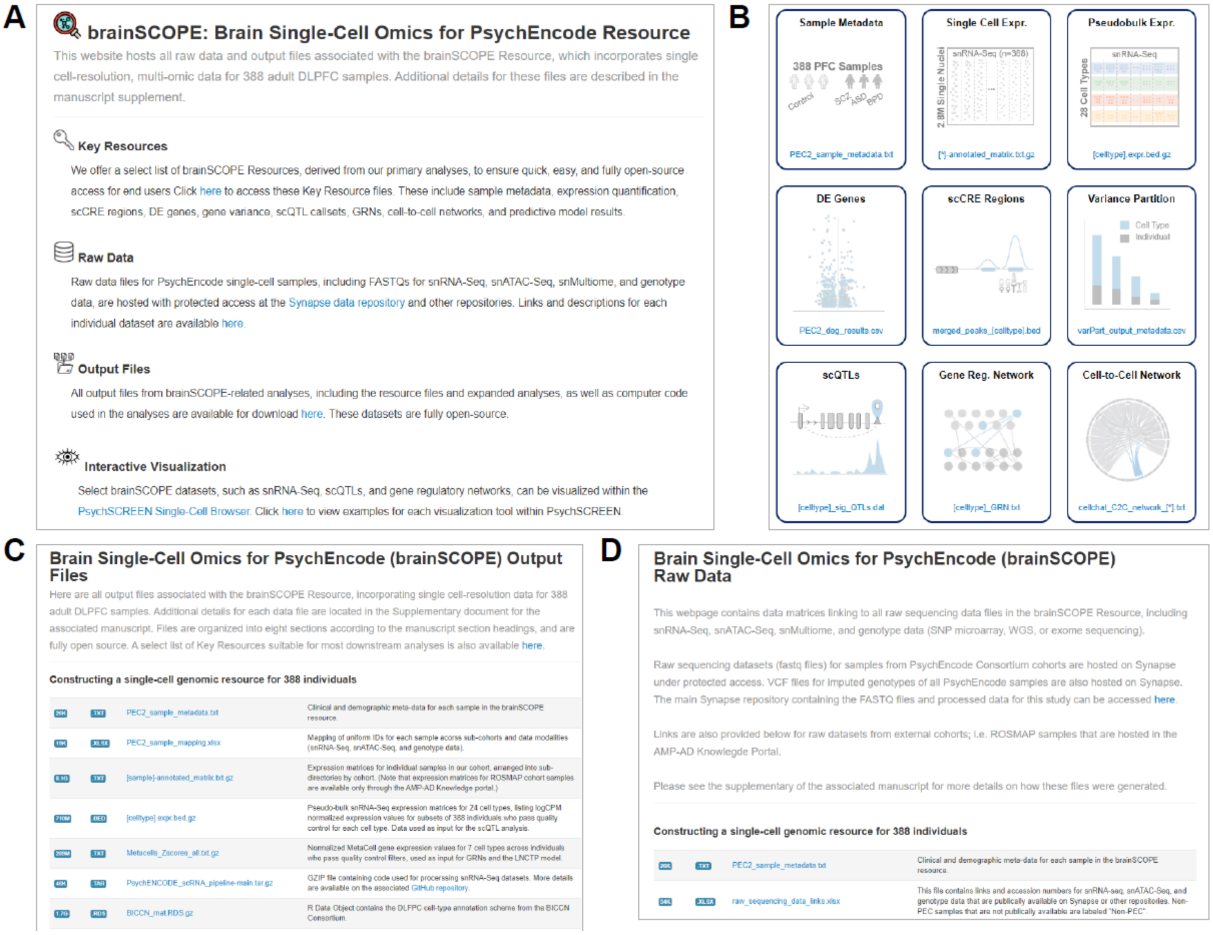


**Fig. S3. Screenshots from the brainSCOPE portal.**

Screenshots of the portal hosting output files and links to raw files and data visualization tools, available at http://brainscope.psychencode.org and https://brainscope.gersteinlab.org. **(A)** shows the landing page to the brainSCOPE portal, and **(B)** shows an excerpt of the webpage highlighting selected key resource files. **(C)** shows an excerpt of the webpage that links to all output files described in the supplement, and **(D)** shows an excerpt of the webpage that links to all raw datasets used in the manuscript (including protected-access datasets).

Further details on the portal are described in the supplementary section "***Portal Overview.***" This supplementary figure relates to **Fig. 1** and main text section "Constructing a single-cell genomic resource for 388 individuals."

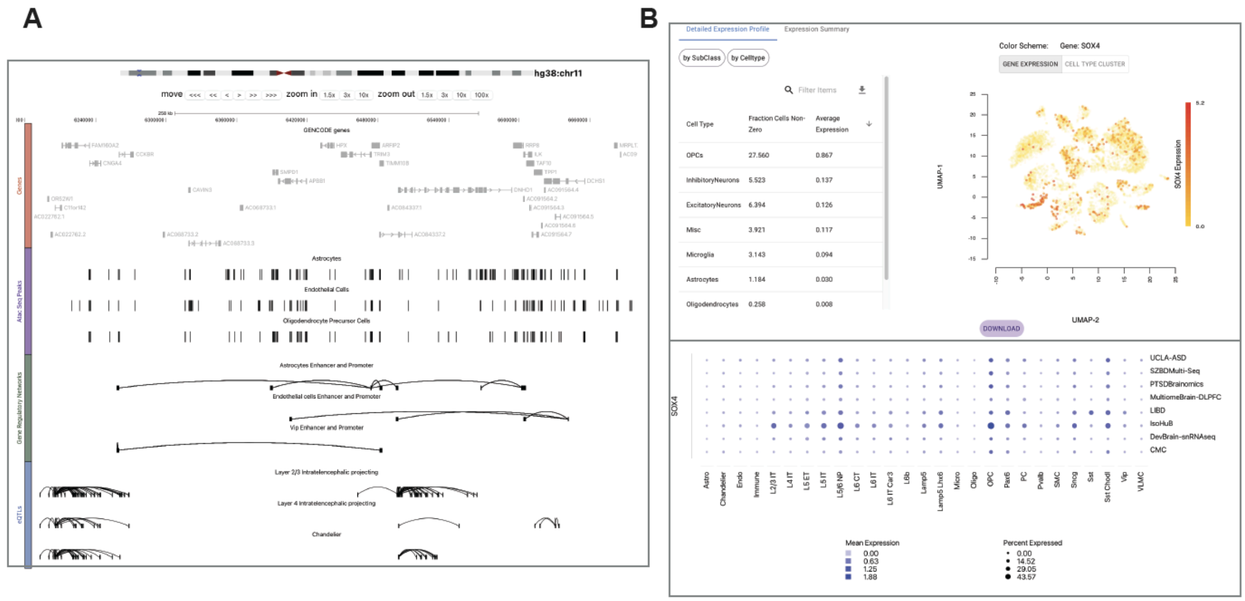


**Fig. S4. Screenshots from the PsychSCREEN data visualization portal.**

This figure shows screenshots of the online visualization tool for key brainSCOPE resources, available at https://psychscreen.wenglab.org/psychscreen/single-cell. **(A)** Integrated genome browser shows (top to bottom) tracks for genes, cell-type-specific snATAC-seq peaks, enhancer and promoter regions within cell-type GRNs (arcs represent links between enhancers and promoters), and links between eGenes and eSNPs for cell-type-specific scQTLs. Page available at https://psychscreen.wenglab.org/psychscreen/single-cell/datasets/scATAC-Seq-peaks. **(B)** Panel shows interactive UMAP tool used to visualize expression of *SOX4* across cells in the SZBDMulti-Seq cohort, and a dot plot visualizing *SOX4* expression across all subcohorts per cell type. Page available at https://psychscreen.wenglab.org/psychscreen/gene/SOX4.

Further details on PsychSCREEN are described in the supplementary section "***Portal Overview.***" This supplementary figure relates to **Fig. 1** and main text section "Constructing a single-cell genomic resource for 388 individuals."

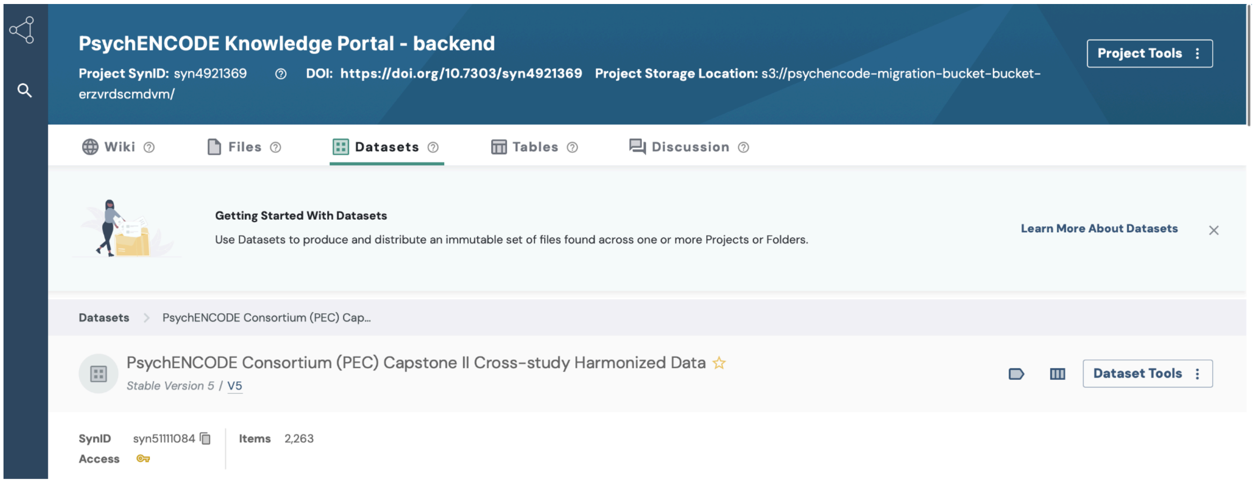


**Fig. S5. Screenshots from the protected-access repository for raw brainSCOPE datasets.**

Screenshot of the PsychENCODE knowledge portal for storage of protected raw sequencing data. This page is available at https://www.synapse.org/#!Synapse:syn51111084.5 at the time of publication.

Further details on the data portal are described in the supplementary section "***Portal Overview.***" This supplementary figure relates to **Fig. 1** and main text section "Constructing a single-cell genomic resource for 388 individuals."

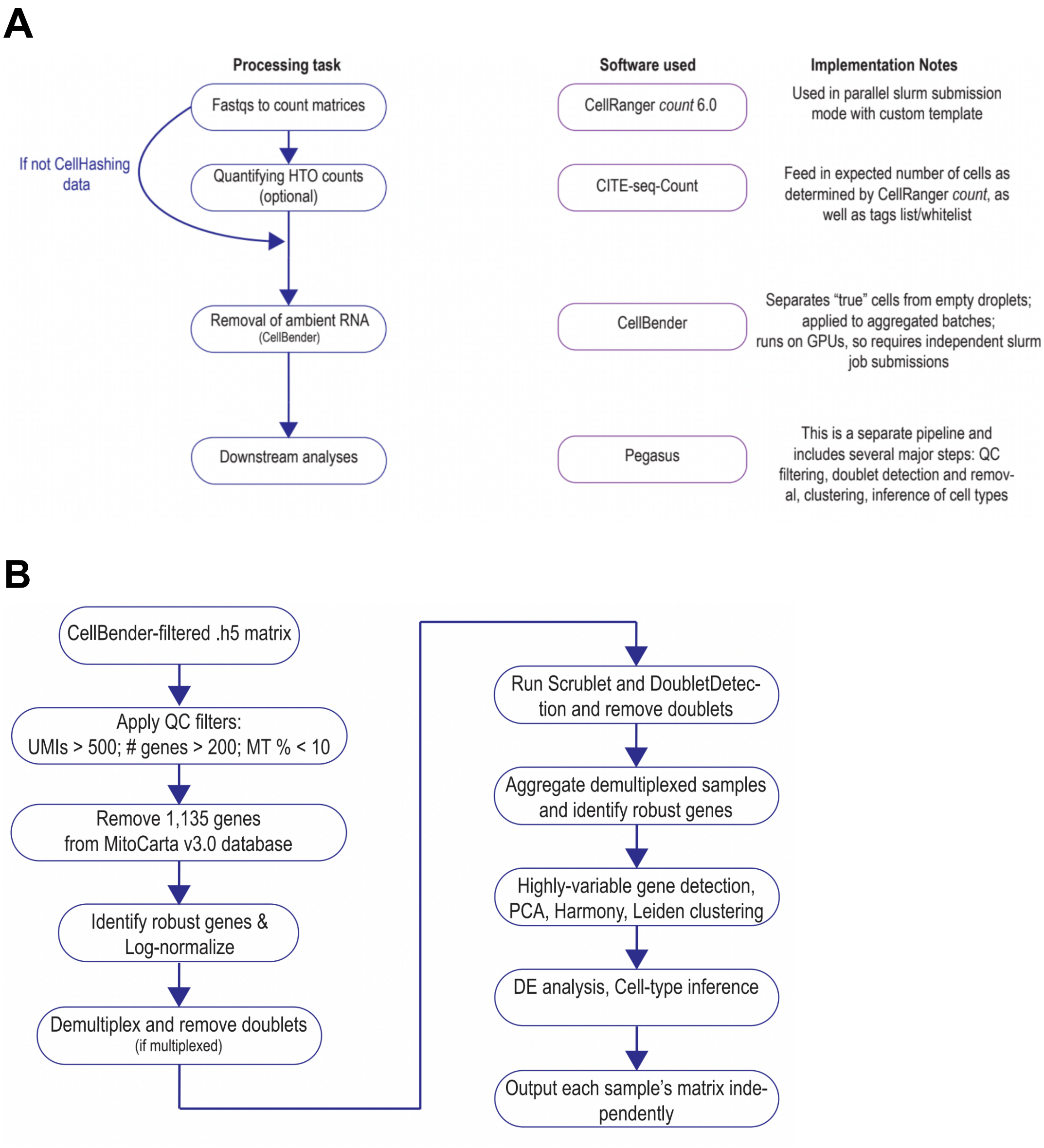


**Fig. S6. Pegasus snRNA-seq workflows.**

Schematic diagrams for **(A)** single-cell processing workflow and **(B)** analysis in Pegasus.

More detail in the supplementary section "***snRNA-seq Processing*.**" This supplementary figure relates to **Fig. 1** and main text section "Constructing a single-cell genomic resource for 388 individuals."

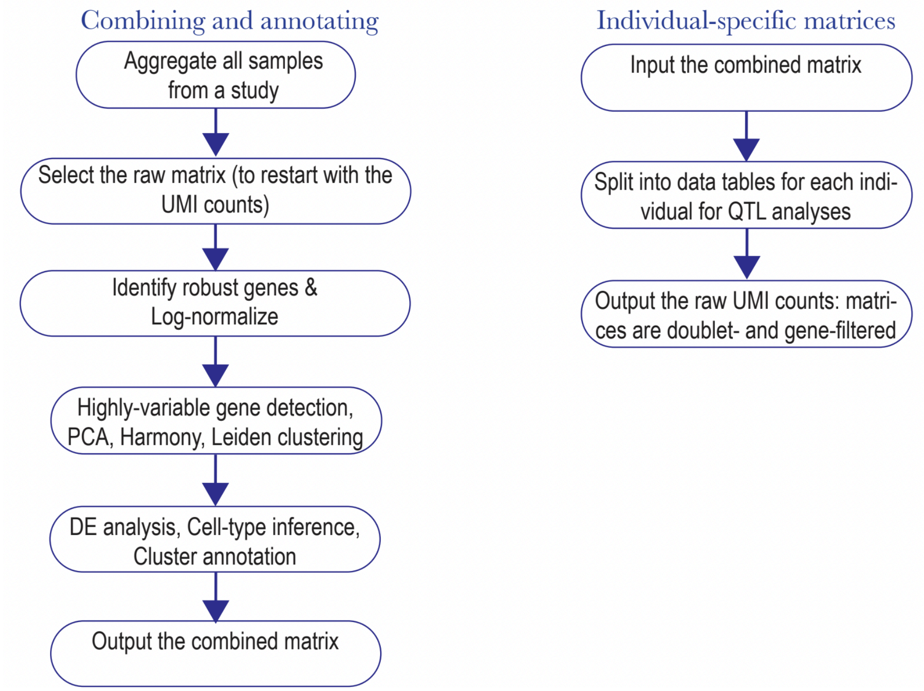


**Fig. S7. Schematic for per-study-based single-cell analysis.**
More detail in the supplementary section "***snRNA-seq Processing*.**" This supplementary figure relates to **Fig. 1** and main text section "Constructing a single-cell genomic resource for 388 individuals."

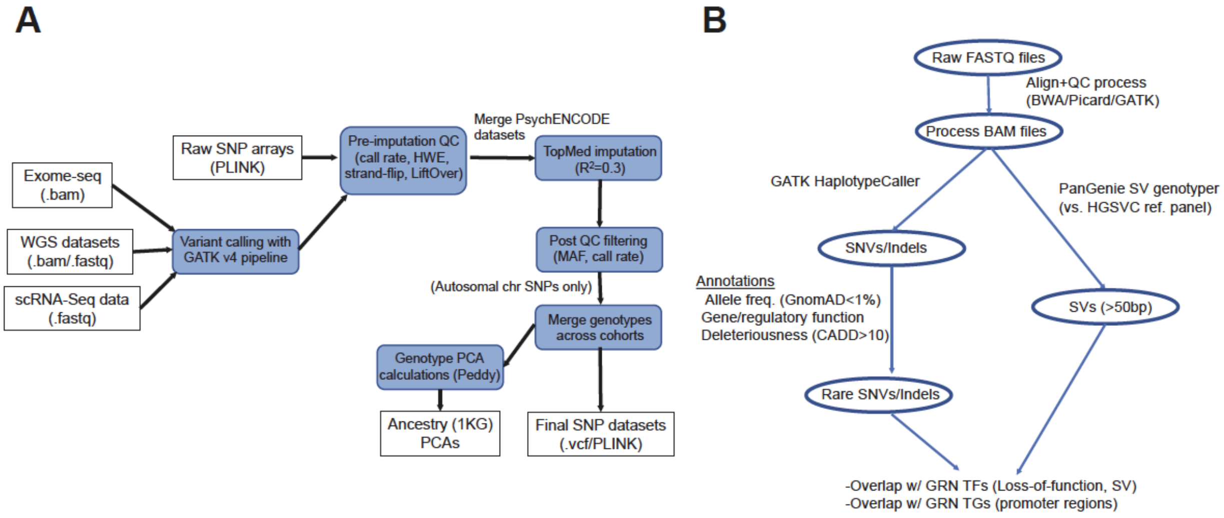


**Fig. S8. Genotype processing and rare variant calling pipeline.**
**(A)** Workflow for brainSCOPE cohort genotype processing, QC, imputation, and PC/ancestry calculation. **(B)** Workflow for detecting rare SNVs, indels, and SVs in samples with available WGS data.

More detail in the supplementary section "***Genotype Processing.***" This supplementary figure relates to **Fig. 1** and main text section "Constructing a single-cell genomic resource for 388 individuals."

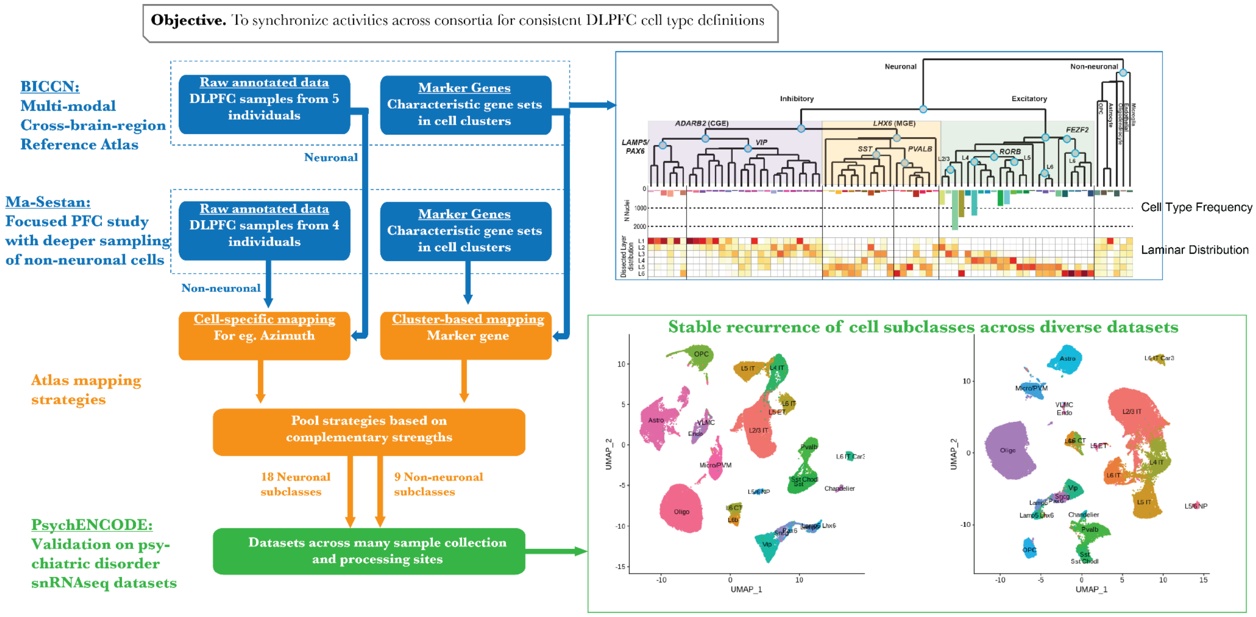


#### Fig. S9. Hybrid cell annotation scheme.

Outline of methods used to generate the hybrid cell annotation scheme, merging neuronal types from the BICCN and the non-neuronal types from (*19*).

More detail in the supplementary section "***snRNA-seq Processing***." This supplementary figure relates to **Fig. 1** and main text section "Constructing a single-cell genomic resource for 388 individuals."

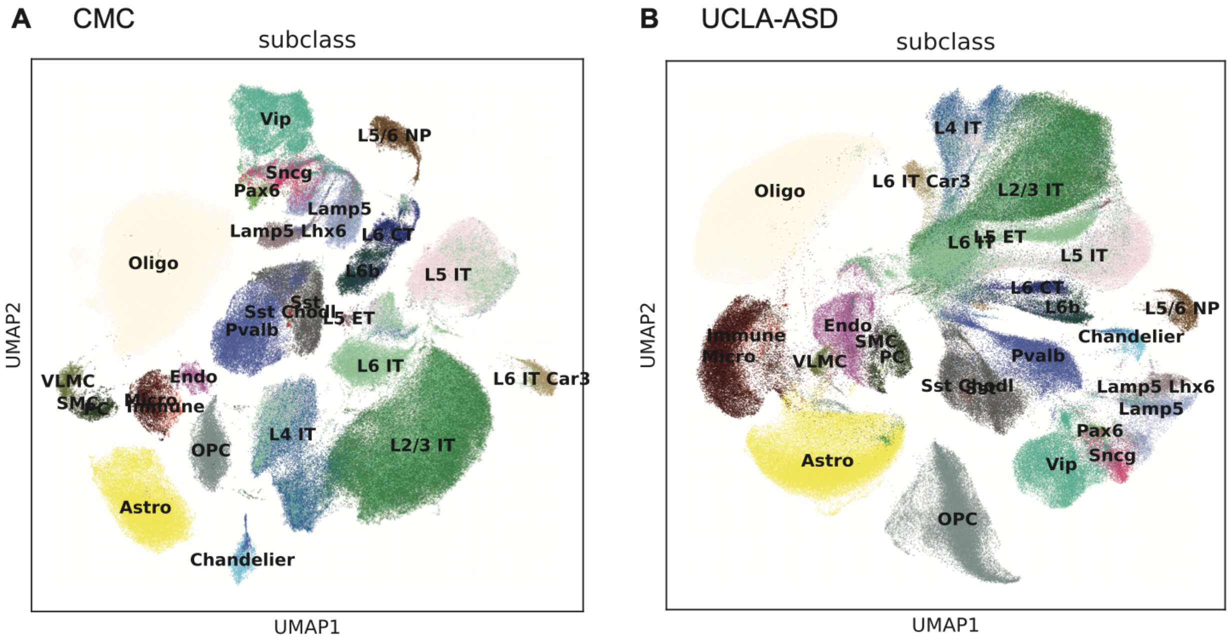


**Fig. S10. UMAPs based on the CMC and UCLA-ASD snRNA-seq cohorts.**

The UMAPs for two study cohorts are shown with the corresponding cell types annotated over the clusters. The same processing and annotation scheme is used as for the SZBDMulti-Seq cohort (shown in **Fig. 1B**). Since we did not generate a unified UMAP across all study cohorts due to expected inter-study batch effects, we present a few examples here. Note that the cell types are well-represented in multiple independent annotation results. Results are shown for the **(A)** CMC and **(B)** UCLA-ASD cohorts.

More detail in the supplementary section "***snRNA-seq Processing***." This supplementary figure relates to **Fig. 1** and main text section "Constructing a single-cell genomic resource for 388 individuals."

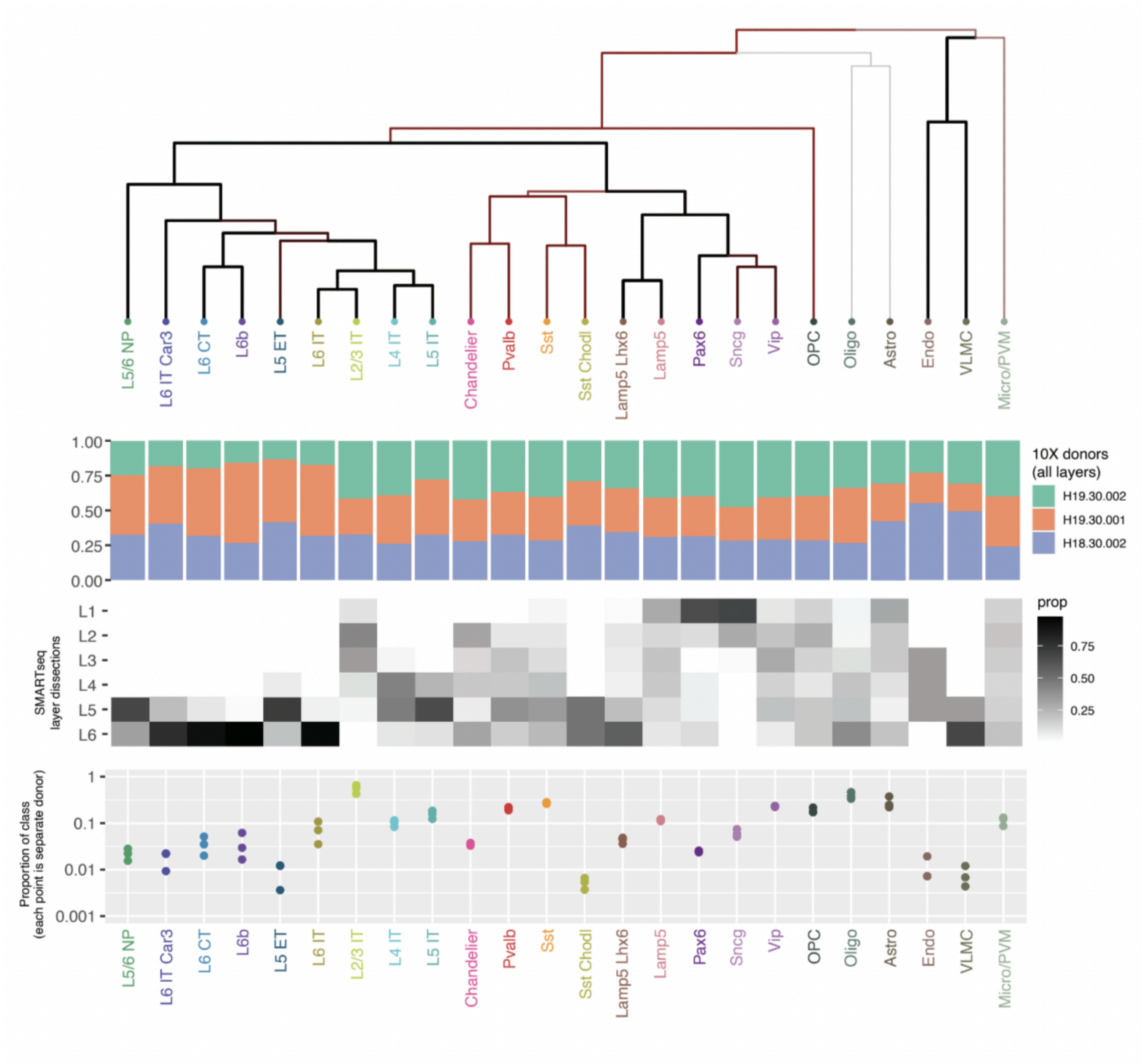


**Fig. S11. BICCN PFC subclass taxonomy.**

The schematic shows the taxonomy used in the harmonized cell annotation scheme to label the neuronal subclasses (*128*). The top panel shows a taxonomy of neuronal and non-neuronal subclasses in the PFC. The second panel indicates the fractional contributions of the donors to the total number of cells in each subclass. The third panel indicates the spatial distribution of the subclasses (or the proportion of cells in each subclass that were dissected from the respective cortical layers based on Smart-seq v4 profiling). The fourth panel indicates the abundance of each subclass within each cell class (excitatory neurons, inhibitory neurons, and non-neuronal cells) where each donor is represented by a separate point.

More detail in the supplementary section "***snRNA-seq Processing***." This supplementary figure relates to **Fig. 1** and main text section "Constructing a single-cell genomic resource for 388 individuals."

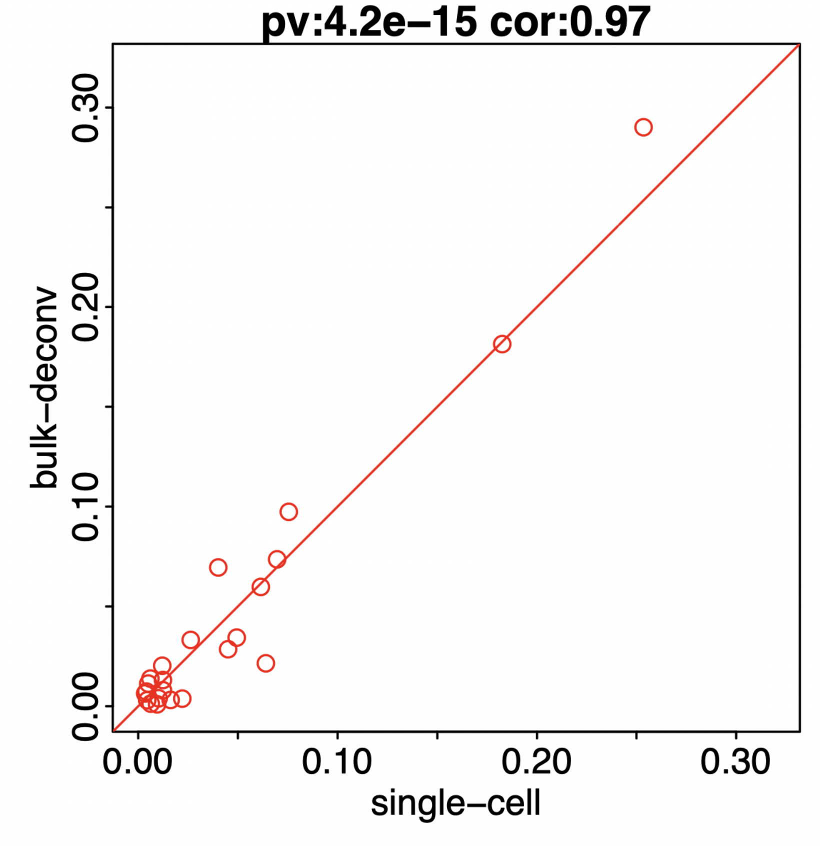


**Fig. S12. Correlation between snRNA-seq cell-type fraction and deconvolved bulk-RNA-seq cell-type fraction.**

Each dot represents a cell type. The red line is the diagonal (y=x) line. The correlation of the cell-type fraction of common samples estimated by snRNA-seq and CMC bulk RNA-seq is 0.97 with $p=4.2\times10^{-15}$. The p-value is based on a correlation test.

More detail in the supplementary section "***Cell-type Fractions***." This supplementary figure relates to **Fig. 1** and main text section "Constructing a single-cell genomic resource for 388 individuals."

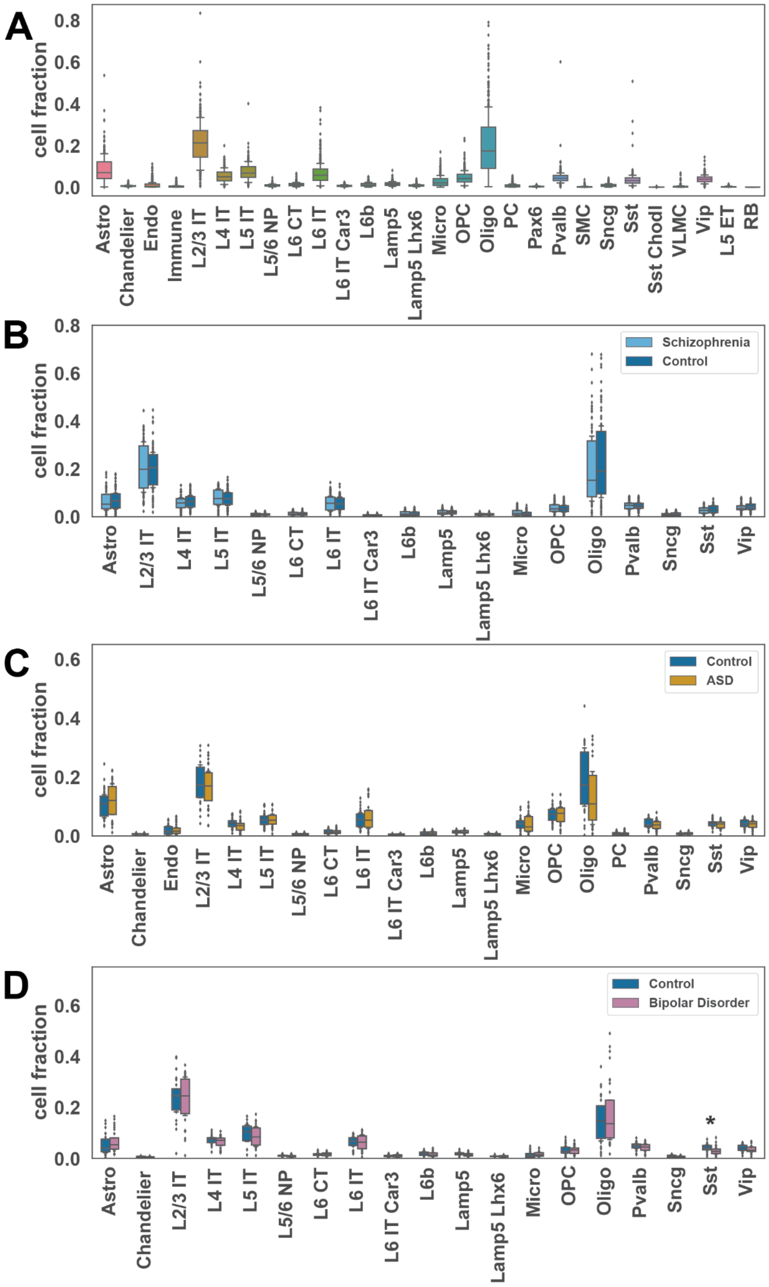


**Fig. S13. Single-cell fractions by cell type and primary diagnosis.**

**(A)** Single-cell cell-type fraction distribution for different cell types. **(B)** Comparison of single-cell cell-type fractions for schizophrenia and control samples. **(C)** Comparison of single-cell cell-type fraction for ASD and control samples. **(D)** Comparison of single-cell cell-type fraction for bipolar disorder and control samples. Points represent individual samples. Asterisks denote a significant difference based on two sided Welch's t test and an FDR Benjamini–Hochberg-corrected adjusted p-value <0.05.

More detail in the supplementary section "***Cell-type Fractions*.**" This supplementary figure relates to **Fig. 1** and main text section "Constructing a single-cell genomic resource for 388 individuals."

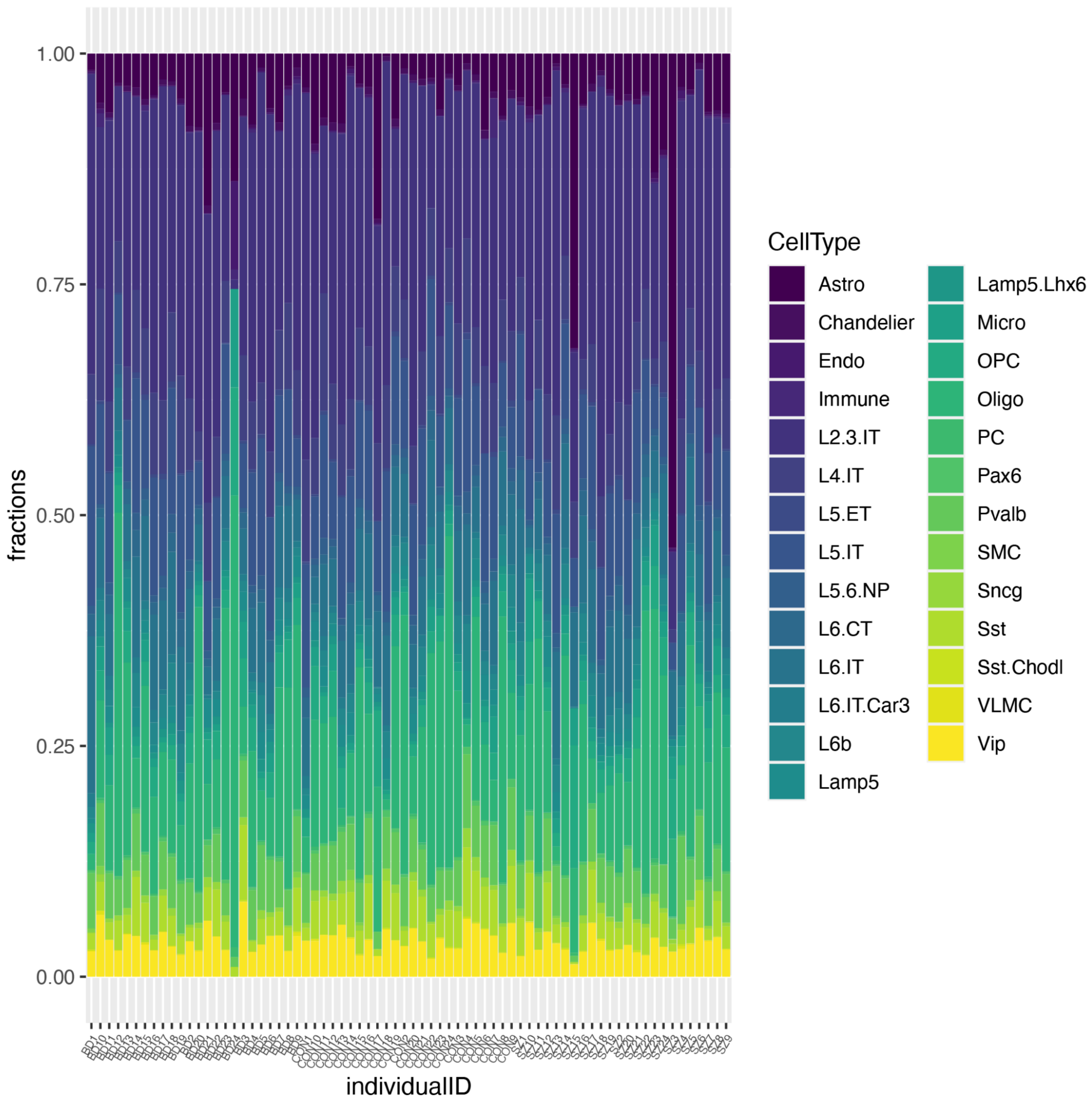


**Fig. S14. Cell-fraction distributions for individuals in the SZBDMulti-seq cohort.**
Cell fractions per individual (72 individuals along the x-axis) in the SZBDMulti-seq cohort are shown for 27 cell types (note that the RB cell type does not appear in any individuals in this cohort).

More detail in the supplementary section "***Cell-type Fractions.***" This supplementary figure relates to **Fig. 1** and main text section "Constructing a single-cell genomic resource for 388 individuals."

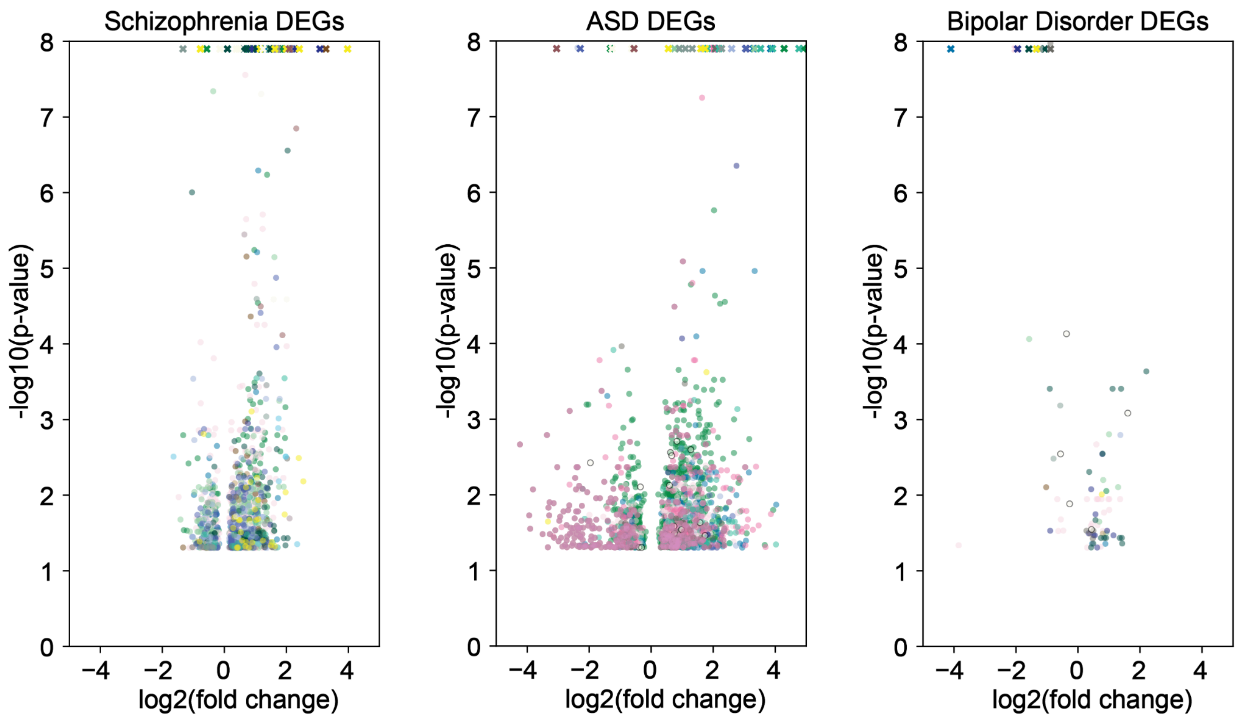


**Fig. S15. Volcano plots show DE genes for schizophrenia, ASD, and bipolar disorder.** (Left) DE genes between schizophrenia and control samples. (Middle) DE genes between ASD and control samples. (Right) DE genes between bipolar disorder and control samples. Genes with an abs($log_2$fold) > 0.1 and adjusted p < 0.05 are defined as DE genes. Each dot represents one DE gene in each cell type, colored following the color code shown in **Fig. 1A**. Values where -log(p) are >8 are shown as an "x".

More detail in the supplementary section "***DE analysis*.**" This supplementary figure relates to **Fig. 1** and main text section "Constructing a single-cell genomic resource for 388 individuals."

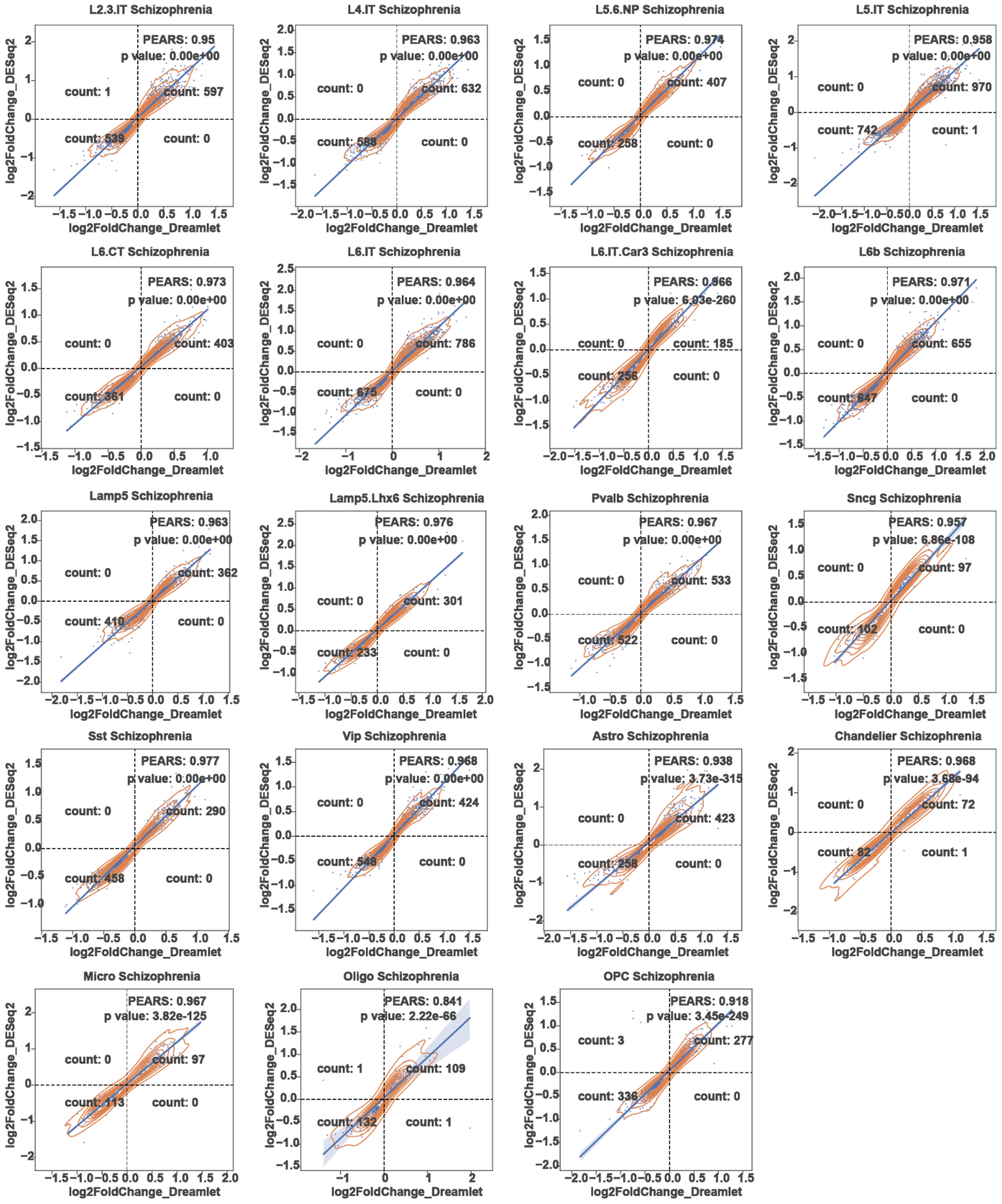


**Fig. S16. Comparison of schizophrenia DE Genes from DESeq2 and Dreamlet.**

Scatter plots show inner join of genes with p-value < 0.05 calculated by two methods, DESeq2 and Dreamlet. x-axis represents the $\log_2$ fold change calculated by Dreamlet. y-axis represents the $\log_2$ fold change calculated by DESeq2. Pearson correlations, p-values, and gene counts in each quadrant are labeled in each panel.

More detail in the supplementary section "***DE analysis*.**" This supplementary figure relates to **Fig. 1** and main text section "Constructing a single-cell genomic resource for 388 individuals."

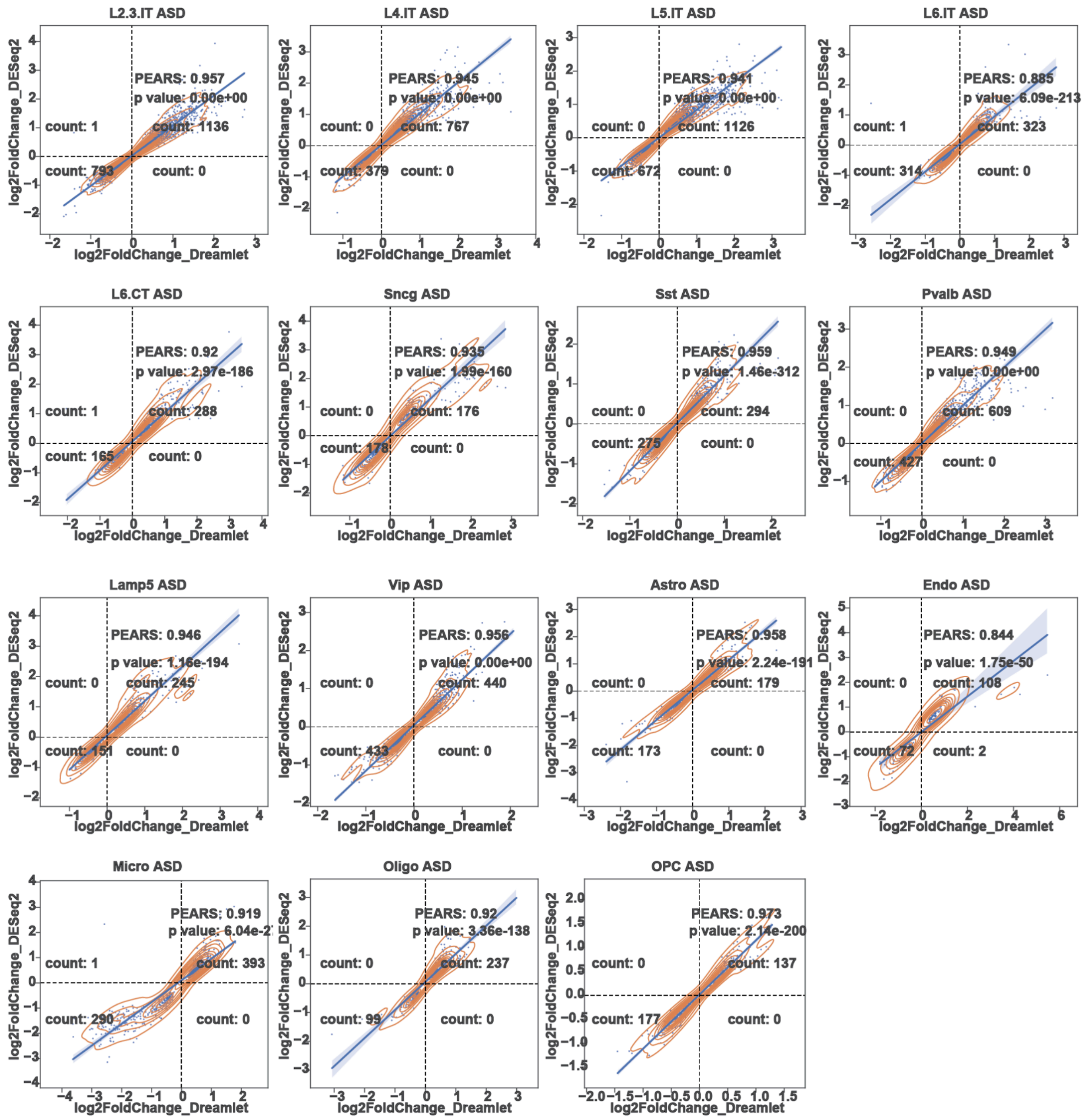


**Fig. S17. Comparison of ASD DE Genes from DESeq2 and Dreamlet.**

Scatter plots show inner join of genes with p-value < 0.05 calculated by two methods, DESeq2 and Dreamlet. x-axis represents the $\log_2$ fold change calculated by Dreamlet. y-axis represents the $\log_2$ fold change calculated by DESeq2. Pearson correlations, p-values, and gene counts in each quadrant are labeled in each panel.

More detail in the supplementary section "***DE analysis.***" This supplementary figure relates to **Fig. 1** and main text section "Constructing a single-cell genomic resource for 388 individuals."

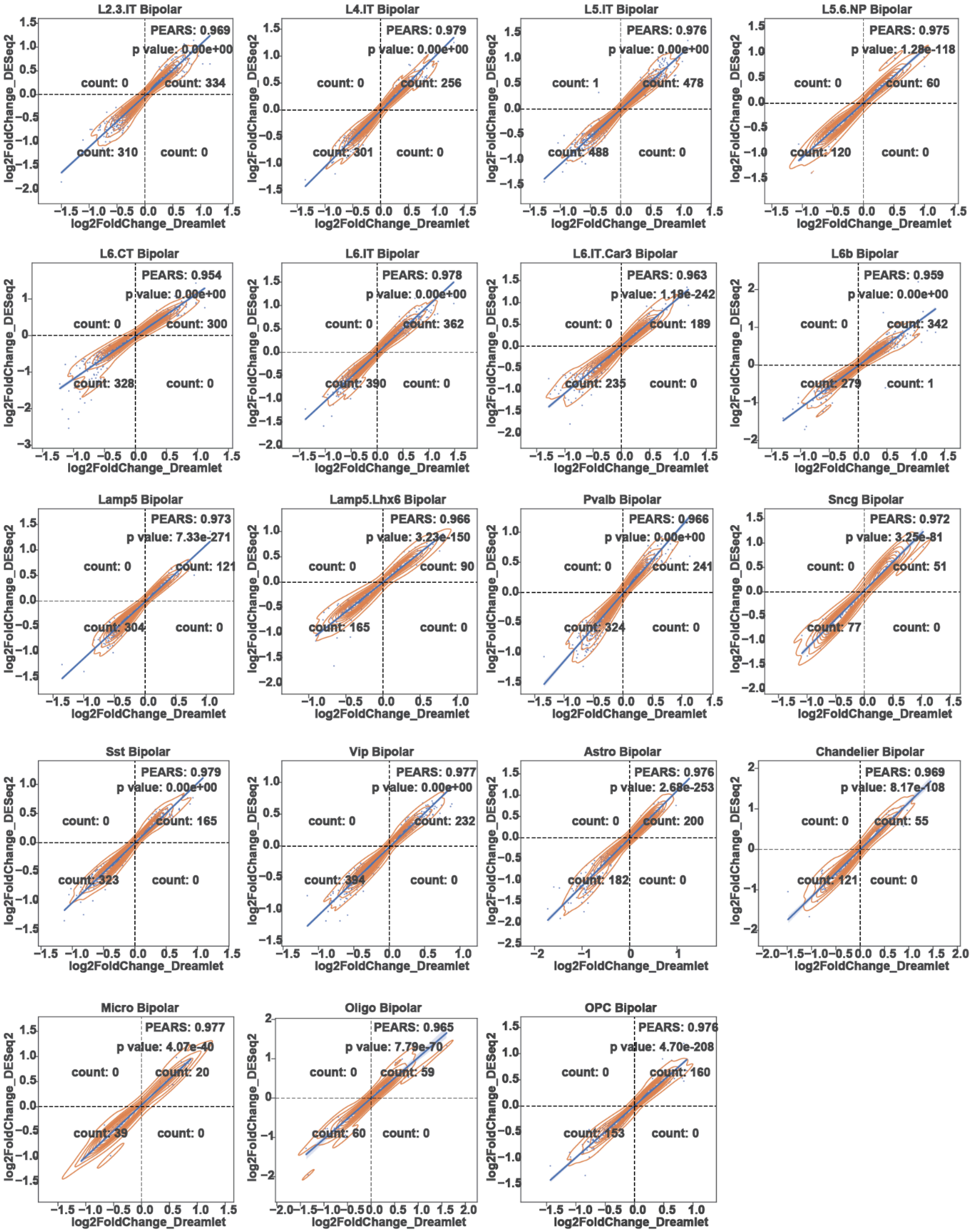


**Fig. S18. Comparison of bipolar DE Genes from DESeq2 and Dreamlet.**

Scatter plots show inner join of genes with p-value < 0.05 calculated by two methods, DESeq2 and Dreamlet. x-axis represents the $\log_2$ fold change calculated by dreamlet. y-axis represents the $\log_2$ fold change calculated by DESeq2. Pearson correlations, p-values, and gene counts in each quadrant are labeled in each panel.

More detail in the supplementary section "***DE analysis.***" This supplementary figure relates to **Fig. 1** and main text section "Constructing a single-cell genomic resource for 388 individuals."

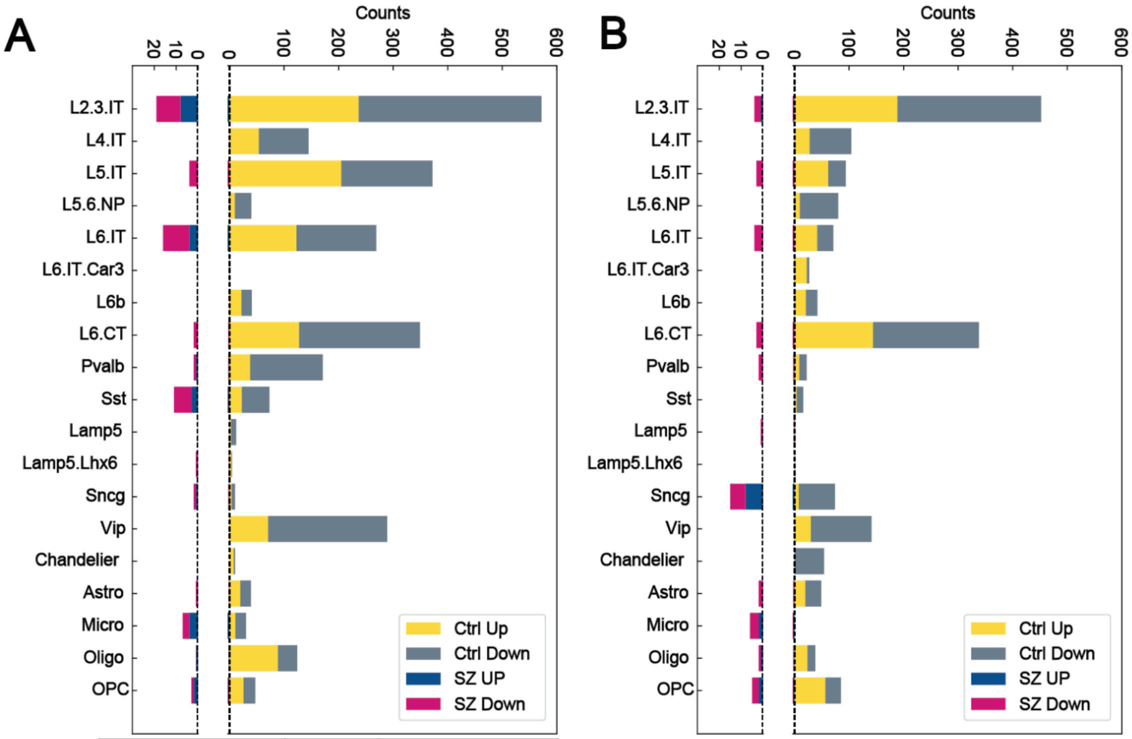


| Num of Sample | Control | Schizophrenia |
|---|---|---|
| old>=70 | 41 | 37 |
| young<70 | 35 | 34 |
| | 76 | 71 |

| Num of Sample | Control | Schizophrenia |
|---|---|---|
| old>=70 | 36 | 32 |
| young<70 | 30 | 29 |
| | 66 | 61 |

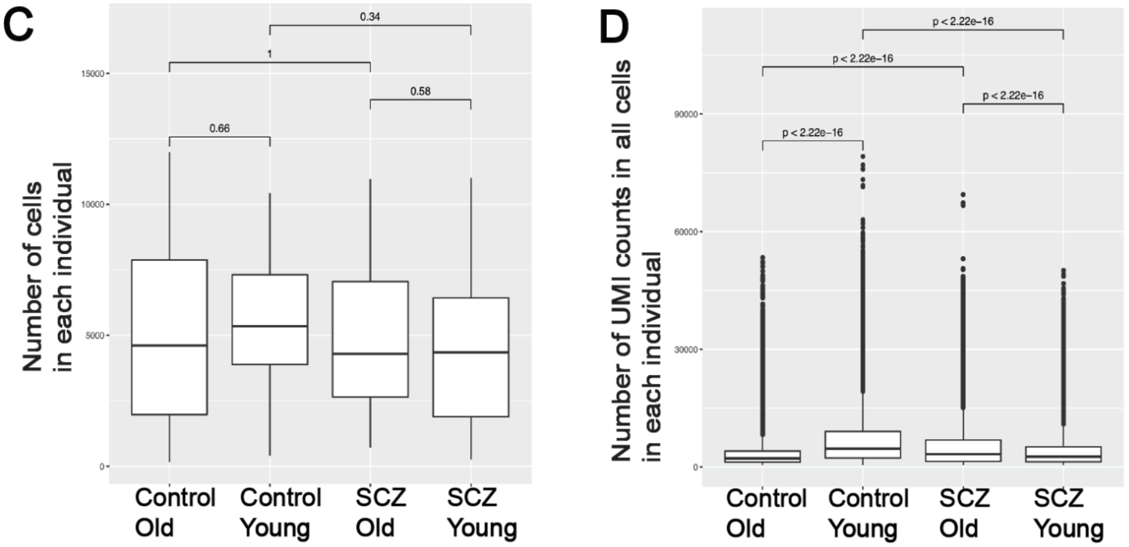


**Fig. S19. Comparison of aging DE Genes between schizophrenia patients and healthy individuals.**

(**A**) Number of up (yellow) and down (gray) regulated aging DE genes (old vs. young) in each cell type in healthy individuals. The number of up (blue) and down (red) regulated aging DE genes in the schizophrenia patient group is smaller than in the control group. Number of samples used in the calculation is showed at the bottom. (**B**) For a permutation analysis, we dropped five samples in each comparison group (control young, control old, schizophrenia young, schizophrenia old) and recalculated the aging DE genes. We observed a similar pattern, where schizophrenia patients have less aging DE genes compared with healthy groups. The number of samples used in the permutation analysis is presented at the bottom.

In addition to the permutation analysis, we explored technical and biological covariates that might potentially affect the results. We explored the distribution of the number of cells per individual and the UMI count. (**C-D**) Technical metrics for the CMC snRNA-seq cohort as a function of age and schizophrenia diagnosis. The groups of individuals are: controls in the younger (age < 70 yr) age group, “control_Younger”; controls in the older (age >= 70 yr) age group, “control_Older”; individuals diagnosed with schizophrenia in the younger age group, “Schizophrenia_Younger”;  individuals diagnosed with schizophrenia in the older age group, “Schizophrenia_Older". (**C**) Distributions of the number of cells in the expression matrices for each individual (across all cell types). (**D**) Distributions of the number of UMI counts in all cells in the expression matrices for each individual (across all cell types). P-values for comparisons in **C** and **D** are obtained from the Wilcoxon rank-sum test. We did not observe a substantial change in the number of cells between different groups. We do observe a difference in UMI counts per individual among the groups. Differences in the UMI counts among groups could be due to biological or technical reasons; thus, we included this as a covariate when we characterized the aging DE genes.

More details in supplementary section 7.3 "***Aging DE.***" This supplementary figure relates to **Fig. 1** and main text section "Constructing a single-cell genomic resource for 388 individuals."

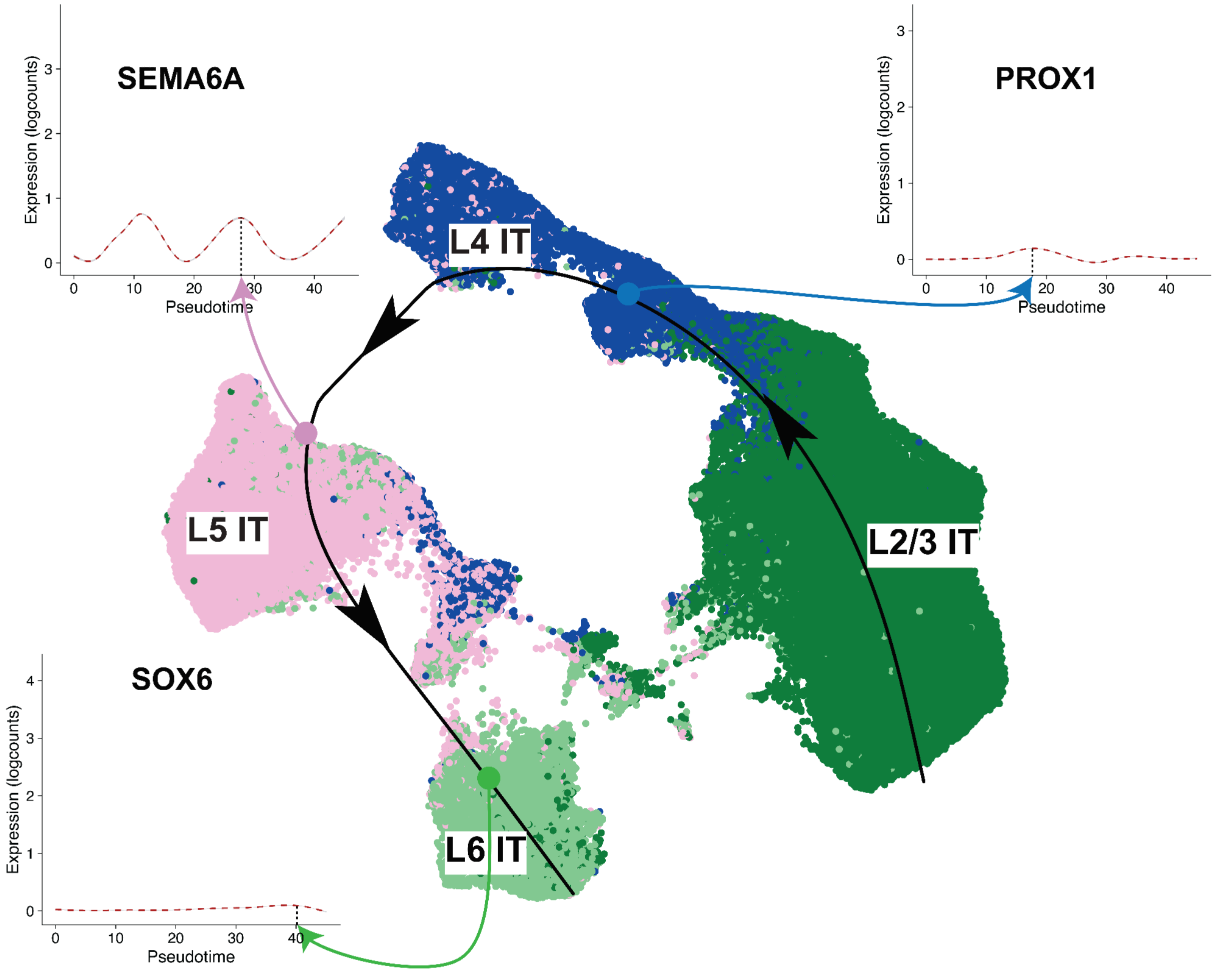


**Fig. S20. Pseudotime trajectory patterns for three significant genes in the SZBDMulti-seq cohort.**

UMAP plot shows predicted trajectory for excitatory IT neurons in adult control samples from the SZBDMulti-Seq cohort. The predicted trajectory proceeds along the cortical layer dimension from L2/3 to L6 in the PFC. Smoothed line plot insets highlight log-normalized gene expression in cells along the pseudotime axis for three genes: *SEMA6A*, *PROX1*, and *SOX6*. Significance was assessed as FDR < 0.05 (Wald test) and overlapped across 5 cohorts.

More detail in the supplementary section "***Trajectory Analysis***." This supplementary figure relates to **Fig. 1** and main text section "Constructing a single-cell genomic resource for 388 individuals."

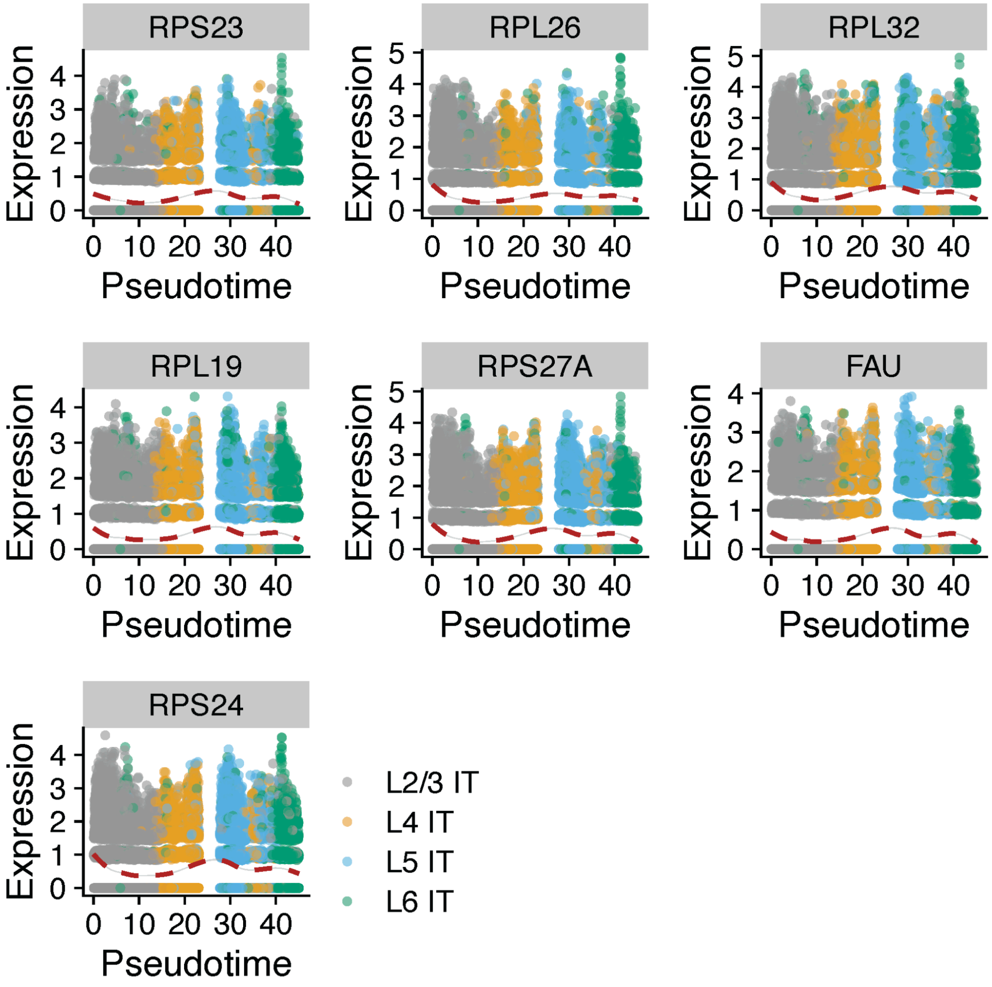


**Fig. S21. Pseudotime trajectory patterns for ribosomal genes in the SZBDMulti-seq cohort.**

Gene expression profiles for seven ribosomal protein genes whose expression significantly varies along the trajectory line across IT neurons. Dashed pseudotime plots and scatter plots of gene expression in individual cells are shown for individuals in the SZBDMulti-seq cohort. Significance was assessed as FDR < 0.05 (Wald test) and overlapped across 5 cohorts.

More detail in the supplementary section "***Trajectory Analysis*.**" This supplementary figure relates to **Fig. 1** and main text section "Constructing a single-cell genomic resource for 388 individuals."

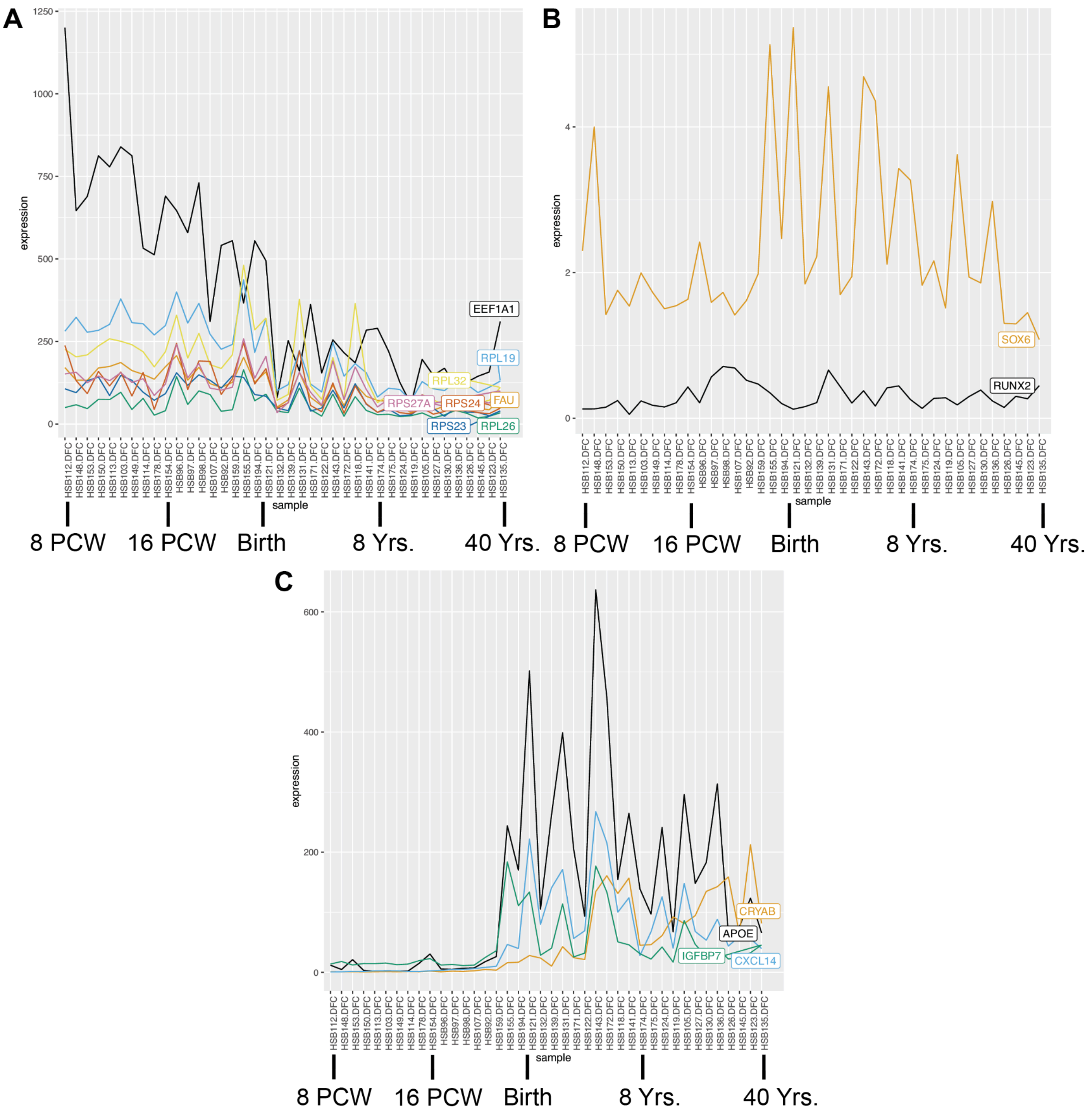


**Fig. S22. Bulk-tissue gene expression patterns from BrainSpan for select trajectory analysis genes, as a function of developmental stage.**

Shown are the bulk-tissue expressions of genes (in Transcripts Per Million) for samples ordered along the developmental trajectory. The sample ages at time of death range from 8 Post-Conception Weeks (PCW) to 40 years of age. On each panel, we mark a few points along the developmental trajectory. Note the different scales for each panel. **(A)** Genes related to translation: *EEF1A1*, *RPS23*, *RPL26*, *RPL32*, *RPL19*, *RPS27A*, *FAU*, and *RPS24*, where the last seven are ribosomal protein-encoding genes (see also **fig. S21**). **(B)** TF-encoding genes: *SOX6* and *RUNX2*. **(C)** Additional genes from the significant trajectory gene set with notable changes in expression along the developmental trajectory: *APOE*, *CRYAB*, *CXCL14*, and

*IGFBP7*. Significance was assessed as FDR < 0.05 (Wald test) and overlapped across 5 cohorts.

More detail in the supplementary section "***Trajectory Analysis*.**" This supplementary figure relates to **Fig. 1** and main text section "Constructing a single-cell genomic resource for 388 individuals."

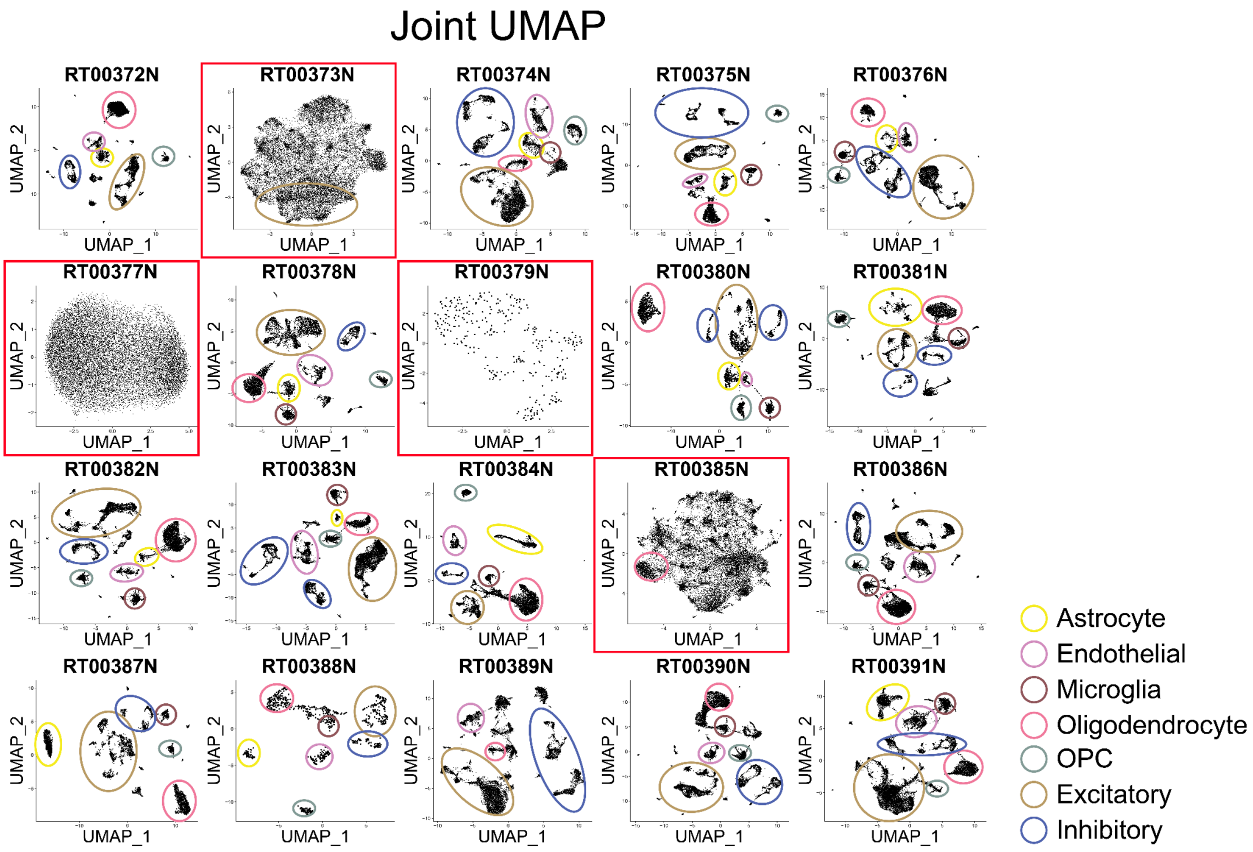


**Fig. S23. snMultiome joint UMAP.**

Comparison of cell-type clustering across individual samples in the snMultiome datasets. Each UMAP shows one sample in the dataset with circled cells representing the cell types found using the corresponding marker genes. Red boxes denote UMAPs with suboptimal snATAC-seq resolution where some or all of the cell types could not be distinguished.

More detail in the supplementary section "***snATAC-seq Processing*.**" This supplementary figure relates to **Fig. 2** and main text section "Determining regulatory elements for cell types from snATAC-seq."

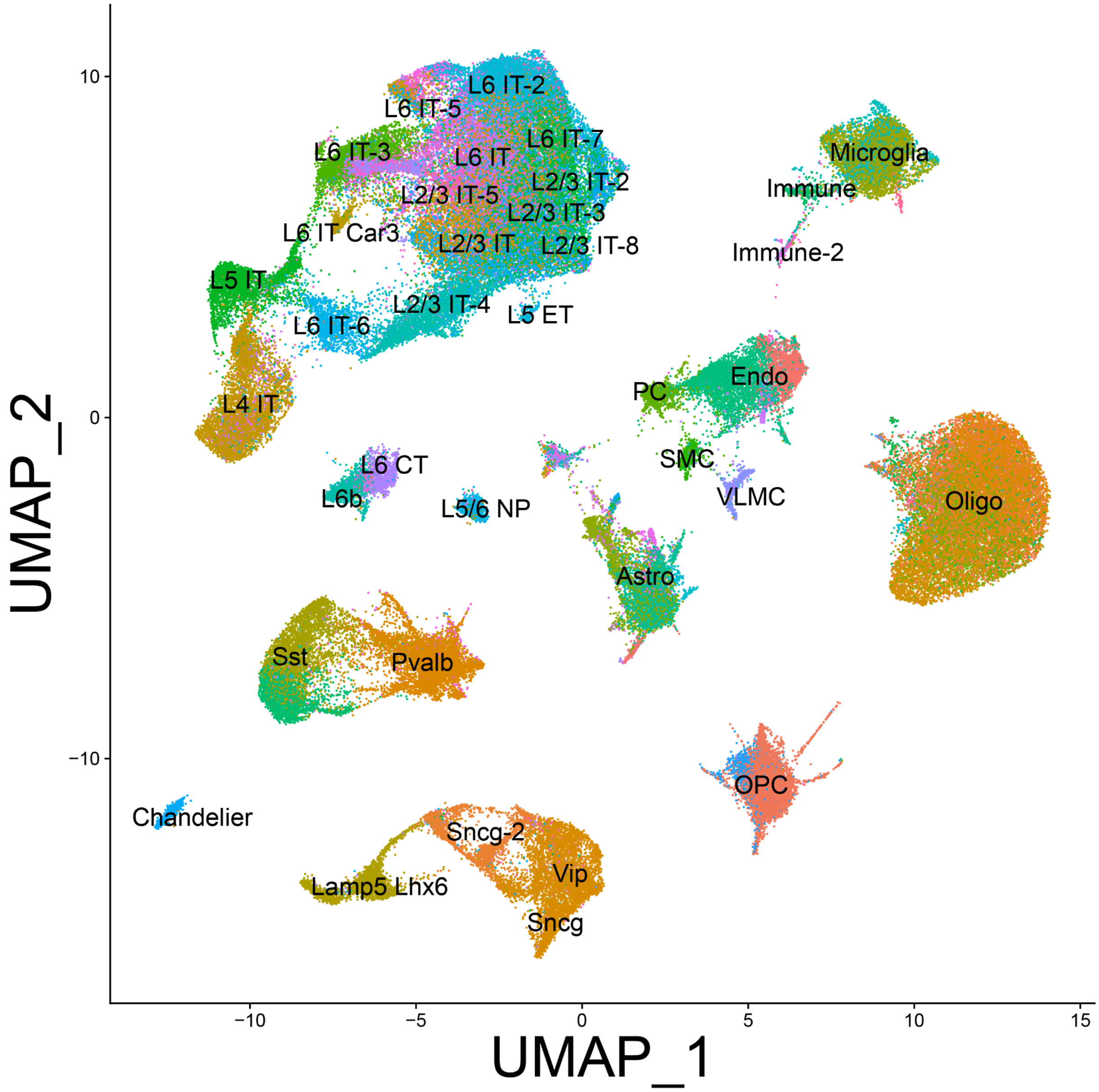


**Fig. S24. snMultiome integrated UMAP.**
UMAP visualization of the integrated snMultiome dataset using only snATAC-seq information with latent semantic indexing.

More detail in the supplementary section "***snATAC-seq Processing***." This supplementary figure relates to **Fig. 2** and main text section "Determining regulatory elements for cell types from snATAC-seq."

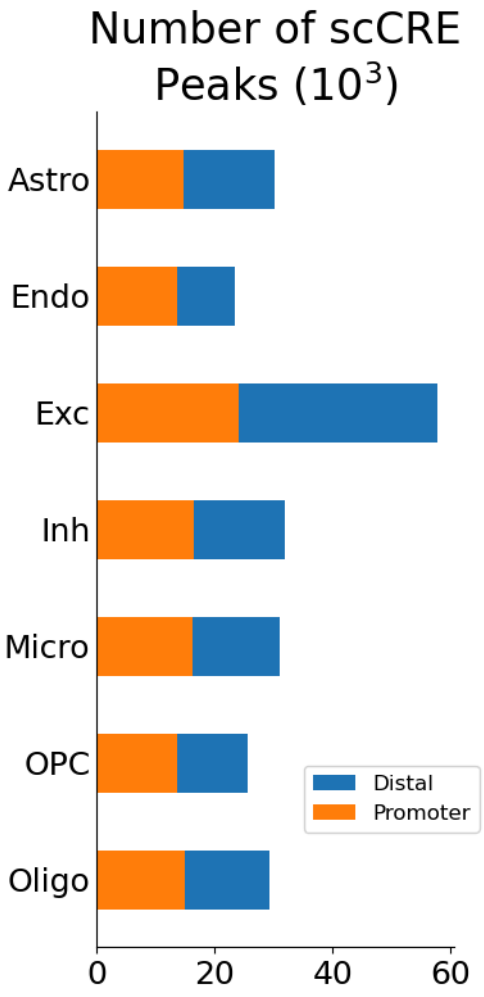


**Fig. S25. scCRE distributions by cell types.**
Distribution of scCRE locations colored by promoter or non-promoter (distal) and grouped by the seven major cell types.

More detail in the supplementary section "***snATAC-seq Processing*.**" This supplementary figure relates to **Fig. 2** and main text section "Determining regulatory elements for cell types from snATAC-seq."

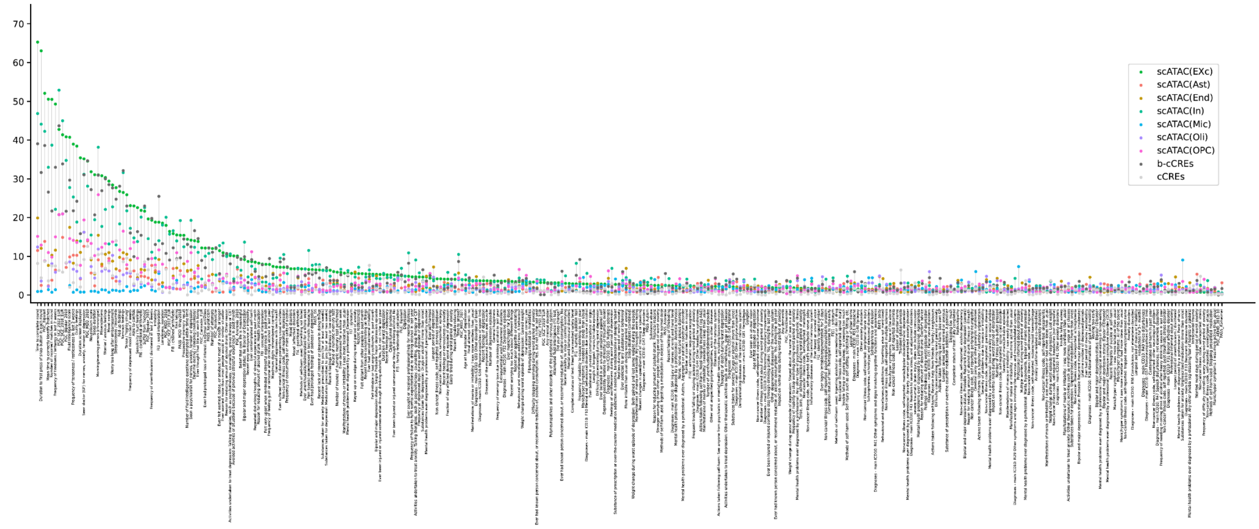


**Fig. S26. LDSC analysis results for all brain-related traits.**

This plot shows -log(p-value) LDSC enrichment scores (y-axis) for 333 brain-related traits that we collected from UKBB, PGC, and PASS (x-axis), against snATAC-Seq peaks for Exc, Astro, Endo, Inh, Micro, Oligo, and OPC cells, as well as b-cCREs and cCREs (colored dots). Data are sorted according to the -log(p-value) of snATAC-Seq for Exc neurons. See data S9 and S10 for a list of enrichment values for all traits.

More detail in the supplementary section "***LDSC***." This supplementary figure relates to **Fig. 2** and main text section "Determining regulatory elements for cell types from snATAC-seq."

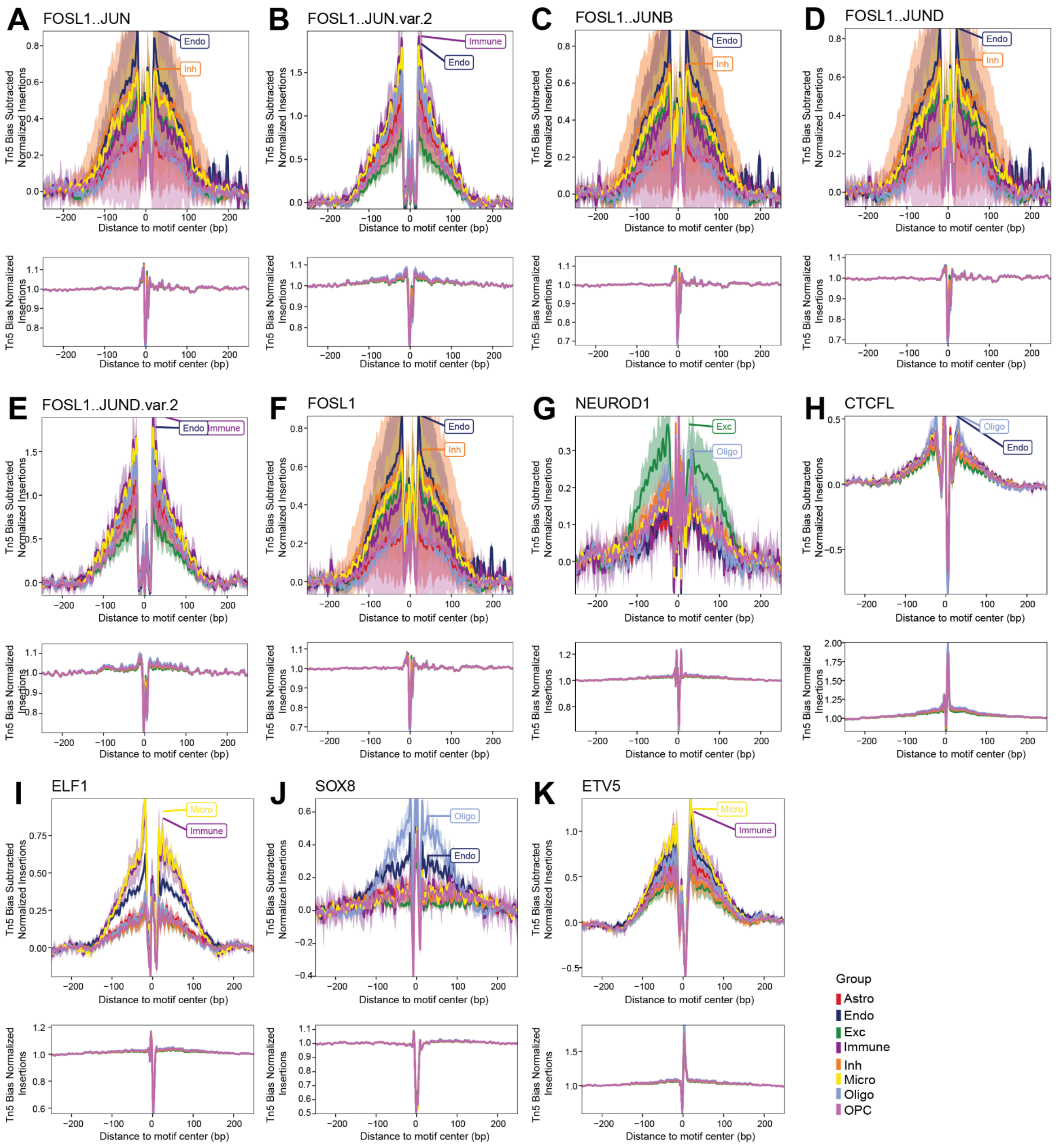


**Fig. S27. Additional footprints for top TFs.**

Subfigures **A-K** show additional footprints of the TFs shown in **Fig. 2F**, drawn in a similar way as the selected footprints in **Fig. 2G**. Note that these plots include an additional cell type (immune) not shown in **Figs. 2F-2G**.

More detail in the supplementary section "***snATAC-seq Processing***." This supplementary figure relates to **Fig. 2** and main text section "Determining regulatory elements for cell types from snATAC-seq."

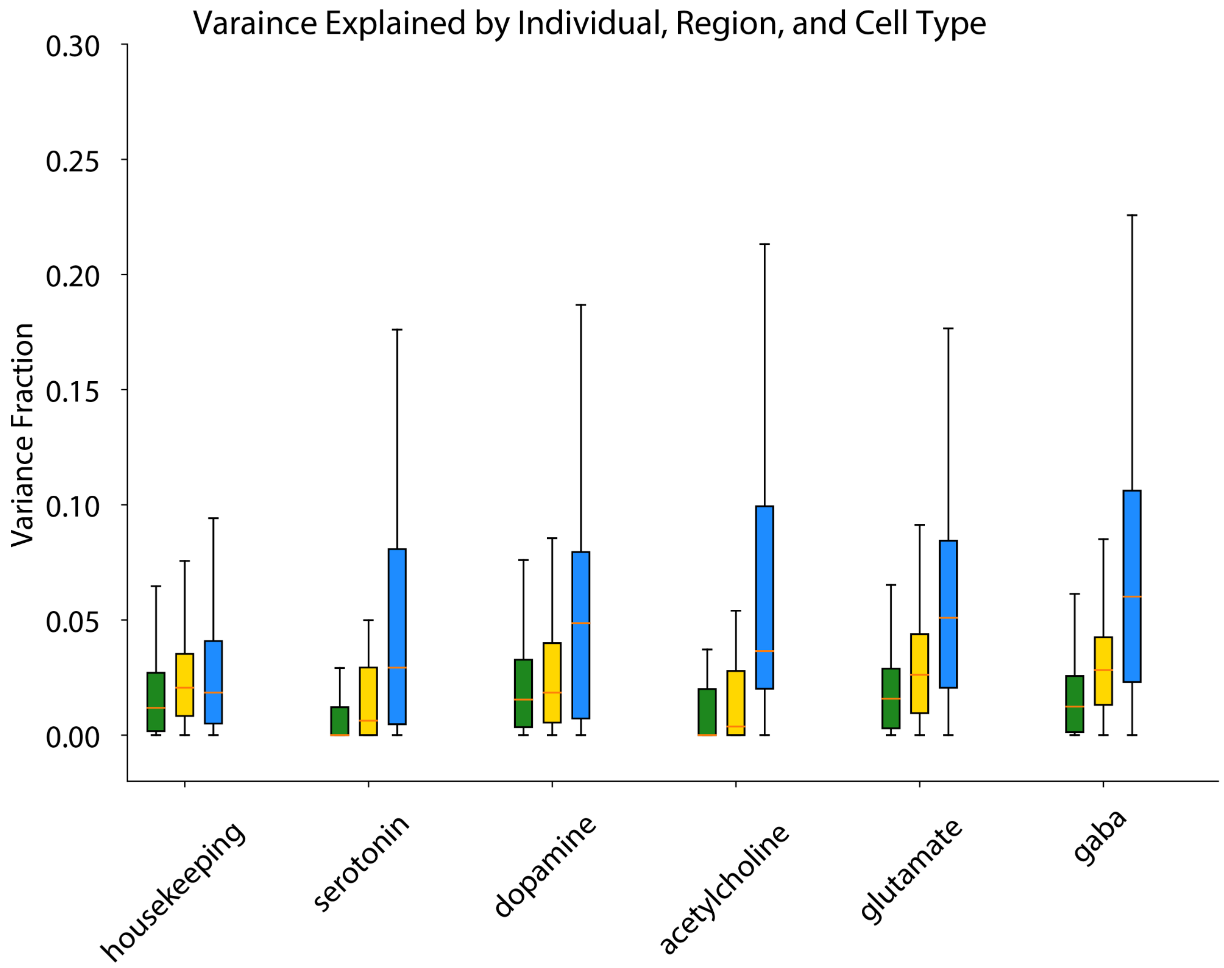


**Fig. S28. Variation partition over three different brain regions.**

Using a similar strategy as in **Fig. 3A-C**, we considered the partitioning of variation among individuals, different brain regions (PFC, parietal lobule, and occipital lobule), and cell types. The box plot shows the distribution of variation fraction explained by individuals (green), brain region (yellow), and cell type (blue) within each group of genes. Due to the constraint of the number of samples, only six individuals with snRNA-seq data for three brain regions collected from the same donor (from Gandal-UCLA and UCLA-ASD cohorts) were included in this analysis.

More detail in the supplementary section "***Variance Partition.***" This supplementary figure relates to **Fig. 3** and main text section "Measuring transcriptome and epigenome variation across the cohort at the single-cell level."

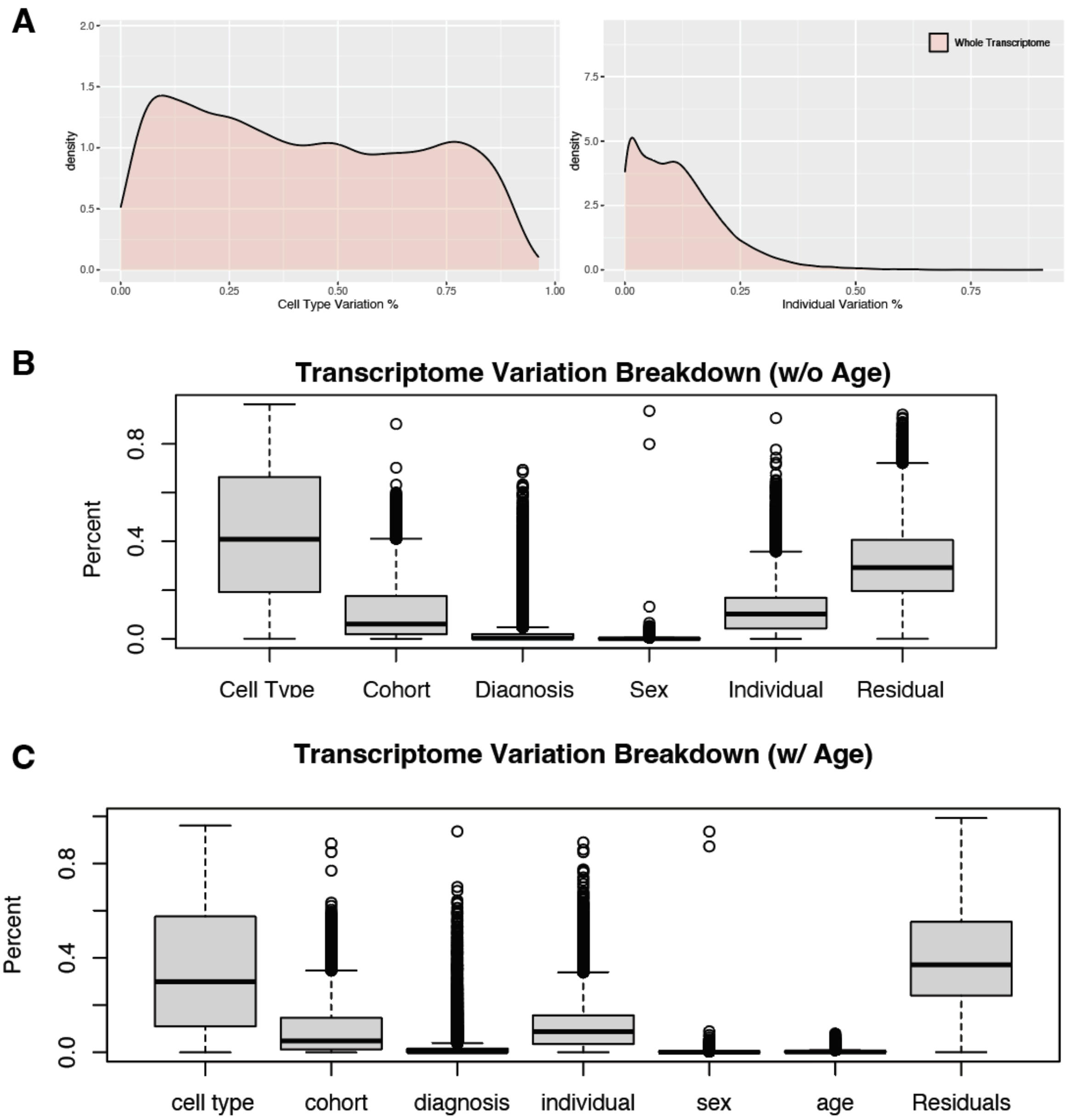


**Fig. S29. Variation partition.**

**(A)** Overall distribution of inter-individual and inter-cell type variation across all genes in the transcriptome. (**B)** Breakdown of transcriptome variation across all genes for cell type, individual, covariates (cohort, diagnosis, and sex), and residual. (**C**) Breakdown of transcriptome variation across all genes, including covariates in **(B)** as well as age.

More detail in the supplementary section "***Variance Partition.***" This supplementary figure relates to **Fig. 3** and main text section "Measuring transcriptome and epigenome variation across the cohort at the single-cell level."

A

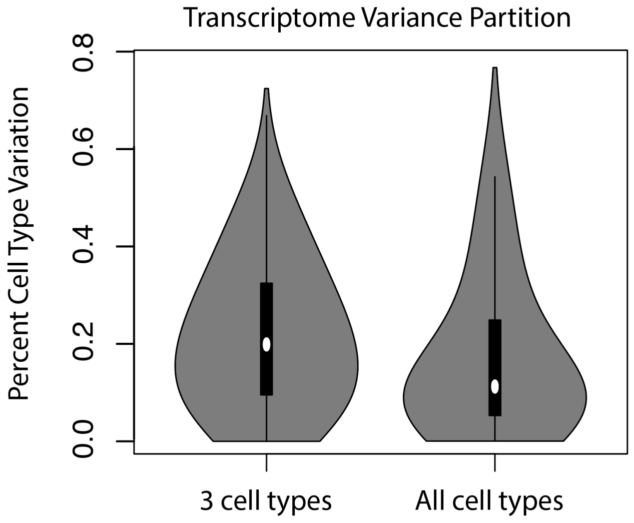


B

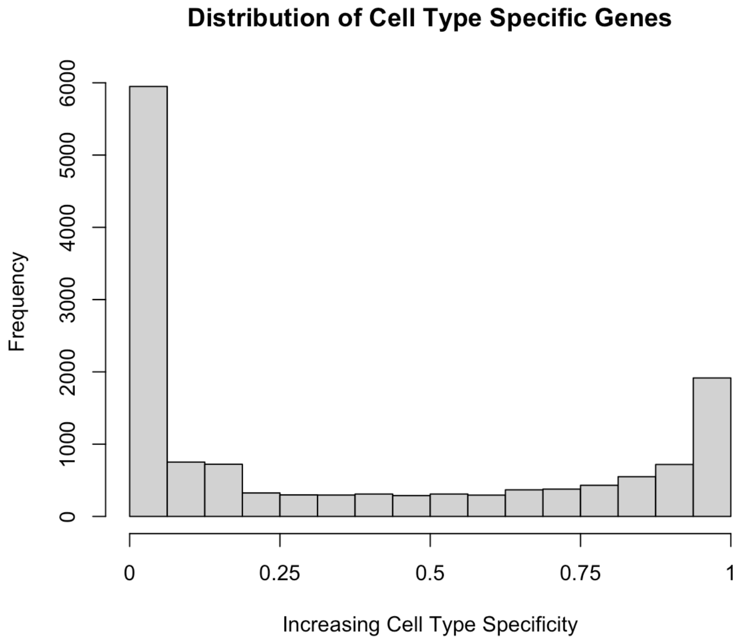


**Fig. S30. Comparing cell-type variation using different numbers of cell types.**

**(A)** Transcriptome variation partition shows increased overall cell type variation when using fewer cell types (three major cell types: Exc, Inh, Non-neuronal) as compared to all cell types. **(B)** Histogram of cell-type-specific genes. Of the 3,216 genes that have the majority of their variation coming from cell type variation, 423 are cell-type-specific (meaning expressed in only one cell type). X-axis represents [1 - fraction of cell types in which a gene is expressed], with a larger number representing a more cell-type-specific gene.

More detail in the supplementary section "***Variance Partition***." This supplementary figure relates to **Fig. 3** and main text section "Measuring transcriptome and epigenome variation across the cohort at the single-cell level."

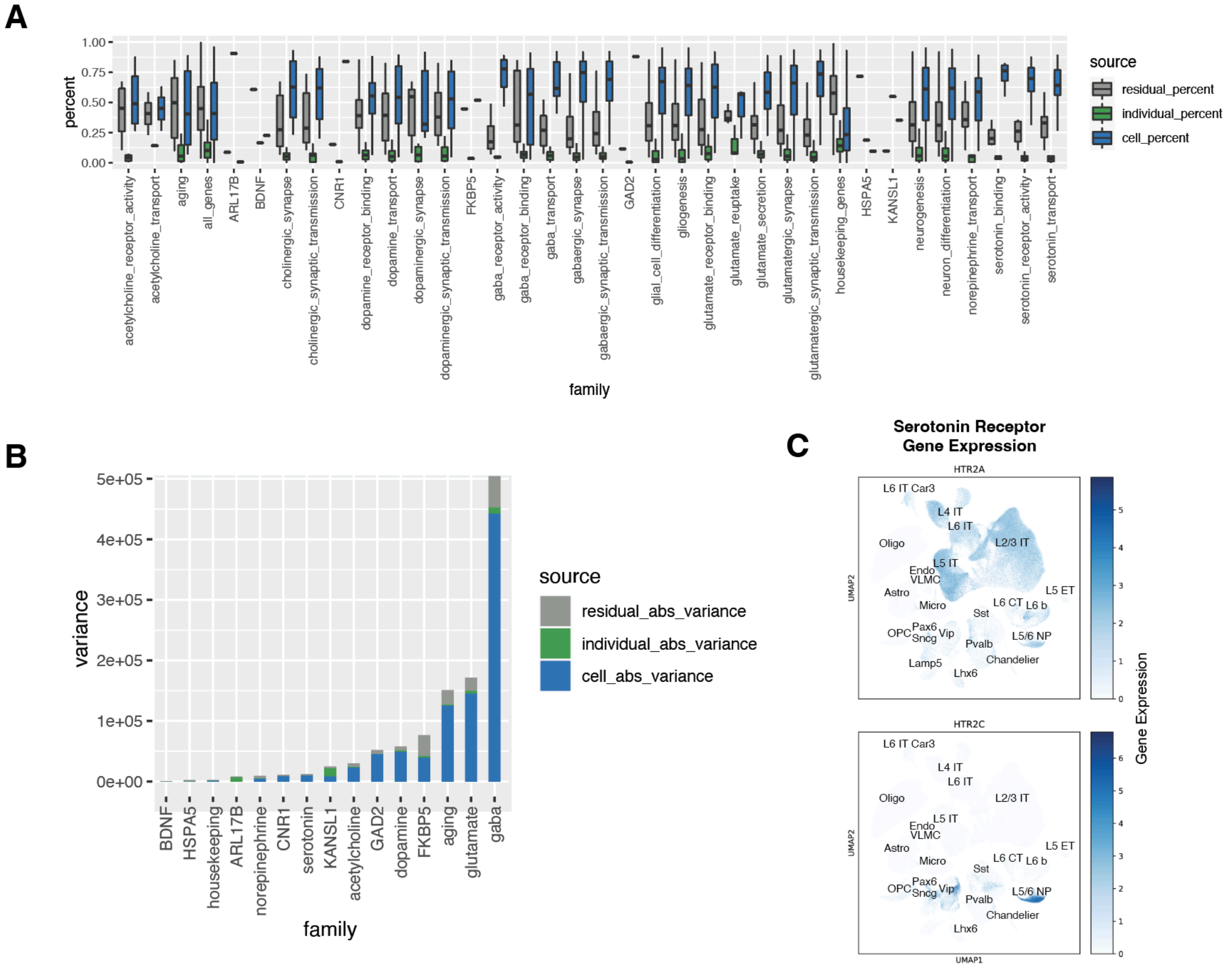


**Fig. S31. Serotonin-related gene variation.**

**(A)** Breakdown of expression variation in gene sets relating to neurotransmitters (grouped by GO annotations) and selected genes highlighted in **Fig. 3C**. **(B)** Total expression variance and breakdown of variance for select genes and gene families/categories highlighted in **Fig. 3C**. **(C)** Serotonin genes *HTR2A* and *HTR2C* demonstrate cell-type variation, and their transcriptomic profiles across cell types are shown in example UMAPs.

More detail in the supplementary section "***Variance Partition***." This supplementary figure relates to **Fig. 3** and main text section "Measuring transcriptome and epigenome variation across the cohort at the single-cell level."

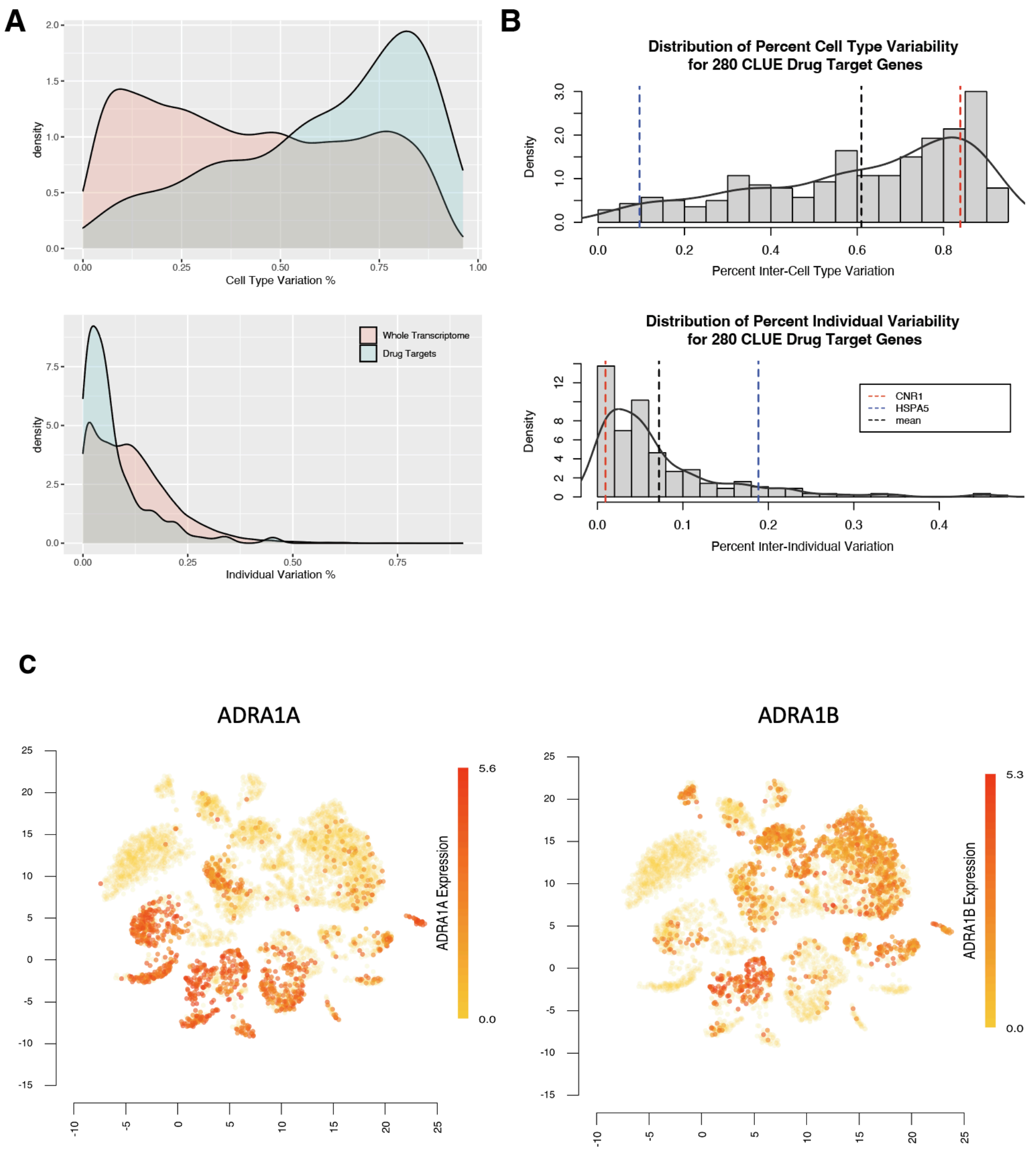


**Fig. S32. Cell-type and individual variability for drug targets.**

**(A)** Comparison of inter-individual and inter-cell-type variation for the whole transcriptome (red) and drug target genes (blue). (**B)** Assessment of inter-individual and inter-cell-type variation for two drug target genes, *CNR1* and *HSPA5*, relative to the distribution of 280 CLUE database drug targets. Dashed red line represents *CNR1*, dashed blue line represents *HSPA5*, and dashed black line represents mean value across 280 drug target genes. **(C)** UMAP of two additional example drug target genes (*ADRA1A* and *ADRA1B*) with high cell-type variation but distinct patterns across which cell types show high expression.

More detail in the supplementary section "***Variance Partition***." This supplementary figure relates to **Fig. 3** and main text section "Measuring transcriptome and epigenome variation across the cohort at the single-cell level."

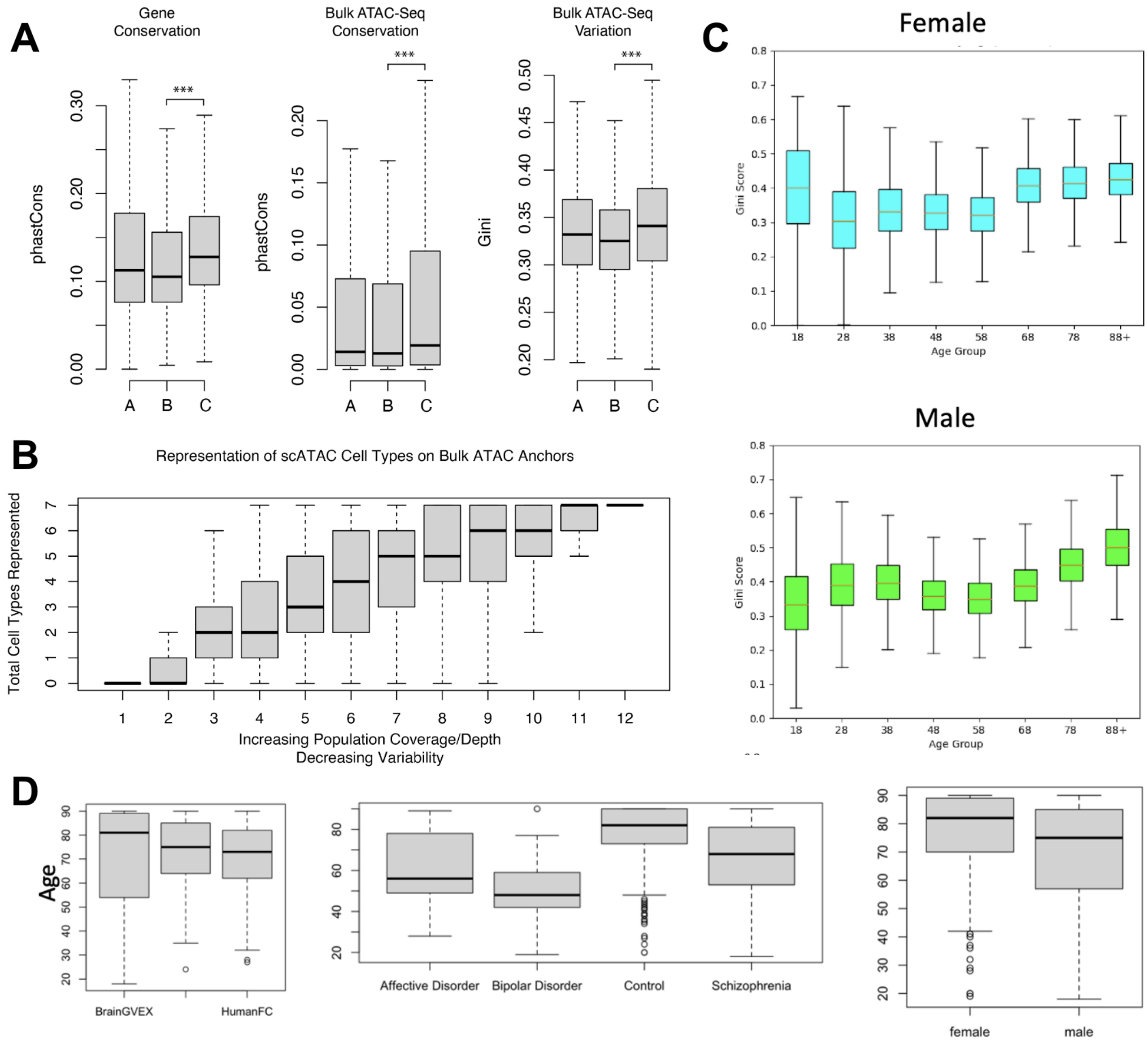


**Fig. S33. Relationships between conservation and coding regions, bulk ATAC-seq peaks, and variability.**

**(A)** Conservation and variation of protein-coding genes and bulk ATAC-seq peaks. A: random sample of genes; B: genes varying by cell type; C: genes varying by individual. **(B)** Number of cell types (cell specificity) versus decreasing population-scale variability for open chromatin regions. **(C)** Gini index of open chromatin regions for male and female samples shows increasing variability during aging. **(D)** Cell-type-specific open chromatin regions from snATAC-seq data show relatively balanced covariates (age, diagnosis, and biological sex).

More detail in the supplementary section "***Conservation***." This supplementary figure relates to **Fig. 3** and main text section "Measuring transcriptome and epigenome variation across the cohort at the single-cell level."

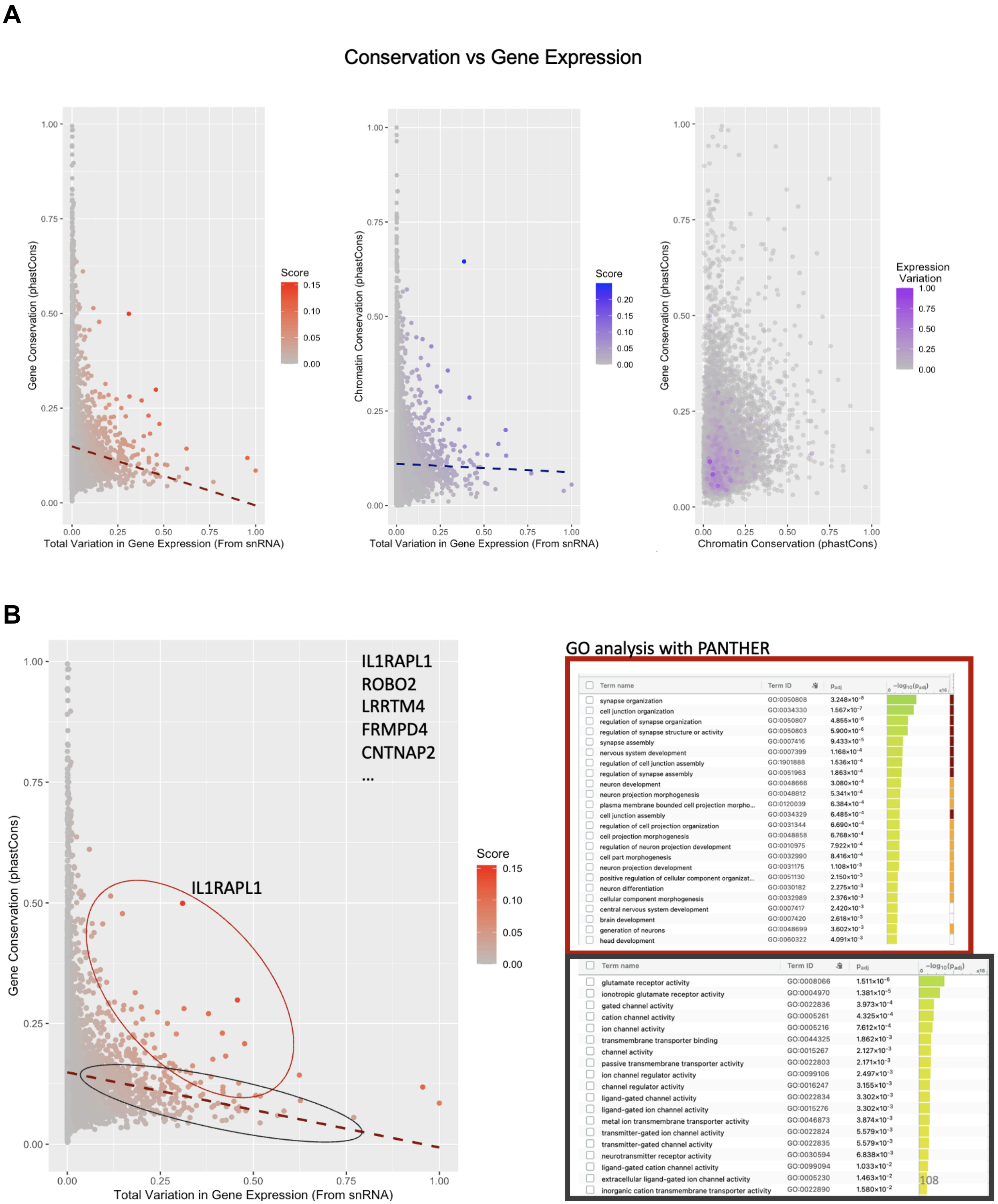


**Fig. S34. Conservation and gene expression patterns.**
**(A)** Scatter plots compare gene conservation with gene expression variation (left), chromatin conservation with gene expression variation (middle), and gene conservation with matched chromatin conservation (right). **(B)** Left scatter plot shows gene conservation versus total variation in gene expression. Genes deviating from expected patterns of variation are colored in

red. Right shows GO analysis of genes within the red and black clusters, demonstrating enrichment in brain functional pathways among genes with abnormal variation patterns.

More detail in the supplementary section "***Conservation*.**" This supplementary figure relates to **Fig. 3** and main text section "Measuring transcriptome and epigenome variation across the cohort at the single-cell level."

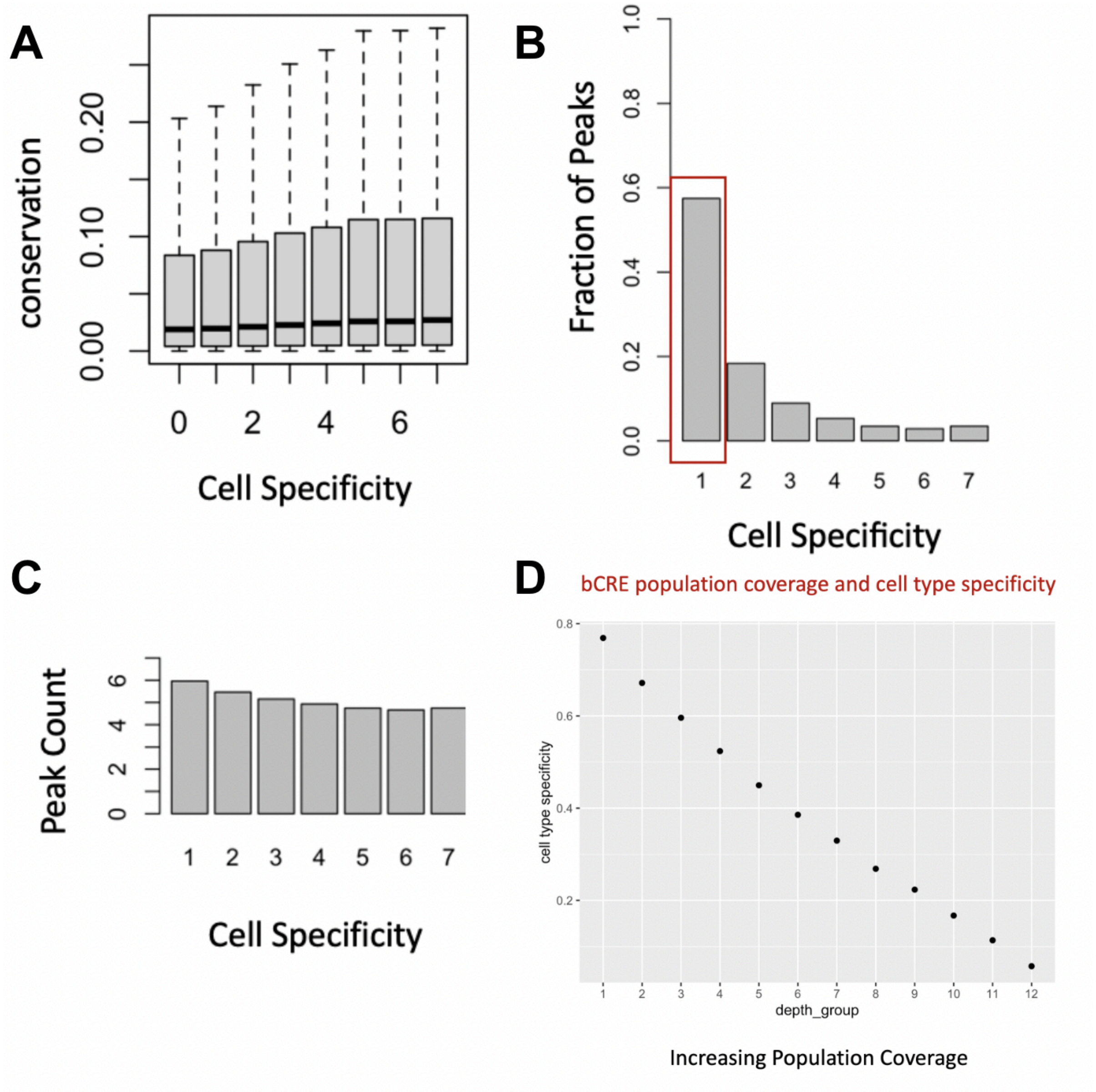


**Fig. S35. Cell-type specificity, variability, and conservation of snATAC-seq peaks.**
**(A)** Conservation versus cell-type specificity of snATAC-Seq peaks. **(B)** Fraction of peaks that correspond to 1 to 7 cell types (ordered by decreasing cell specificity). Red box denotes cell-type-specific peaks unique to a single cell type. **(C)** Number of peaks associated with number of cell types. **(D)** Relationship between cell-specific b-cCREs (defined by overlaying snATAC-seq peaks with b-cCREs) and population coverage (bulk ATAC-seq).

More detail in the supplementary section "***Variance Partition.***" This supplementary figure relates to **Fig. 3** and main text section "Measuring transcriptome and epigenome variation across the cohort at the single-cell level."

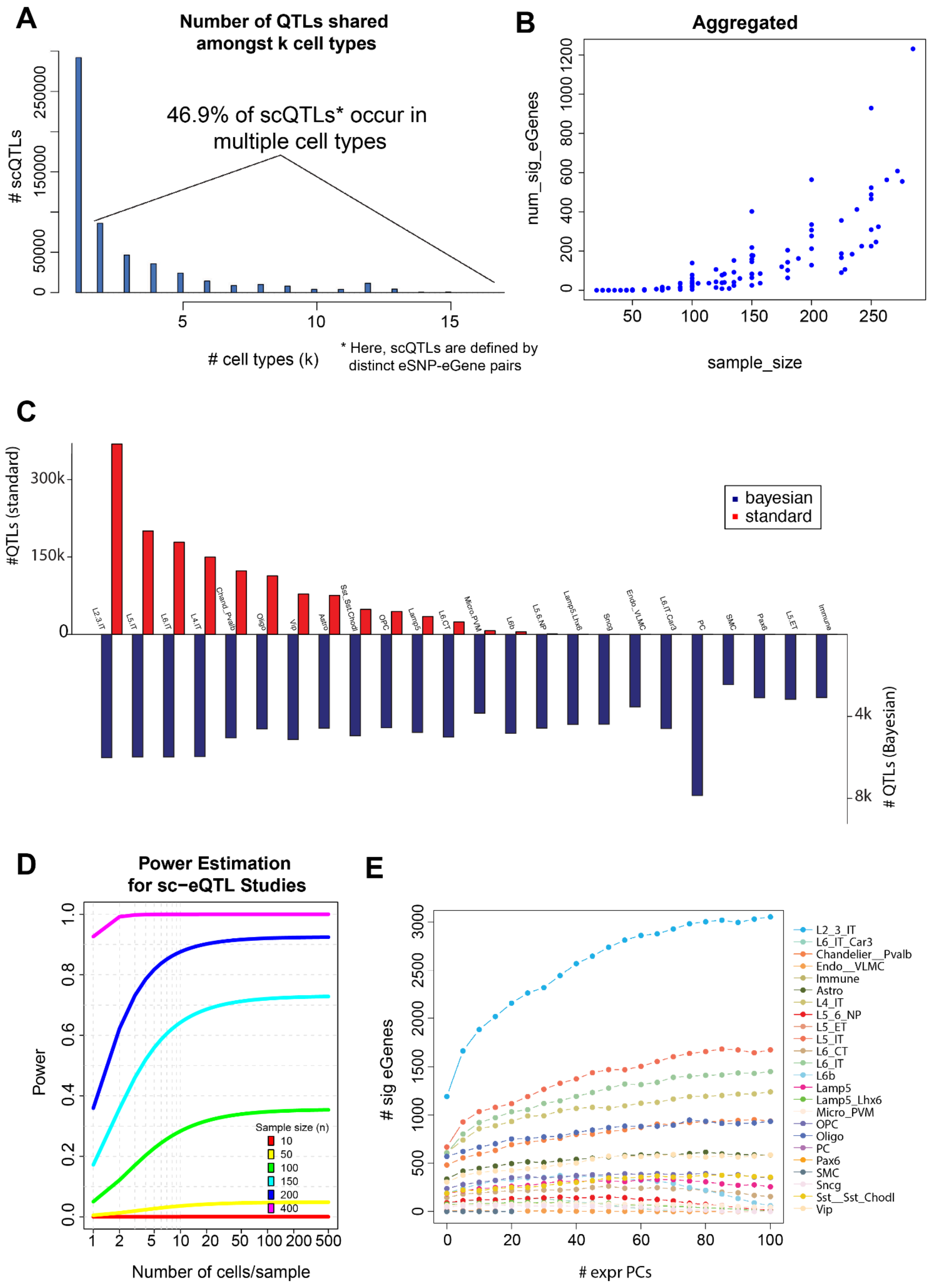
A
Number of QTLs shared amongst k cell types
46.9% of scQTLs* occur in multiple cell types
### scQTLs
### cell types (k)
* Here, scQTLs are defined by distinct eSNP-eGene pairs
B
Aggregated
num_sig_eGenes
sample_size
C
#QTLs (standard)
### QTLs (Bayesian)
bayesian
standard
L2.3.IT
L5.IT
L6.IT
L4.IT
Chand._Pvalb
Oligo
Vip
Astro
Sst_Sst.Chodl
OPC
Lamp5
L6.CT
Micro.PVM
L6b
L5.6.NP
Lamp5.Lhx6
Sncg
Endo._VLMC
L6.IT.Car3
PC
SMC
Pax6
L5.ET
Immune
D
Power Estimation for sc−eQTL Studies
Power
Number of cells/sample
Sample size (n)
E
### sig eGenes
### expr PCs
L2_3_IT
L6_IT_Car3
Chandelier__Pvalb
Endo__VLMC
Immune
Astro
L4_IT
L5_6_NP
L5_ET
L5_IT
L6_CT
L6_IT
L6b
Lamp5
Lamp5_Lhx6
Micro_PVM
OPC
Oligo
PC
Pax6
SMC
Sncg
Sst__Sst_Chodl
Vip

**Fig. S36. Number of eQTLs among cell types and sc-eQTL power estimation.**

**(A)** Distribution of the number of scQTLs that are shared among exactly k cell types (with k in the range of 1 to 19 cell types, inclusive). **(B)** Aggregation of power analysis results. For each cell type, random subsets were selected of varying sizes, and the number of eGenes was recorded (see supplementary section 4 for details on significance testing). The results for all subsets in all cell types are shown. **(C)** Counts of identified eGenes from the Bayesian-based scheme for QTL detection, compared with counts from the primary scQTL analysis. **(D)** Power calculation estimates for scQTL studies. The statistical power for finding a significant scQTL ($p<0.05$, one-way unbalanced ANOVA) at 0.5 error rate increases to >0.8 for n=200 samples at >50 cells/sample, as calculated using the powerEQTL software. **(E)** Number of eGenes identified as a function of the number of expression PCs.

More detail in the supplementary section "***scQTLs.***" This supplementary figure relates to **Fig. 4** and main text section "Determining cell-type-specific eQTLs from single-cell data."

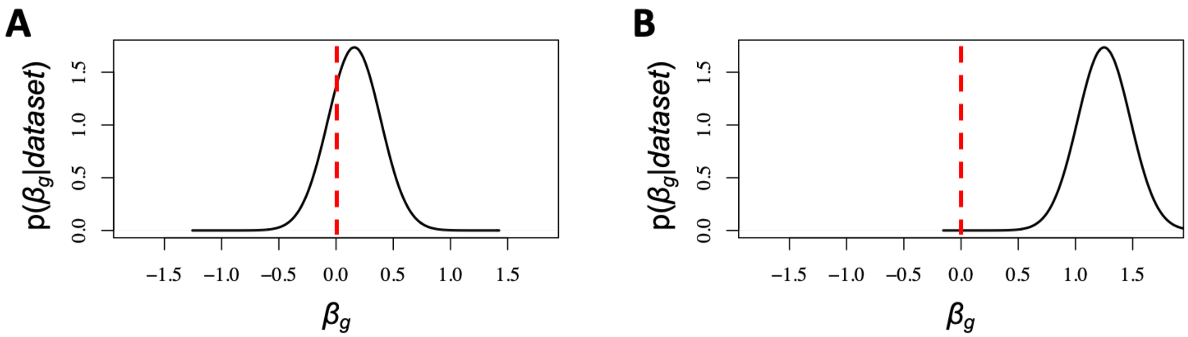


**Fig. S37. scQTL identification using Bayesian linear mixed effects models.**
**(A)** The posterior distribution p($\beta_g$|*dataset*) of the effect size is such that the value $\beta_g$ = 0 lies within a high-density region of the distribution. Such a posterior distribution suggests the absence of a strong scQTL. **(B)** The posterior distribution p($\beta_g$|*dataset*) of the effect size is such that the value $\beta_g$ = 0 lies within a very low-density region of the distribution (very far away from $\beta_g$ = 0). Such a posterior distribution more likely suggests a strong scQTL.

More detail in the supplementary section "***Bayesian scQTLs.***" This supplementary figure relates to **Fig. 4** and main text section "Determining cell-type-specific eQTLs from single-cell data."

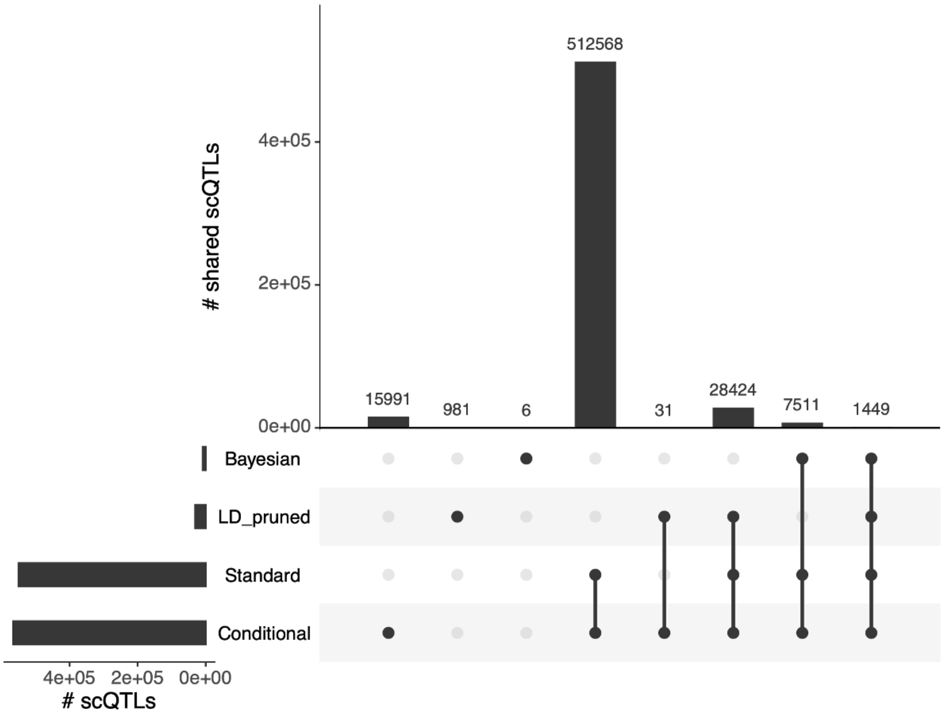


**Fig. S38. Comparisons across different methods for scQTL calling (standard linear regression, Bayesian linear mixed effects models, conditional calling, and linear regression with LD-pruned variants).**

The UpSet plot represents the extent to which scQTLs (aggregated across all cell types) are identified by four different calling methods. In this analysis, a given scQTL is defined as a distinct eSNP-eGene pair.

More detail in the supplementary section "***scQTLs.***" This supplementary figure relates to **Fig. 4** and main text section "Determining cell-type-specific eQTLs from single-cell data."

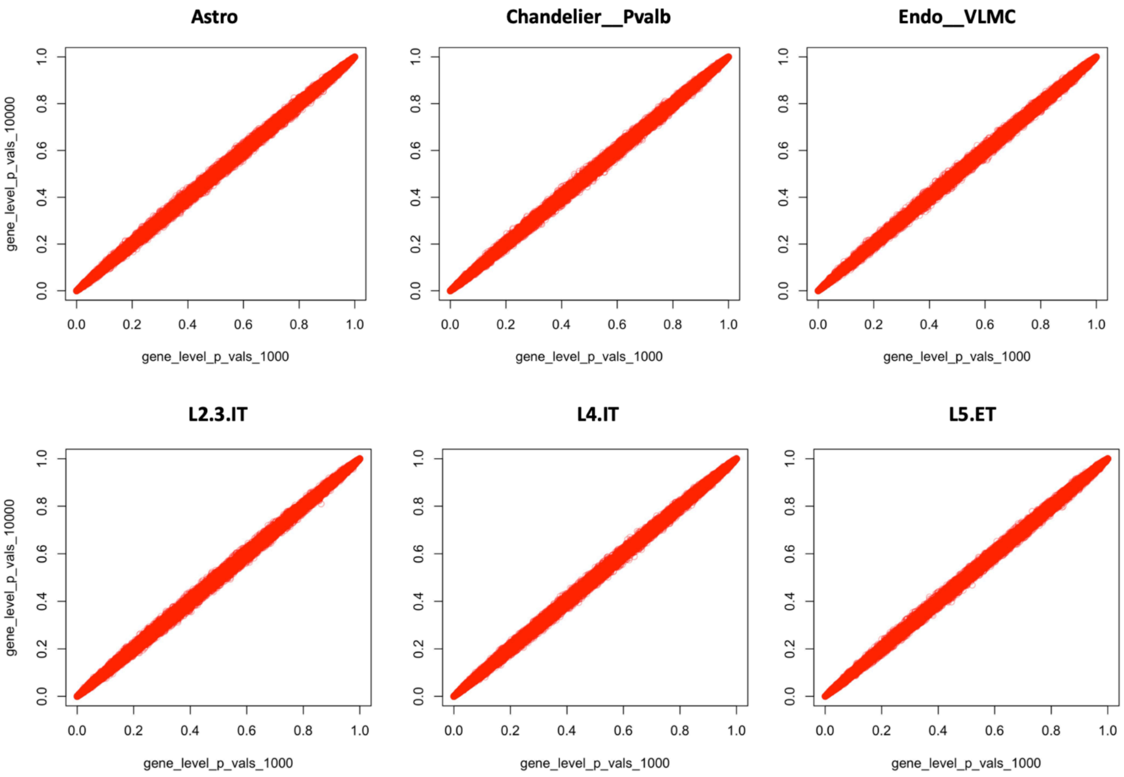


**Fig. S39. Comparison of gene-level p-values from different numbers of permutations.** Shown are the gene-level p-values from 1,000 permutations compared to those derived from 10,000 permutations when calculating significant eGenes associated with scQTLs.

More detail in the supplementary section "***scQTLs.***" This supplementary figure relates to **Fig. 4** and main text section "Determining cell-type-specific eQTLs from single-cell data."

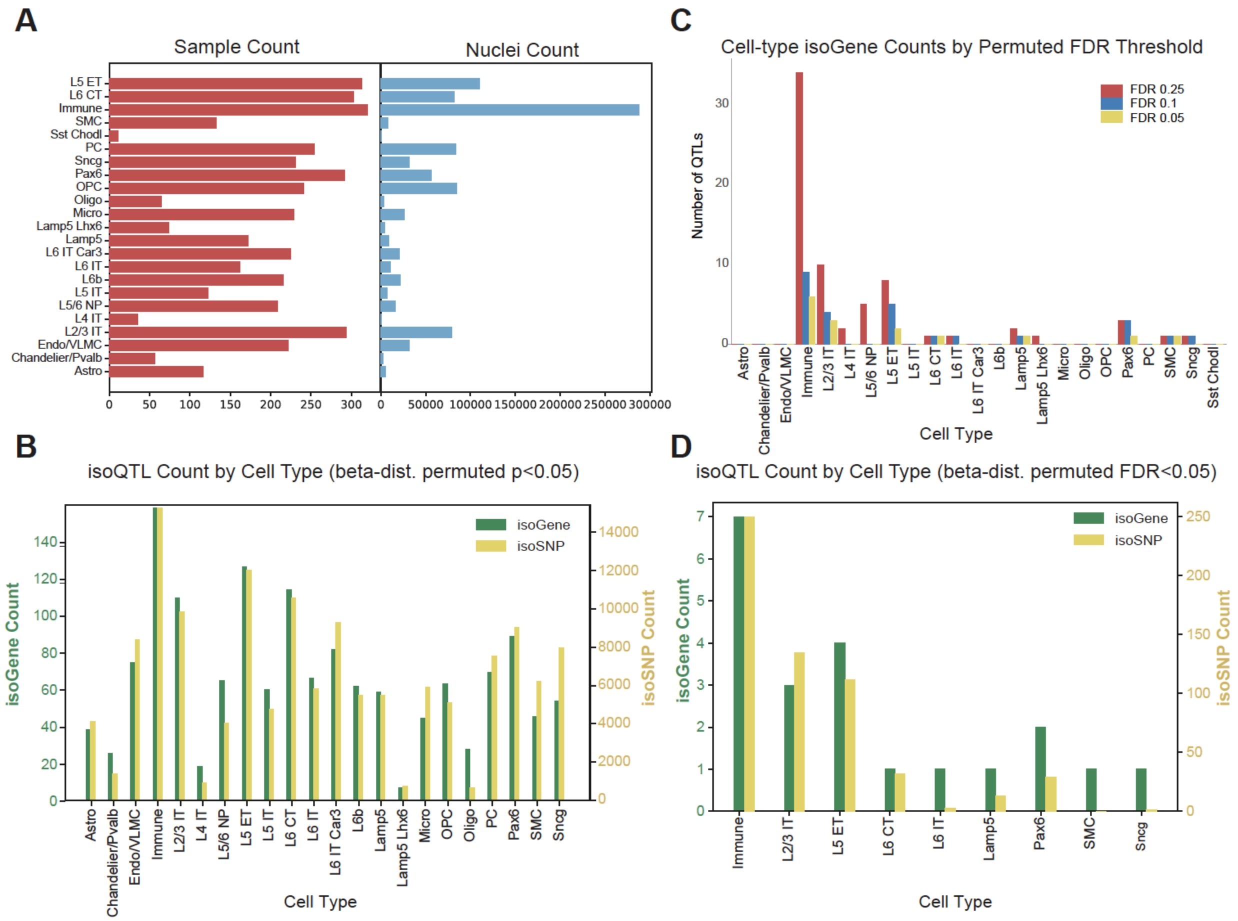


**Fig. S40. IsoQTL calculation.**

**(A)** Bar plots show the distribution of samples and nuclei assessed for isoQTLs after QC filtering. **(B)** Bar plots show the distribution of nominally significant (beta distribution permuted p<0.05 without FDR correction) isoGenes (green) and isoSNPs (pink) per cell type. **(C)** Bar plots show the counts of significant isoQTLs in each cell type with nominal beta distribution permuted p<0.05 at different FDR thresholds. **(D)** Bar plots show the distribution of significant (beta distribution permuted FDR<0.05) isoGenes (green) and isoSNPs (pink) per cell type.

More detail in the supplementary section "***Isoform QTLs.***" This supplementary figure relates to **Fig. 4** and the main text section "Determining cell-type-specific eQTLs from single-cell data."

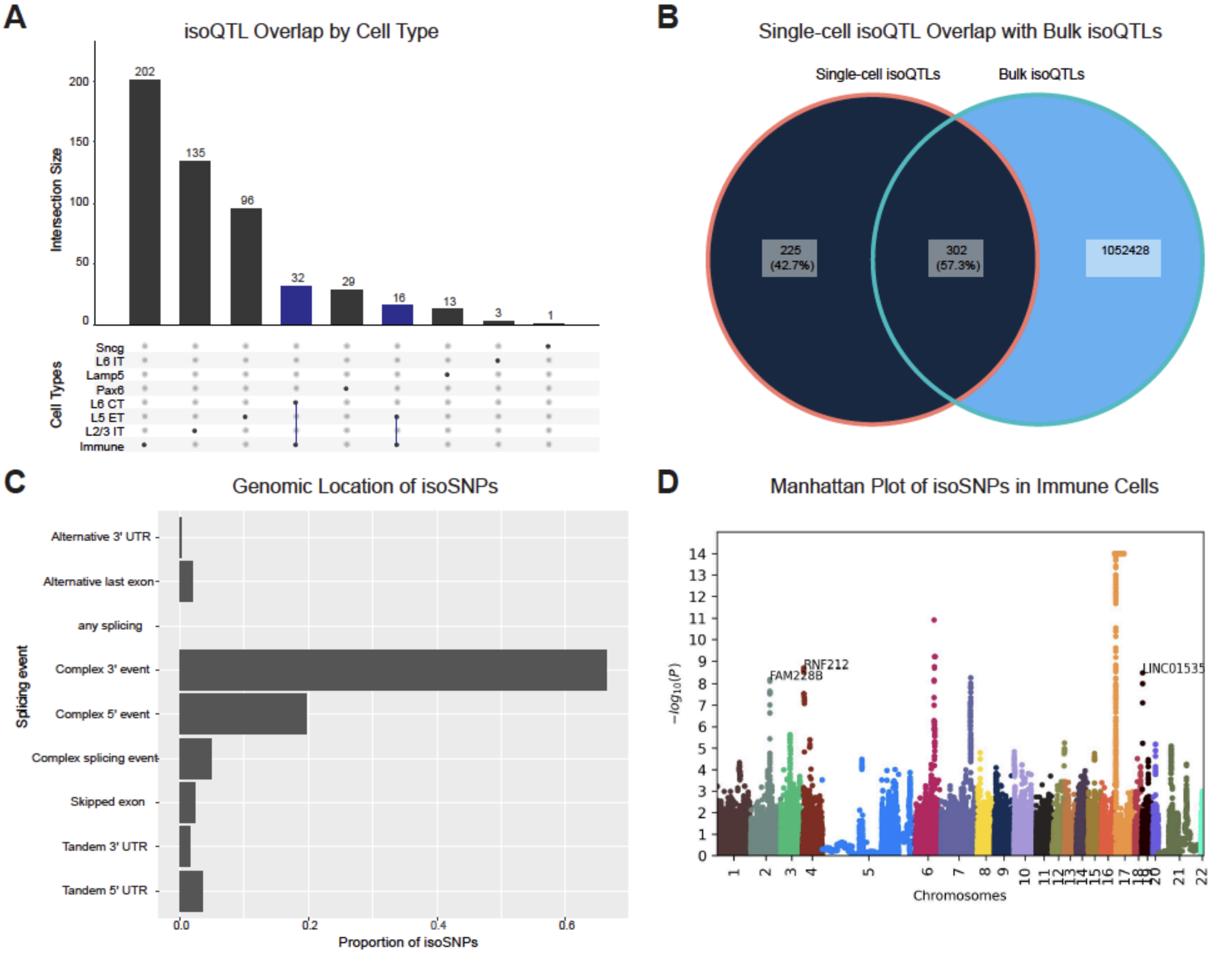


**Fig. S41. IsoQTL characterization.**

**(A)** UpSet plot shows the overlap of significant isoQTLs that pass multiple-testing correction (FDR<0.05, beta-distribution permuted p-values) between cell types. Blue bars represent isoSNPs that are observed in multiple cell types. **(B)** Venn diagram shows the overlap of all significant isoQTLs with isoQTLs identified from bulk RNA-seq in the adult brain(*4*). **(C)** Bar plots show the proportion of significant isoQTLs by genomic location. **(D)** Manhattan plot shows identified isoQTLs within immune cells. isoSNPs for three significant isoGenes (FDR<0.05) are highlighted.

More detail in the supplementary section "***Isoform QTLs.***" This supplementary figure relates to **Fig. 4** and main text section "Determining cell-type-specific eQTLs from single-cell data."

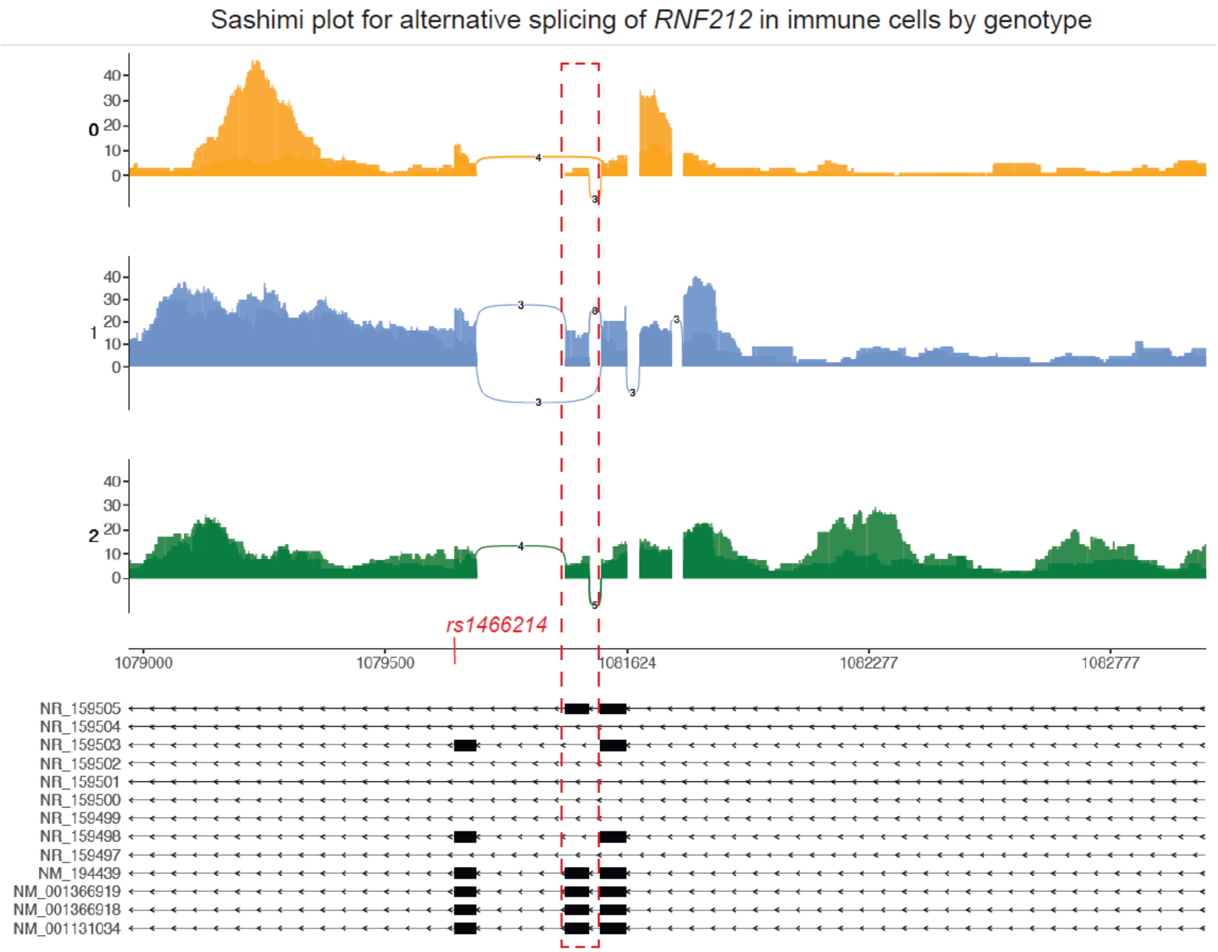


**Fig. S42. Example of alternative isoform usage due to an isoQTL.**

Sashimi plot shows alternative splicing of *RNF212* in immune cells due to the isoSNP rs4166214. Red box highlights an *RNF212* isoform with increased expression among individuals with the alternate allele (blue, heterozygous; green, homozygous alternate) vs. those homozygous for the reference allele (orange). Sashimi plot was generated using the ggsashimi package (*183*).

More detail in the supplementary section "***Isoform QTLs.***" This supplementary figure relates to **Fig. 4** and main text section "Determining cell-type-specific eQTLs from single-cell data."

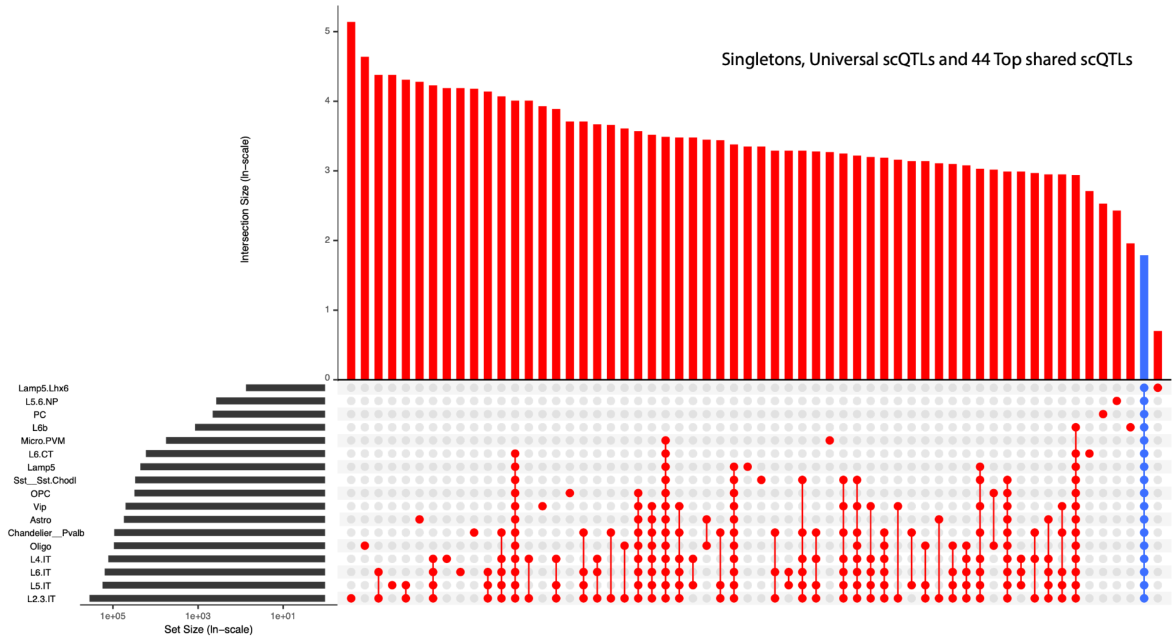


**Fig. S43. eQTL UpSet plot.**

UpSet plot for core scQTLs showing singleton scQTLs (scQTLs unique to a given cell type), universal scQTLs (core scQTLs in all 17 cell types in which scQTLs occur, highlighted in blue), and the top 44 combinations of cell types with the largest numbers of shared scQTLs.

More detail in the supplementary section "***scQTLs.***" This supplementary figure relates to **Fig. 4** and main text section "Determining cell-type-specific eQTLs from single-cell data."

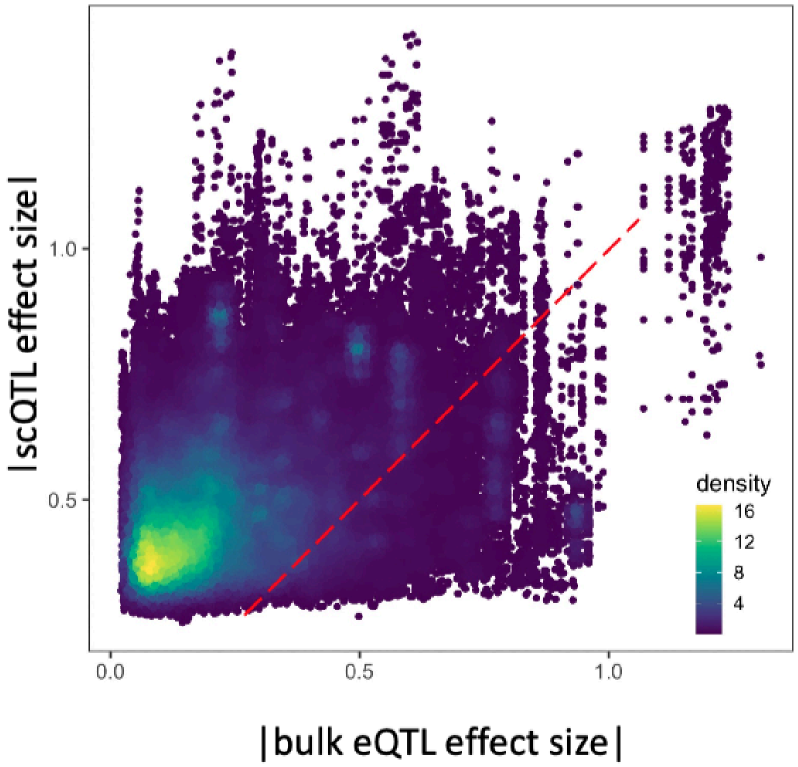


**Fig. S44. Comparison of scQTL and bulk eQTL.**

Density map of the scQTL effect sizes (y-axis) against matched bulk cis-eQTL effect sizes (x-axis). The dashed red line denotes the plot's diagonal (ie, "x = y") line.

More detail in the supplementary section "***scQTLs.***" This supplementary figure relates to **Fig. 4** and main text section "Determining cell-type-specific eQTLs from single-cell data."

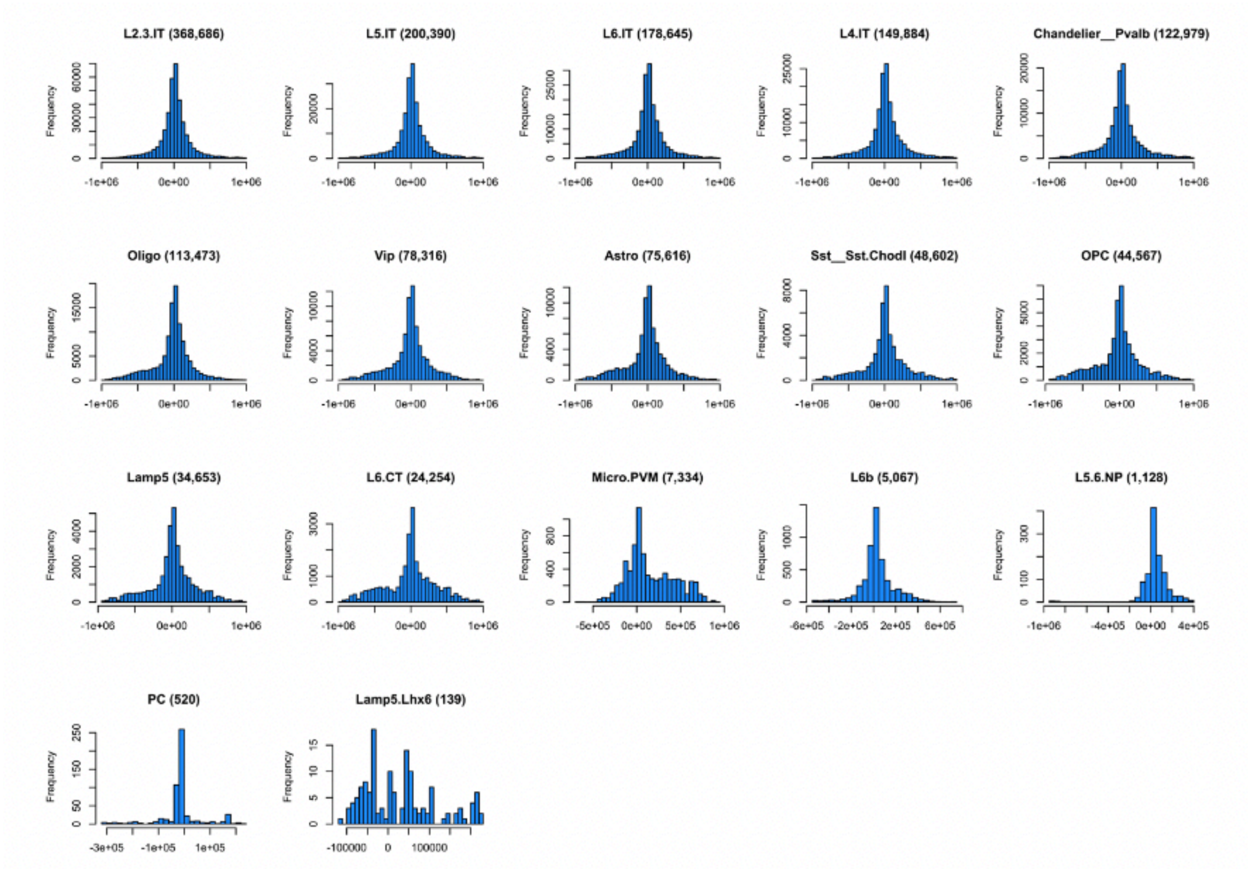


**Fig. S45. Distance to TSS.**

Distance between eGenes and their respective eSNPs for each individual cell type (scQTLs FDR < 0.05).

More detail in the supplementary section "***scQTLs.***" This supplementary figure relates to **Fig. 4** and main text section "Determining cell-type-specific eQTLs from single-cell data."

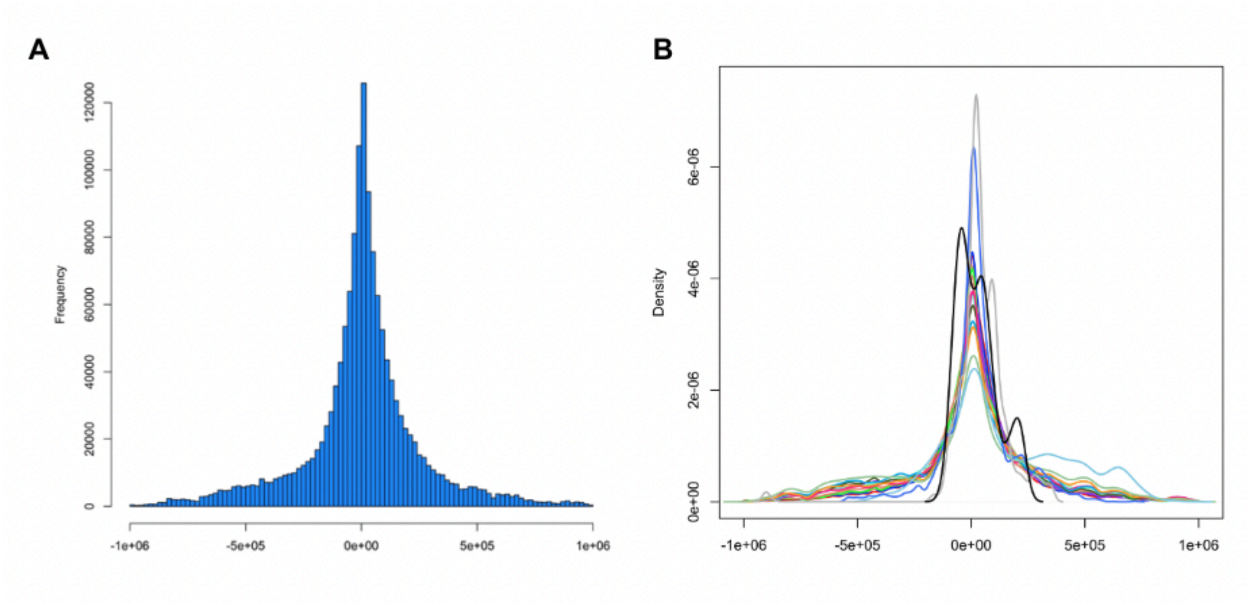


**Fig. S46. Aggregated distances between eGenes and eSNPs.**
**(A)** Aggregated distances (among 17 cell types with scQTLs) between eGenes and their respective eSNPs for each individual cell type (scQTL FDR < 0.05). **(B)** Smoothed distribution profiles of the distances between eGenes and their respective eSNPs for scQTLs in all individual cell types. (The cell type PC is omitted from this graph.)

More detail in the supplementary section "***scQTLs***." This supplementary figure relates to **Fig. 4** and main text section "Determining cell-type-specific eQTLs from single-cell data."

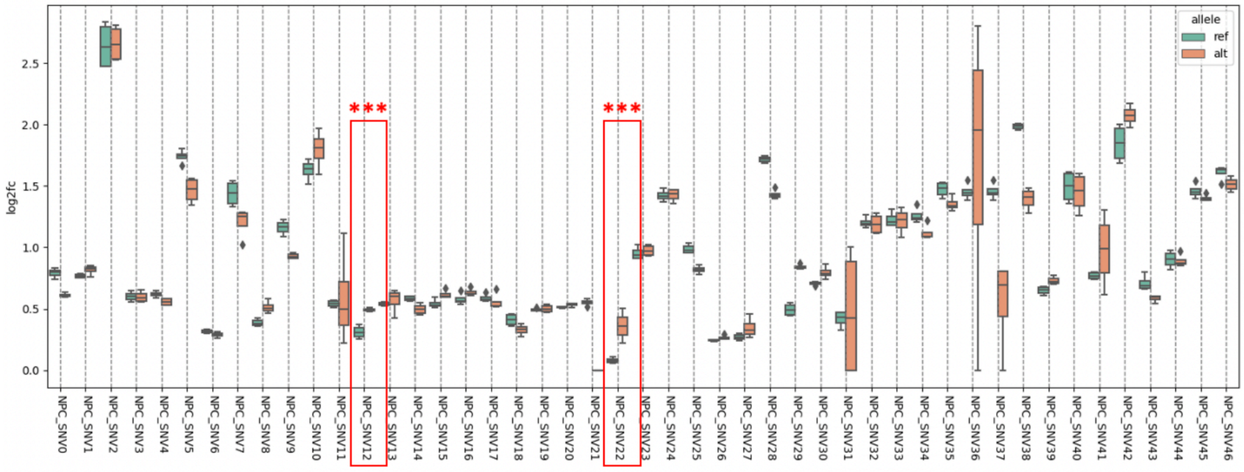


NPC_SNV12: chr17:45894571:C:A, p=0.0046
NPC_SNV22: chr17:45490914:G:A, p=0.00011

**Fig. S47. Quantifying the modulation of enhancer activity by mutSTARR-seq.**

Box plots show the $log_2fc$ STARR-seq signal change before and after mutation of target SNPs. Highlighted in red are two key significant SNPs based on scQTLs and validated enhancers tested, both demonstrating increasing STARR-seq assay signal after mutation. Asterisks represent $p<0.005$ (Chi-squared test). We note that the mut-STARR-seq experiments were done in NPC cells.

More detail in the supplementary section "***mut-STARR-Seq.***" This supplementary figure relates to **Fig. 4** and main text section "Determining cell-type-specific eQTLs from single-cell data."

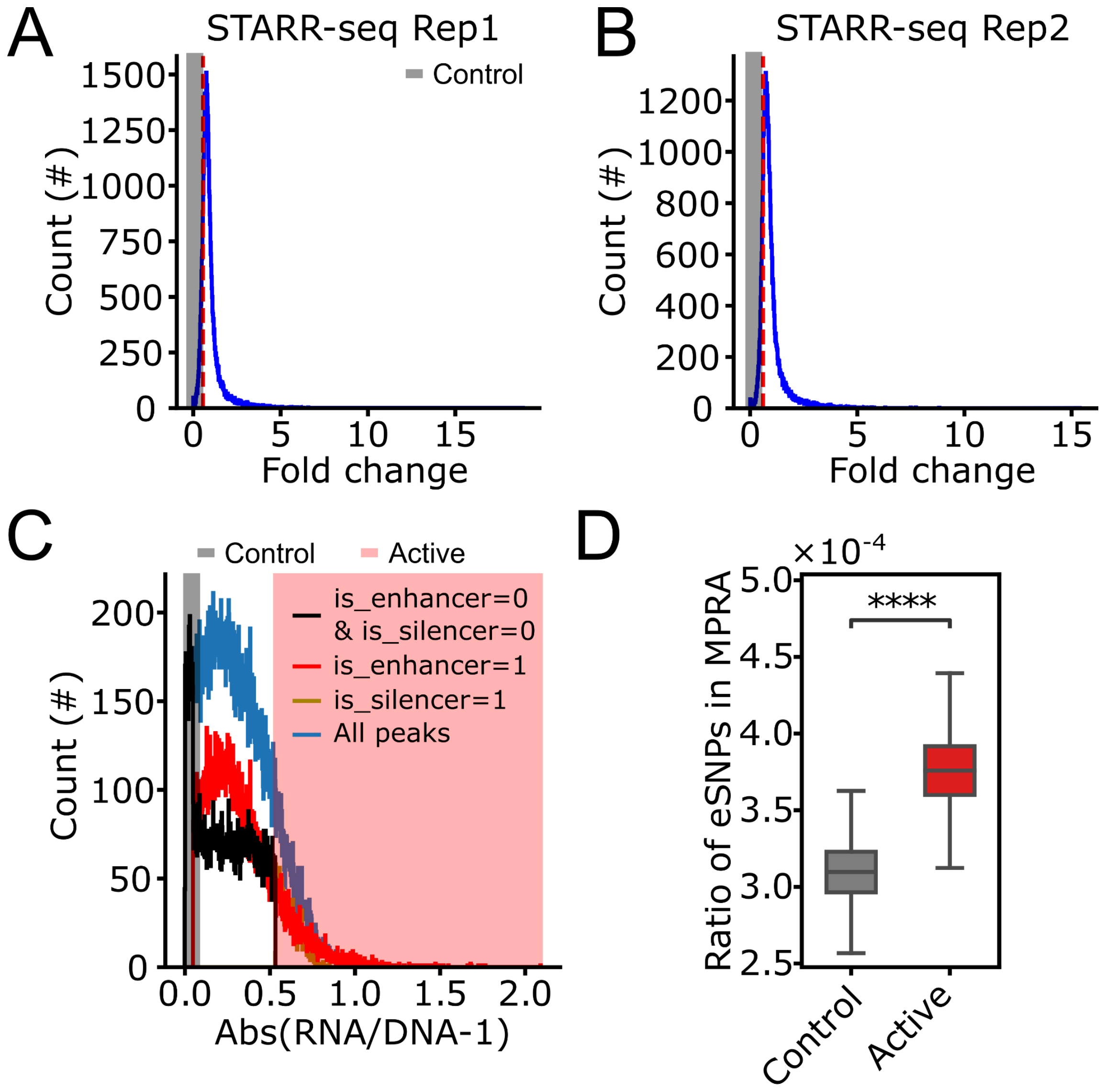


**Fig. S48. Validation of scQTLs using STARR-seq and MPRA.**

**(A-B)** Fold-change signal distribution of the STARR-seq candidate regions across two replicates; gray regions denote the selected candidate regions serving as the control set. **(C)** Distribution of the absolute values of the ratio between RNA and DNA-1 in MPRA experiments. Red regions denote the selected candidate regions serving as the active enhancer set, while gray regions denote the candidate regions serving as the control set. Distributions are shown for all peaks, defined enhancers, defined silencers, and non-enhancer/silencer peaks. **(D)** Comparison of the ratio of eSNPs in active (red) or control (gray) regions as defined by MPRA experiments. **** indicates p-value $<1.0\times10^{-4}$, Mann-Whitney Wilcoxon test.

More detail in the supplementary section "***STARR-seq and MPRA Validation.***" This supplementary figure relates to **Fig. 4** and main text section "Determining cell-type-specific eQTLs from single-cell data."

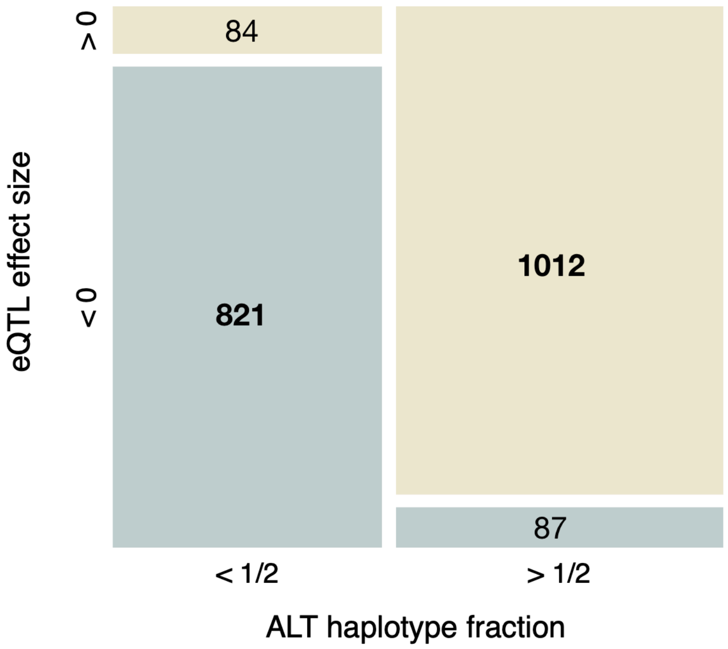


**Fig. S49. Matching concordance between allele-specific expression data and scQTLs.**
Plot showing numbers of concordant (eQTL effect size < 0 and ALT haplotype fraction < 0.5, or eQTL effect size > 0 and ALT haplotype fraction > 0.5) and discordant (eQTL effect size < 0 and ALT haplotype fraction > 0.5, or eQTL effect size > 0 and ALT haplotype fraction < 0.5) eQTL-ASE eGene pairs found in the MultiomeBrain cohort samples (see **Fig. 4F**).

More detail in the supplementary section "***Allele-specific expression.***" This supplementary figure relates to **Fig. 4** and main text section "Determining cell-type-specific eQTLs from single-cell data."

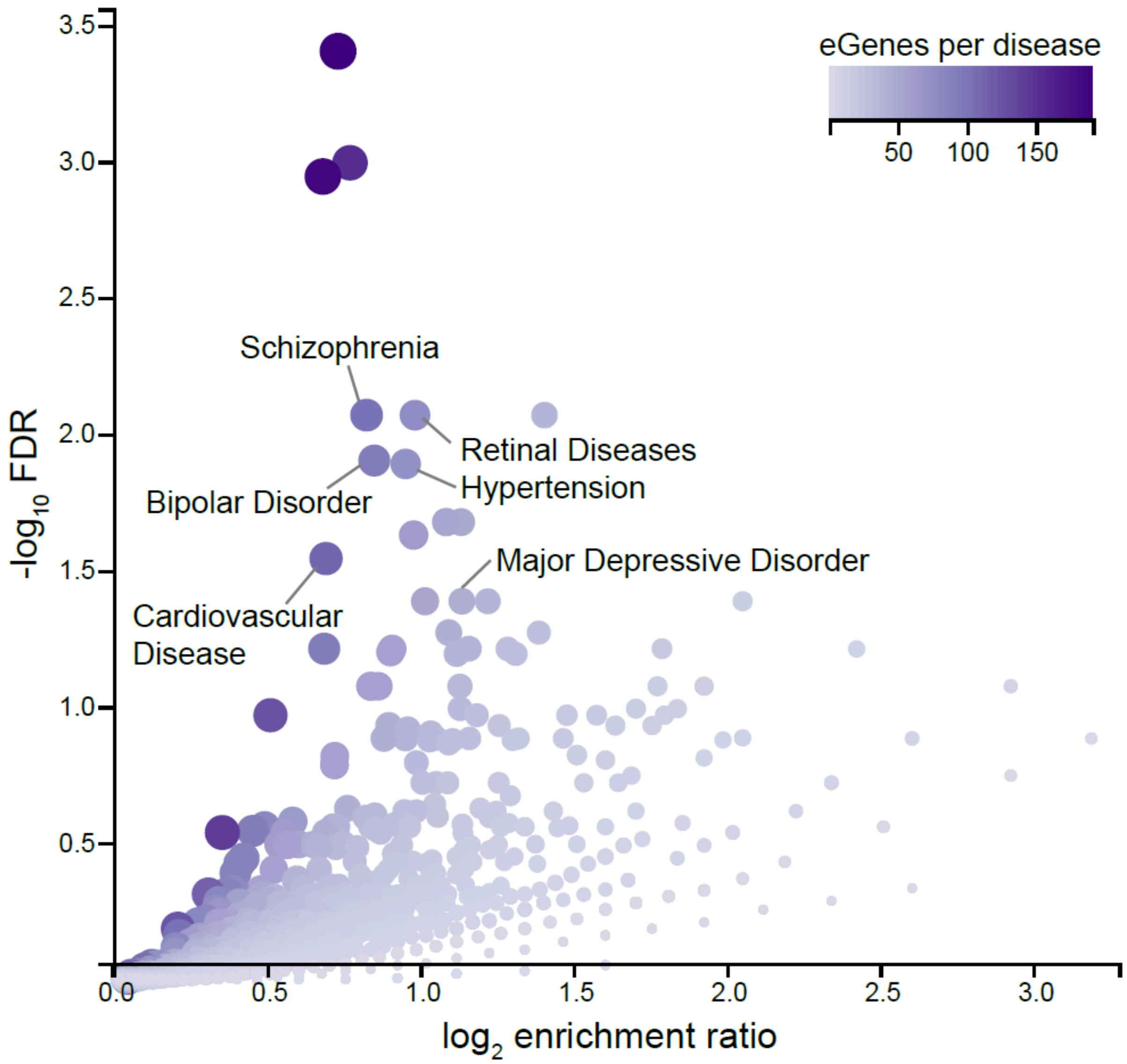


**Fig. S50. Enrichment of eGenes in scQTLs for disease-associated gene sets.**
Volcano plot shows the enrichment of eGenes found across all cell types in the primary scQTL dataset for disease trait annotations. Plot was generated using the WebGestalt gene set enrichment toolkit (*230*) for genes annotated for disease association in the GLAD4U database (*231*). Labeled diseases on the plot show select disorders with significant (FDR<0.05, Fisher's exact test) enrichment among the eGenes. Circle color indicates the number of eGenes annotated for each disease category.

Related to the supplementary section "***scQTLs***." This supplementary figure relates to **Fig. 4** and main text section "Determining cell-type-specific eQTLs from single-cell data."

**A**

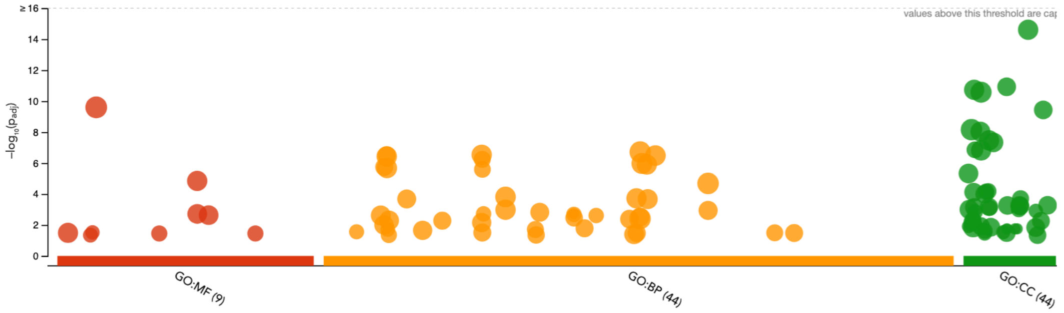


**B**

| GO:BP | | stats | | |
|---|---|---|---|---|
| Term name | Term ID | $p_{adj}$ | $-\log_{10}(p_{adj})$ 0 … ≤16 | Show evidence codes |
| regulation of cellular process | GO:0050794 | $2.417\times10^{-7}$ | | |
| signaling | GO:0023052 | $3.347\times10^{-7}$ | | |
| cell adhesion | GO:0007155 | $3.363\times10^{-7}$ | | |
| cellular response to stimulus | GO:0051716 | $3.850\times10^{-7}$ | | |
| cell communication | GO:0007154 | $4.323\times10^{-7}$ | | |
| actin filament-based process | GO:0030029 | $6.078\times10^{-7}$ | | |
| response to stimulus | GO:0050896 | $1.257\times10^{-6}$ | | |
| localization | GO:0051179 | $1.413\times10^{-6}$ | | |
| cytoskeleton organization | GO:0007010 | $1.905\times10^{-6}$ | | |
| signal transduction | GO:0007165 | $2.407\times10^{-6}$ | | |
| actin cytoskeleton organization | GO:0030036 | $2.671\times10^{-6}$ | | |
| biological regulation | GO:0065007 | $1.812\times10^{-5}$ | | |
| multicellular organismal process | GO:0032501 | $1.755\times10^{-4}$ | | |
| anatomical structure development | GO:0048856 | $2.093\times10^{-4}$ | | |
| anatomical structure morphogenesis | GO:0009653 | $2.223\times10^{-4}$ | | |
| establishment of localization | GO:0051234 | $2.323\times10^{-4}$ | | |
| regulation of molecular function | GO:0065009 | $8.252\times10^{-4}$ | | |
| developmental process | GO:0032502 | $1.161\times10^{-3}$ | | |
| intracellular signal transduction | GO:0035556 | $1.595\times10^{-3}$ | | |
| DNA methylation involved in gamete generation | GO:0043046 | $1.814\times10^{-3}$ | | |
| extracellular matrix organization | GO:0030198 | $1.946\times10^{-3}$ | | |
| positive regulation of catalytic activity | GO:0043085 | $2.043\times10^{-3}$ | | |
| extracellular structure organization | GO:0043062 | $2.115\times10^{-3}$ | | |
| regulation of catalytic activity | GO:0050790 | $2.137\times10^{-3}$ | | |
| external encapsulating structure organization | GO:0045229 | $2.495\times10^{-3}$ | | |
| transport | GO:0006810 | $2.646\times10^{-3}$ | | |
| cell development | GO:0048468 | $4.385\times10^{-3}$ | | |
| regulation of biological process | GO:0050789 | $4.637\times10^{-3}$ | | |
| vesicle-mediated transport | GO:0016192 | $5.534\times10^{-3}$ | | |
| multicellular organism development | GO:0007275 | $5.550\times10^{-3}$ | | |
| regulation of signaling | GO:0023051 | $7.454\times10^{-3}$ | | |
| positive regulation of molecular function | GO:0044093 | $1.062\times10^{-2}$ | | |
| organelle organization | GO:0006996 | $1.067\times10^{-2}$ | | |
| positive regulation of cytosolic calcium ion concentration | GO:0007204 | $1.974\times10^{-2}$ | | |
| tube morphogenesis | GO:0035239 | $2.004\times10^{-2}$ | | |
| regulation of cell communication | GO:0010646 | $2.347\times10^{-2}$ | | |
| regulation of GTPase activity | GO:0043087 | $2.605\times10^{-2}$ | | |
| vascular process in circulatory system | GO:0003018 | $2.794\times10^{-2}$ | | |
| cell projection organization | GO:0030030 | $3.213\times10^{-2}$ | | |
| supramolecular fiber organization | GO:0097435 | $3.282\times10^{-2}$ | | |
| cell motility | GO:0048870 | $3.459\times10^{-2}$ | | |
| plasma membrane bounded cell projection organization | GO:0120036 | $3.465\times10^{-2}$ | | |
| system development | GO:0048731 | $4.273\times10^{-2}$ | | |
| small GTPase mediated signal transduction | GO:0007264 | $4.514\times10^{-2}$ | | |
| tube development | GO:0035295 | $4.553\times10^{-2}$ | | |

## C

| GO:MF | | stats | |
|---|---|---|---|
| Term name | Term ID | $p_{adj}$ | $-\log_{10}(p_{adj})$ |
| protein binding | GO:0005515 | $3.784\times10^{-10}$ | |
| ion binding | GO:0043167 | $1.600\times10^{-5}$ | |
| cation binding | GO:0043169 | $2.083\times10^{-3}$ | |
| metal ion binding | GO:0046872 | $2.471\times10^{-3}$ | |
| GTPase regulator activity | GO:0030695 | $1.877\times10^{-2}$ | |
| nucleoside-triphosphatase regulator activity | GO:0060589 | $1.877\times10^{-2}$ | |
| extracellular matrix structural constituent | GO:0005201 | $3.046\times10^{-2}$ | |
| catalytic activity | GO:0003824 | $3.461\times10^{-2}$ | |
| guanyl-nucleotide exchange factor activity | GO:0005085 | $4.610\times10^{-2}$ | |

Show evidence codes

## D

| GO:CC | | stats | |
|---|---|---|---|
| Term name | Term ID | $p_{adj}$ | $-\log_{10}(p_{adj})$ |
| cell periphery | GO:0071944 | $1.733\times10^{-15}$ | |
| cell projection | GO:0042995 | $1.294\times10^{-11}$ | |
| plasma membrane | GO:0005886 | $1.348\times10^{-11}$ | |
| membrane | GO:0016020 | $2.155\times10^{-11}$ | |
| plasma membrane bounded cell projection | GO:0120025 | $4.002\times10^{-10}$ | |
| cytoplasm | GO:0005737 | $8.845\times10^{-9}$ | |
| endomembrane system | GO:0012505 | $1.042\times10^{-8}$ | |
| intrinsic component of membrane | GO:0031224 | $3.859\times10^{-8}$ | |
| vesicle | GO:0031982 | $5.096\times10^{-8}$ | |
| cilium | GO:0005929 | $1.446\times10^{-7}$ | |
| integral component of membrane | GO:0016021 | $1.740\times10^{-7}$ | |
| extracellular region | GO:0005576 | $5.150\times10^{-6}$ | |
| extracellular matrix | GO:0031012 | $6.844\times10^{-5}$ | |
| external encapsulating structure | GO:0030312 | $7.303\times10^{-5}$ | |
| cytoskeleton | GO:0005856 | $8.038\times10^{-5}$ | |
| cell junction | GO:0030054 | $1.071\times10^{-4}$ | |
| anchoring junction | GO:0070161 | $2.073\times10^{-4}$ | |
| extracellular vesicle | GO:1903561 | $5.573\times10^{-4}$ | |
| extracellular membrane-bounded organelle | GO:0065010 | $5.741\times10^{-4}$ | |
| extracellular organelle | GO:0043230 | $5.741\times10^{-4}$ | |
| extracellular exosome | GO:0070062 | $6.365\times10^{-4}$ | |
| intrinsic component of plasma membrane | GO:0031226 | $7.139\times10^{-4}$ | |
| integral component of plasma membrane | GO:0005887 | $7.513\times10^{-4}$ | |
| cell leading edge | GO:0031252 | $8.058\times10^{-4}$ | |
| collagen-containing extracellular matrix | GO:0062023 | $9.882\times10^{-4}$ | |
| extracellular space | GO:0005615 | $1.046\times10^{-3}$ | |
| non-motile cilium | GO:0097730 | $1.248\times10^{-3}$ | |
| cell cortex | GO:0005938 | $1.469\times10^{-3}$ | |
| microtubule organizing center | GO:0005815 | $4.932\times10^{-3}$ | |
| supramolecular complex | GO:0099080 | $5.687\times10^{-3}$ | |
| Golgi apparatus | GO:0005794 | $7.086\times10^{-3}$ | |
| microtubule cytoskeleton | GO:0015630 | $1.227\times10^{-2}$ | |
| collagen trimer | GO:0005581 | $1.235\times10^{-2}$ | |
| cytoplasmic vesicle | GO:0031410 | $1.452\times10^{-2}$ | |
| cytosol | GO:0005829 | $1.504\times10^{-2}$ | |
| intracellular vesicle | GO:0097708 | $1.522\times10^{-2}$ | |
| P granule | GO:0043186 | $1.779\times10^{-2}$ | |
| germ plasm | GO:0060293 | $1.779\times10^{-2}$ | |
| pole plasm | GO:0045495 | $1.779\times10^{-2}$ | |
| cell-substrate junction | GO:0030055 | $2.435\times10^{-2}$ | |
| sperm flagellum | GO:0036126 | $2.978\times10^{-2}$ | |
| neuron projection | GO:0043005 | $3.394\times10^{-2}$ | |
| lamellipodium | GO:0030027 | $3.481\times10^{-2}$ | |
| plasma membrane region | GO:0098590 | $4.662\times10^{-2}$ | |

Show evidence codes

**Fig. S51. GO enrichment analysis for scQTLs.**

**(A)** Summary scatter plot for GO terms with significant enrichment among the identified eGenes.

The y-axis shows $-\log_{10}(p_{adj})$ (p-values calculated as in (*170*)). **(B-D)** Tables list GO terms for biological process **(B)**, molecular function **(C)**, and cellular component **(D)** that are the most enriched among eGenes. BP = biological process; MF = molecular function; CC = cellular component.

Related to the supplementary section "***scQTLs***." This supplementary figure relates to **Fig. 4** and main text section "Determining cell-type-specific eQTLs from single-cell data."

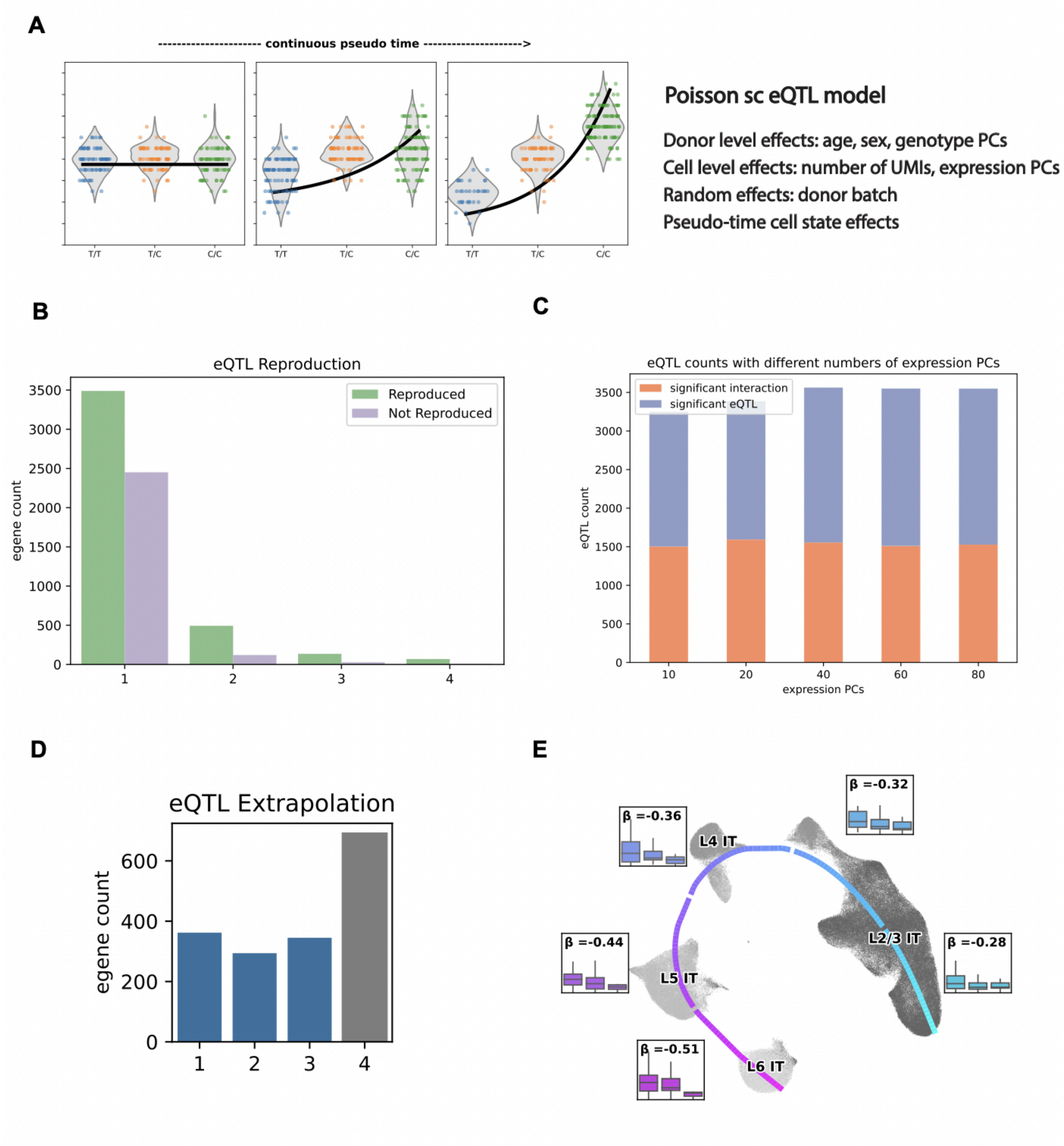


**Fig. S52. Pseudo-time trajectory-dependent dynamic scQTL analysis.**

**(A)** Schematic of Poisson regression model for pseudo-time trajectory-dependent dynamic scQTL analysis. We used this model to assess the interaction between genotype and the continuous pseudo-time. The plots show an example of an eSNP whose effect size increases over pseudo-time, represented by violin plots showing expression in cells of individuals who have homozygous reference (blue), heterozygous (orange), and homozygous alternate (green) genotypes at the eSNP locus. **(B)** Bar plot illustrates the number of top eQTLs for IT neurons (L2/3, L4, L5, and L6) that were successfully replicated by the PME model in the

SZBDMulti-Seq cohort. The x-axis represents the number of cell types where the scQTL was detected using the conventional pseudo-bulk method. **(C)** Bar plot illustrates the quantity of eQTLs associated with varying numbers of expression PCs. The orange labels signify the number of eQTLs with significant interaction terms (significance was determined by the likelihood ratio test). **(D)** Bar plot illustrates the number of eGenes for IT neurons (L2/3, L4, L5, and L6) that were identified with the pseudo-bulk approach. The x-axis represents the number of cell types in which the eGene was detected. The bars colored in blue indicate cases where eGenes were not detected in all four cell types. The top eQTLs corresponding to these eGenes were extrapolated using the PME model. **(E)** UMAP plot shows predicted trajectory for excitatory neurons in samples from the SZBDMulti-seq cohort. Box plots highlight the expression of *MGAM2*, stratified by eSNP genotype in each sample, for cell types in each cortical layer; effect size ($\beta$) values for the eSNP increase over pseudotime.

More detail in the supplementary section "***Dynamic scQTLs.***" This supplementary figure relates to **Fig. 4** and main text section "Determining cell-type-specific eQTLs from single-cell data."

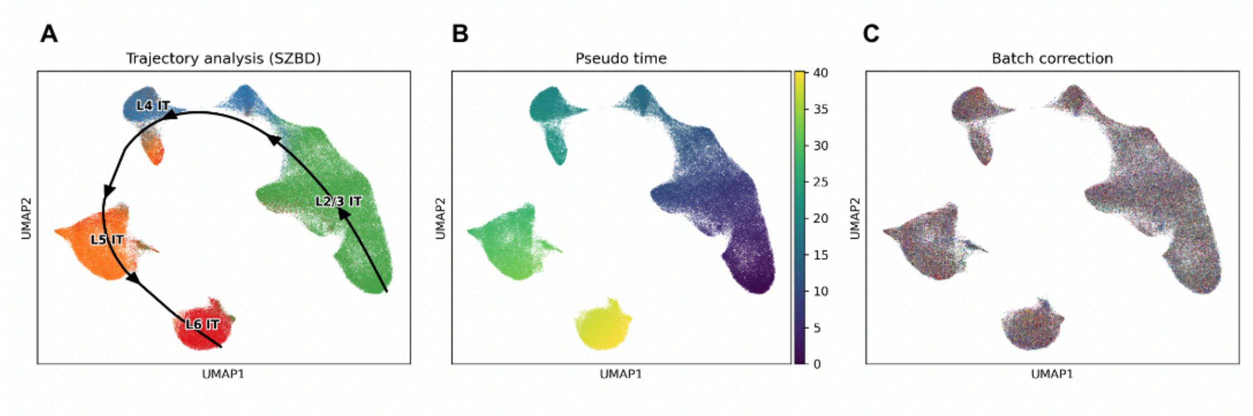


**Fig. S53. Dynamic scQTLs.**
This figure acts as a "shadow figure" for **Fig. 4H**, showing additional detail regarding the trajectory used for dynamic scQTLs. UMAP illustrates **(A)** the trajectory line; **(B)** the trajectory pseudo-time; and **(C)** batch correction conducted with SCALEX.

More detail in the supplementary section "***Dynamic scQTLs.***" This supplementary figure relates to **Fig. 4** and main text section "Determining cell-type-specific eQTLs from single-cell data."

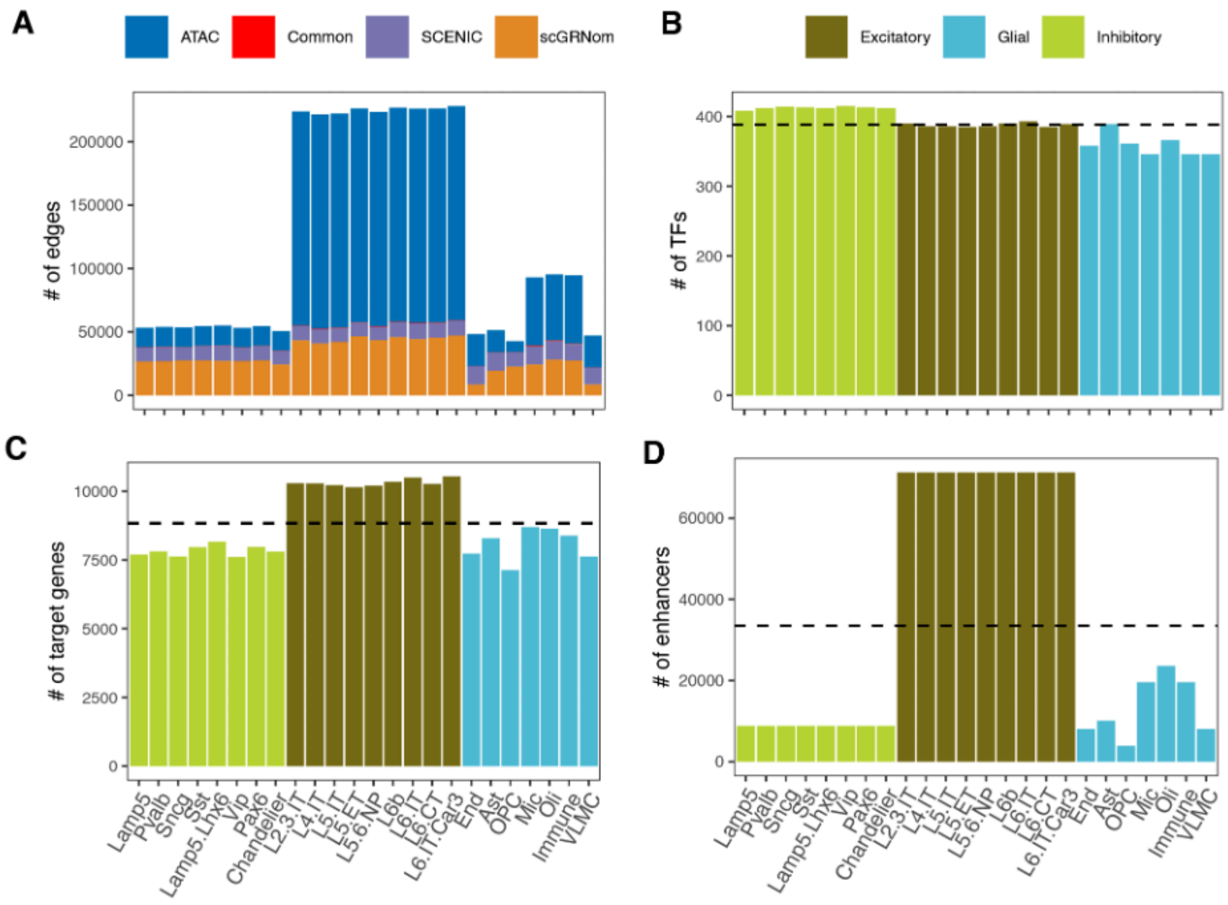


**Fig. S54. Summary of cell-type GRNs.**

**(A)** Distribution of edge sources (TF-target gene links) in the final GRNs with bars colored uniquely for each of the three sources used to construct the GRNs. Red bars show common distal (snATAC-seq) and proximal (SCENIC + scGRNom) edges. **(B-D)** Plots show the distribution of TFs **(B)**, target genes **(C)**, and enhancers **(D)** in the cell-type GRNs, colored according to three broad cell type groups. Dashed black lines represent the average number of elements identified in all cell types.

More detail in the supplementary section "***GRN evaluation.***" This supplementary figure relates to **Fig. 5** and main text section "Building a gene regulatory network for each cell type."

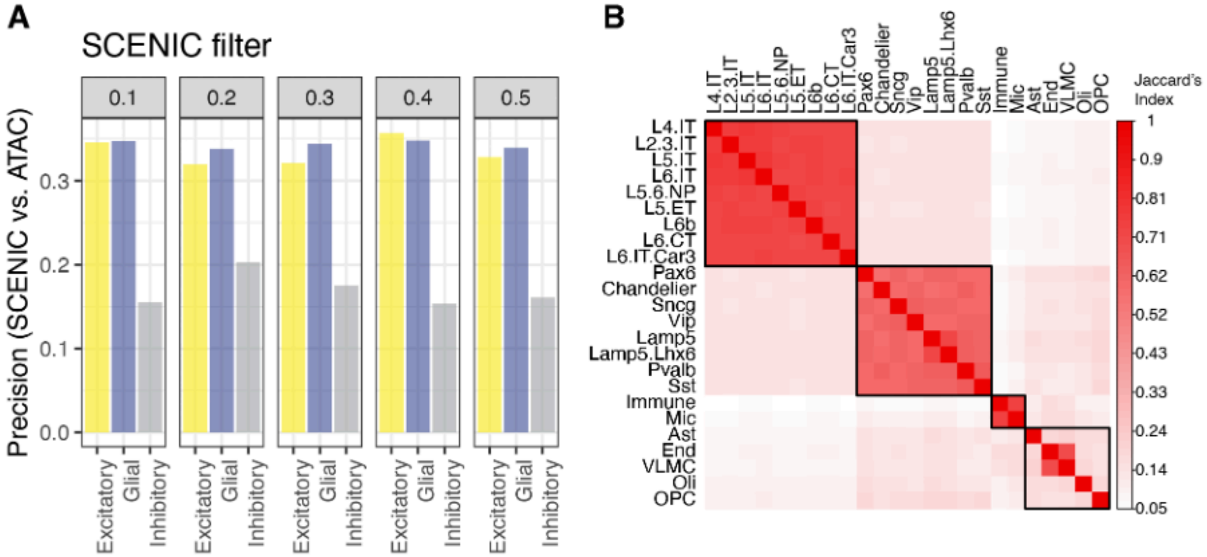


**Fig. S55. GRN visualization and evaluation.**

**(A)** Evaluation of SCENIC runs to determine edge-trimming threshold. The bar plot shows the precision (true positives/true positives + false positives; y-axis) of SCENIC runs at various edge weight thresholds (fraction of all predicted edges selected) using TF promoters identified in snATAC-seq as the benchmark. **(B)** Heatmap shows that cell types with the same parent class annotations are more similar to each other than those with different parent classes. The similarity between every possible pair of GRNs is estimated as the Jaccard Index and shaded along a low-to-high red gradient with darker colors showing greater similarity.

More detail in the supplementary section "***GRN evaluation.***" This supplementary figure relates to **Fig. 5** and main text section "Building a gene regulatory network for each cell type."

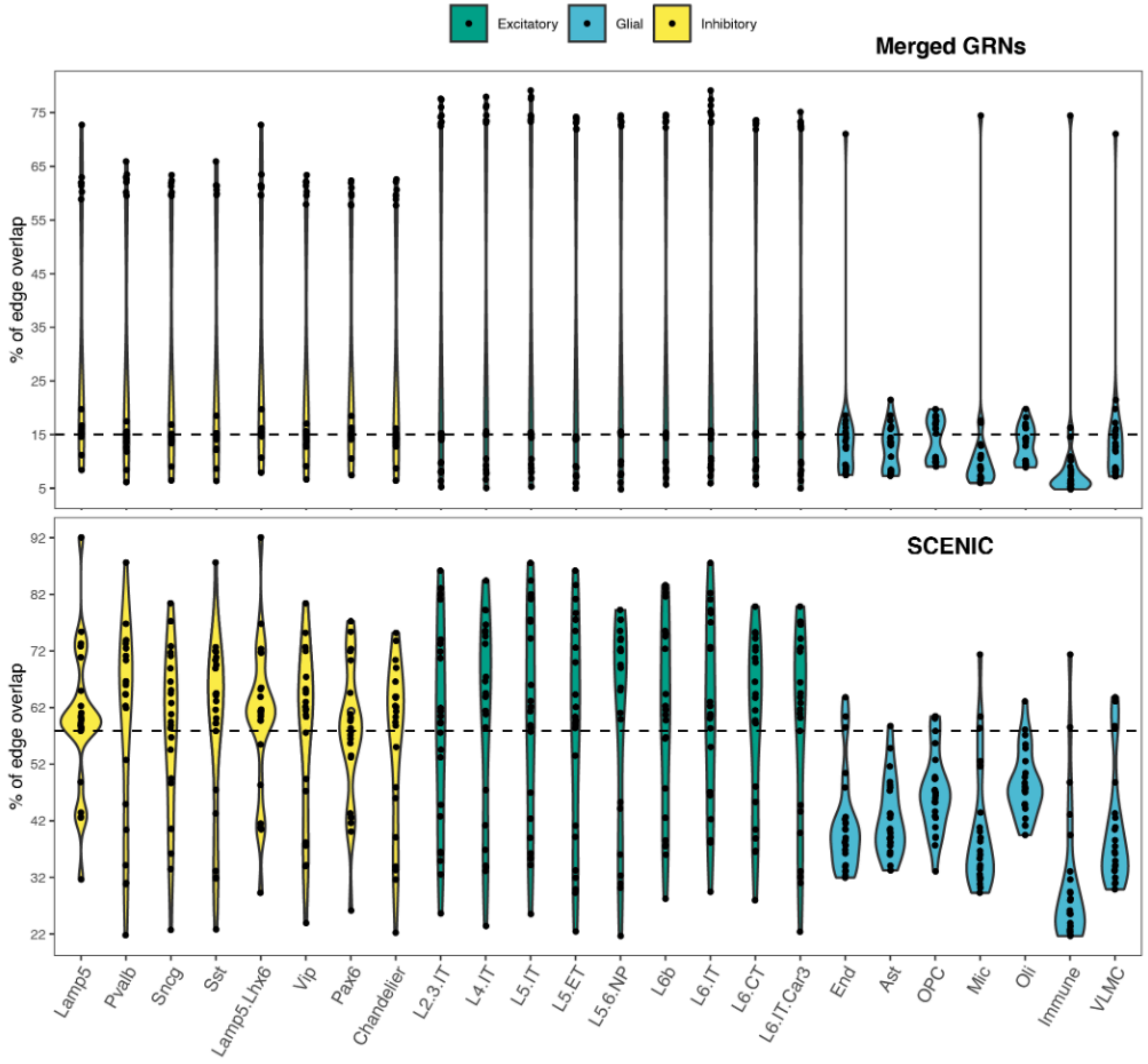


**Fig. S56. Overlaps between cell-type GRNs.**

For every pair of cell-type GRNs, the overlap between predicted edges was calculated as the fraction of all predicted edges that are common in both GRNs. The overlap is expressed as a percentage shown on the y-axis of the violin plot colored red, green, and blue for inhibitory, excitatory, and glial cell types, respectively. The dots in the jitter plots represent individual cell types. The black dashed line corresponds to the mean overlap across all GRNs.

More detail in the supplementary section "***GRN evaluation.***" This supplementary figure relates to **Fig. 5** and main text section "Building a gene regulatory network for each cell type."

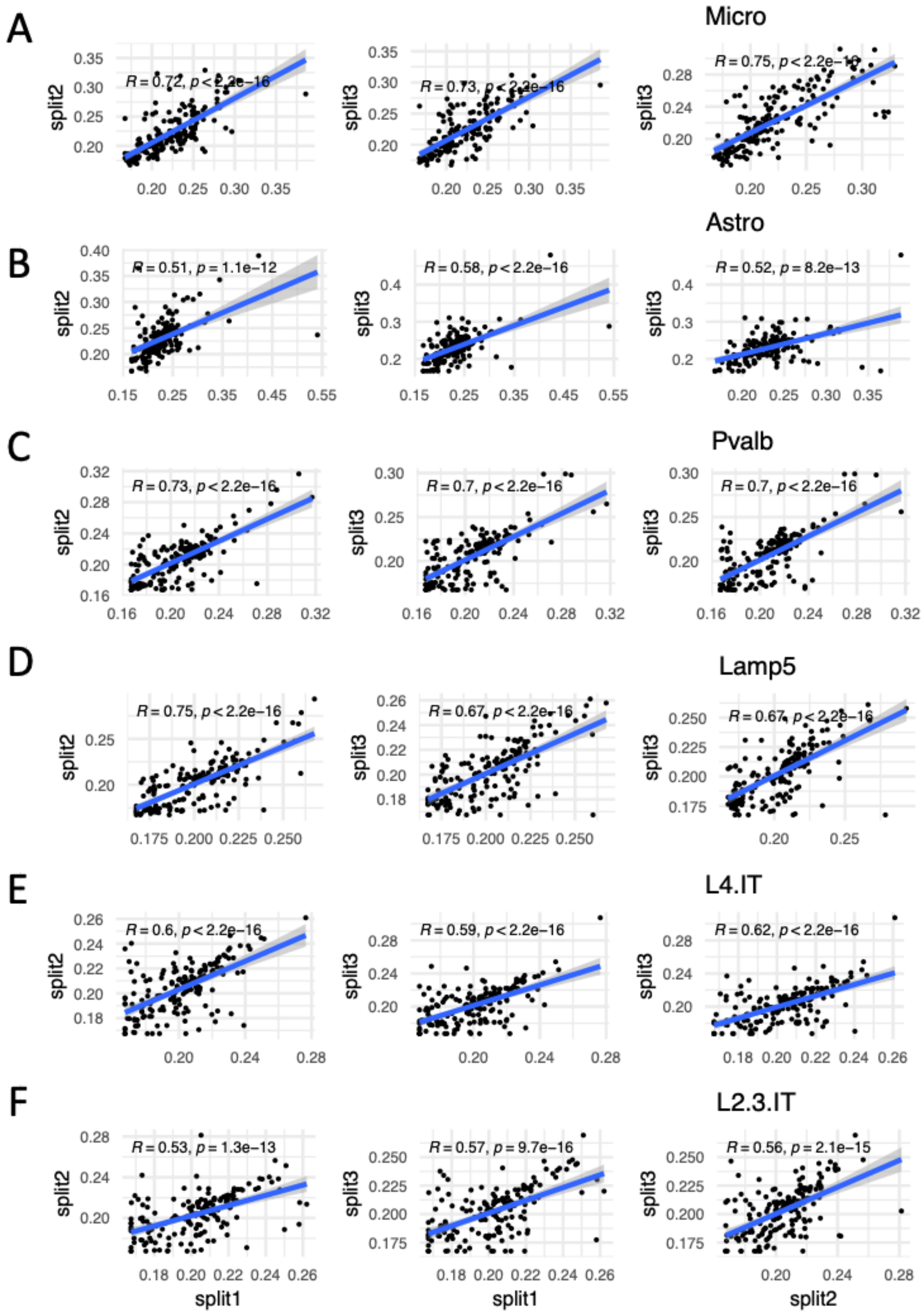


**Fig. S57. GRN stability.**

**(A-F)** Scatter plots show statistically significant (t-test) Pearson's correlation coefficient between the RSS scores from SCENIC across three random splits of the CMC cohort.

More detail in the supplementary section "***GRN evaluation.***" This supplementary figure relates to **Fig. 5** and the main text section "Building a gene regulatory network for each cell type."

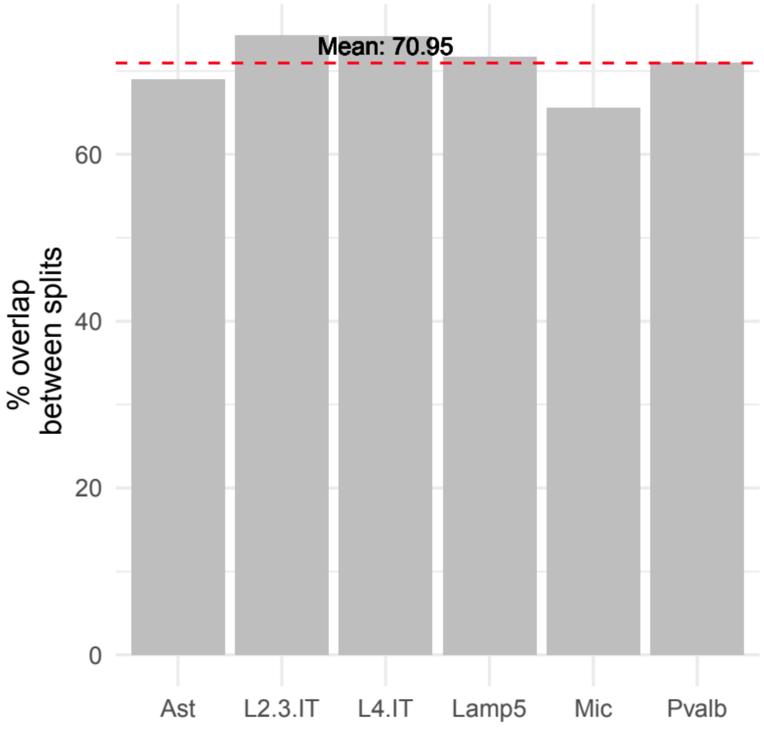


**Fig. S58. GRN stability.**

The plot shows the mean overlap between hub TFs identified across the three splits of the CMC cohort.

More detail is available in the supplementary section "***GRN evaluation*.**" This supplementary figure relates to **Fig. 5** and the main text section "Building a gene regulatory network for each cell type."

**A**

| TF | TG | Unified Score | Ast | Chandelie | End | Immune | L2.3.IT | L4.IT |
|---|---|---|---|---|---|---|---|---|
| RXRG | SLC4A2 | 4.92E-05 | 0.00140002 | 0.00089362 | | 0.0056757 | 0.00053856 | 0.00063063 |
| RXRG | PLIN3 | 4.46E-05 | | 0.00089362 | | 0.00567568 | 0.00053906 | 0.00063146 |
| RXRG | TBL3 | 3.95E-05 | | | 0.00100478 | 0.00567568 | 0.00053906 | 0.00063063 |
| RXRG | ADAMTS4 | 3.81E-05 | 0.0014 | | | 0.00567568 | 0.00053846 | 0.00063063 |
| RXRG | HIRIP3 | 3.73E-05 | | 0.00089362 | | 0.00567568 | 0.00053854 | 0.00063063 |
| RXRG | LINC01494 | 3.65E-05 | | | | 0.00567568 | | |
| RXRG | TNNC2 | 3.56E-05 | | | | 0.00567568 | 0.00053869 | 0.00063075 |
| RXRG | MINDY1 | 3.32E-05 | | 0.00089362 | | | 0.00053912 | 0.00063147 |
| RXRG | CLDN15 | 3.06E-05 | | 0.00089363 | 0.00100479 | | 0.00053846 | |
| RXRG | CRYBB2 | 3.06E-05 | | 0.00090939 | | | 0.00053869 | 0.00063076 |
| RXRG | TCAP | 3.02E-05 | 0.00140059 | 0.00089362 | | | 0.00053853 | |
| RXRG | GRWD1 | 3.00E-05 | | 0.00089362 | | | 0.00053854 | 0.00063063 |
| RXRG | CHRNB1 | 2.98E-05 | | | | | 0.00053847 | 0.00063063 |
| RXRG | LINC01484 | 2.91E-05 | | | | 0.00567568 | | |
| RXRG | NPM2 | 2.83E-05 | | 0.00089362 | | 0.00567568 | 0.00053869 | 0.00063076 |
| RXRG | ZNF229 | 2.77E-05 | | 0.0008971 | | | 0.00053847 | 0.00063064 |
| RXRG | LEFTY1 | 2.62E-05 | | | | 0.00567569 | | |
| RXRG | LOXL3 | 2.51E-05 | | 0.00089362 | | | | 0.00063063 |
| RXRG | CHST3 | 2.48E-05 | | 0.00089362 | | | 0.00053862 | 0.00063149 |
| RXRG | DNAH2 | 2.46E-05 | | 0.00089363 | | 0.00567568 | 0.00053852 | |
| RXRG | SPAG8 | 2.45E-05 | 0.0014 | | | 0.00567568 | | |
| RXRG | CDK5RAP2 | 2.45E-05 | | | | | 0.00053846 | 0.00063063 |
| RXRG | CCPG1 | 2.44E-05 | | | | | | |
| RXRG | BICD1 | 2.37E-05 | | 0.00089362 | | | | 0.00063063 |
| RXRG | PLCB2 | 2.36E-05 | | 0.00089423 | | | 0.00053846 | |
| RXRG | SLC12A5 | 2.35E-05 | | | | 0.00567568 | | |
| RXRG | SLC9A3R1 | 2.31E-05 | | 0.00089362 | | | | 0.00063158 |
| RXRG | ARMC7 | 2.30E-05 | | | | | 0.00053853 | 0.00063063 |
| RXRG | TTLL7 | 2.28E-05 | | 0.00089362 | 0.00100479 | | 0.00053846 | 0.00063063 |
| RXRG | NME5 | 2.27E-05 | | 0.00089787 | | 0.00567568 | 0.00053876 | |
| RXRG | CASKIN2 | 2.27E-05 | 0.0014 | | | | | 0.00063066 |

**B**

| TF | TG | Unified Score | Ast | Chandelie | End | Immune | L2.3.IT | L4.IT |
|---|---|---|---|---|---|---|---|---|
| RXRG | EGFR | 7.86E-06 | | | | | | |
| NR2F1 | EGFR | 5.84E-06 | | 0.00155556 | | | | |
| KLF15 | EGFR | 1.65E-06 | | | | | | |
| SP3 | EGFR | 1.31E-06 | | | | | | |
| CEBPG | EGFR | 6.15E-06 | | 0.0025 | | | | |
| STAT1 | EGFR | 2.60E-06 | | | | | | |
| FOXP1 | EGFR | 4.82E-06 | | | | | | |
| RXRA | EGFR | 2.74E-06 | | | | | | |
| TEAD1 | EGFR | 2.76E-06 | | | | | | |
| ZNF148 | EGFR | 3.89E-06 | 0.00024073 | | 0.00023423 | | | |
| STAT2 | EGFR | 4.00E-06 | | | | | | |
| NFIC | EGFR | 1.74E-06 | 0.00020937 | | | | | |
| EBF1 | EGFR | 9.46E-07 | 0.0003799 | | | | | |
| PLAG1 | EGFR | 2.53E-06 | | | 0.00051598 | | | |
| TBX19 | EGFR | 2.89E-05 | | | | | | |
| ZNF263 | EGFR | 2.35E-06 | | 0.00024823 | 0.00023397 | | | |
| TCF7L1 | EGFR | 3.00E-06 | | | | | | |
| TCF7L2 | EGFR | 1.19E-05 | | | | | | |
| SP2 | EGFR | 1.02E-06 | | | 0.00020782 | | | |
| PRDM1 | EGFR | 4.71E-06 | | | | | | |
| HNF4G | EGFR | 8.53E-06 | | | | | | |
| ESR2 | EGFR | 8.97E-06 | | | | | | |
| NR1H2 | EGFR | 1.49E-05 | | 0.00396226 | | | | |
| ZNF140 | EGFR | 6.01E-06 | | | | | | |
| EGR1 | EGFR | 8.82E-07 | | | | | | |
| NR3C1 | EGFR | 1.42E-06 | 0.00057221 | | | | | |
| ASCL1 | EGFR | 1.41E-06 | | | | | | |
| ZKSCAN5 | EGFR | 4.22E-06 | | | | | | |
| CEBPA | EGFR | 1.65E-05 | | 0.00272727 | | | | |
| SP1 | EGFR | 2.01E-06 | 0.00028632 | | | | | |
| MAZ | EGFR | 3.26E-06 | 0.00026569 | | 0.00016402 | | | |

**Fig. S59. Excerpts of the unified GRN diffusion score file.**

**(A)** An excerpt of the unified GRN diffusion score file. **(B)** An excerpt of the unified GRN diffusion score file that shows the up-regulators of the EGFR gene.

More detail in the supplementary section "***Unifying TF-target Regulons***." This supplementary figure relates to **Fig. 5** and the main text section "Building a gene regulatory network for each cell type."

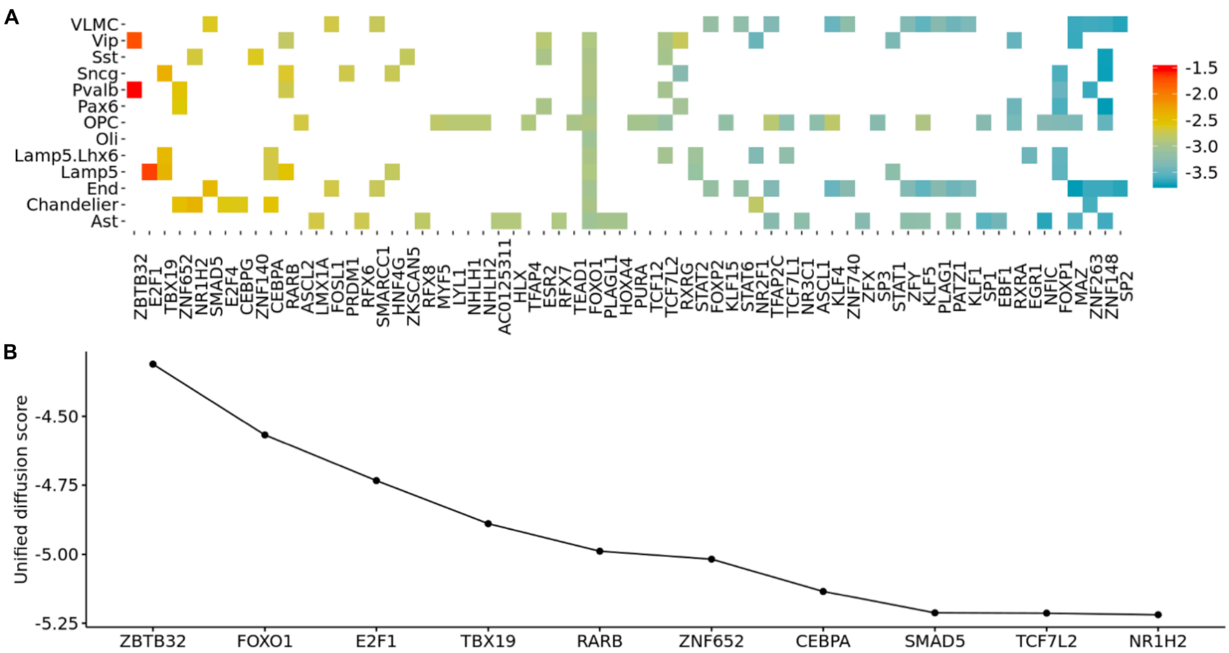


**Fig. S60. Upregulators of the EGFR gene, leveraging log-base-ten diffusion scores.**
**(A)** Upregulators in the cell-type-specific GRNs. Left to right, TFs are ordered in descending order by their diffusion score. **(B)** Top 10 upregulators in the unified GRN.

More detail in the supplementary section "***Unifying TF-target Regulons*.**" This supplementary figure relates to **Fig. 5** and main text section "Building a gene regulatory network for each cell type."

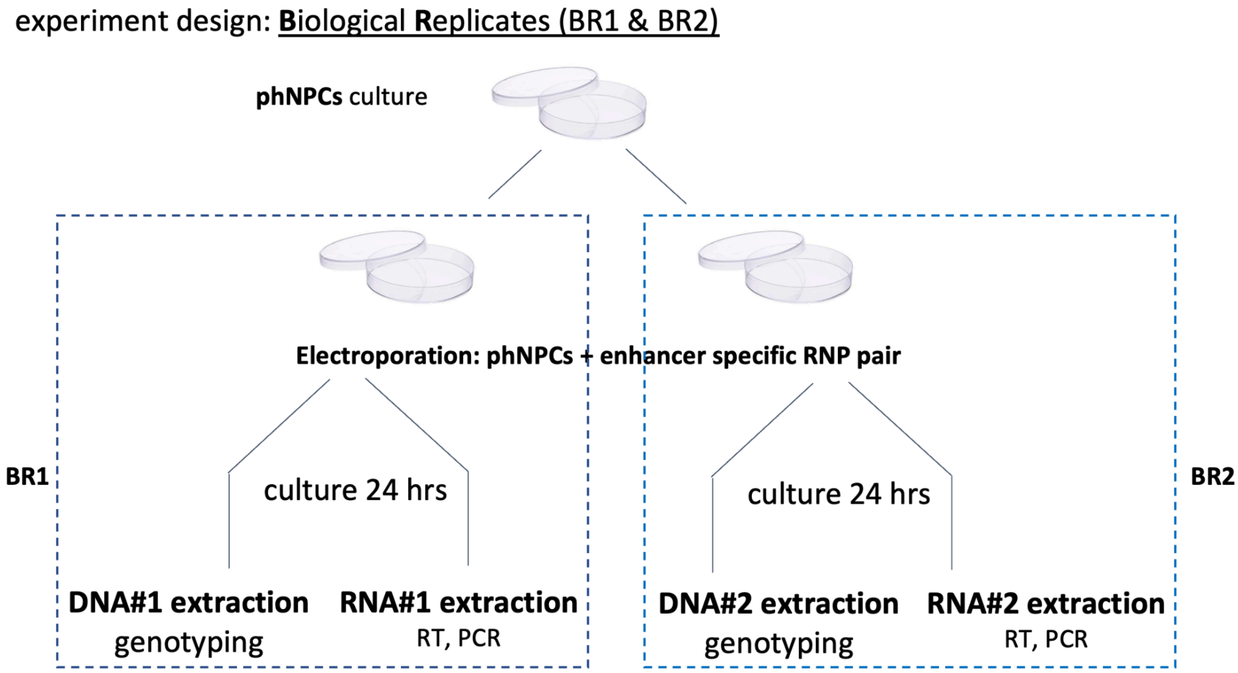


**Fig. S61. Experimental design for biological replicates in CRISPR enhancer KO experiments.**
Schematic of biological replicates and experimental design for CRISPR experiments in phNPCs.

More detail in the supplementary section "***CRISPR Validation*.**" This supplementary figure relates to **Fig. 5** and main text section "Building a gene regulatory network for each cell type."

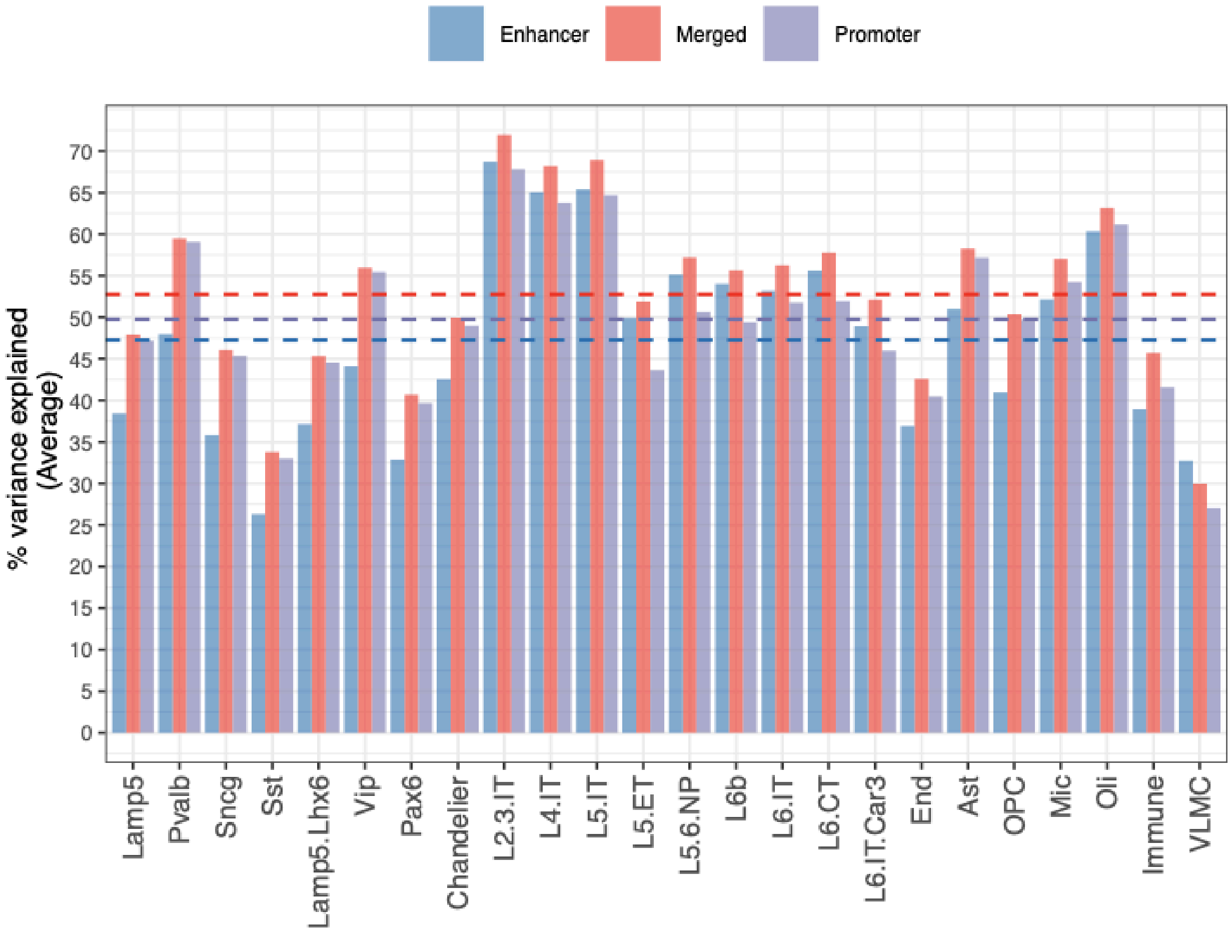


**Fig. S62. Network-based regression.**

Network-based regression was used to estimate the average percentage of gene expression variance (y-axis) explained by the cell-type GRN models (x-axis). The dotted lines colored blue, purple, and red show the average across enhancer, promoter, and merged edges, respectively.

More detail in the supplementary section "***GRN evaluation.***" This supplementary figure relates to **Fig. 5** and main text section "Building a gene regulatory network for each cell type."

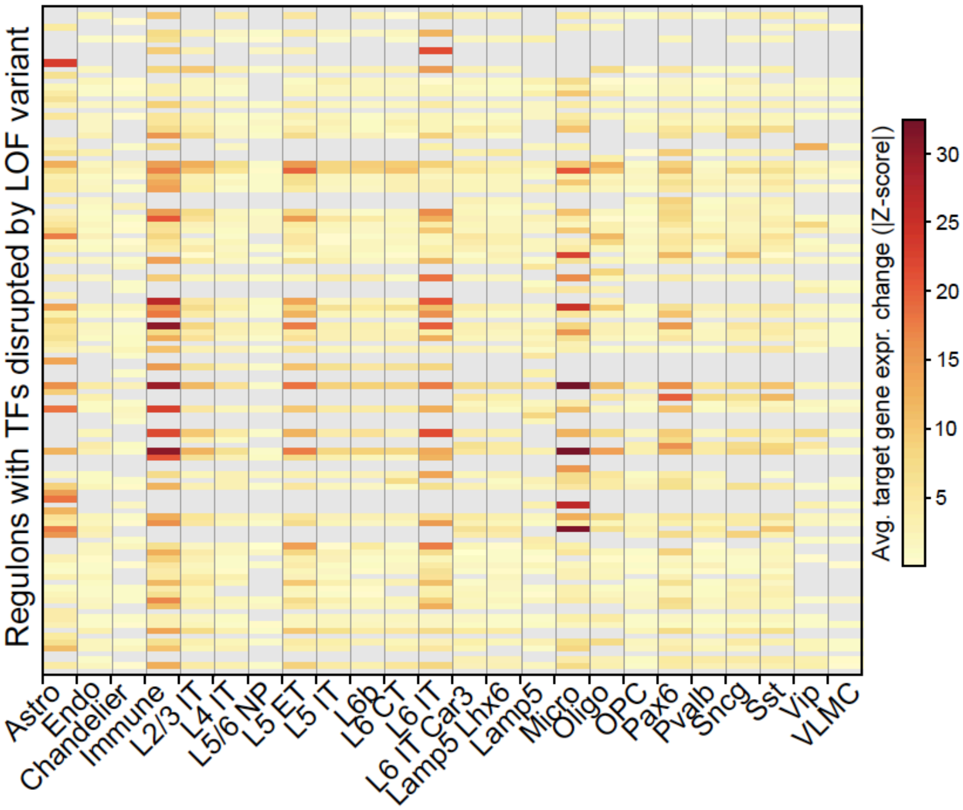


**Fig. S63. Effect of LOF variants in TF on regulon gene expression.**

Heatmap showing the average absolute z-score changes in expression of target genes in 112 regulons whose TFs are affected by LOF variants (y-axis) across cell types (x-axis). Gray boxes indicate that a particular TF does not have an active regulon in the cell type. z-scores were calculated by comparing expression of target genes in the regulon among individual cells from samples with and without the LOF variant in the regulon TF.

More detail in the supplementary section "***Genotype processing.***" This supplementary figure relates to **Fig. 5** and main text section "Building a gene regulatory network for each cell type."

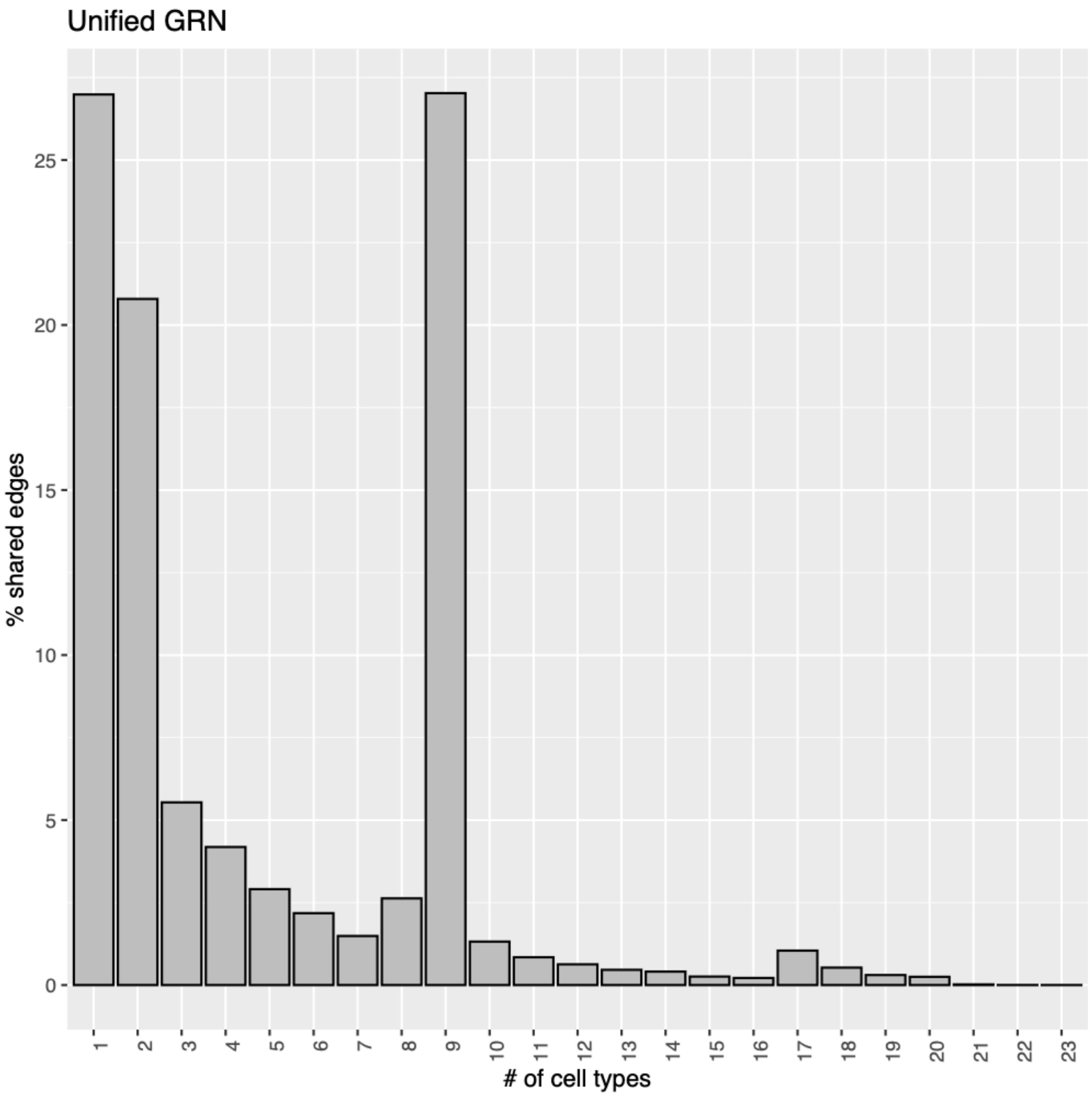


**Fig. S64. Unified GRN.**
The barplot illustrates the proportion of edges shared among different cell types within the unified GRN, which is constructed by merging individual cell-type-specific GRNs. The y-axis represents the percentage of edges (corresponding to TF-target gene links) shared across varying numbers of cell types, as indicated on the x-axis.

More detail in the supplementary section "***Network Characterization***." This supplementary figure relates to **Fig. 5** and main text section "Building a gene regulatory network for each cell type."

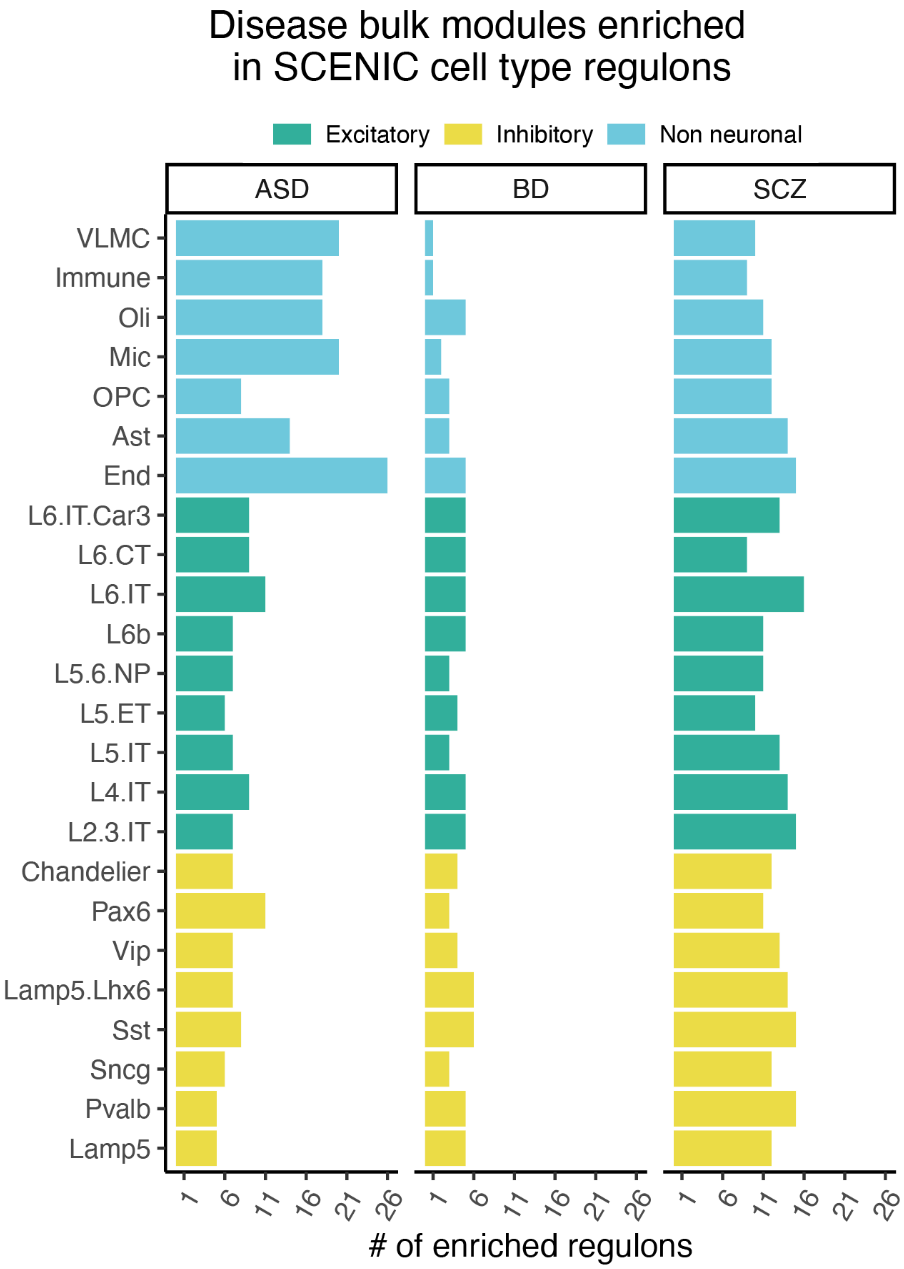


**Fig. S65. Comparison with bulk co-expression.**

The barplots show the enrichment of bulk disease co-expression modules from (*100*) Gandal et al., 2018, within our cell-type GRNs. Within each cell type shown on the y-axis, the number of regulons enriched with co-expression modules for ASD, bipolar disorder (BD), and schizophrenia (SCZ) are shown on the x-axis.

More detail in the supplementary section "***Network Characterization.***" This supplementary figure relates to **Fig. 5** and main text section "Building a gene regulatory network for each cell type."

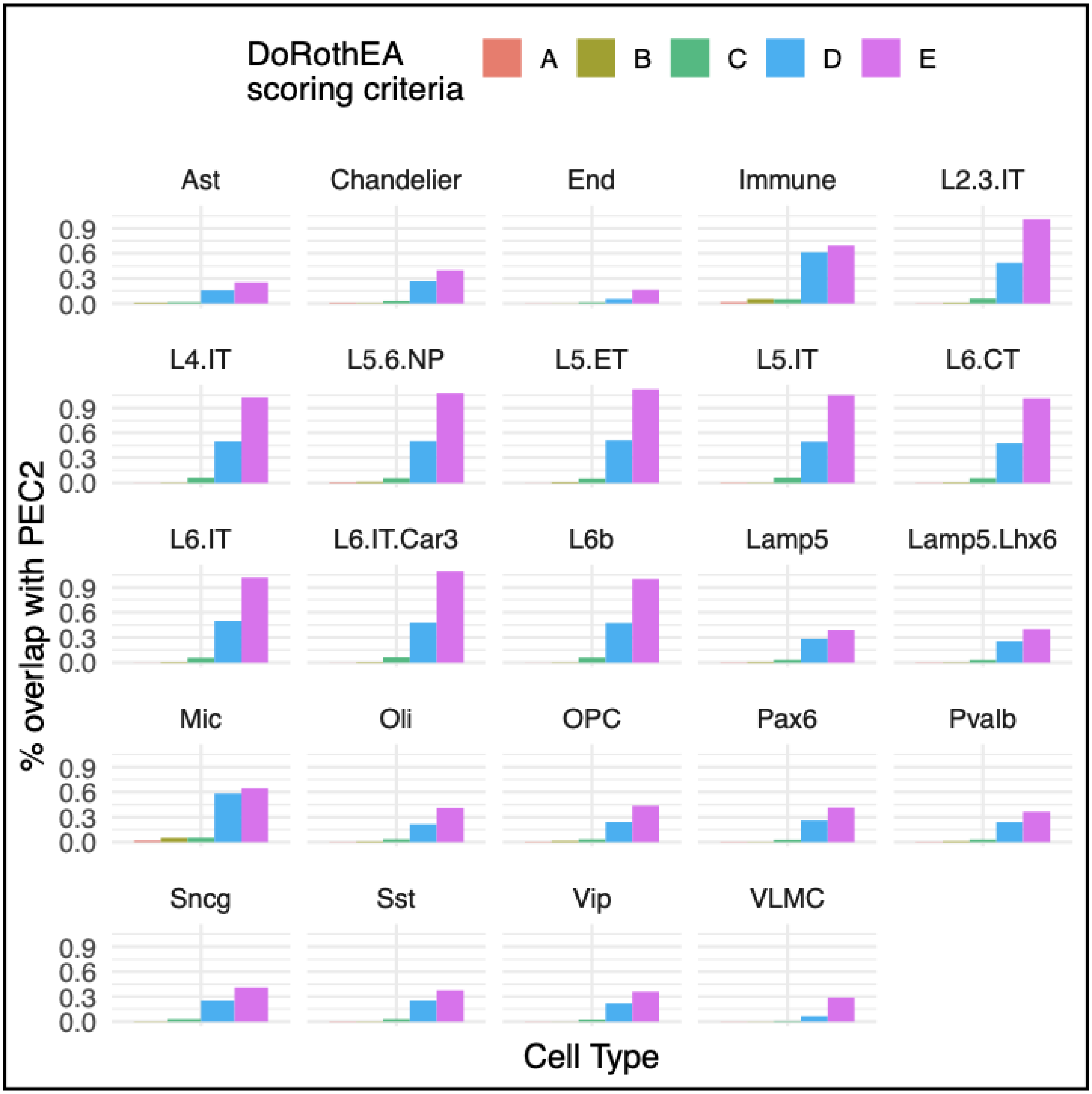


**Fig. S66. Comparison of cell-type GRNs with tissue-naive GRNs.**

The percentage overlap (y-axis) between each of the cell-type GRNs with different scoring criteria of tissue-naive GRNs in the DoRothEA database shows the largest overlap with predicted edges.

More detail in the supplementary section "***GRN evaluation.***" This supplementary figure relates to **Fig. 5** and main text section "Building a gene regulatory network for each cell type."

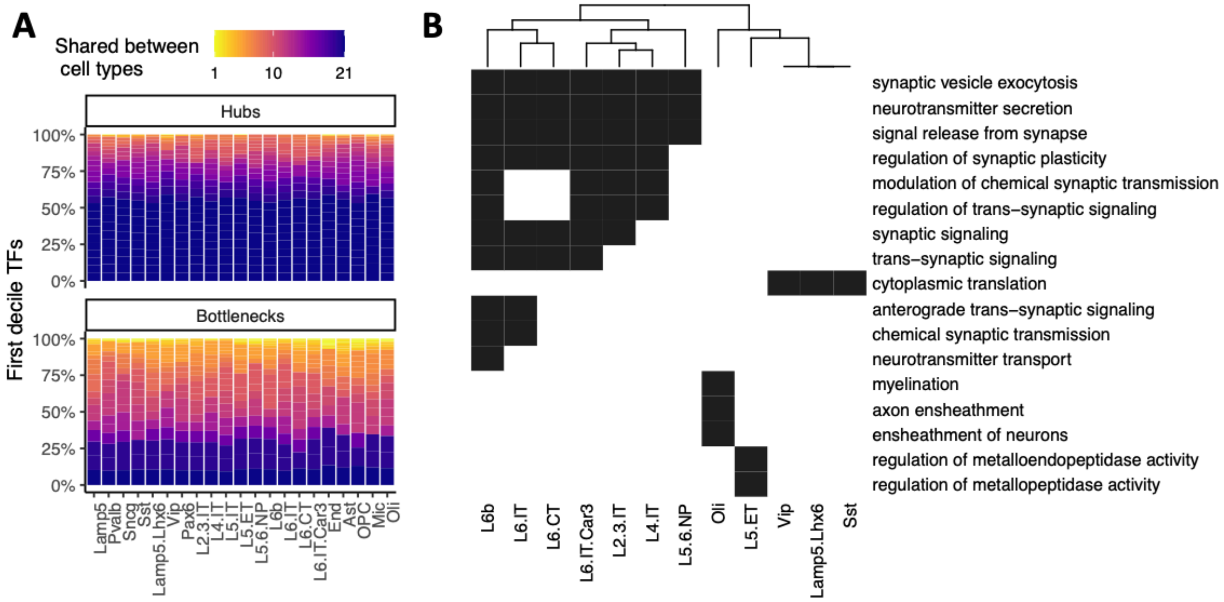


**Fig. S67. Comparison of GRN centralities and GO targets of bottlenecks.**

**(A)** Similarity between centralities across cell types. For each cell-type GRN, the first decile TFs, ranked based on the centrality scores (out-degree or betweenness), were selected. The y-axis of the stacked bar plot shows the fraction of TFs within each cell type that are shared with other cell types. Lighter shades show more uniqueness and darker shades indicate more commonness. **(B)** GO biological process (y-axis) enrichment results of bottleneck TFs identified across all cell types (x-axis). Cells of the heatmap are shaded along a gradient representing corrected p-values resulting from the hypergeometric tests used for testing enrichment. Darker shades indicate stronger enrichment and vice versa.

More detail in the supplementary section "***Network Characterization.***" This supplementary figure relates to **Fig. 5** and main text section "Building a gene regulatory network for each cell type."

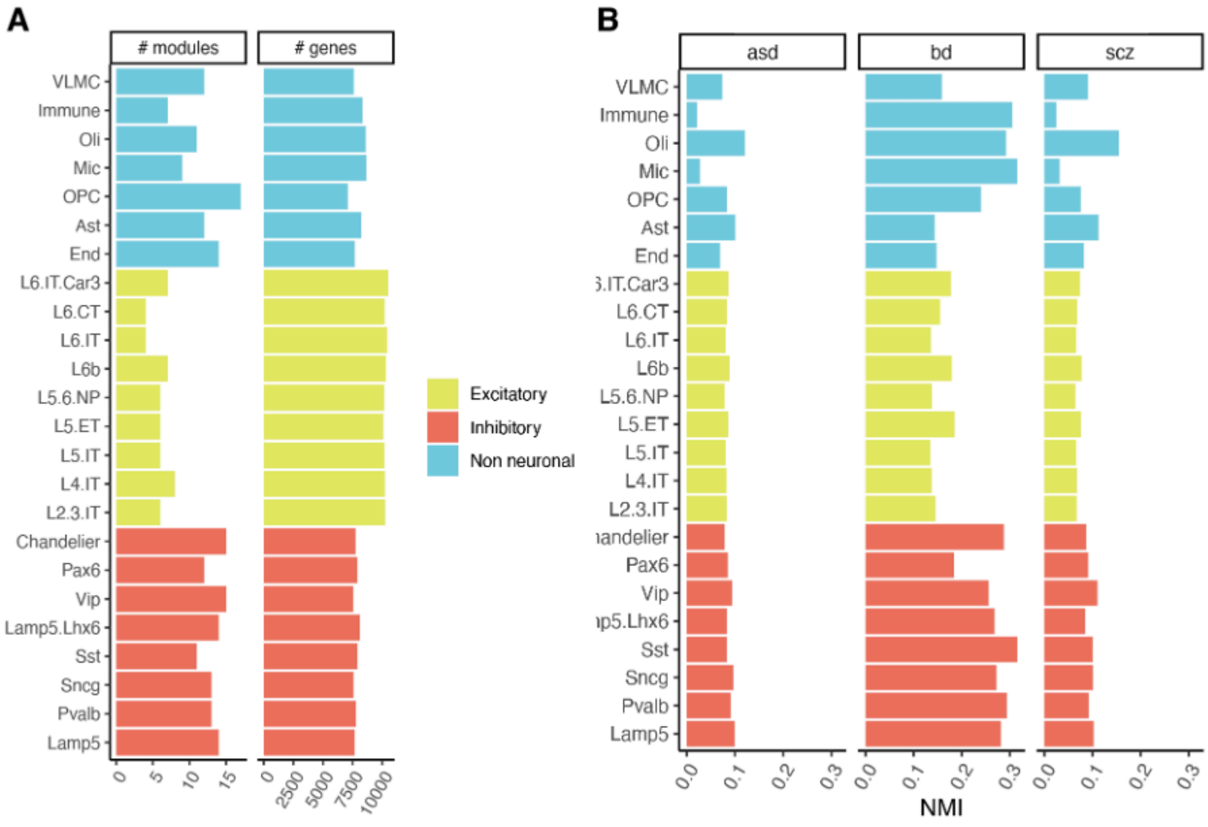


**Fig. S68. Summary of the gene module analysis.**

**(A)** The left facet of the bar plot shows the number of modules identified within cell-type GRNs (y-axis) and the right facet shows the number of genes within those modules. **(B)** Normalized mutual information (NMI; x-axis) is used as a metric to gauge similarity between cell-type modules and bulk disease modules (left facet: ASD; middle facet: bipolar disorder [BD]; right facet: schizophrenia [SCZ]).

More detail in the supplementary section "***Network Characterization.***" This supplementary figure relates to **Fig. 5** and the main text section "Building a gene regulatory network for each cell type."

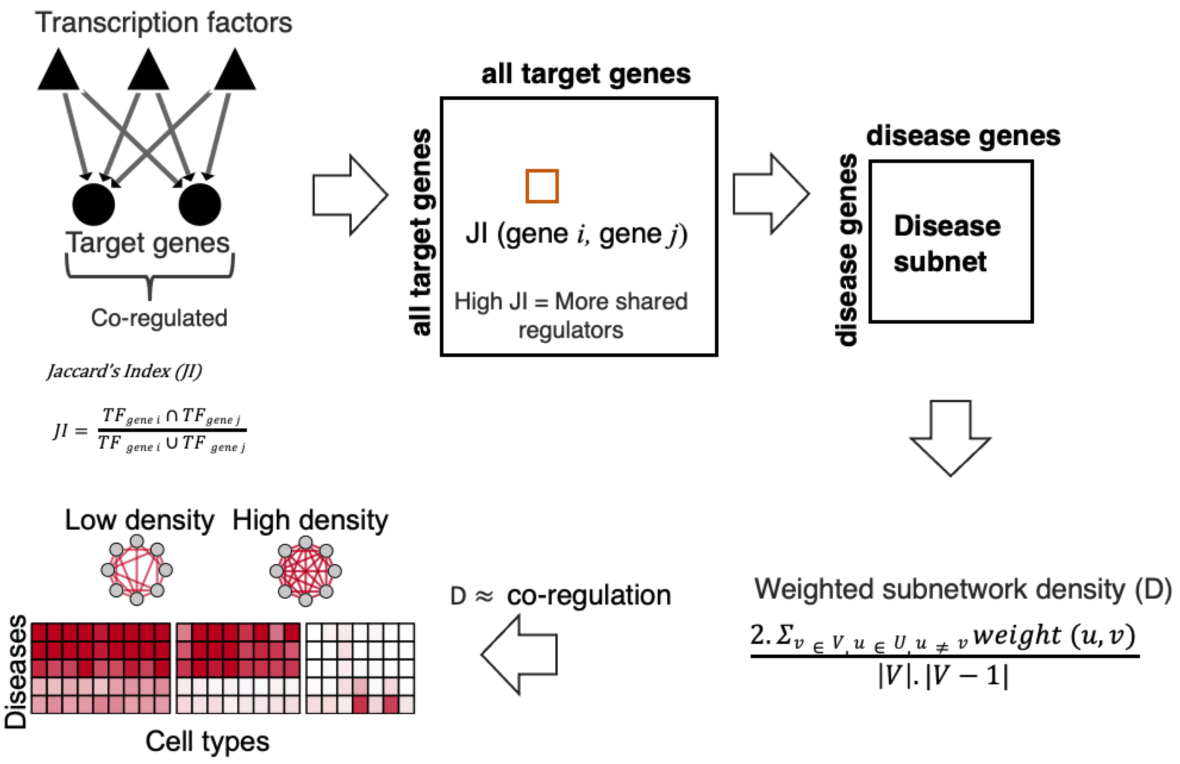


**Fig. S69. Schematic showing analysis of disease-gene co-regulatory subnetworks.**
For each cell type, the directed unweighted TF→target gene interactions in GRNs are converted to undirected weighted TG→target gene 'co-regulatory' networks based on the similarity between the pair's predicted regulators (TFs). This similarity is measured as the Jaccard Index. Then, for a given list of disease-risk genes, a subnetwork consisting of only those genes is extracted. The weighted density of the subnetwork is recorded and used as a proxy for 'coregulation'. The p-value is calculated based on randomly picking disease genes 1,000 times and counting the number of times the random density is greater than or equal to the observed density. The -1*$\log_{10}$(p-values) are used to generate the heatmap.

More detail in the supplementary section "***Network Characterization.***" This supplementary figure relates to **Fig. 5** and main text section "Building a gene regulatory network for each cell type."

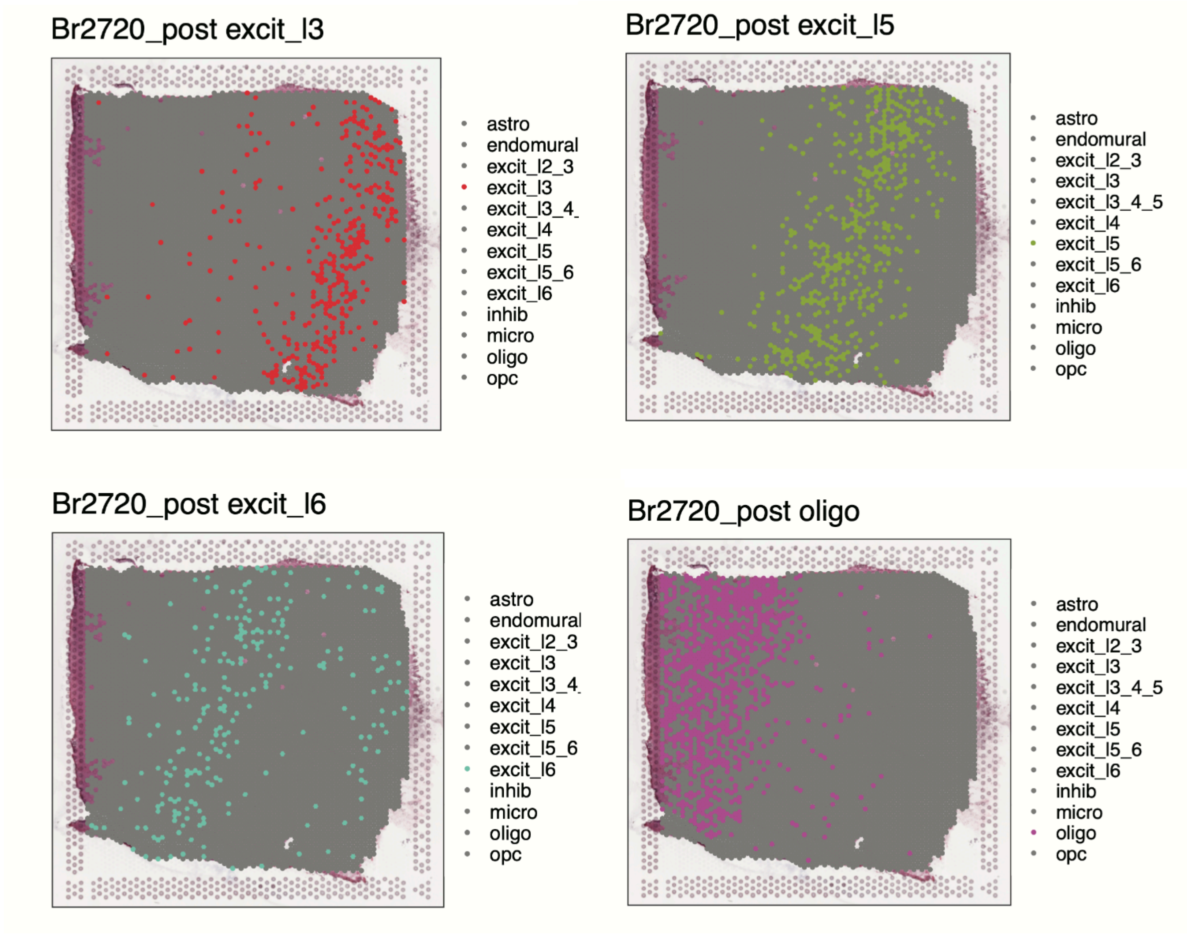


**Fig. S70. Spatial locations of specific snRNA-seq-labeled cell types.**
Shown are the spatial locations of specific snRNA-seq-labeled cell types, showing layer specificity. We obtained the specific cell type through both cell-type-specific and sample-specific normalization (details in section 6.3). Examples show specifically “excit_l3”, “excit_l5”, “excit_l6”, and “oligo” annotated cells.

More detail in the supplementary section "***Cell-to-Cell Network*.**" This supplementary figure relates to **Fig. 6** and the main text section "Constructing a cell-to-cell communication network."

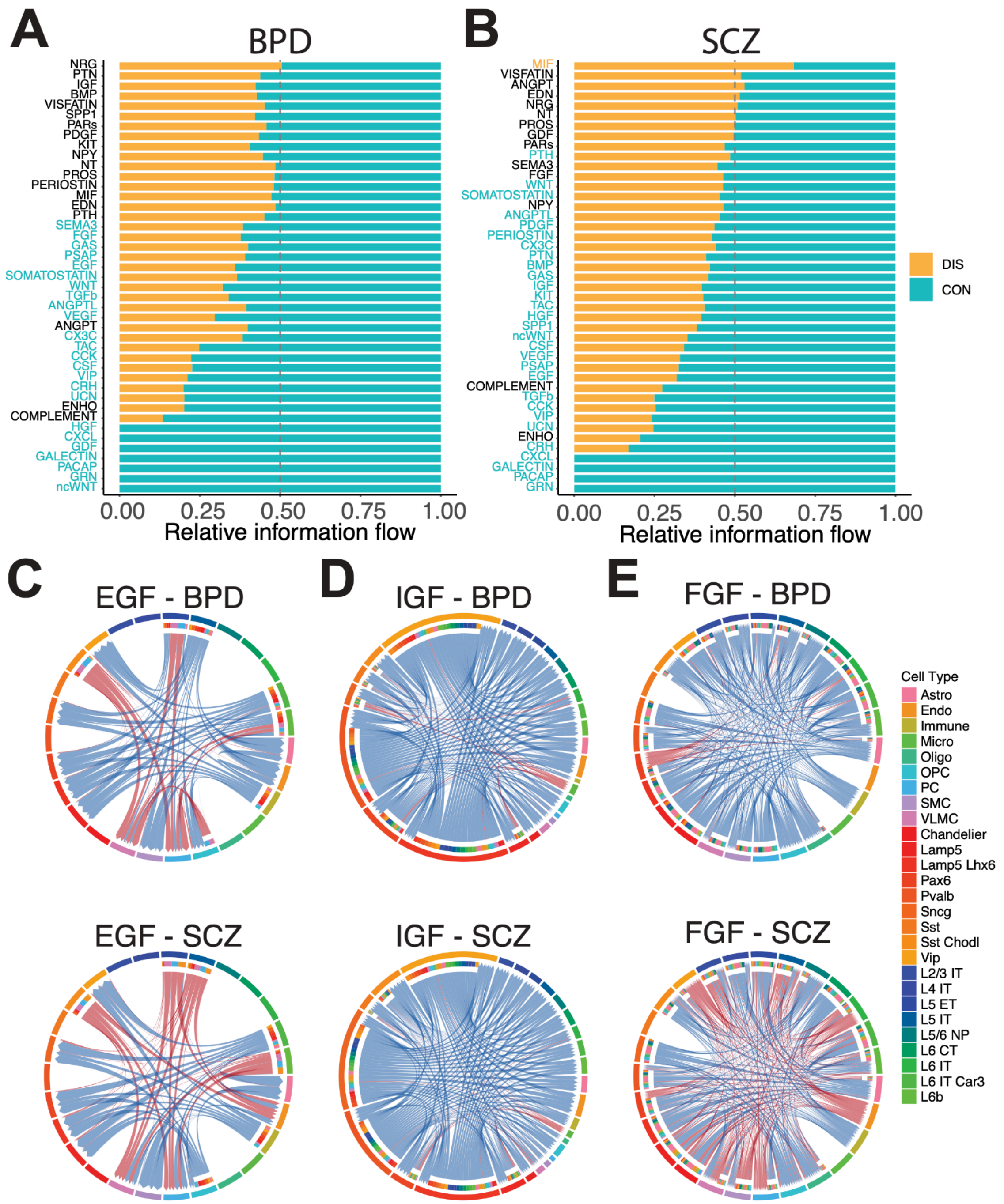


**Fig. S71. Cell-to-cell communication analysis.**

**(A)** Dysregulated ligand-receptor signaling pairs in bipolar disorder (BPD). **(B)** Dysregulated ligand-receptor signaling pairs in schizophrenia (SCZ). **(C)** Cell-type-specific differential interaction in the EGF signaling pathway for bipolar disorder and schizophrenia individuals. **(D)** Cell-type-specific differential interaction in the IGF signaling pathway for bipolar disorder and schizophrenia individuals. **(E)** Cell-type-specific differential interaction in the FGF signaling pathway for individuals with bipolar disorder or schizophrenia.

More detail in the supplementary section "***Cell-to-Cell Network*.**" This supplementary figure relates to **Fig. 6** and the main text section "Constructing a cell-to-cell communication network."

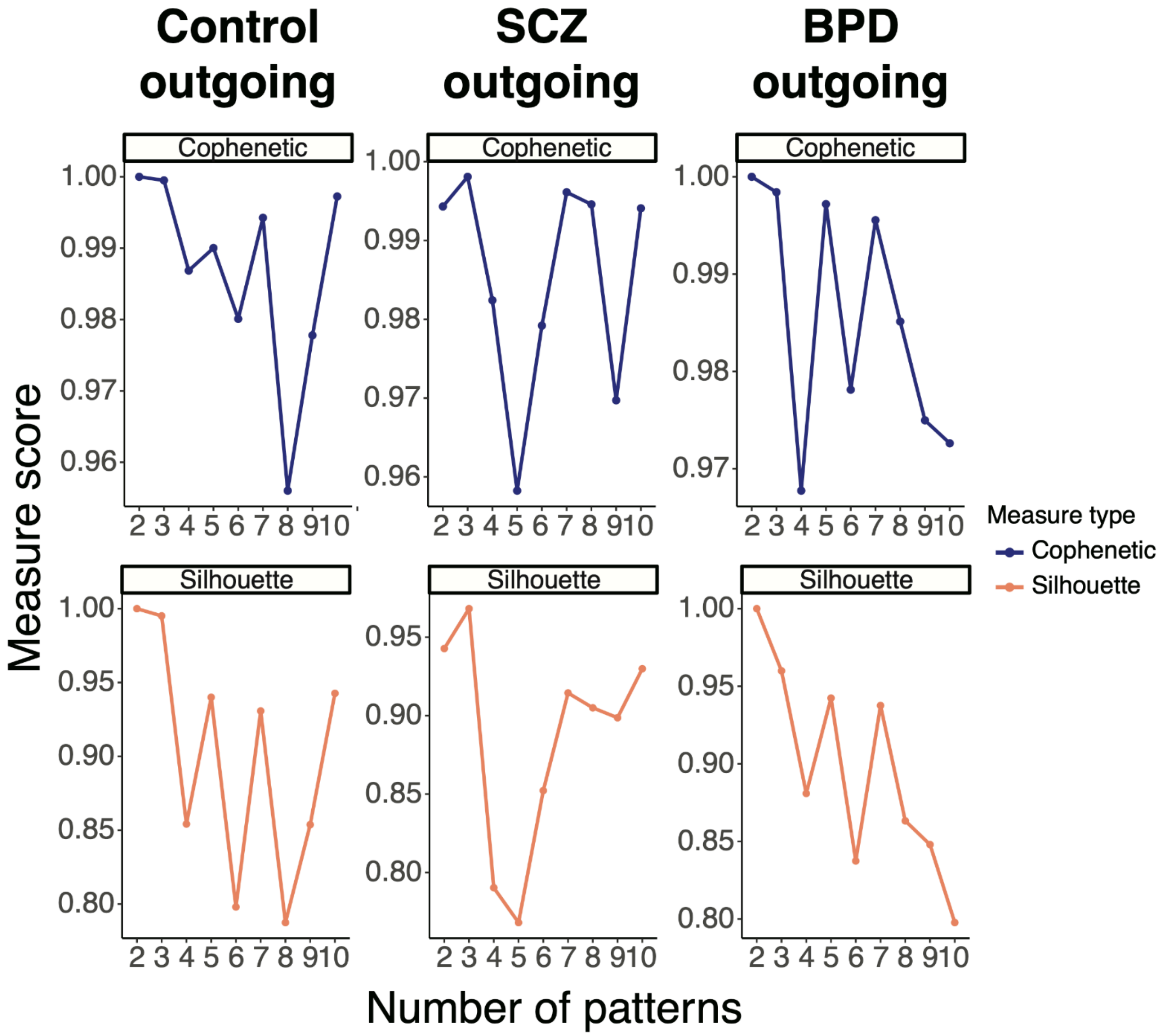


**Fig. S72. Co-phenetic and silhouette scores for various ranks of NMF.**
We calculated co-phenetic and silhouette scores repeatedly for different pattern numbers in control, bipolar disorder, and schizophrenia individuals to determine the optimal scores. We also wanted to further clarify NMF latent patterns (**Fig. 6B**). For example, we see that the inhibitory Vip cell type and the Vip signaling pathway (specifically, VIP-VIPR1) both belong to pattern 2. The pattern represents a hidden underlying structure to the cell-to-cell communication network that is directly observed through cell-type and signaling pathways. The fact that both belong to pattern 2 makes sense, as the VIP interneurons are predominantly characterized by the Vip gene and its associated signaling pathway.

More detail in the supplementary section "***Cell-to-Cell Network.***" This supplementary figure relates to **Fig. 6** and the main text section "Constructing a cell-to-cell communication network."

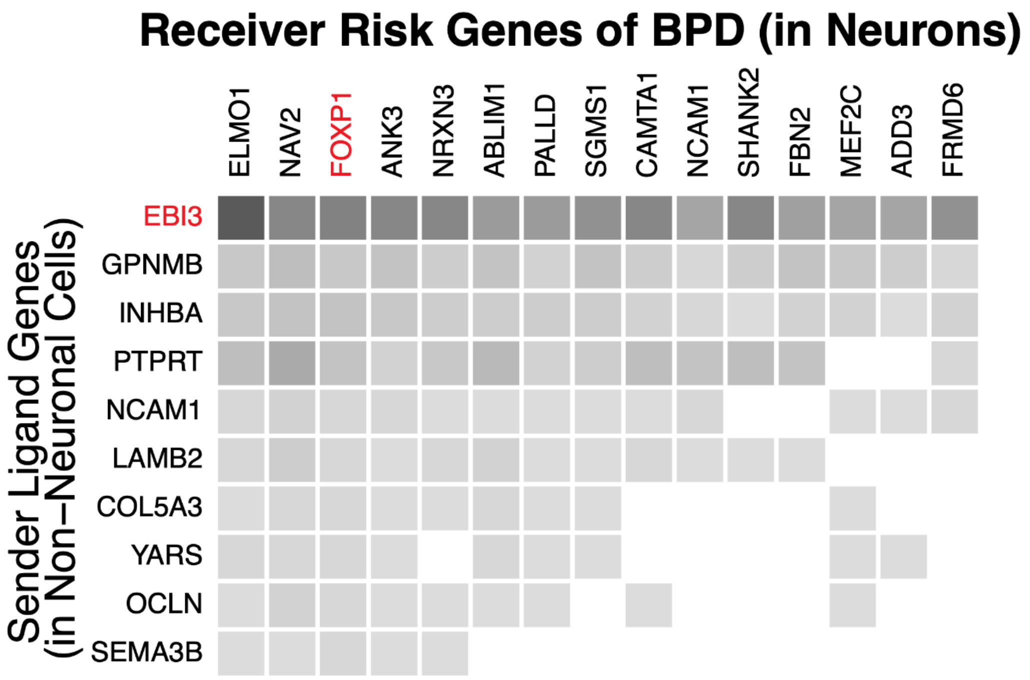


**Fig. S73. Ligand-target risk gene regulation in bipolar disorder.**
Shown are the predicted likelihoods that ligand genes in non-neuronal cells (y-axis) regulate bipolar disorder-associated risk genes (x-axis) in neuronal cell types, with the neurological risk gene *FOXP1* highlighted in red.

More detail in the supplementary section "***Cell-to-Cell Network*.**" This supplementary figure relates to **Fig. 6** and the main text section "Constructing a cell-to-cell communication network."

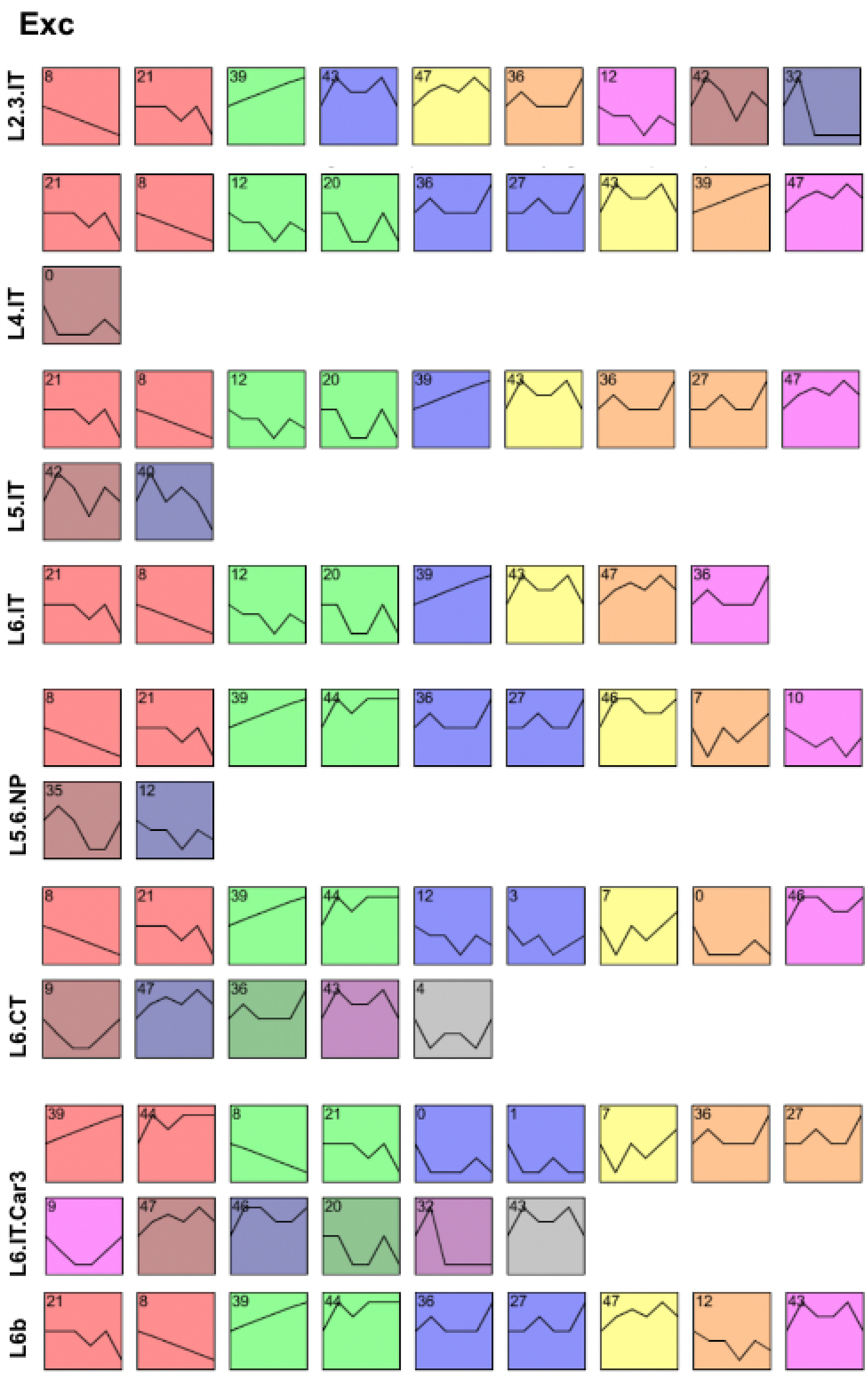
Exc
L2.3.IT
L4.IT
L5.IT
L6.IT
L5.6.NP
L6.CT
L6.IT.Car3
L6b

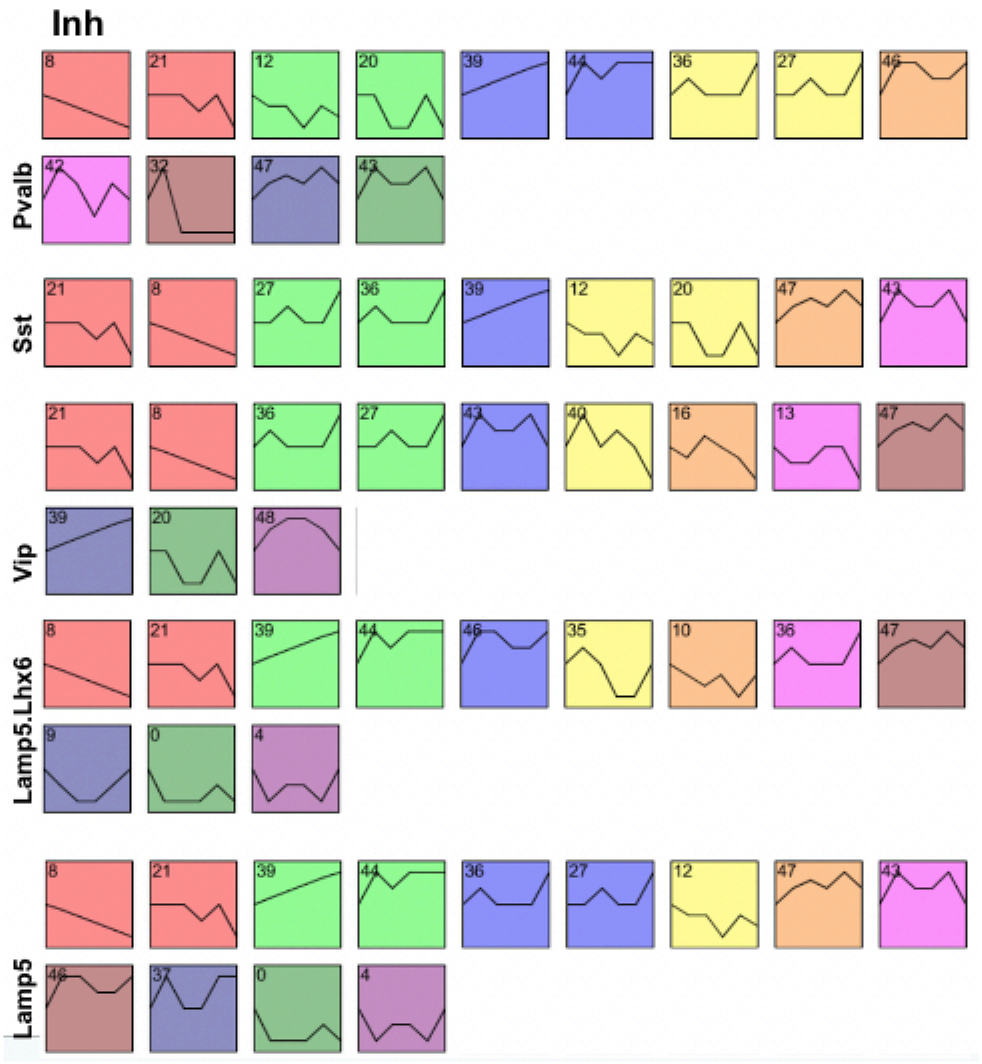
Inh
8 21 12 20 39 44 36 27 46
Pvalb
42 32 47 43
Sst
21 8 27 36 39 12 20 47 43
Vip
21 8 36 27 43 40 16 13 47
39 20 48
Lamp5.Lhx6
8 21 39 44 46 35 10 36 47
9 0 4
Lamp5
8 21 39 44 36 27 12 47 43
46 37 0 4

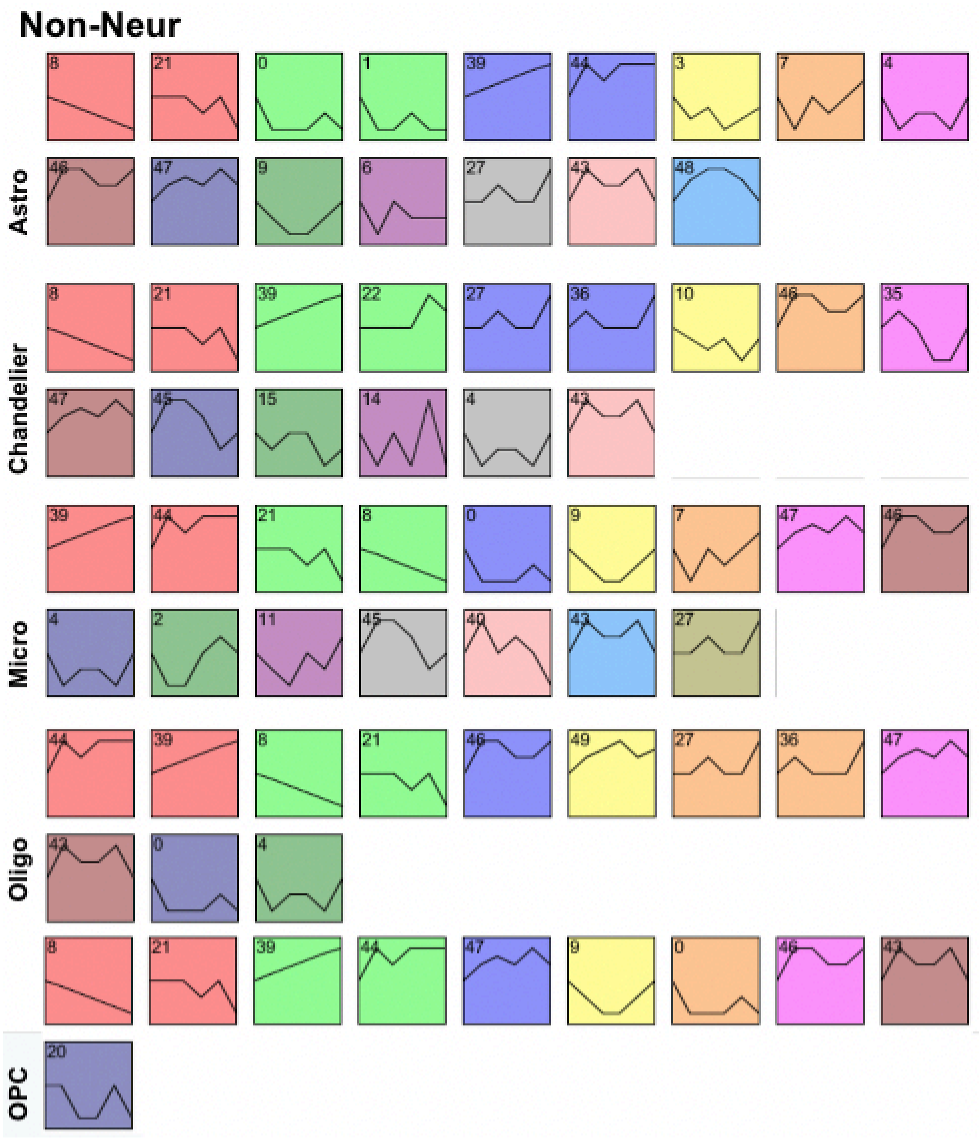


**Fig. S74. STEM analysis.**

STEM clustering analysis was applied to model gene expression changes across age for each cell type. Each cell type exhibited a variety of gene expression patterns. The x-axis represents six aging intervals (25-40, 40-50, 50-60, 60-70, 70-80, and 80-90 years old); the y-axis depicts gene expression levels characterized as steady, increasing, or decreasing.

More detail in the supplementary section "***Aging STEM.***" This supplementary figure relates to **Fig. 7** and main text section "Assessing cell-type-specific transcriptomic and epigenetic changes in aging."

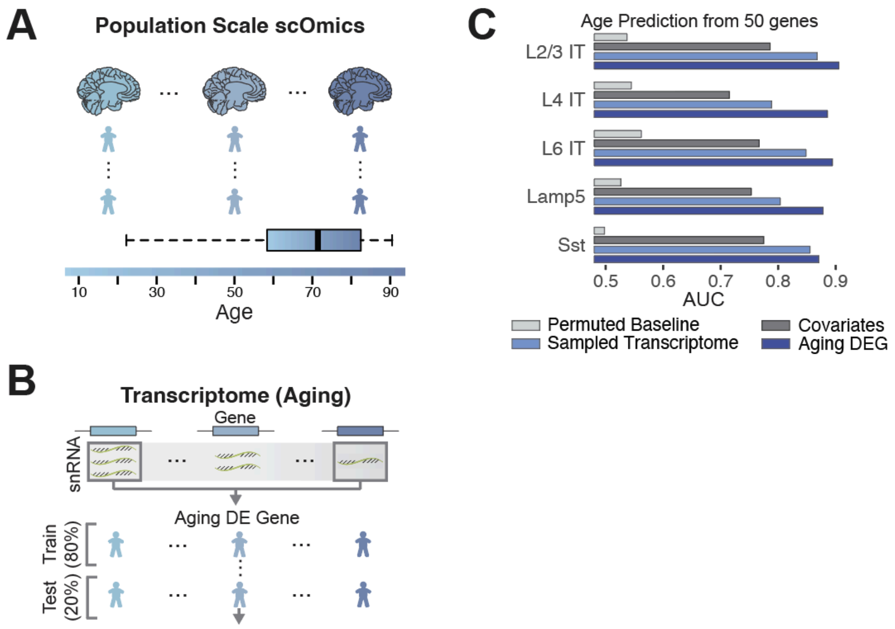


**Fig. S75. A simple aging classification model.**

**(A)** Age distribution of individuals used in a simple aging classification model. **(B)** Schematic of a simple classification model for aging based on the single-cell transcriptome. **(C)** AUC performance of the simple aging model by cell type. Comparisons are shown for permuted baseline, covariates, sampled transcriptome, and aging DE genes.

More detail in the supplementary section "***Aging Model***." This supplementary figure relates to **Fig. 7** and main text section "Assessing cell-type-specific transcriptomic and epigenetic changes in aging."

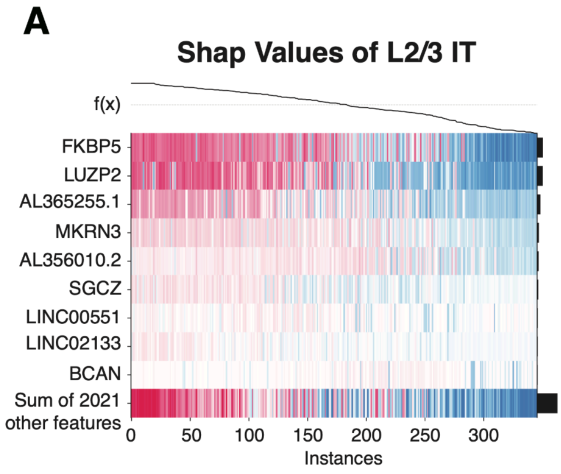


**Fig. S76. Aging prediction SHAP plot.**

**(A)** SHAP values of the aging prediction model for the top nine genes in L2/3 IT neurons across individuals. **(B)** SHAP values of the aging prediction model for the top nine genes in oligodendrocytes across individuals. The last row displays the sum of SHAP values for other assessed genes and covariates.

More detail in the supplementary section "***Aging Model*.**" This supplementary figure relates to **Fig. 7** and main text section "Assessing cell-type-specific transcriptomic and epigenetic changes in aging."

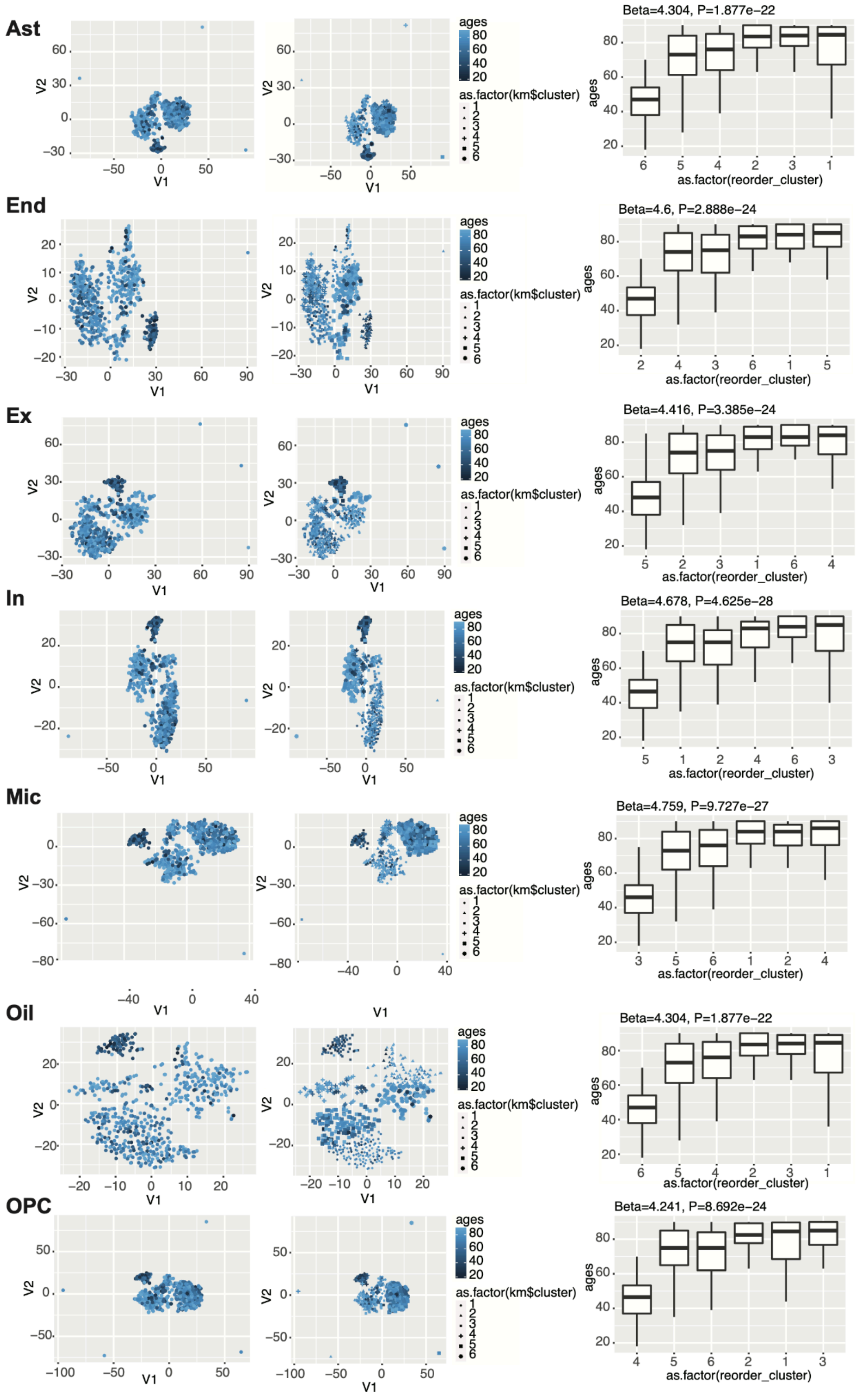


**Fig. S77. Clustering of cell-specific open chromatin regions reveals distinct clusters based on age.**

UMAPs of chromatin accessibility by age (**Fig. 7D**) for each cell type, as well as boxplots showing age distribution for each distinct UMAP cluster, are shown. These highlight open chromatin region signals that show patterns of clustering for each age group.

More detail in the supplementary section "***Aging Chromatin.***" This supplementary figure relates to **Fig. 7** and main text section "Assessing cell-type-specific transcriptomic and epigenetic changes in aging."

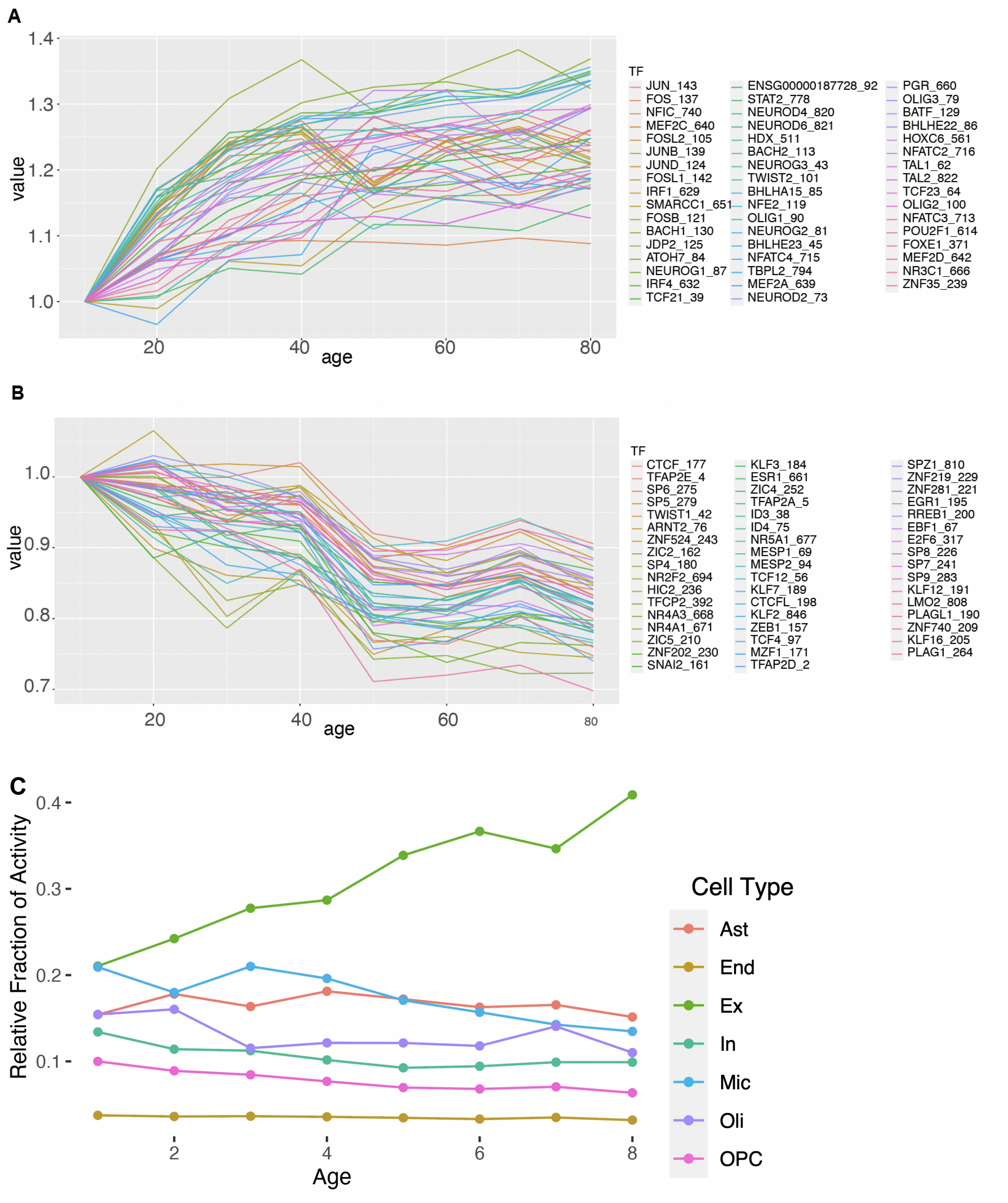
A
1.4
1.3
1.2
1.1
1.0
value
20
40
60
80
age
TF
JUN_143
FOS_137
NFIC_740
MEF2C_640
FOSL2_105
JUNB_139
JUND_124
FOSL1_142
IRF1_629
SMARCC1_651
FOSB_121
BACH1_130
JDP2_125
ATOH7_84
NEUROG1_87
IRF4_632
TCF21_39
ENSG00000187728_92
STAT2_778
NEUROD4_820
NEUROD6_821
HDX_511
BACH2_113
NEUROG3_43
TWIST2_101
BHLHA15_85
NFE2_119
OLIG1_90
NEUROG2_81
BHLHE23_45
NFATC4_715
TBPL2_794
MEF2A_639
NEUROD2_73
PGR_660
OLIG3_79
BATF_129
BHLHE22_86
HOXC6_561
NFATC2_716
TAL1_62
TAL2_822
TCF23_64
OLIG2_100
NFATC3_713
POU2F1_614
FOXE1_371
MEF2D_642
NR3C1_666
ZNF35_239
B
1.0
0.9
0.8
0.7
value
20
40
60
80
age
TF
CTCF_177
TFAP2E_4
SP6_275
SP5_279
TWIST1_42
ARNT2_76
ZNF524_243
ZIC2_162
SP4_180
NR2F2_694
HIC2_236
TFCP2_392
NR4A3_668
NR4A1_671
ZIC5_210
ZNF202_230
SNAI2_161
KLF3_184
ESR1_661
ZIC4_252
TFAP2A_5
ID3_38
ID4_75
NR5A1_677
MESP1_69
MESP2_94
TCF12_56
KLF7_189
CTCFL_198
KLF2_846
ZEB1_157
TCF4_97
MZF1_171
TFAP2D_2
SPZ1_810
ZNF219_229
ZNF281_221
EGR1_195
RREB1_200
EBF1_67
E2F6_317
SP8_226
SP7_241
SP9_283
KLF12_191
LMO2_808
PLAGL1_190
ZNF740_209
KLF16_205
PLAG1_264
C
0.4
0.3
0.2
0.1
Relative Fraction of Activity
2
4
6
8
Age
Cell Type
Ast
End
Ex
In
Mic
Oli
OPC

**D**

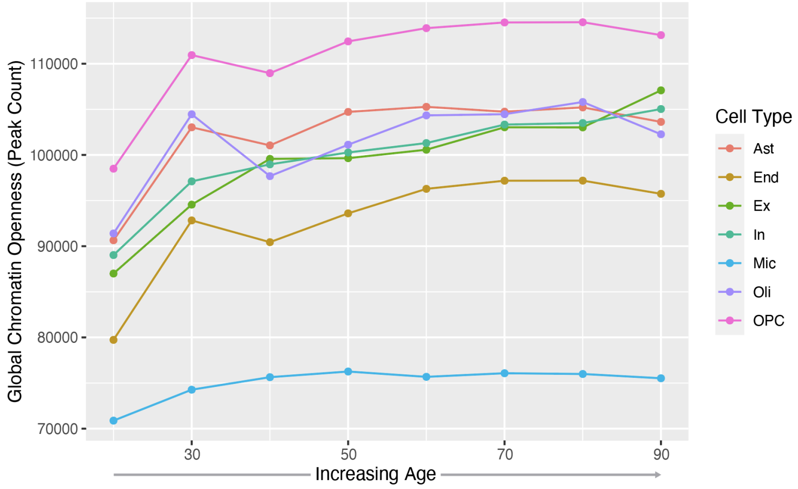


**Fig. S78. TF binding motif enrichment analysis for aging.**

**(A)** Line plot shows TF binding motif enrichment (top 50) in different age groups demonstrating increasing enrichment from young to old. **(B)** Line plot shows TF binding motif enrichment (top 50) showing decreasing enrichment from young to old. **(C)** Relative fraction of peaks represented in each cell type for ATAC-seq peaks across age. **(D)** Total number of open chromatin peaks per cell type across age.

More detail in the supplementary section "***Aging Chromatin***." This supplementary figure relates to **Fig. 7** and main text section "Assessing cell-type-specific transcriptomic and epigenetic changes in aging."

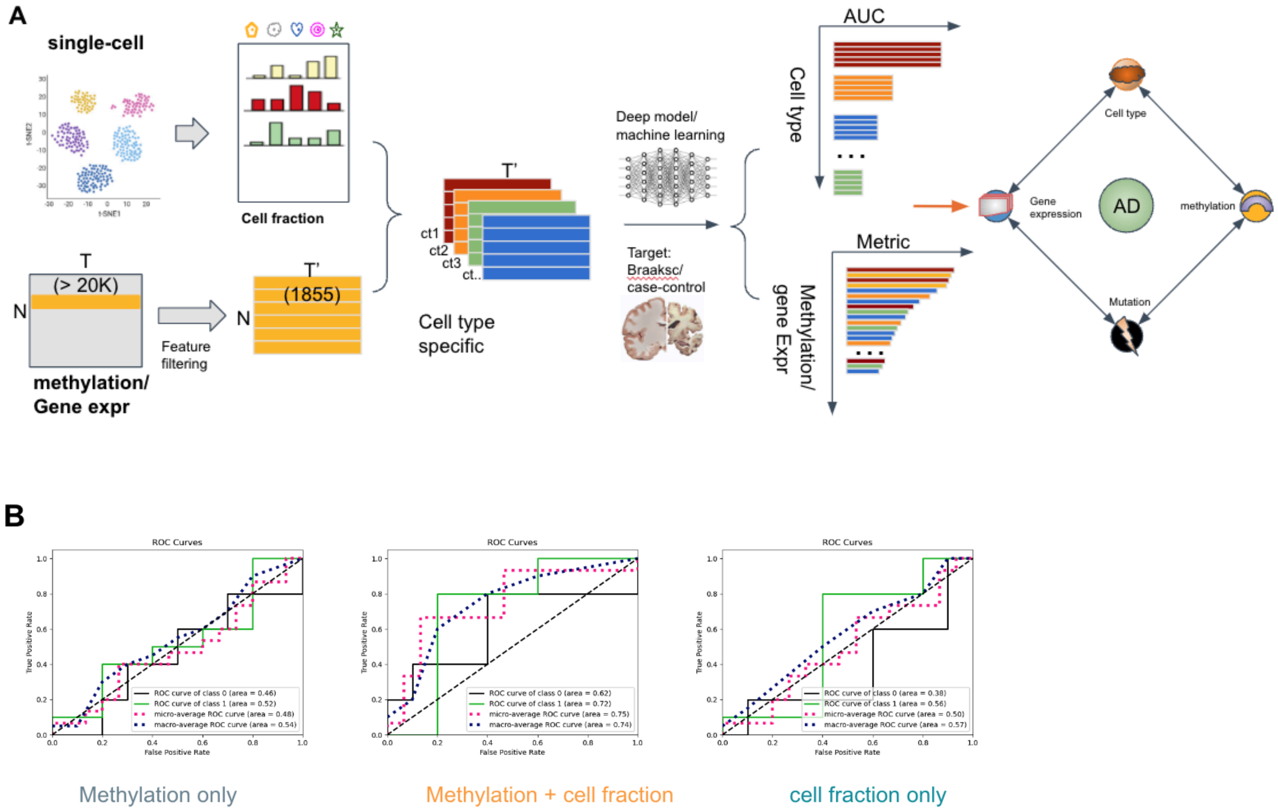


**Fig. S79. AD modeling.**

**(A)** Diagram provides an overview of the deep learning model for prediction of AD phenotype (Braak Score) from deconvolved cell fraction, methylation, and gene expression datasets. **(B)** The addition of cell fraction data helps improve the performance of cell-type signature (methylation) towards the prediction of AD phenotypes. The diagonal lines represent AUC scores (0.5) based on random guess.

More detail in the supplementary section "***AD Model***." This supplementary figure relates to **Fig. 7** and main text section "Assessing cell-type-specific transcriptomic and epigenetic changes in aging."

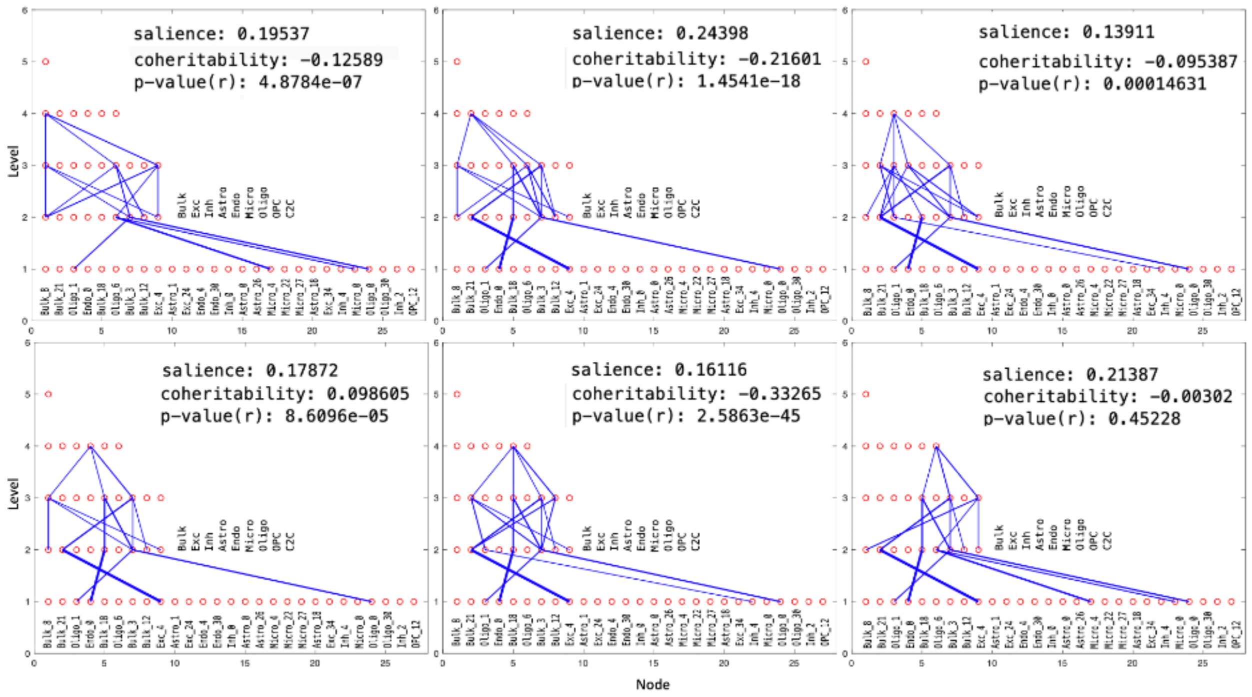


**Fig. S80. LNCTP-prioritized subgraphs for schizophrenia.**

Diagrams show prioritized consensus subgraphs of the schizophrenia LNCTP model, using the approach in Algorithm 1, where all links in the subgraphs appear in at least 4 out of 10 trained models (edge thickness corresponds to number of models containing link). Shown are the salience, coheritability, and p-value (Pearson correlation) of the intermediate phenotype corresponding to the activation of the upper-most node in each subgraph. Further annotated on levels 2 and 1 are the cell types and cell-type gene-module assignments of nodes, respectively. Gene modules are defined in the WGCNA analysis in (*100*). Graphs 1 and 2 in the upper row are those shown in **Fig. 8D** for schizophrenia prioritization.

More detail in the supplementary section "***LNCTP Interpretation***." This supplementary figure relates to **Fig. 8** and the main text section "Imputing gene expression and prioritizing disease genes across cell types with an integrative model."

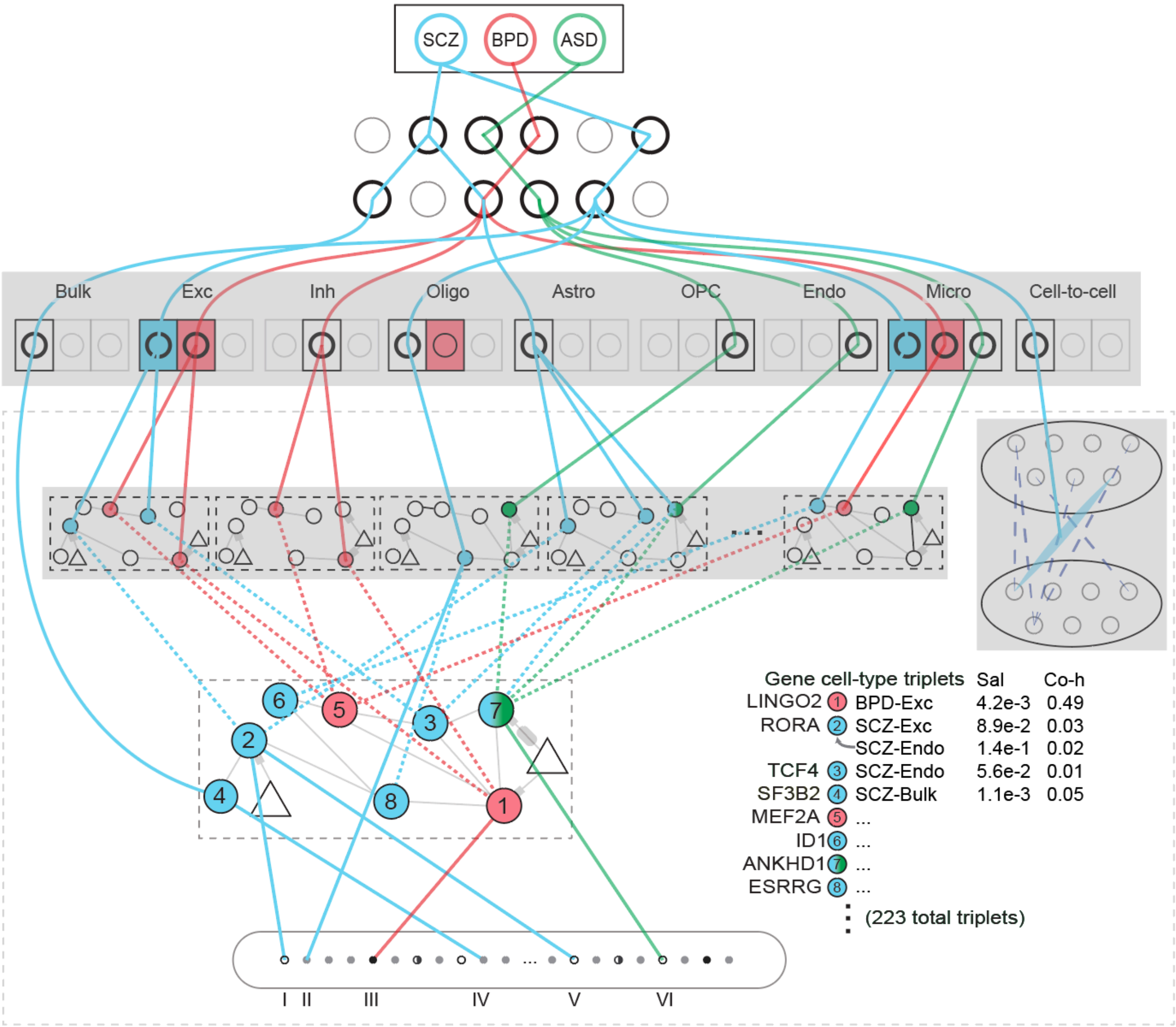


**Fig. S81. Comprehensive network connections and Information flow for cell-type-specific GRNs and bulk GRNs in LNCTP.**

This supplementary figure serves as an extension to **Fig. 8D**. The dotted lines represent the consistency potential links for specific genes, illustrating the flow of information from the bulk QTLs through the network. This flow traces from the bulk SNPs to cell-specific expression levels, culminating in high-level trait prediction. While this supplementary figure captures all connections, including the less important ones between cell-type-specific GRNs and bulk GRNs, the primary figure, which is a subgraph of this supplementary figure, includes only the strongest connections between individual cell types and bulk networks. We note that the highlighted genes here may be prioritized in the cell-type-specific networks and/or the bulk network (see also **Fig. 8E**). The bulk prioritized genes are thus based on the same LNCTP models as the other prioritized genes in our analyses; they are also based on the same training data (including both bulk and single-cell cohorts), and their salience arises through their combined effect across cell types.

More detail in the supplementary section "***LNCTP Interpretation*.**" This supplementary figure relates to **Fig. 8** and the main text section "Imputing gene expression and prioritizing disease genes across cell types with an integrative model."

# A

**SCZ**

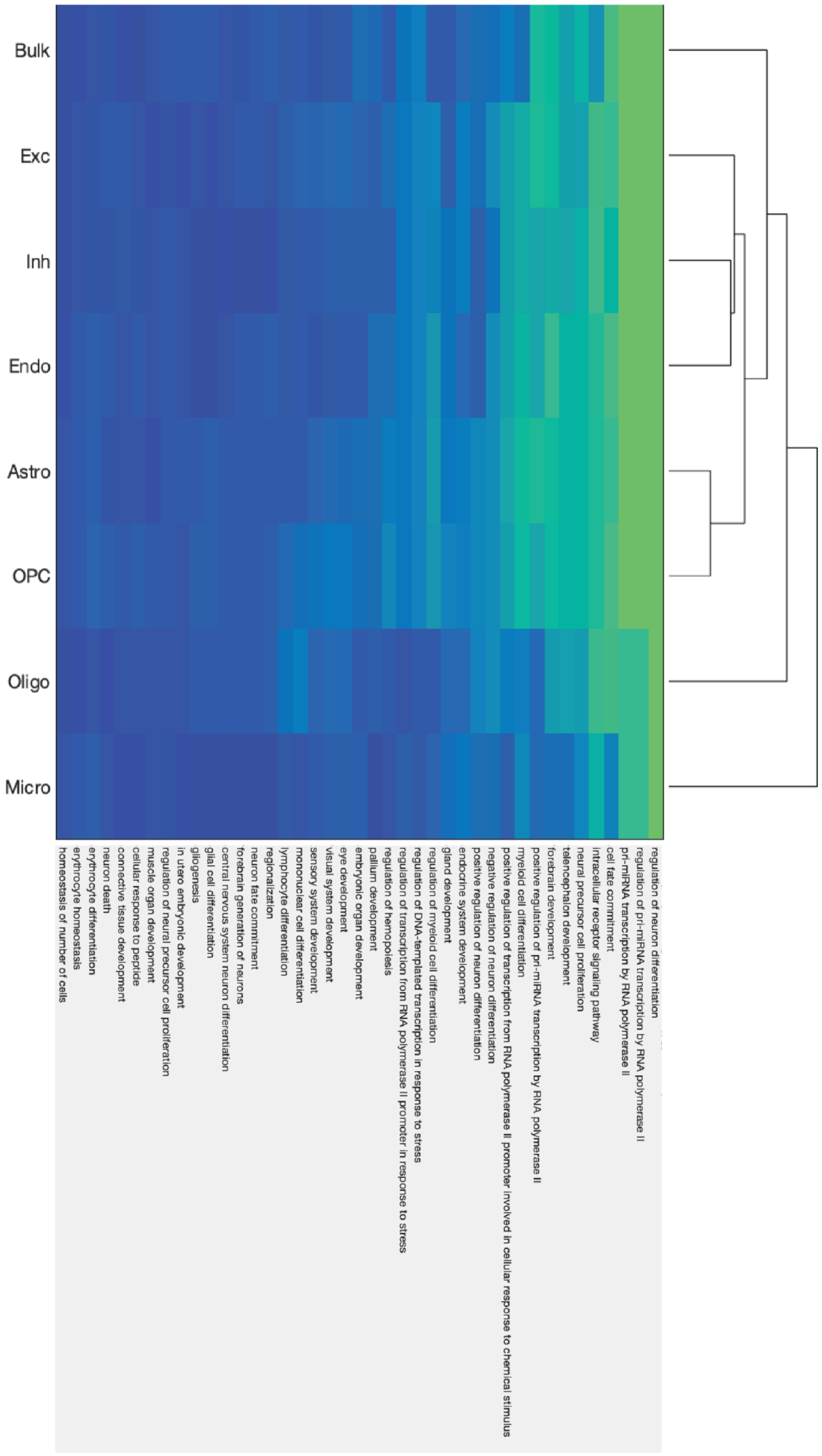

Bulk
Exc
Inh
Endo
Astro
OPC
Oligo
Micro
homeostasis of number of cells
erythrocyte homeostasis
erythrocyte differentiation
neuron death
connective tissue development
cellular response to peptide
muscle organ development
regulation of neural precursor cell proliferation
in utero embryonic development
gliogenesis
glial cell differentiation
central nervous system neuron differentiation
forebrain generation of neurons
neuron fate commitment
regionalization
lymphocyte differentiation
mononuclear cell differentiation
sensory system development
visual system development
eye development
embryonic organ development
pallium development
regulation of hemopoiesis
regulation of transcription from RNA polymerase II promoter in response to stress
regulation of DNA-templated transcription in response to stress
regulation of myeloid cell differentiation
gland development
endocrine system development
positive regulation of neuron differentiation
negative regulation of neuron differentiation
positive regulation of transcription from RNA polymerase II promoter involved in cellular response to chemical stimulus
myeloid cell differentiation
positive regulation of pri-miRNA transcription by RNA polymerase II
forebrain development
telencephalon development
neural precursor cell proliferation
intracellular receptor signaling pathway
cell fate commitment
pri-miRNA transcription by RNA polymerase II
regulation of pri-miRNA transcription by RNA polymerase II
regulation of neuron differentiation

B

**BPD**

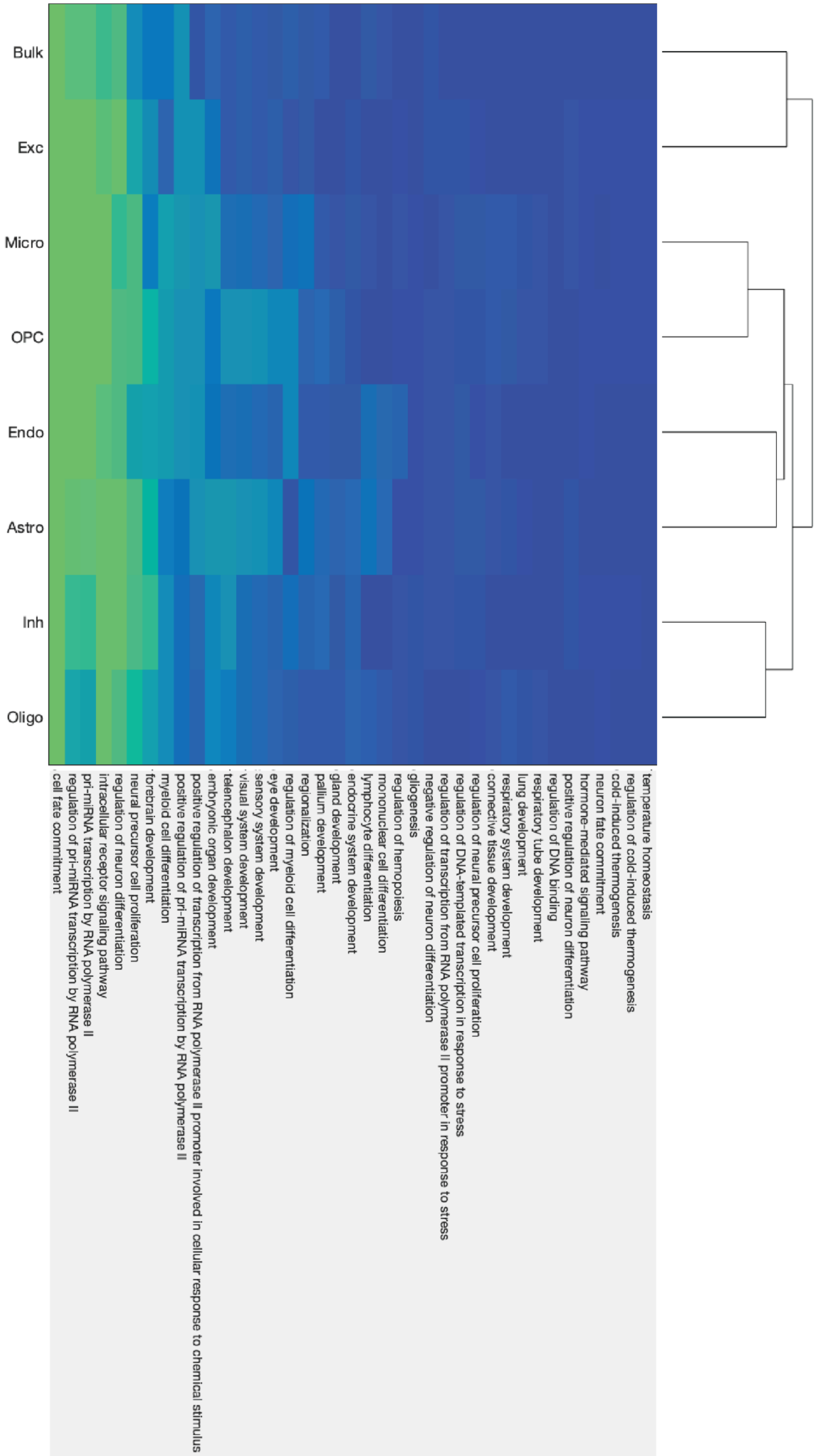

Bulk
Exc
Micro
OPC
Endo
Astro
Inh
Oligo
cell fate commitment
regulation of pri-miRNA transcription by RNA polymerase II
pri-miRNA transcription by RNA polymerase II
intracellular receptor signaling pathway
regulation of neuron differentiation
neural precursor cell proliferation
forebrain development
myeloid cell differentiation
positive regulation of pri-miRNA transcription by RNA polymerase II
positive regulation of transcription from RNA polymerase II promoter involved in cellular response to chemical stimulus
embryonic organ development
telencephalon development
visual system development
sensory system development
eye development
regulation of myeloid cell differentiation
regionalization
pallium development
gland development
endocrine system development
lymphocyte differentiation
mononuclear cell differentiation
regulation of hemopoiesis
gliogenesis
negative regulation of neuron differentiation
regulation of transcription from RNA polymerase II promoter in response to stress
regulation of DNA-templated transcription in response to stress
regulation of neural precursor cell proliferation
connective tissue development
respiratory system development
lung development
respiratory tube development
regulation of DNA binding
positive regulation of neuron differentiation
hormone-mediated signaling pathway
neuron fate commitment
cold-induced thermogenesis
regulation of cold-induced thermogenesis
temperature homeostasis

**C**

**ASD**

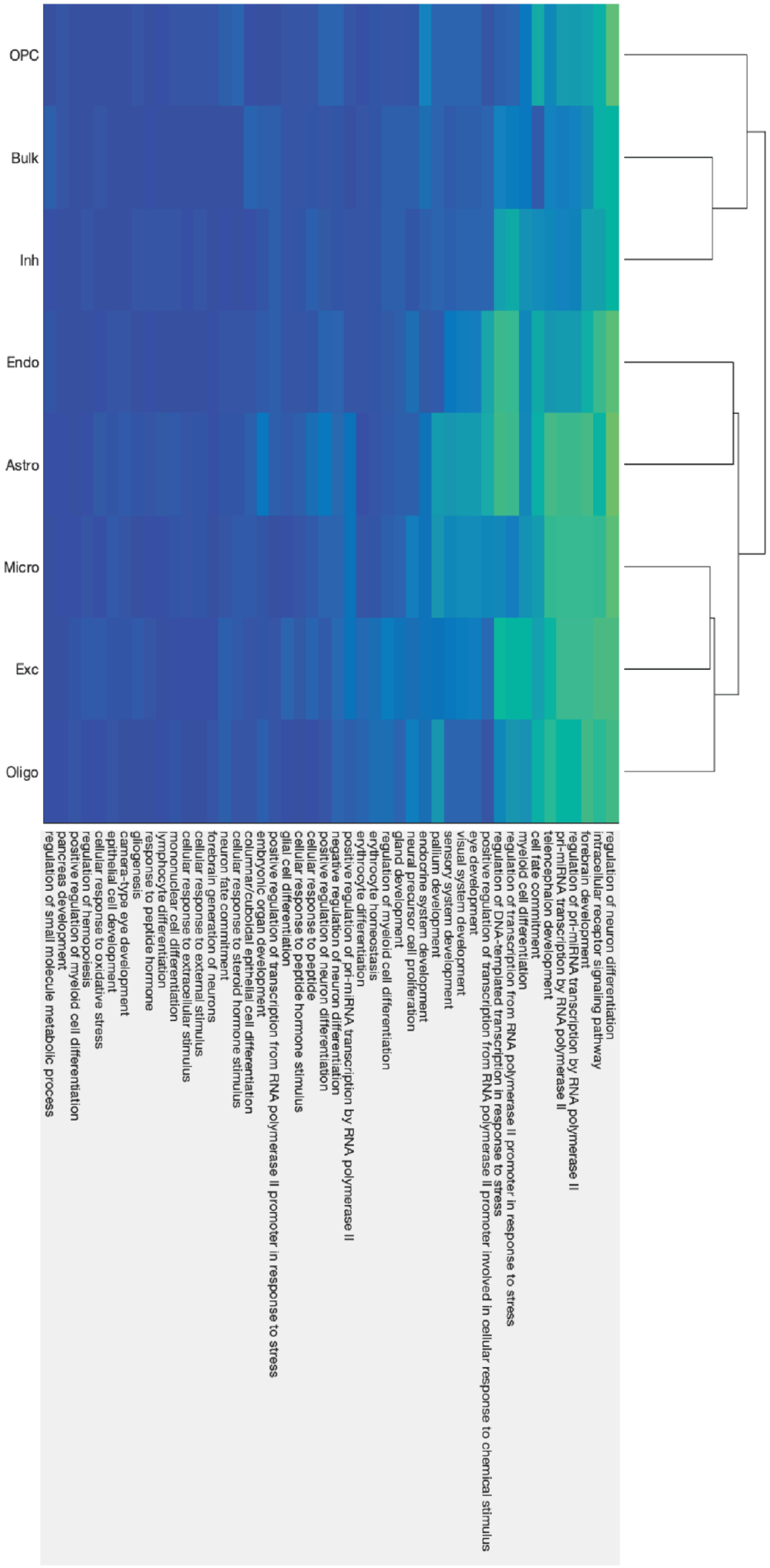

OPC
Bulk
Inh
Endo
Astro
Micro
Exc
Oligo
regulation of neuron differentiation
intracellular receptor signaling pathway
forebrain development
regulation of pri-miRNA transcription by RNA polymerase II
pri-miRNA transcription by RNA polymerase II
telencephalon development
cell fate commitment
myeloid cell differentiation
regulation of transcription from RNA polymerase II promoter in response to stress
regulation of DNA-templated transcription in response to stress
positive regulation of transcription from RNA polymerase II promoter involved in cellular response to chemical stimulus
eye development
visual system development
sensory system development
pallium development
endocrine system development
neural precursor cell proliferation
gland development
regulation of myeloid cell differentiation
erythrocyte homeostasis
erythrocyte differentiation
positive regulation of pri-miRNA transcription by RNA polymerase II
negative regulation of neuron differentiation
positive regulation of neuron differentiation
cellular response to peptide
cellular response to peptide hormone stimulus
glial cell differentiation
positive regulation of transcription from RNA polymerase II promoter in response to stress
embryonic organ development
columnar/cuboidal epithelial cell differentiation
cellular response to steroid hormone stimulus
neuron fate commitment
forebrain generation of neurons
cellular response to external stimulus
cellular response to extracellular stimulus
mononuclear cell differentiation
lymphocyte differentiation
response to peptide hormone
gliogenesis
camera-type eye development
epithelial cell development
cellular response to oxidative stress
regulation of hemopoiesis
positive regulation of myeloid cell differentiation
pancreas development
regulation of small molecule metabolic process

**Fig. S82. LNCTP enrichment of GO and KEGG terms for prioritized genes.**

Heatmaps show -log(p) values (green and blue indicate high and low, respectively) for the enrichment of GO biological process terms and KEGG pathways in prioritized gene sets for LNCTP models across cell types for **(A)** schizophrenia, **(B)** bipolar disorder, and **(C)** ASD.

More detail in the supplementary section "***LNCTP Interpretation*.**" This supplementary figure relates to **Fig. 8** and the main text section "Imputing gene expression and prioritizing disease genes across cell types with an integrative model."

A

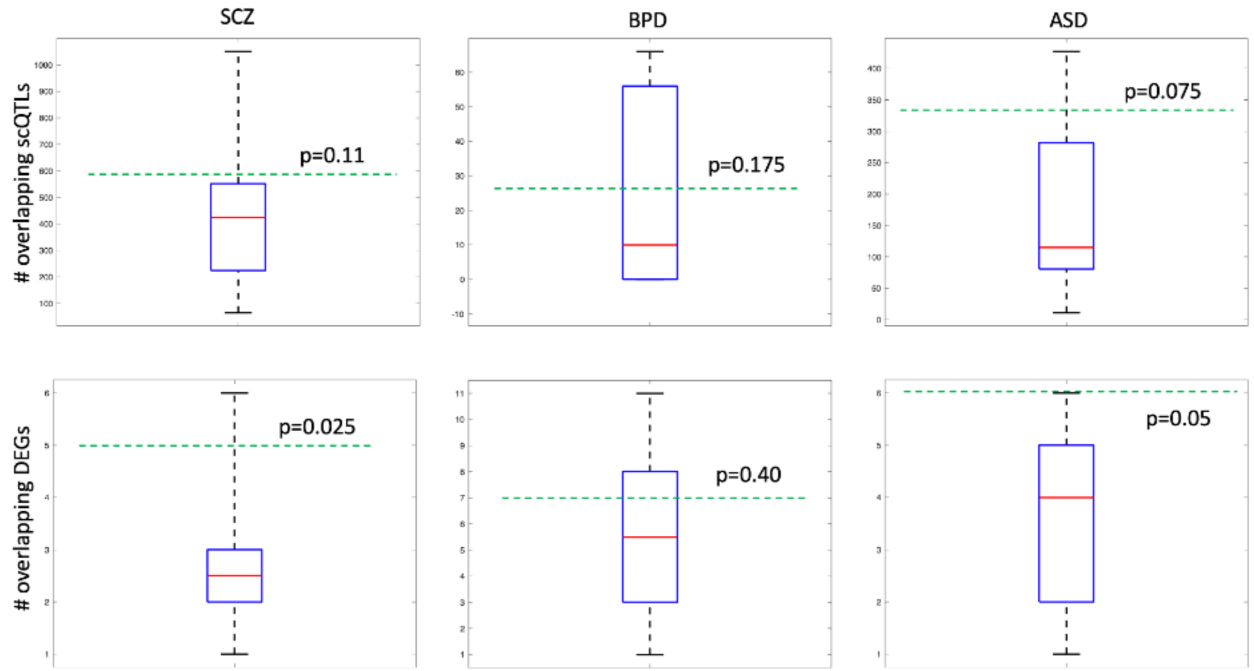


B

| Intersection (LNCTP/DEGs) | LNCTP unique genes | | DEGs unique genes |
|---|---|---|---|
| BACH2 | AKAP6 | NR3C2 | A2M-AS1 |
| BCL11B | ARID5B | NRF1 | AAK1 |
| BCL6 | ARNT2 | OLIG1 | AAMDC |
| FUBP1 | BRINP2 | OLIG2 | AARSD1 |
| HMBOX1 | CREM | POU2F1 | ABCA1 |
| INPP4B | CUX1 | RBM26 | ABCA4 |
| KMT2E | DGKZ | RC3H1 | ABCA7 |
| MEIS2 | EPAS1 | RFTN2 | ABCC12 |
| MSANTD2 | GLIS3 | RFX7 | ABCC2 |
| NR4A3 | HIF1A | SF3B2 | ABCC3 |
| NSD3 | HIVEP2 | SOX9 | ABCD4 |
| PBX1 | HSPA9 | SREBF2 | ABHD12B |
| PHF21A | HSPD1 | SRPK2 | ABHD17C |
| PTPRK | HSPE1 | STAG1 | ABHD2 |
| RORA | KLC1 | STAT5B | ABHD3 |
| TEAD1 | MEF2A | TCF4 | ABI3BP |
| ZFX | NDUFA13 | YBX1 | ⋮ |
| | NFATC3 | ZEB1 | |
| | NR3C1 | ZEB2 | |

**Fig. S83. LNCTP-prioritized genes overlap with sc-eQTLs and DE genes.**
**(A)** Permutation tests for significant overlap of LNCTP prioritized genes with scQTLs (top) and DE genes (bottom) per disorder (one-sample, two-tailed t-tests). Green dotted lines show the number of overlapping scQTLs or DE genes, and box plots show distribution of overlap counts when cell-type labels are permuted 40 times. p-values shown are the fraction of the latter that are greater than the former. In all tests, scQTLs are selected with adjusted p<0.05 (see supplementary section 4.1 for statistical testing), DE genes are selected with adjusted p<0.2, and LTCTP prioritized genes are selected with p<0.1. **(B)** shows the intersection of salient genes with DE genes, and uniquely occurring LNCTP and DE genes for schizophrenia only (thresholds are the same as above).

More detail in the supplementary section "***LNCTP Interpretation*.**" This supplementary figure relates to **Fig. 8** and main text section "Imputing gene expression and prioritizing disease genes across cell types with an integrative model."

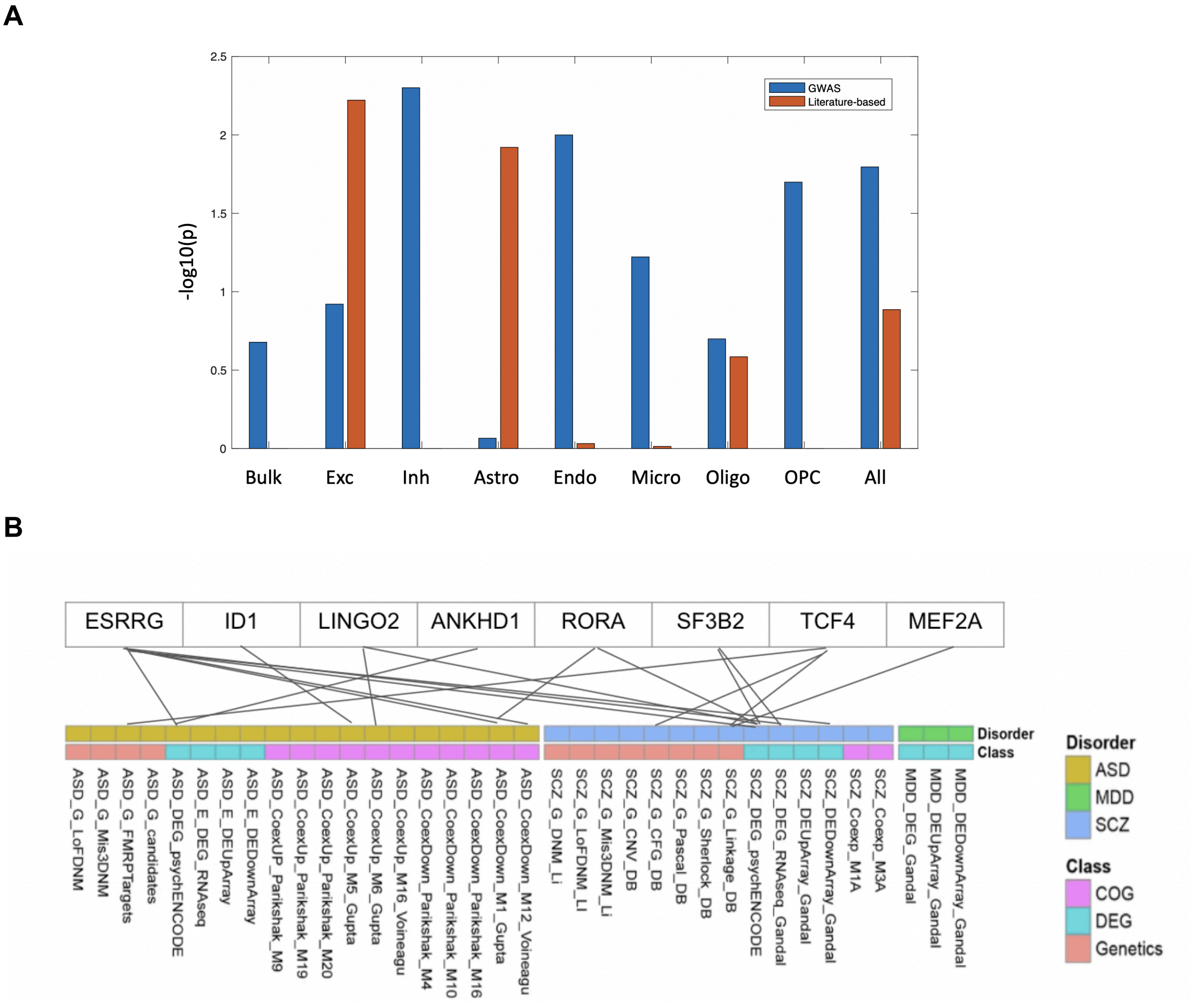


**Fig. S84. Prior literature support for LNCTP-prioritized genes.**

**(A)** Results of hypergeometric tests to compare the enrichment of prioritized genes in each cell type (as well as bulk and all prioritized genes combined) for fine-mapped GWAS genes in schizophrenia and ASD datasets (blue) and prior literature support (red). **(B)** Graph showing GWAS and prior literature support for eight key genes towards disease relevance (ASD, schizophrenia, and MDD) in a curated set of references (see **data S33** for abbreviations and citations). COG: co-expressed gene; DEG: DE gene; Genetics: genetic evidence for association.

More detail in the supplementary section "***LNCTP Validation.***" This supplementary figure relates to **Fig. 8** and main text section "Imputing gene expression and prioritizing disease genes across cell types with an integrative model."

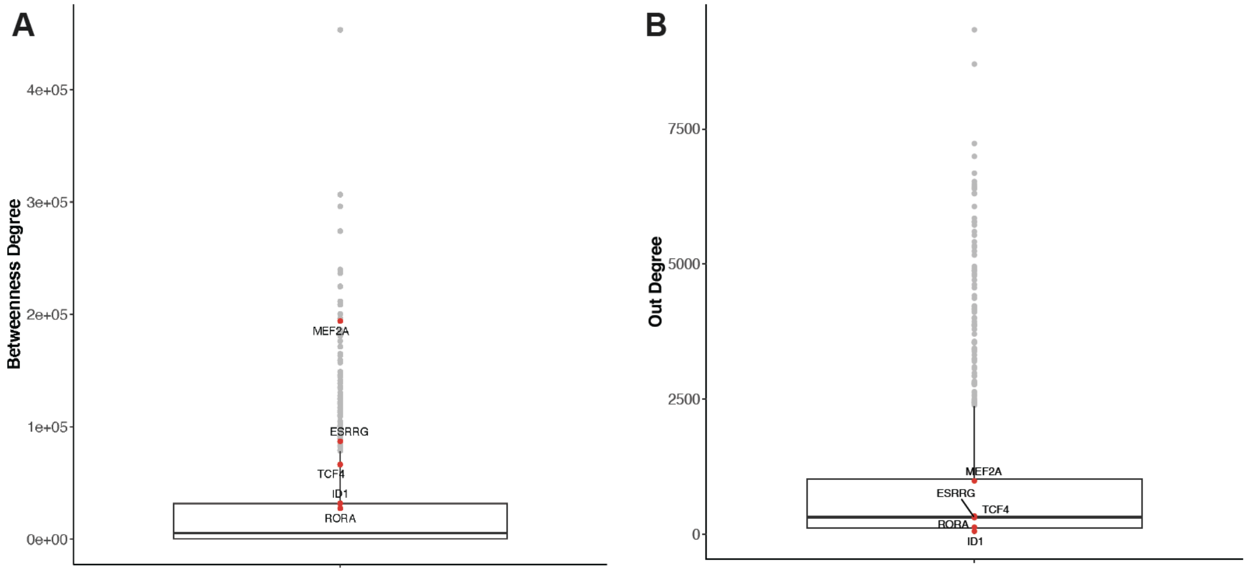


**Fig. S85. Interpretation of LNCTP-prioritized genes.**

Boxplots show the betweenness degree **(A)** and out-degree **(B)** distributions of the TFs that exist in the bulk GRN. Values for the eight key genes are highlighted in red.

Related to the supplementary section "***LNCTP Validation*.**" This supplementary figure relates to **Fig. 8** and the main text section "Imputing gene expression and prioritizing disease genes across cell types with an integrative model."

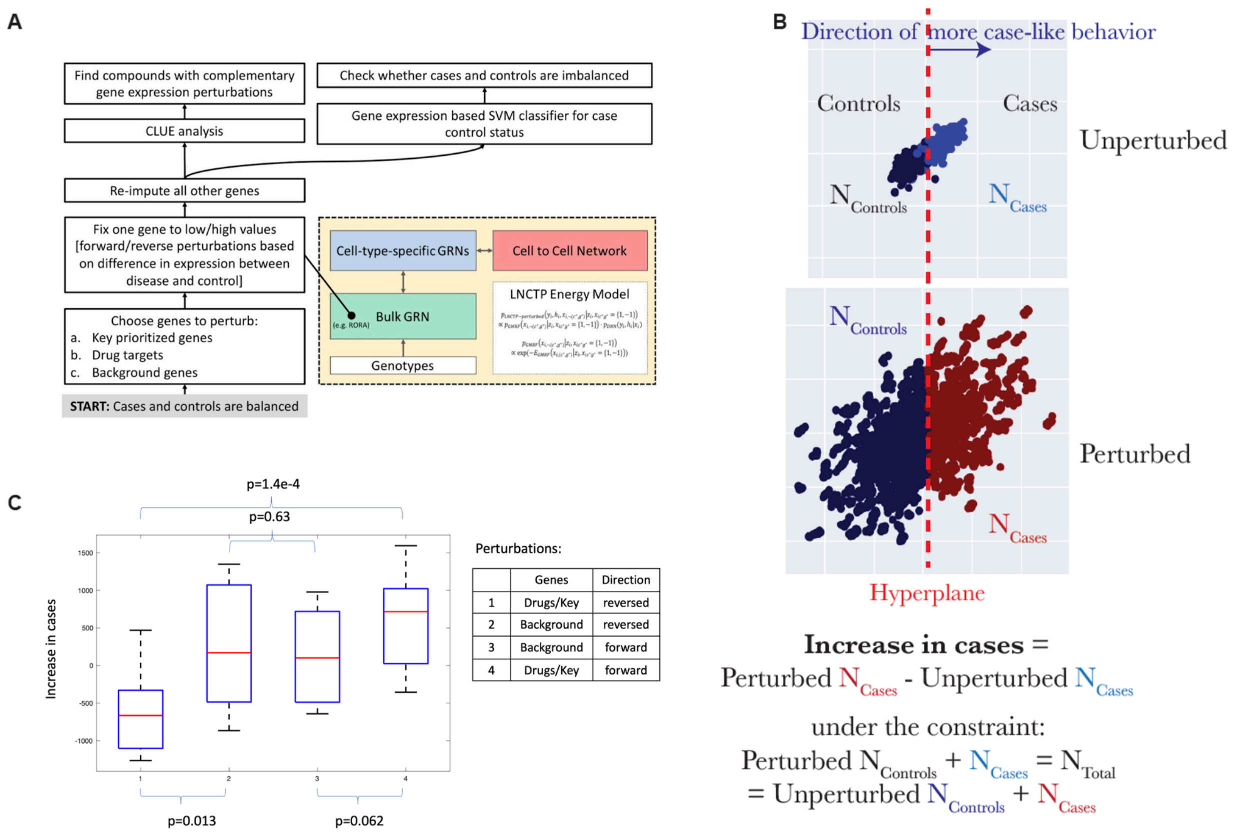


**Fig. S86. In silico perturbation analysis of trained LNCTP models.**

**(A)** Flowchart describing the gene-expression perturbation and downstream analyses. The flowchart represents two branches, one for the SVC-based analysis and the other for the CLUE-based drug analysis. **(B)** A simplified schematic explaining how the SVC analysis helps define "case-like" behavior and how the "Increase in Cases" metric is defined. **(C)** Boxplots of the "Increase in Cases" distributions as a function of the gene sets considered: Drug+Key genes with the forward ("case-like") perturbation; Drug+Key genes with the reverse ("control-like") perturbation; Background genes with the forward ("case-like") perturbation; Background genes with the reverse ("control-like") perturbation. P-values calculated using one-tailed two-sample t-tests,

More detail in the supplementary section "***LNCTP Validation***." This supplementary figure relates to **Fig. 8** and the main text section "Imputing gene expression and prioritizing disease genes across cell types with an integrative model."

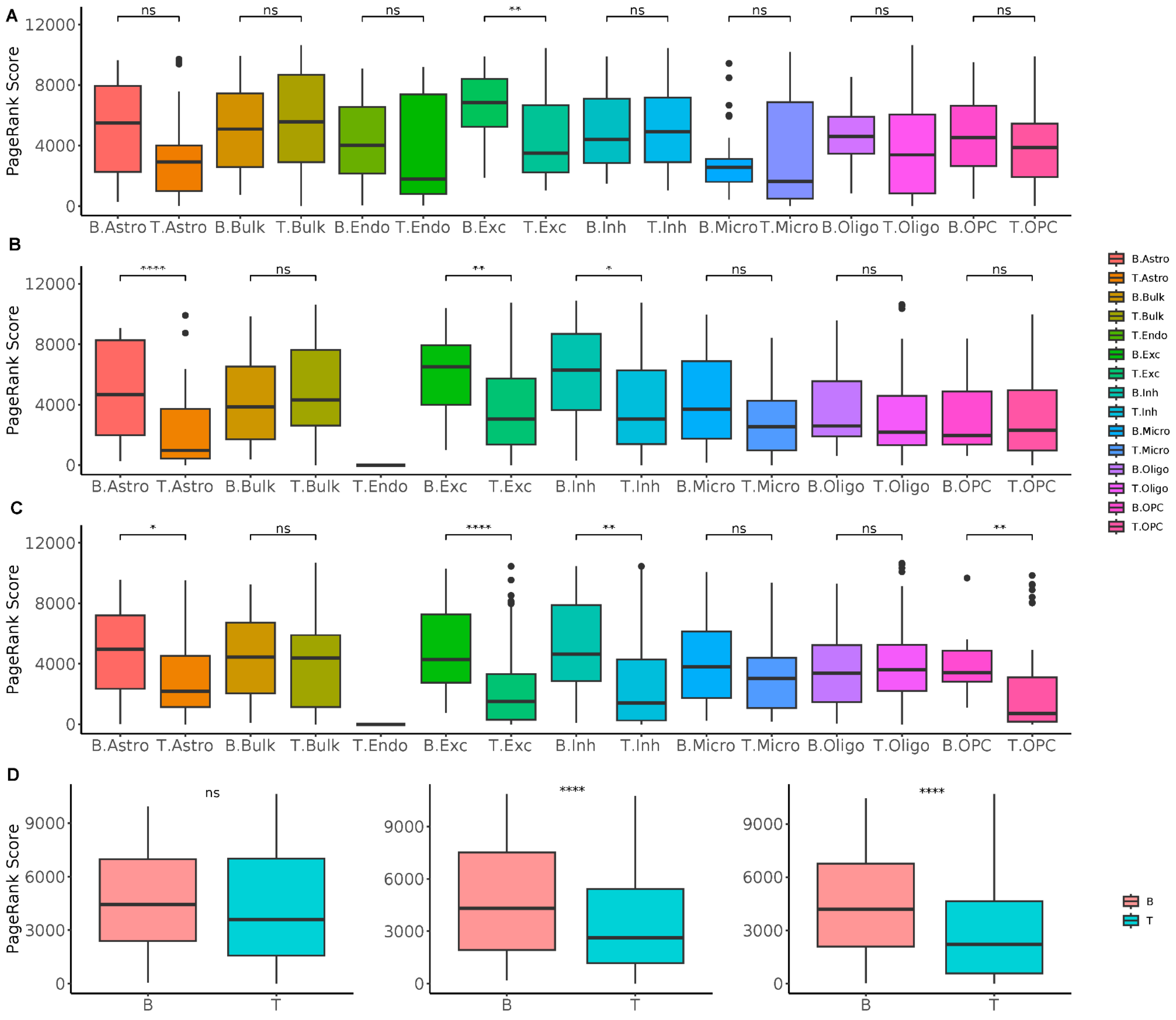


**Fig. S87. Network effects of LNCTP perturbations.**

**(A-C)** Plots for *MEF2A*, *RORA*, and *TCF4* show the PageRank score distribution of bottom (B) and top ranked genes (T) for each cell type with respect to perturbed genes in the GRNs. Imputed genes are ranked by their magnitude of imputed gene expression change. **(D)** The aggregated distribution of the PageRank scores for bottom and top-ranked genes. Left to right distributions are for the *MEF2A*, *RORA*, and *TCF4* perturbed genes.

More detail in the supplementary section "***LNCTP Validation***." This supplementary figure relates to **Fig. 8** and the main text section "Imputing gene expression and prioritizing disease genes across cell types with an integrative model."

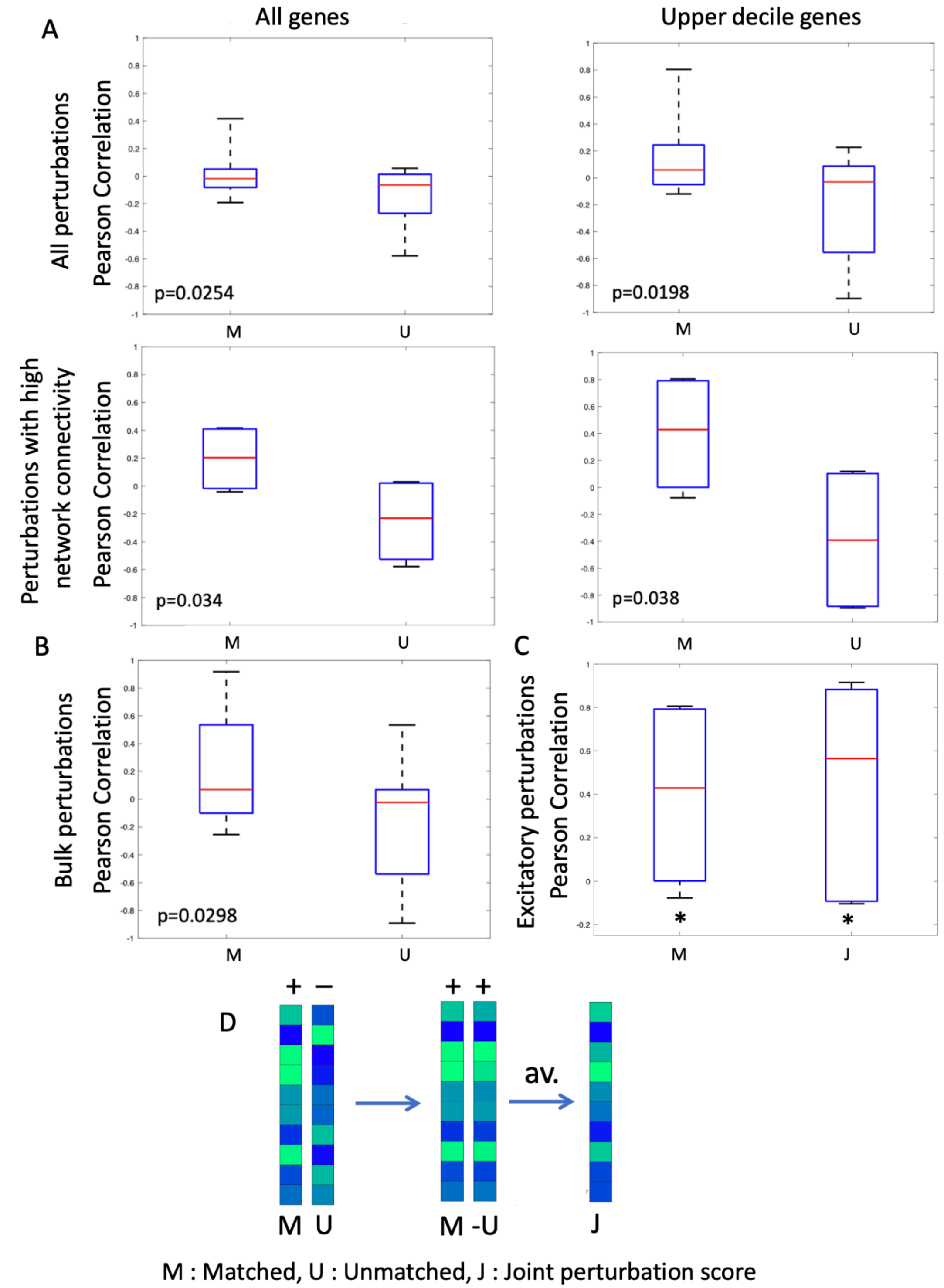


**Fig. S88. Comparing LNCTP and CRISPR perturbations.**

Graphs compare the Pearson correlation values for LNCTP and CRISPR-interference/ CRISPR-activation perturbation vectors for 10 genes, where the perturbation directions are matched vs. unmatched (LNCTP z-score changes are correlated with CRISPR fold-change vectors for all genes other than the perturbed gene). (**A**) shows the correlations resulting from perturbations in the excitatory neuron GRN. Left vs right shows all genes vs. the upper decile of genes according to the absolute LNCTP z-score changes; top vs bottom shows all perturbations vs perturbations whose target gene has at least 10 connected genes in the GRN (5 perturbations). (**B**) shows the correlations resulting from perturbations in the bulk GRN using the upper decile genes. (**C**) compares Pearson correlations of the matched and joint perturbation scores with the CRISPR perturbations for the upper decile genes; calculated as $corr(J, X)$, where $J = 0.5(M - U)$ , and $M, U, X, J$ are the LNCTP matched, unmatched and CRISPR experimental perturbation vectors and joint perturbation score respectively, shown schematically in (**D**). The bottom-right graph in (**A**) corresponds to **Fig. 8F.** p-values calculated using one-tailed paired and one-sample t-tests, in (**A-B**) and (**C**) respectively, *p<0.05.

More detail in the supplementary section "***Independent CRISPR validation of LNCTP*.**" This supplementary figure relates to **Fig. 8** and the main text section "Imputing gene expression and prioritizing disease genes across cell types with an integrative model."

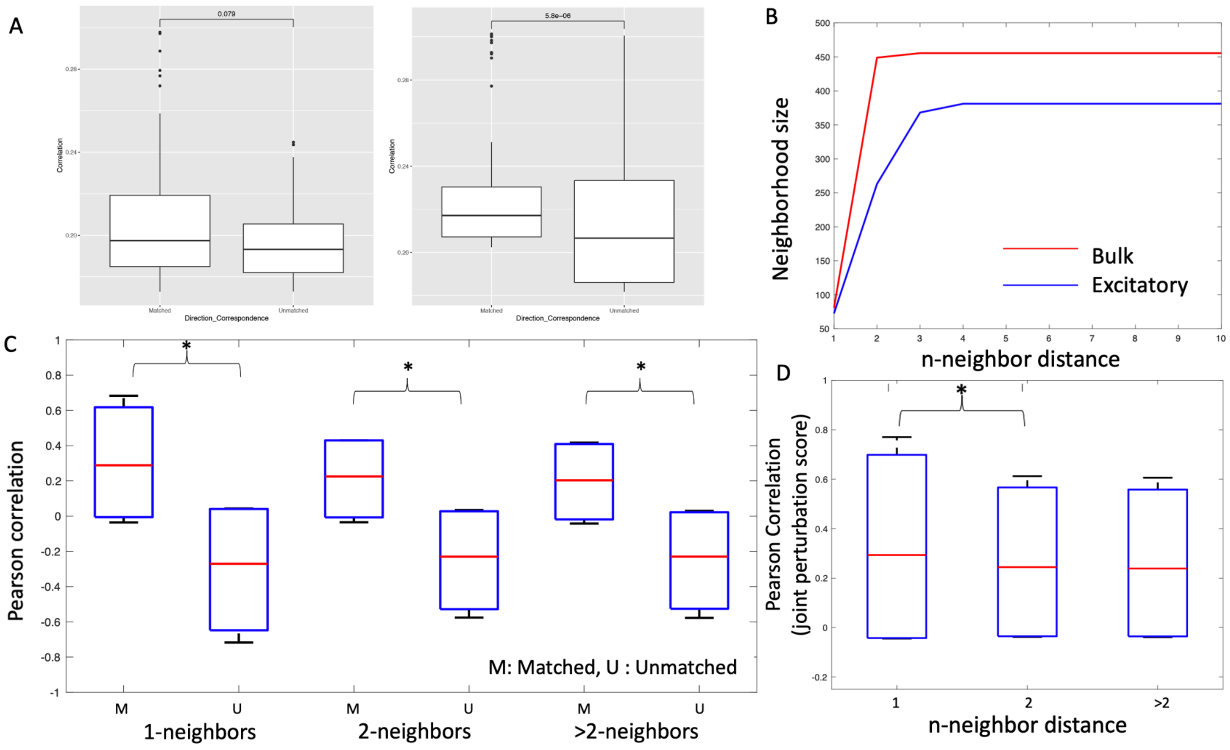


Fig. S89. Effects of genetic background and neighborhood size on LNCTP perturbation effects.

**(A)** Box plots compare Pearson's correlations between LNCTP and CRISPR perturbations analogously to **fig. S88**, but comparing LNCTP perturbations across all individuals. Here, correlations are calculated using the signed z-score differences in both the CRISPR and LNCTP perturbations. Left vs right shows the correlations using LNCTP perturbations in the bulk and excitatory GRNs respectively (perturbed genes have at least 10 connected genes in the GRN). **(B)** shows how neighborhood size varies with distance in bulk and excitatory GRNs. **(C)** plots the Pearson correlations between LNCTP and CRISPR perturbations analogously to **fig. S88** in the excitatory GRN, restricted to the genes within neighborhoods of varying size of the perturbed target gene, and **(D)** shows joint perturbation scores for the same neighborhood sizes. p-values are calculated using one-tailed Wilcoxon rank-sum tests **(A)** and t-tests **(B-D),** *$p<0.05$.

More detail in the supplementary section "***Independent CRISPR validation of LNCTP*.**" This supplementary figure relates to **Fig. 8** and the main text section "Imputing gene expression and prioritizing disease genes across cell types with an integrative model."

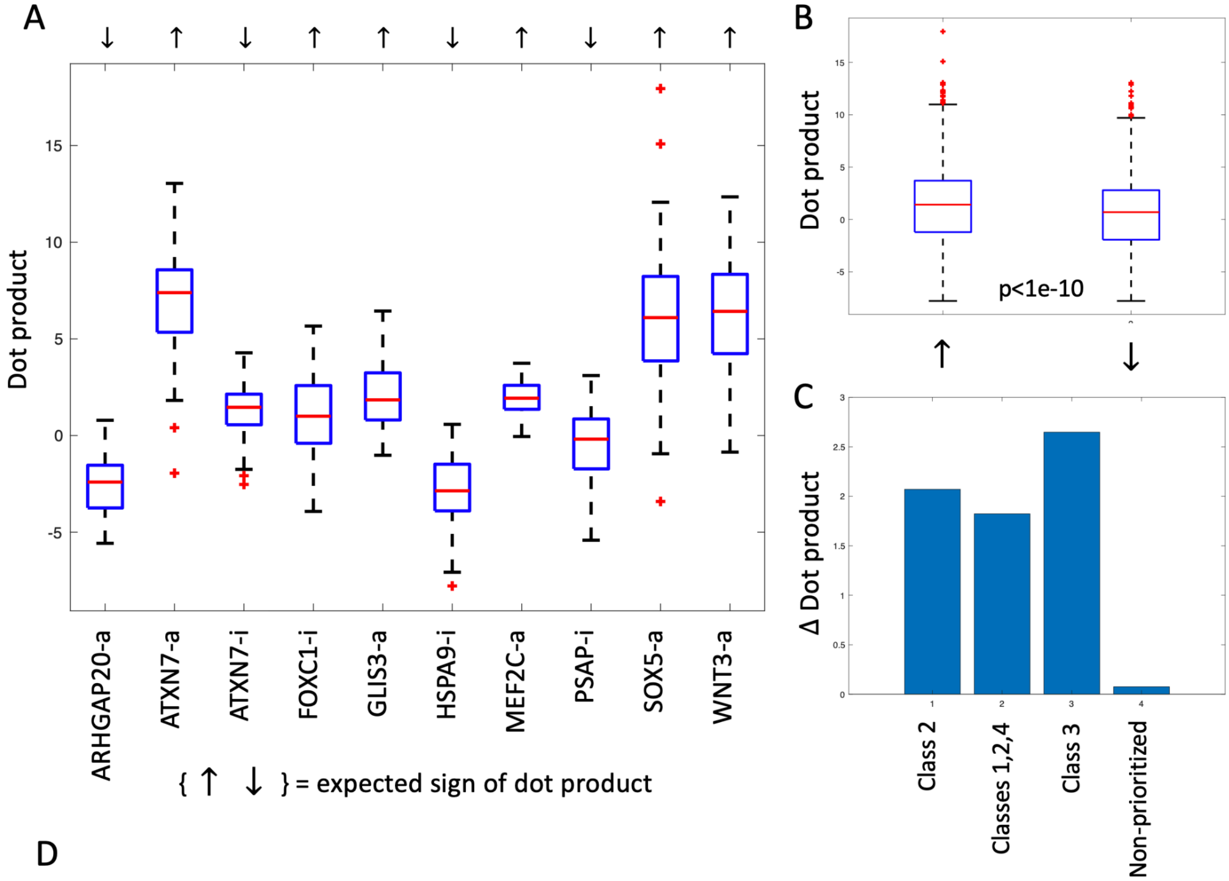


D

| Prioritization classes | CRISPR Gene/Perturbations |
|---|---|
| Class 1 | GLIS3-a, HSPA9-i, MEF2C-a |
| Class 2 | PSAP-i, WNT3-a |
| Class 3 | ARHGAP20-a |
| Non-prioritized | ATXN7-a, ATXN7-i, FOXC1-i, SOX5-a |

**Fig. S90. Comparing case/control-like effects of CRISPR perturbations for LNCTP prioritized genes vs. non-prioritized genes.**

**(A)** Plot shows the distribution of dot products of the CRISPR fold-change vectors for 10 gene-perturbation pairs with 100 unit normal SVC vectors, reflecting the direction of maximum discrimination of case/control status, along with the expected sign of the dot product (via DE analysis). Plots compare **(B)** the distribution of dot products for those pairs whose expected change is positive vs. negative, and **(C)** the mean z-score change for positive and negative pairs in different classes of genes. **(D)** Table summarizes the CRISPR gene-perturbation pairs belonging to each prioritization class (Classes 1, 2, and 3 correspond to LNCTP prioritized and not DE genes, cell-to-cell network prioritized, and LNCTP and DE gene prioritized without extensive prior support, respectively; see section 8.7 for further details). p-values calculated using one-tailed t-tests.

More detail in the supplementary section "***Independent CRISPR validation of LNCTP*.**" This supplementary figure relates to **Fig. 8** and the main text section "Imputing gene expression and prioritizing disease genes across cell types with an integrative model."

### Supplementary Tables

**Table S1. Summary of the snATAC-Seq/snMultiome processing and filtering for each of the cohorts with snATAC data.**

| **Study** | **No. of cells after initial processing** | **No. of cells after QC filtering** |
|---|---:|---:|
| UCLA-ASD snATAC-Seq | 88,677 | 66,946 |
| Girgenti snMultiome | 295,434 | 125,991 |
| MultioneBrain snMultiome | 181,374 | 80,565 |
| **Total snATAC-Seq** | **565,485** | **273,502** |

More detail in the supplementary section "***snMultiome Dataset.***" This supplementary table relates to **Fig. 2** and main text section "Determining regulatory elements for cell types from snATAC-seq."

**Table S2. Dataset-by-dataset numbers of cells that remain after important steps in the snRNA-seq processing.**

| Study | No. of cells after initial processing and QC | No. of cells after cluster-based filtering of unannotated cells | No. of cells after Seurat-based annotation and reconciliation of the two annotations |
|---|---|---|---|
| SZBDMulti-Seq | 616,032 | 605,360 | 484,249 |
| CMC | 519,887 | 505,442 | 502,021 |
| UCLA-ASD | 704,548 | 509,101 | 448,524 |
| IsoHuB | 45,388 | 30,270 | 29,675 |
| DevBrain-snRNAseq | 131,237 | 108,708 | 102,936 |
| PTSDBrainomics | 226,099 | 200,427 | 198,572 |
| LIBD | 93,165 | 58,659 | 52,214 |
| MultiomeBrain-DLPFC | 160,439 | 140,361 | 134,645 |
| Velmeshev | 75,409 | 63,925 | 63,635 |
| AMP-AD_ROSMAP | 70,594 | 63,962 | 63,228 |
| Ma-Sestan | 247,415 | 206,269 | 201,574 |
| Girgenti-snMultiome | 311,200 | 291,074 | 276,018 |
| **Total snRNA-Seq** | **3,201,413** | **2,783,558** | **2,557,291** |

Table shows (1) numbers of cells after the initial processing and QC filtering; (2) numbers of cells remaining after the removal of cells that were unannotated in the cluster-based annotation step; and (3) numbers of cells remaining after cell-by-cell annotation using the Seurat label transfer pipeline viausing the BICCN and Ma-Sestan schemes, and subsequent reconciliation between the schemes. The last column represents the final numbers of cells used in the downstream analyses.

More detail in the supplementary section "***snRNA-seq Processing.***" This supplementary table relates to **Fig. 1** and main text section "Constructing a single-cell genomic resource for 388 individuals."

**Table S3. Cell types and their full titles and abbreviations.**

| Class | Cell Types | Expanded title (if applicable) | Abbreviation |
|---|---|---|---|
| Excitatory | L2/3 IT | Layer 2/3 Intratelencephalic projecting | **L2/3 IT** |
| | L4 IT | Layer 4 Intratelencephalic projecting | **L4 IT** |
| | L5 IT | Layer 5 Intratelencephalic projecting | **L5 IT** |
| | L6 IT | Layer 6 Intratelencephalic projecting | **L6 IT** |
| | L6 IT Car3 | Layer 6 Intratelencephalic projecting Car3 | **L6 IT Car3** |
| | L5 ET | Layer 5 Extratelencephalic projecting | **L5 ET** |
| | L5/6 NP | Layer 5/6 Near-projecting | **L5/6 NP** |
| | L6b | Layer 6b | **L6b** |
| | L6 CT | Layer 6 Corticothalamic projecting | **L6 CT** |
| Inhibitory | SST | | **SST** |
| | SST CHODL | | **SST CHODL** |
| | PVALB | | **PVALB** |
| | Chandelier | | **Chlr** |
| | LAMP5 LHX6 | | **LAMP5 LHX6** |
| | LAMP5 | | **LAMP5** |
| | SNCG | | **SNCG** |
| | VIP | | **VIP** |
| | PAX6 | | **PAX6** |
| Non-neuronal | Astro | Astrocytes | **Ast** |
| | Oligo | Oligodendrocytes | **Oli** |
| | OPC | Oligodendrocyte Precursor Cells | **OPC** |
| | Micro | Microglia | **Mic** |
| | Endo | Endothelial cells | **End** |
| | VLMC | Vascular Leptomeningeal Cells | **VLMC** |
| | PC | Pericytes | **PC** |
| | SMC | Smooth Muscle Cells | **SMC** |
| | Immune | Immune cells | **Imm** |
| | RB | Red Blood lineage cells | **RB** |

Related to the supplementary section "***snRNA-seq Processing*.**" This supplementary table relates to **Fig. 1** and main text section "Constructing a single-cell genomic resource for 388 individuals."

**Table S4. Subclasses/cell types and the color schemes used in the manuscript (including hex codes).**

| Class color scheme for Figs. 1-4, 5-8 | | Cell Type | | Subclass color scheme for Figs. 1-4, 5-8 | | | Color scheme for Fig. 6 | |
|---|---|---|---|---|---|---|---|---|
| | | | | **Hex code** | **Color** | | **Hex code** | **Color** |
| **Exc** | | L2/3 IT | | #078d46 | | | #3954a4 | |
| #078d46 | | L4 IT | | #0073ab | | | #384fa1 | |
| | | L5 IT | | #fbdbe6 | | | #1562a0 | |
| | | L6 IT | | #8ecda0 | | | #36b44a | |
| | | L6 IT Car3 | | #ba9c66 | | | #51b949 | |
| | | L5 ET | | #d388b1 | | | #2f50a2 | |
| | | L5/6 NP | | #7b4c1e | | | #0b8281 | |
| | | L6b | | #004d45 | | | #6bbd46 | |
| | | L6 CT | | #29348c | | | #13a060 | |
| **Inh** | | SST | | #6b6a64 | | | #f37d21 | |
| #bb2028 | | SST CHODL | | #bc2025 | | | #f7901e | |
| | | PVALB | | #5066b0 | | | #f05726 | |
| | | Chandelier | | #64cce9 | | | #ec2327 | |
| | | LAMP5 LHX6 | | #ae98a1 | | | #f26f51 | |
| | | LAMP5 | | #a1b6de | | | #ec2928 | |
| | | SNCG | | #f175aa | | | #f16a23 | |
| | | VIP | | #35bba0 | | | #f9a31f | |
| | | PAX6 | | #67be62 | | | #f26f51 | |
| **Non-Neur** | | Astro | | #f5ed1f | | | #f2799c | |
| #f3eb1a | | Oligo | | #fdfded | | | #37ba88 | |
| | | OPC | | #869c98 | | | #34c1d2 | |
| | | Micro | | #92575d | | | #5fbb46 | |
| | | Endo | | #d490bf | | | #f39528 | |
| | | VLMC | | #717c33 | | | #d47eb4 | |
| | | PC | | #29471f | | | #46b1e4 | |
| | | SMC | | #413c42 | | | #af91c3 | |
| | | Immune | | #f15c5a | | | #bdb235 | |
| | | RB | | #050304 | | | #050304 | |

Related to the supplementary section "***snRNA-seq Processing*.**" This supplementary table relates to **Fig. 1** and the main text section "Constructing a single-cell genomic resource for 388 individuals."

**Table S5. List of genes identified as significant (FDR < 0.05, Wald test, overlapped across 5 cohorts) in the IT neuron trajectory analysis.**

| | | | |
|---|---|---|---|
| PENK | ANKRD62 | APOE | CCN2 |
| CDH19 | ARHGAP6 | RXFP2 | DENND2A |
| RPL26 | FAT4 | ELN | CNGA3 |
| TACR3 | PRSS12 | TFEC | PLSCR4 |
| CCN4 | TMEM132C | SLC6A1 | CAMKMT |
| SOX6 | ZFHX4 | RASGEF1B | IGSF1 |
| BCHE | CRYAB | RASSF6 | FAU |
| RPS27A | RPS23 | SLC5A8 | EEF1A1 |
| CARD18 | CA8 | DPP4 | ROR1 |
| NPNT | SKAP1 | RERG | ENPP1 |
| PTGER3 | CXCL14 | RPL19 | DOCK8 |
| RPS24 | FBXL7 | EYA4 | RUNX2 |
| VAV3 | IGFBP7 | SCN7A | RGS22 |
| ADAMTS6 | LRIG3 | SEMA6A | GPR149 |
| LONRF3 | MAF | OR3A2 | SLFN11 |
| SLC7A2 | VRK2 | HIF3A | RYR3 |
| RPL32 | TOX3 | NXPH2 | SCML4 |
| DSP | PROX1 | BTNL9 | CCDC178 |
| MYO1E | CNDP1 | ERG | EMCN |

Related to the supplementary section "***Trajectory Analysis*.**" This supplementary table relates to **Fig. 1** and the main text section "Constructing a single-cell genomic resource for 388 individuals."

**Table S6. LDSC enrichment table for the top 15 brain-related traits.**

| | Ex | cCRE | adult b-cCRE |
|---|---|---|---|
| **Time to complete round** | 65.2771 | 8.1477 | 39.0413 |
| **Duration to first press of snap-button in each round** | 63.0204 | 4.4477 | 31.6167 |
| **BipolarI** | 52.0586 | 8.6706 | 38.5841 |
| **Mean time to correctly identify matches** | 50.5290 | 3.6502 | 26.5568 |
| **Number of incorrect matches in round** | 50.4897 | 7.5351 | 21.6810 |
| **Frequency of tiredness / lethargy in last 2 weeks** | 49.2801 | 7.4888 | 32.9555 |
| **SCZ** | 41.3638 | 5.9104 | 33.8396 |
| **Fluid intelligence score** | 40.7174 | 2.0124 | 29.6504 |
| **Frequency of tenseness / restlessness in last 2 weeks** | 38.9374 | 3.1992 | 21.7368 |
| **Guilty feelings** | 38.4306 | 7.7506 | 24.1987 |
| **Duration screen displayed** | 35.4009 | 2.0756 | 15.8289 |
| **Seen doctor (GP) for nerves, anxiety, tension or depression** | 35.1478 | 9.1612 | 31.0113 |
| **MDD** | 34.5334 | 12.5635 | 34.6495 |
| **Neuroticism score** | 31.8168 | 9.5395 | 27.1935 |
| **Sleep duration** | 31.0456 | 5.6908 | 20.6848 |

LDSC enrichment against snATAC-seq peaks in Exc neurons, cCREs, and b-cCREs, with a scale of -log(p-value). The top 15 (sorted according to the enrichment in Ex) brain-related traits among UKBB, PGC, and PASS are listed. Values for all traits are available on the brainSCOPE portal.

More detail in the supplementary section "***LDSC.***" This supplementary table relates to **Fig. 2** and main text section "Determining regulatory elements for cell types from snATAC-seq."

**Table S7. scQTL counts per cell type (without LD pruning).**

**scQTL calls w/o LD pruning**

| Cell type | Sample size | # expr PCs | # Sig eGenes | Tot # sig scQTLs | # eSNPs/eGene |
|---|---|---|---|---|---|
| **L2.3.IT** | 346 | 100 | 3,049 | 368,686 | 120.9203017 |
| **Oligo** | 339 | 100 | 934 | 113,473 | 121.4914347 |
| **L5.IT** | 333 | 100 | 1,671 | 200,390 | 119.9222023 |
| **L6.IT** | 328 | 100 | 1,446 | 178,645 | 123.54426 |
| **L4.IT** | 319 | 100 | 1,220 | 149,884 | 122.8557377 |
| **Astro** | 315 | 100 | 581 | 75,616 | 130.1480207 |
| **Chandelier__Pvalb** | 313 | 100 | 943 | 122,979 | 130.4125133 |
| **OPC** | 295 | 100 | 350 | 44,567 | 127.3342857 |
| **Vip** | 294 | 100 | 591 | 78,316 | 132.5143824 |
| **Sst__Sst.Chodl** | 278 | 100 | 351 | 48,602 | 138.4672365 |
| **Lamp5** | 223 | 100 | 251 | 34,653 | 138.059761 |
| **Micro.PVM** | 207 | 100 | 43 | 7,334 | 170.5581395 |
| **L6.CT** | 202 | 100 | 157 | 24,254 | 154.4840764 |
| **L6b** | 171 | 100 | 59 | 5,067 | 85.88135593 |
| **Lamp5.Lhx6** | 163 | 100 | 5 | 139 | 27.8 |
| **L5.6.NP** | 160 | 100 | 11 | 1,128 | 102.5454545 |
| **Sncg** | 145 | 100 | 0 | 0 | - |
| **Endo__VLMC** | 128 | 100 | 0 | 0 | - |
| **L6.IT.Car3** | 123 | 100 | 0 | 0 | - |
| **PC** | 97 | 20 | 5 | 520 | 104 |
| **Immune** | 36 | 20 | 0 | 0 | - |
| **Pax6** | 36 | 20 | 0 | 0 | - |
| **L5.ET** | 28 | 20 | 0 | 0 | - |
| **SMC** | 23 | 20 | 0 | 0 | - |
| ***Avgs*** | *204.25* | | *486.125* | *60,593.875* | *~120** |

Summary statistics on the scQTL callset without LD pruning (FDR < 0.05). For each cell type (column 1), this table lists the sample size used in scQTL calling (column 2), the number of expression PCs used among the covariates (column 3), the number of significant eGenes discovered at an FDR of 0.05 (column 4), the total number of scQTLs (column 5), and the average number of eSNPs per eGene (column 6) (see supplementary section 4 for significance determination testing).

More detail in the supplementary section "***scQTLs.***" This supplementary table relates to **Fig. 4** and main text section "Determining cell-type-specific eQTLs from single-cell data."

**Table S8. Number of Bayesian scQTLs per cell type.**

| Cell Type | # Bayesian scQTLs |
|---|---|
| **Astro** | 4577 |
| **Chandelier__Pvalb** | 5047 |
| **Endo__VLMC** | 3535 |
| **Immune** | 3082 |
| **L2.3.IT** | 6017 |
| **L4.IT** | 5964 |
| **L5.6.NP** | 4574 |
| **L5.ET** | 3165 |
| **L5.IT** | 5982 |
| **L6.CT** | 5011 |
| **L6.IT** | 5982 |
| **L6.IT.Car3** | 4591 |
| **L6b** | 4830 |
| **Lamp5** | 4783 |
| **Lamp5.Lhx6** | 4393 |
| **Micro.PVM** | 3843 |
| **Oligo** | 4614 |
| **OPC** | 4558 |
| **Pax6** | 3093 |
| **PC** | 7861 |
| **SMC** | 2453 |
| **Sncg** | 4380 |
| **Sst__Sst.Chodl** | 4942 |
| **Vip** | 5130 |

Table summarizing the number of Bayesian scQTLs in each of 24 cell types (including very rare cell types). These data are also plotted within **fig. S36C**.

Related to the supplementary section "***scQTLs.***" This supplementary table relates to **Fig. 4** and main text section "Determining cell-type-specific eQTLs from single-cell data."

**Table S9. scQTL counts per cell type (with LD pruning).**

**scQTL calls w/LD pruning (~90% loss in # of SNPs in cis windows)**

| | | | w/LD pruning (~90% reduction in # SNPs) | | |
|---|---|---|---|---|---|
| **Cell type** | **Sample size** | **# expr PCs** | **# Sig eGenes** | **Tot # sig scQTLs** | **# eSNPs/eGene** |
| **L2.3.IT** | 346 | 100 | 2,732 | 18,326 | 6.707906296 |
| **Oligo** | 339 | 100 | 784 | 4,830 | 6.160714286 |
| **L5.IT** | 333 | 100 | 1,407 | 9,058 | 6.437810945 |
| **L6.IT** | 328 | 100 | 1,236 | 7,729 | 6.253236246 |
| **L4.IT** | 319 | 100 | 1,043 | 6,301 | 6.041227229 |
| **Astro** | 315 | 100 | 480 | 2,980 | 6.208333333 |
| **Chandelier__Pvalb** | 313 | 100 | 807 | 5,092 | 6.309789343 |
| **OPC** | 295 | 100 | 300 | 1,736 | 5.786666667 |
| **Vip** | 294 | 100 | 490 | 2,971 | 6.063265306 |
| **Sst__Sst.Chodl** | 278 | 100 | 287 | 1,734 | 6.041811847 |
| **Lamp5** | 223 | 100 | 198 | 1,044 | 5.272727273 |
| **Micro.PVM** | 207 | 100 | 39 | 233 | 5.974358974 |
| **L6.CT** | 202 | 100 | 136 | 677 | 4.977941176 |
| **L6b** | 171 | 100 | 41 | 162 | 3.951219512 |
| **Lamp5.Lhx6** | 163 | 100 | 4 | 7 | 1.75 |
| **L5.6.NP** | 160 | 100 | 9 | 39 | 4.333333333 |
| **Sncg** | 145 | 100 | 0 | 0 | - |
| **Endo__VLMC** | 128 | 100 | 0 | 0 | - |
| **L6.IT.Car3** | 123 | 100 | 0 | 0 | - |
| **PC** | 97 | 20 | 4 | 17 | 4.25 |
| **Immune** | 36 | 20 | 0 | 0 | - |
| **Pax6** | 36 | 20 | 0 | 0 | - |
| **L5.ET** | 28 | 20 | 0 | 0 | - |
| **SMC** | 23 | 20 | 0 | 0 | - |
| **Avgs** | 204.25 | | 416.5416667 | 2,622.333333 | ~5.4* |

Summary statistics on the scQTL callset with LD pruning ($r^2$ = 0.5; FDR < 0.05). Details on how LD pruning was performed are provided in supplementary section 4.1 (scQTLs - Cell-type-specific eQTL analysis). For each cell type (column 1), this table lists the sample size used in scQTL calling (column 2), the number of expression PCs used among the covariates (column 3), the number of eGenes discovered at an FDR of 0.05 (column 4), the total number of scQTLs (column 5), and the average number of eSNPs per eGene (column 6).

More detail in the supplementary section "***scQTLs.***" This supplementary table relates to **Fig. 4** and main text section "Determining cell-type-specific eQTLs from single-cell data."

**Table S10. Comparison of scQTLs and bulk eQTLs.**

A

Enrichment of scQTL eSNPs w/bulk eSNPs

| Cell type | # scQTL eSNPs overlap w/bulk eSNPs | Expected # scQTL eSNPs overlap w/bulk eSNPs | Observed/Expected |
|---|---|---|---|
| L2.3.IT | 135,484 | 119.6 | 1,132.81 |
| L5.IT | 76,165 | 67.3 | 1,131.70 |
| L6.IT | 67,118 | 58.8 | 1,141.50 |
| L4.IT | 58,676 | 50.5 | 1,161.90 |
| Chandelier__Pvalb | 46,968 | 41.6 | 1,129.00 |
| Oligo | 41,384 | 36.2 | 1,143.20 |
| Vip | 29,945 | 25.6 | 1,169.70 |
| Astro | 25,952 | 23.1 | 1,123.50 |
| Sst__Sst.Chodl | 19,539 | 16.7 | 1,170.00 |
| OPC | 16,157 | 13.8 | 1,170.80 |
| Lamp5 | 12,829 | 11.2 | 1,145.40 |
| L6.CT | 7,862 | 6.9 | 1,139.40 |
| Micro.PVM | 3,026 | 3.3 | 9,17.0 |
| L6b | 2,790 | 2.1 | 1,328.60 |
| L5.6.NP | 776 | 0.5 | 1,552.00 |
| PC | 94 | 0.2 | 470 |
| Lamp5.Lhx6 | 75 | 0.1 | 750 |

B

Relating of scQTLs to bulk cis-eQTLs

| Cell type | # scQTLs overlap w/bulk cis-eQTLs | # scQTLs not found in bulk | Fraction scQTLs overlap w/bulk |
|---|---|---|---|
| Lamp5.Lhx6 | 75 | 64 | 0.54 |
| L6b | 2,111 | 2956 | 0.42 |
| Sst__Sst.Chodl | 15,936 | 32665 | 0.33 |
| L4.IT | 48,704 | 101176 | 0.32 |
| L5.IT | 63,702 | 136683 | 0.32 |
| L6.IT | 56,785 | 121858 | 0.32 |
| Oligo | 35,247 | 78224 | 0.31 |
| Chandelier__Pvalb | 37,389 | 85585 | 0.3 |
| L2.3.IT | 111,961 | 256718 | 0.3 |
| Lamp5 | 10,462 | 24191 | 0.3 |
| Astro | 22,819 | 52796 | 0.3 |
| OPC | 13,394 | 31173 | 0.3 |
| Vip | 23,427 | 54887 | 0.3 |
| L6.CT | 6,998 | 17255 | 0.29 |
| Micro.PVM | 1,749 | 5584 | 0.24 |
| L5.6.NP | 215 | 913 | 0.19 |
| PC | 94 | 426 | 0.18 |

**(A)** Enrichment statistics associated with the overlap of scQTL eSNPs and bulk cis-eQTL eSNPs. **(B)** Per-cell-type summary statistics on the overlap between scQTLs and bulk cis-eQTLs.

Related to the supplementary section "***scQTLs***." This supplementary table relates to **Fig. 4** and main text section "Determining cell-type-specific eQTLs from single-cell data."

**Table S11. Pooling of snRNA-seq cell types to agree with spatial data annotations.**

| **Spatial (Validation)** | **snRNA-seq (Own)** |
|---|---|
| opc | OPC |
| excit_l4 | L4 IT |
| astro | Astro |
| inhib | (all) Inhibitory Neurons |
| endomural | Endo, SMC, PC |
| excit_l3 | |
| excit_l5_6 | L5/6 NP |
| excit_l2_3 | L2/3 IT |
| micro | Micro |
| oligo | Oligo |
| excit_l5 | L5 IT, L5 ET |
| excit_l6 | L6 IT, L6 IT Car3, L6 CT, L6b |
| excit_l3_4_5 | |
| | VLMC, Immune |

The table represents how we harmonized spatial transcriptomic and snRNA-seq cell-type annotations to perform validations of our cell-to-cell communication network.

More detail in the supplementary section "***Cell-to-Cell Network*.**" This supplementary table relates to **Fig. 6** and the main text section "Constructing a cell-to-cell communication network."

**Table S12. Correlation and significance for spatial C2C validation.**

| | **Coef** | **Pval** |
|---|---|---|
| Pearson | -0.179 | 0.06 |
| Kendall | -0.158 | 0.01 |
| Spearman | -0.241 | 0.01 |

This table represents the correlation coefficients and their significance between the spatial distances and the communication strengths across all pairs of cell types. The negative correlation values validate the spatial requirement of our communication network. Specifically, the farther apart the cell types are, the less likely they are communicating with one another.

More detail in the supplementary section "***Cell-to-Cell Network.***" This supplementary table relates to main figure 6 and main text section "Constructing a cell-to-cell communication network."

**Table S13. Model accuracy and associated common SNP heritability estimates for LNCTP models.**

| Model | PRS (*232*) | PrediXcan (*92*) | DBM(*4*) | LNCTP (w/o c2c) | LNCTP (full) | DBM (*4*) |
|---|---|---|---|---|---|---|
| Gene Expression | None | Continuous+ imputed | Binarized+ imputed | Continuous+ imputed | Continuous+ imputed | Binarized + Ground truth |
| Networks (GRN/C2C) | None | None | Bulk | Bulk + Single-cell | Bulk + Single-cell + Cell-to-cell | Bulk |
| SCZ | 56.9 (0.009) | 54.3 (0.003) | 59.0 (0.018) | 60.2 ± 0.047 (0.0287) | 60.8 ± 0.065 (0.0593) | 73.6 (0.328) |
| BPD | 57.0 (0.069) | 51.7 (0.0026) | 67.2 (0.107) | 70.6 ± 0.091 (0.161) | 72.8 ± 0.055 (0.136) | 76.7 (0.374) |
| ASD | 50.0 (0.0001) | 50.0 (0.0) | 58.8 (0.032) | 65.8 ± 0.092 (0.174) | 64.0 ± 0.086 (0.104) | 68.3 (0.113) |
| AD | 58.0 (0.013) | – | – | 69.3 ± 0.032 (0.192) | 69.5 ± 0.0203 (0.108) | – |

Table compares predictive accuracy of LNCTP with polygenic risk score (PRS), PrediXcan, and deep Boltzmann machine (DBM) models (*4*, *92*, *232*). Table shows model accuracy on a balanced test set for classification of case/control status (chance performance = 0.5). All datasets are balanced for covariates as in (*4*) and averaged across 10 data splits, as described in Supplementary Methods section 8.4. Classification accuracy is shown ± SD values for LNCTP models; additionally, associated estimates of heritability on the liability scale are quoted in brackets. Results for the DBM models are quoted directly from (*4*). SCZ = schizophrenia, BPD = bipolar disorder, ASD = autism spectrum disorder, AD = Alzheimer's disease.

More detail in the supplementary section "***LNCTP validation***." This supplementary table relates to main figure 8 and main text section "Imputing gene expression and prioritizing disease genes across cell types with an integrative model."

**Table S14. Comparison of activation functions for LNCTP (schizophrenia prediction).**

| Activation | sigmoid | ReLU | tanh | linear |
|---|---|---|---|---|
| Accuracy (SCZ) | 0.598 ± 0.036 | 0.597 ± 0.020 | 0.601 ± 0.031 | 0.602 ± 0.047 |

Table compares performances of LNCTP models using different activation functions for the MLP layers. Shown are the mean accuracy and the standard deviation across 10 balanced data splits, as described in **table S13.**

More detail in the supplementary section "***LNCTP Motivation*.**" This supplementary table relates to main figure 8 and main text section "Imputing gene expression and prioritizing disease genes across cell types with an integrative model."

**Table S15. Comparison of LNCTP (AD prediction) and other related works.**

| **Paper ID** | (*233*) | (*234*) | (*234*) | (*235*) | (*236*) | **Ours** |
|---|---|---|---|---|---|---|
| **Data Source** | Unspecified | ADNI | ANM2 (GSE63061) | IGAP dataset | ROSMAP | ROSMAP Genotypes |
| **Data Types** | Gene Expr. + DNA Methylation | Gene Expression | ` | Genotypes | Gene Expr. | Genotypes |
| **Classifiers Used** | RF | DNN | ` | CPRS | JDINAC | LNCTP |
| **AUC (Internal Validation)** | 0.683 | 0.657 | 0.804 | 0.78 | 0.84 | 0.72 |
| **AUC (External Validation)** | N/A | 0.697 (ANM1), 0.764 (ANM2) | 0.655 (ADNI), 0.859 (ANM1) | 82% in ADNI, 90% sensitivity in high-risk group | N/A | N/A |
| **Dataset Size** | 20,376 AD, 11,178 controls | 11,276 gene probes | 22,338 gene probes | 17,008 AD, 37,154 controls, 87,000 variants | 193 AD, 172 NCI, 158 genes | 366 AD, 179 control |
| **Cell-type-Specific Insights** | N/A | N/A | - | Yes - via cited papers | N/A | Yes |
| **Multi-Omics** | Dual-omics | No | - | Yes | No | Tri-omics |
| **Interpretability** | Feature Selection Methods | Feature Selection Methods, Pathway Analysis | - | N/A | Identified Hub Genes, Gene pairs | Hierarchical Linear Architecture, Feature Selection Methods, Highlighted Omics Data Contribution, etc. |
| **Multi-tasks Robustness** | N/A | N/A | - | Claimed for potential | N/A | ASD, BPD, SCZ, and AD |
| **Potential Utility in Medicine** | AD diagnosis | Early AD diagnosis | - | Candidate selection for clinical traits | Identify genes for AD | Tailored therapeutic strategies guides |

This table summarizes key information from various studies focused on using computational methods for AD research. Metrics such as AUC for internal and external validations, the dataset sizes, potential utilities in medicine are included, as well as various qualitative traits.

Related to the supplementary section "***LNCTP Training.***" This supplementary table relates to main figure 8 and main text section "Imputing gene expression and prioritizing disease genes across cell types with an integrative model."

**Table S16. Network analysis of LNCTP perturbations.**

| Disease | Perturbed Gene | Proximity Type | Correlation | p-value |
|---|---|---|---|---|
| ASD | *ANKHD1-EIF4EBP3* | Shortest Path (In) | -0.19 | 0.09 |
| BPD | *LINGO2* | Shortest Path (In) | -0.05 | 0.57 |
| SCZ | *SF3B2* | Shortest Path (In) | -0.05 | 0.27 |
| BPD | *MEF2A* | PageRank | -0.13 | 0 |
| | | Shortest Path (Out) | -0.1 | 0 |
| | | Shortest Path (In) | -0.08 | 0.15 |
| SCZ | *RORA* | PageRank | -0.18 | 0 |
| | | Shortest Path (Out) | -0.17 | 0 |
| | | Shortest Path (In) | 0 | 0.95 |
| SCZ | *TCF4* | PageRank | -0.18 | 0 |
| | | Shortest Path (Out) | -0.14 | 0 |
| | | Shortest Path (In) | -0.13 | 0.03 |

This table shows the correlation of different network proximity metrics of the perturbed genes to the magnitude of the LNCTP-imputed gene expressions in the cell-type-specific GRNs.

More detail in the supplementary section "***LNCTP Validation*.**" This supplementary table relates to main figure 8 and main text section "Imputing gene expression and prioritizing disease genes across cell types with an integrative model."

**Table S17. CLUE analysis of LNCTP perturbations.**

| Gene | Bulk | Astro | Endo | Exc | Inh | Micro | Oligo | OPC |
|---|---|---|---|---|---|---|---|---|
| *ANKHD1* | 797 | 234 | 15 | 405 | 375 | 13 | 1808 | 253 |
| *ESRRG* | 184 | 52 | 105 | 403 | 89 | 413 | 325 | 206 |
| *ID1* | 400 | 402 | 756 | 389 | 119 | 9073 | 594 | 1005 |
| *LINGO2* | 431 | 791 | 346 | 25 | 16 | 1688 | 492 | 1454 |
| *MEF2A* | 335 | 283 | 118 | 8 | 11 | 2670 | 661 | 349 |
| *RORA* | 242 | 196 | 301 | 156 | 79 | 1289 | 673 | 4605 |
| *SF3B2* | 176 | 100 | 1101 | 421 | 1014 | 4688 | 774 | 722 |
| *TCF4* | 489 | 185 | 7 | 253 | 96 | 1063 | 347 | 385 |

This table shows the number of significant results (calculated using the Computing similarities by Weighted Connectivity Score (WTCS) (*42*)) per gene per cell type (or bulk datasets).

More detail in the supplementary section "***LNCTP Validation*.**" This supplementary table relates to main figure 8 and main text section "Imputing gene expression and prioritizing disease genes across cell types with an integrative model."

**Table S18. Significance of overlaps between CRISPR differentially expressed genes (DEGs) and genes that are downstream of the 9 target TFs in (A) our weighted diffusion network and (B) a hop-distance-based network.**

(**A**) Diffusion-network-based results

| TF → | GLIS3 | NR2F2 | SOX5 | PPARGC1A | TAF1 | MEF2C | THAP1 | EGR2 | SPI1 |
|---|---|---|---|---|---|---|---|---|---|
| **DEGs in CRISPR data** | 218 | 642 | 1997 | 39 | 456 | 3 | 86 | 87 | 75 |
| **Overlaps out of 11,489 genes** | 189 | 538 | 1643 | 26 | 364 | 2 | 60 | 62 | 55 |
| **-Log$_{10}$(FDR) of TF in experiment** | 33.5 | 26.8 | 16.8 | 1.8 | 0.55 | 0.003 | 0.002 | N/A | N/A |
| Downstream Overlaps / Total (Quantile = 0.5) | 128/5745 p-value: 0.0 | 341/5745 p-value: 0.0 | 1067/5745 p-value: 0.0 | 17/5745 p-value: 0.084 | 233/5745 p-value: 0.0 | 1/5745 p-value: 0.75 | 36/5745 p-value: 0.077 | 30/5745 p-value: 0.649 | 22/5745 p-value: 0.948 |
| Downstream Overlaps / Total (Quantile = 0.6) | 110/4596 p-value: 0.0 | 272/4596 p-value: 0.0 | 924/4596 p-value: 0.0 | 15/4596 p-value: 0.052 | 196/4596 p-value: 0.0 | 0/4596 p-value: 1.0 | 28/4596 p-value: 0.177 | 29/4596 p-value: 0.168 | 15/4596 p-value: 0.983 |
| Downstream Overlaps / Total (Quantile = 0.7) | 89/3447 p-value: 0.0 | 229/3447 p-value: 0.0 | 731/3447 p-value: 0.0 | 14/3447 p-value: 0.009 | 147/3447 p-value: 0.0 | 0/3447 p-value: 1.0 | 20/3447 p-value: 0.331 | 23/3447 p-value: 0.14 | 11/3447 p-value: 0.966 |
| Downstream Overlaps / Total (Quantile = 0.8) | 59/2298 p-value: 0.0 | 151/2298 p-value: 0.0 | 471/2298 p-value: 0.0 | 9/2298 p-value: 0.059 | 108/2298 p-value: 0.0 | 0/2298 p-value: 1.0 | 13/2298 p-value: 0.424 | 18/2298 p-value: 0.057 | 9/2298 p-value: 0.798 |

(**B**) Hop-network-based results

| TF → | GLIS3 | NR2F2 | SOX5 | PPARGC1A | TAF1 | MEF2C | THAP1 | EGR2 | SPI1 |
|---|---|---|---|---|---|---|---|---|---|
| Downstream DEG Overlap / Downstream genes vs Total Overlap / Total genes | 131/6846 vs 191/11667 p-value: 0.003 | 297/5160 vs 549/11708 p-value: 0.0 | 515/2573 vs 1680/11799 p-value: 0.0 | 14/3812 vs 26/11769 p-value: 0.019 | 261/6884 vs 368/11682 p-value: 0.0 | 2/9841 vs 2/11564 p-value: 0.724 | 42/6925 vs 60/11673 p-value: 0.058 | 32/6610 vs 63/11682 p-value: 0.855 | 42/8699 vs 56/11613 p-value: 0.565 |

The nine TFs (column headings) were selected based on intersecting our GRN TFs with the TFs that were targets in the CRISPR experiments (*103*).

(**A**) For each target TF in our GRNs and in the CRISPR experiments, the diffusion scores to all genes were calculated using the weighted GRNs for the 9 excitatory cell types as layers in a framework that combines cross-layer information to get net “excitatory” diffusion scores for TF to gene pairs. The scores for each of the 9 TFs were then thresholded at quantile values ranging from 0.5 to 0.8 to determine which genes are “upstream” and which are “downstream” (see rows 4-7). We indicate how many downstream genes overlap with the DEGs out of the total number of downstream genes. Also, shown are the total numbers of DEGs in the CRISPR experiments; the numbers of those DEGs that are found in our 11,489 GRN genes; and the -Log$_{10}$(FDR) value for the DEG effect size (log$_2$ Fold-change) for the target TF in the CRISPR experiment. The columns are ordered based on decreasing values for the $-\text{Log}_{10}$(FDR), which indicates the degree to which the intended CRISPR effect (either activation or interference) was observed. (**B**) We used the hop-distance network generated by applying the *igraph* program on a combined excitatory neuron GRN, which in turn was created as the union of all the GRN connections in the excitatory subtypes: “downstream” and “upstream” directions are determined by whether a target gene is reachable from a TF by following the directed connections in the GRN (downstream) or if the TF is reachable from the gene (upstream); the distance between a TF and up-/down-stream genes are found as the total number of steps needed within the GRN to reach the target (a gene in the downstream case, the TF in the upstream case). We chose to consider all genes within a hop-distance of ≤ 2 as downstream, and pooled all the downstream genes with hop-distance > 2 and all upstream genes together as “upstream”. This is because the high interconnectedness of the network meant that most genes were labeled as downstream in *igraph*. Since the goal is to observe whether more proximal downstream genes show an enrichment in CRISPR DEGs, we chose a hop-distance cutoff of 2 as reasonable.

The numbers of overlaps between the DEGs and the diffusion-based downstream genes in our network, versus those in the total upstream+downstream list are then tested for significance using the one-sided Fisher’s exact test (testing for a greater effect size in downstream genes). We color-coded results as: p-values < 0.05 in red; p-values < 0.1 in blue; the remaining in black.

More detail in the supplementary section "***LNCTP Validation*.**" This supplementary table relates to main figure 8 and main text section "Imputing gene expression and prioritizing disease genes across cell types with an integrative model."

### Supplementary Data Files

**Data files related to Fig. 1 and main text section "Constructing a single-cell genomic resource for 388 individuals":**

**Data S1**. **Clinical and demographic metadata, cohort, and data modalities for all samples.**

**Data S2**. **Mapping of uniform IDs for each sample across sub-cohorts and data modalities.**
This file includes mapping between different IDs across snRNA-Seq, snATAC-Seq, and genotype datasets.

**Data S3**. **brainSCOPE input datasets and output resources generated for main and supplemental figures.**

**Data S4**. **Normalized cell counts and fractions per individual from snRNA-Seq data.**
Columns indicate cell type, cell counts, cell fraction, and relevant meta-data for each individual.

**Data S5**. **Normalized cell fractions per individual from deconvolved bulk RNA-Seq data.**

**Data S6**. **Correlations between deconvolved and single-cell derived cell type fractions.**

**Data S7**. **Cell-type-specific DE genes for ASD, schizophrenia, and bipolar disorder.**
This file lists significant (p<0.05, DESeq2 likelihood ratio test)DE genes in each comparison (across all cell types for individuals with a particular disorder vs. control individuals); full results are available on the brainSCOPE portal.

**Data S8**. **Gene ontology functions of genes identified in pseudotime analysis.**

**Data files related to Fig. 2 and main text section "Determining regulatory elements for cell types from snATAC-seq":**

**Data S9**. **LDSC enrichment of UKBiobank GWAS traits for cCREs, b-cCREs, and scCREs.**
The file is indexed by trait ID and includes -log(p-value) from the LDSC test. The column 'UK Biobank trait' refers to the trait name/description in UKBB. The column 'HPO phenotype category' refers to the phenotype ontology category. The column 'brain' refers to whether the trait is brain-related. File .

**Data S10**. **LDSC enrichment of PGC and PASS GWAS for cCREs, b-cCREs, and scCREs.**
The file is indexed by trait ID and includes -log(p-value) from the LDSC test. The column 'brain' refers to whether the trait is brain-related.

**Data S11**. **Enrichment Z-scores of TF binding motifs in distal and proximal scCREs**.

**Data files related to Fig. 3 and main text section "Measuring transcriptome and epigenome variation across the cohort at the single-cell level":**

**Data S12**. **Total expression variation and variation from sample and cell type from VariancePartition.**
Note that this includes ~13k genes that meet the minimum QC requirements.

**Data S13**. **Total expression variation and variation from sample, cell type, and brain region from VariancePartition.**

**Data S14**. **Mapping between genes and gene families for variation analysis**.

**Data files related to Fig. 4 and main text section "Determining cell-type-specific eQTLs from single-cell data":**

**Data S15**. **Single-cell eGenes for 17 cell types from the core scQTL analysis.**
This gene set was used for QTL-related functional analyses. The full set of scQTLs and eSNPs are available on the brainSCOPE portal.

**Data S16**. **Single-cell isoSNPs for 22 cell types from the isoQTL analysis.**
isoSNPs were filtered for isoGene-specific nominal p-value thresholds (permuted beta distribution-derived $p<0.05$ filter).

**Data S17**. **Annotation of core scQTL eGenes for brain-related diseases and traits.**
Diseases and traits include ASD, schizophrenia, bipolar disorder, and Alzheimer's disease/aging. "X" annotations in each of the four disease columns indicate if an eGene is associated with a disease.

**Data S18**. **Dynamic eQTLs identified with the PME model for SNP terms only.**

**Data S19**. **Dynamic eQTLs identified with the PME model for SNP and interaction terms.**

**Data files related to Fig. 5 and main text section "Building a gene regulatory network for each cell type":**

**Data S20**. **SCENIC-derived scores for regulons in 24 cell types**.
Scores for regulons (TF and all target genes) were used as inputs for constructing the final GRNs.

**Data S21**. **Overlap of scQTLs with enhancer and promoter elements in GRNs.**

**Data S22**. **Validation of GRN-predicted enhancers in targeted CRISPR knockout experiments**.

This file details expression of the target genes before and after CRISPR knockout of the linked enhancers, as predicted by peak2gene linkages.

**Data S23**. **Cell-type-specific "in-hub", “out-hub”, and “bottleneck” genes in GRNs.**
The matrix lists whether a TF was identified as an in-hub, out-hub, or bottleneck (1) or not (0) in each cell type.

**Data S24**. **Gene ontology enrichment for bottleneck genes in cell-type-specific GRNs.**
Columns represent gene ontology terms, p-value, FDR, signature, gene set count, overlap count, background, count cell type, TF, and enrichment score.

**Data files related to Fig. 6 and main text section "Constructing a cell-to-cell communication network”:**

**Data S25**. **Ligand-receptor signaling patterns across cell types for control, schizophrenia, and bipolar disorder**.
File lists all interactions between ligand-receptors in different cell types, along with the strength of interaction and annotations for interaction type and pathway.

**Data S26. Signaling pathway patterns across cell types for control, schizophrenia, and bipolar disorder.**
These files contain sets of ligand-receptor signaling patterns across cell types, summarized by signaling pathway. File lists all signaling pathway interactions in different cell types, along with the strength of interaction and annotations for interaction type.

**Data files related to Fig. 7 and main text section "Constructing a cell-to-cell communication network”:**

**Data S27. Correlation and linear model associations between cell-type fraction and age**.
File lists correlations and p-values/FDRs from GLMs (with age, biological sex, and genotype ancestry covariates) comparing age and cell-type fractions from bulk RNA-Seq deconvolution (table A) and scRNA-Seq (table B).

**Data S28**. **Comparison of cell-type-specific DE genes in aging and AD**.
File contains an inner join of cell-type-specific aging DE genes (identified in this study) and AD DE genes from (*115*).

**Data S29**. **Performance of AD model predictions by cell type and data modality**.
File contains AUPRC values for cell types and data modalities (rf.meth=methylation, rf.expr=expression) from the AD model predictions.

**Data files related to Fig. 8 and main text section "Imputing gene expression and prioritizing disease genes across cell types with an integrative model":**

**Data S30**. **Salience and coheritability estimates for LNCTP prioritized genes by disorder.**
File contains gene lists with salience and coheritability values for schizophrenia, bipolar disorder, ASD, and AD (p-values based on Pearson Correlation).

**Data S31**. **Salience and coheritability estimates for prioritized cell types by disorder.**
P-values in the file are based on Pearson Correlation.

**Data S32**. **Salience and coheritability estimates for prioritized cell-to-cell interactions by disorder.**
P-values in the file are based on Pearson Correlation.

**Data S33**. **Prior literature and GWAS support for LNCTP prioritized genes.**
This file contains citations for literature supporting LNCTP gene prioritization results as follows: (*100*, *237–249*)
